# Computational discovery of hidden breaks in 28S ribosomal RNAs across eukaryotes and consequences for RNA Integrity Numbers

**DOI:** 10.1101/773226

**Authors:** Paschalis Natsidis, Philipp H. Schiffer, Irepan Salvador-Martínez, Maximilian J. Telford

**Affiliations:** Centre for Life’s Origins and Evolution, Department of Genetics, Evolution and Environment, University College London, Gower Street, London WC1E 6BT, UK

## Abstract

In some eukaryotes, a ‘hidden break’ has been described in which the 28S ribosomal RNA molecule is cleaved into two subparts. The break is common in protostome animals (arthropods, molluscs, annelids etc.) but a break has also been reported in some vertebrates and non-metazoan eukaryotes. We present a new computational approach to determine the presence of the hidden break in 28S rRNAs using mapping of RNA-Seq data. We find a homologous break is present across protostomes although has been lost in a small number of taxa. We show that rare breaks in vertebrate 28S rRNAs are not homologous to the protostome break. A break is found in just 4 out of 331 species of non-animal eukaryotes studied and three of these are located in the same position as the protostome break suggesting a striking instance of convergent evolution. RNA Integrity Numbers (RIN) rely on intact 28s rRNA and will be consistently underestimated in the great majority of animal species with a break.

## INTRODUCTION

Ribosomes are made up of up to 80 ribosomal proteins and three (in prokaryotes) or four (in eukaryotes) structural ribosomal RNAs named according to their sizes: in eukaryotes these are the 5S (~120 nucleotides), the 5.8S (~150 nucleotides), the 18S (~1800 nucleotides) and the 28S (~4000 to 5000 nucleotides). The 5.8S, 18S and 28S rRNAs are initially transcribed as a single RNA operon (the 5S is at a separate locus in eukaryotes). The 18S and 5.8S are separated by the Internal Transcribed Spacer 1 (ITS1) and 5.8S and 28S are separated by ITS2. The initial transcript is cleaved into three functional RNAs by removing ITS1 and ITS2 (Fig. 1A).

**Figure 1.**
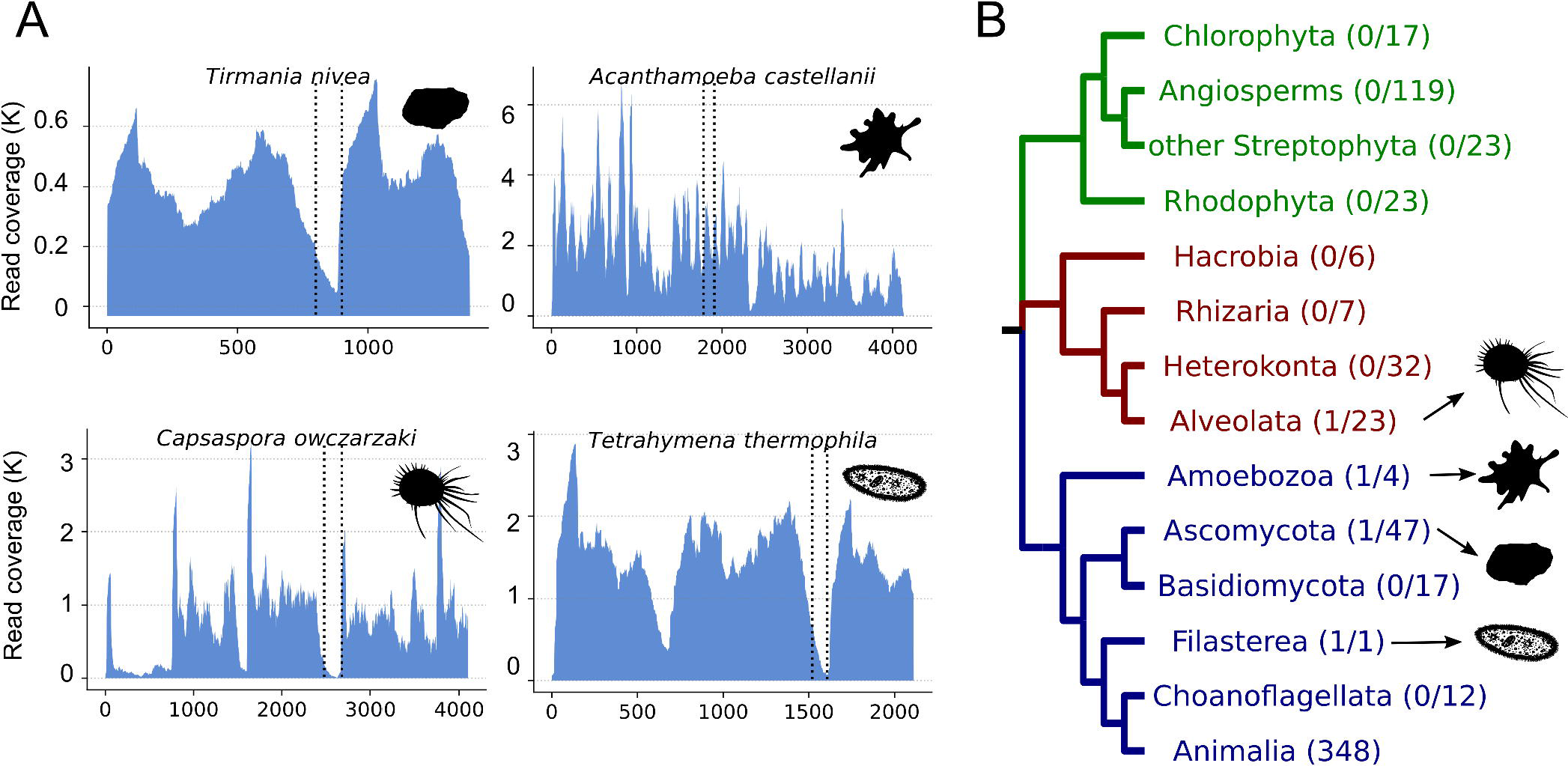
The hidden break and how to diagnose it. A) After post-transcriptional processing, the eukaryotic pre-rRNA molecule is cleaved to produce the 18S, 5.8S and 28S subunits. The RIN works normally in species that follow this rule. B) In some species, the 28S subunit gets cleaved into 28Sα and 28Sβ, a phenomenon known as the hidden break. This phenomenon can be detected via electrophoretic graphs where, in species with break, only one peak, instead of two, is observed. C,D) Our method allows the computational discovery of hidden breaks, by mapping RNA-Seq reads onto the 28S rRNA sequence and measuring i) the read coverage and ii) the ratio of forward/reverse reads mapped in each position of the 28S. In species with a break, e.g. *Bombyx mori*, a drop in coverage is observed in the middle of the break region. Additionally, the majority of reads mapped before and after the break region are reverse and forward reads respectively. These features are not observed in *Aporometra wilsoni*, an animal that does not possess the break. The vertical lines represent the conserved regions flanking the hidden break site.

In some species this picture is complicated by observations of a ‘hidden break’ in the 28S ribosomal RNA. In organisms with a hidden break, the 28S rRNA molecule itself is cleaved into two approximately equal sized molecules of ~2000 nucleotides each (Fig. 1C) [1] These two RNAs (the 5’ 28Sα and the 3’ 28Sβ) are nevertheless intimately linked by inter-molecular hydrogen bonding within the large subunit of the ribosome, just as the 28S and 5.8S rRNAs, as well as different regions of the intact 28S rRNA are in species without a break.

The hidden break, first described in pupae of the silkmoth *Hyalophora cecropia* [2], manifests itself experimentally when total RNA is extracted and separated according to size using gel electrophoresis. If the RNA is first denatured by heating in order to separate hydrogen bonded stretches of RNAs before electrophoresis, the hydrogen bonded 28Sα and 28Sβ subunits separate. These two molecules, being approximately the same size, then migrate on a gel at the same rate as each other and, coincidentally, at the same rate as the almost identically sized 18S rRNA molecule. The effect is that, rather than observing two distinct ribosomal RNA bands on the gel of ~2000 and ~4000 nucleotides, a single (and more intense) band composed of three different molecules each approximately 2kb long (18S, 28Sα and 28Sβ) is seen [3]. This also applies to electropherograms where we normally expect 2 distinct peaks for 18S and 28S molecules (Fig. 1A), but in species that possess the hidden break only one peak is observed corresponding to the equally sized 18S, 28Sα and 28Sβ (Fig. 1A & 1C).

This behaviour of rRNA molecules in species with a hidden break can have interesting and potentially detrimental practical consequences. The integrity of 18S and 28S rRNA molecules, as evidenced by the presence of 2kb and 4-5kb bands/peaks following electrophoresis, is widely used as an indication that a total RNA sample is not degraded. This is an important quality control step for experiments that will use the total RNA sample for subsequent experiments such as transcriptome sequencing, Northern blotting or rtPCR. The requirement to observe two rRNA peaks in an intact RNA sample has been formalised as an important part of the RNA Integrity Number (RIN, [4]). For RNA samples extracted from species possessing the hidden break, if conditions allow the separation of the two 28S subunits, a single rRNA band/peak will be observed. The absence of two distinct rRNA peaks will give the impression of a degraded sample with a low RIN, even when the RNA is not degraded.

Since its description in the silkmoth, the 28S hidden break has been found in numerous other species, although it is by no means universal. The most significant systematic attempts to catalogue the hidden break using gel electrophoresis date to the 1970s when Ishikawa published a series of papers (summarised in [5]). Ishikawa observed the break in several arthropods and one or two representatives of other protostome phyla (Annelida, Mollusca, Rotifera, Platyhelminthes, Phoronida/Brachiopoda and Ectoprocta). The hidden break was not found in members of Deuterostomia (Chordata and Echinodermata) ([6] but was also absent from two species of nematodes [7] and from a species of chaetognath [8] (then thought to be deuterostome relatives but now known to be protostomes [9]). The hidden break was recorded as ambiguous in non-bilaterian Cnidaria and absent in Porifera [8, 10]. The essence of Ishikawa’s work was to show that the hidden break was a characteristic of the Protostomia although, surprisingly, a hidden break was also described in several unicellular eukaryotes: *Euglena*, *Acanthamoeba* and *Tetrahymena* [8] as well as in plant chloroplasts [11]. The precise location of the hidden break is not revealed by these studies using electrophoresis.

Despite Ishikawa’s work and other sporadic publications describing the presence of the hidden break in different taxa [1, 3, 12–15], it is still currently unclear which groups of organisms possess the hidden break, whether the feature is homologous in those species that have a split 28S and has been lost in those that lack it, or whether a hidden break has evolved more than once.

Here we describe a new computational method to diagnose the presence of a hidden break in the many taxa for which large-scale RNA sequence data are available. Rather than requiring fresh material to be available for extracting total RNA to be run on an electrophoresis gel, our method is predicated on the expectation that, in taxa with a break, few if any RNA-Seq reads will map to the region that becomes excised from the initial 28S rRNA when this is broken to form the 28Sα and 28Sβ (Fig. 1C). Similarly, in the region immediately before the break we expect a large proportion of the mapped reads to be reverse reads (red in Fig. 1C), while the majority of the reads that mapped immediately after the break will be forward reads (green in Fig. 1C). Lastly, when paired-end sequences are available, we expect there to be few if any pairs of reads spanning the region of the break, as the two sides of the break are on separate molecules.

We have tested our method by detecting known instances of the hidden break and have expanded our investigation to selected members of all metazoan phyla for which suitable RNA-Seq data currently exist. We also use our method to examine reported instances of the hidden break outside of the protostomes to establish whether there is evidence that these patchily distributed observations reveal hidden breaks that are homologous to the break commonly observed in the protostome animals. Finally, we have expanded our search to a diversity of eukaryotes: to verify the described instances of a break; to determine where in the molecule this exists; and to search for novel instances of a hidden break.

## RESULTS AND DISCUSSION

### Testing the method using known examples with and without the break

To date, the presence of the hidden break has been established experimentally using electrophoresis of total RNA [3]. A comparison of untreated and heat denatured total RNA shows the conversion of the large 28S band into two smaller bands running coincident with the 18S band: in effect the 28S band disappears [3]. Our computational method relies, instead, on the expectation that RNA sequencing reads derived from RNA extracted from organisms possessing a 28S rRNA molecule that has been split by a deletion will show two characteristics. First, there will be an obvious absence of reads mapped to the genomic rRNA location precisely at the position of the split; second, we expect that reads mapped right before the break will mostly be reverse reads, in contrast with the region after the break where most mapped reads will be forward reads; third, if paired-end sequencing data are available, there should be no read pairs spanning from one side of the split to the other as these are separate molecules.

The position of the split has been mapped experimentally in very few species, including the silkworm *Bombyx mori* [16] (Fig 1B). Our method makes the accurate mapping of the position of the split a simple process which is of importance: if we are to be confident that this character is homologous between species it is necessary to show that the break occurs in a homologous region of the molecule. The procedure for mapping reads and counting spanning reads are described in Materials and Methods. Figure 1B shows an example of the pattern of read depth and forward/reverse reads ratios that we observe in an organism without a split (*Aporometra wilsoni*) and in an organism in which the split is known to exist (*Bombyx mori*) [16]. We also indicate the position of the break, which is known experimentally in *Bombyx mori*. This experimentally mapped break is surrounded by highly conserved stretches of nucleotides allowing us to identify the homologous region of the gene across eukaryotes.

### Expanding to most protostomian phyla

The hidden break has been characterised in a number of animal species and has largely been considered to be specific to the protostomes. We have used our method to expand the search for a potential hidden break to members of all animal phyla for which we have found suitable RNA-Seq data. In total we have examined 347 metazoan species including members of all but 4 animal phyla (Onychophora, Loricifera, Micrognathozoa and Gnathostomulida). The majority of the species analysed came from the Arthropoda (169) and Mollusca (31). We have also analysed 31 chordates, 28 echinoderms, as well as 2 hemichordates and 7 xenacoelomorphs. We have also examined data from 36 non-bilaterian species (29 cnidarians species, 3 ctenophores, 1 placozoan and 3 poriferans).

For the most part our results confirm those of Ishikawa; in almost all protostomes providing unambiguous results (17 out of the 19 protostomian phyla we were able to test) we observe the existence of the hidden break. Representative examples are shown in Fig. 2A. In each observation of the break we find it is bounded by the same conserved sequence regions of the 28S molecule. Our ability to map the break means we have been able to show that, across the protostomes, it is present in the same position in the molecule and we conclude that this is a homologous character throughout this major group of animals. This is the first time that this finding is supported by such a large taxon sampling and across such a breadth of taxa.

**Figure 2.**
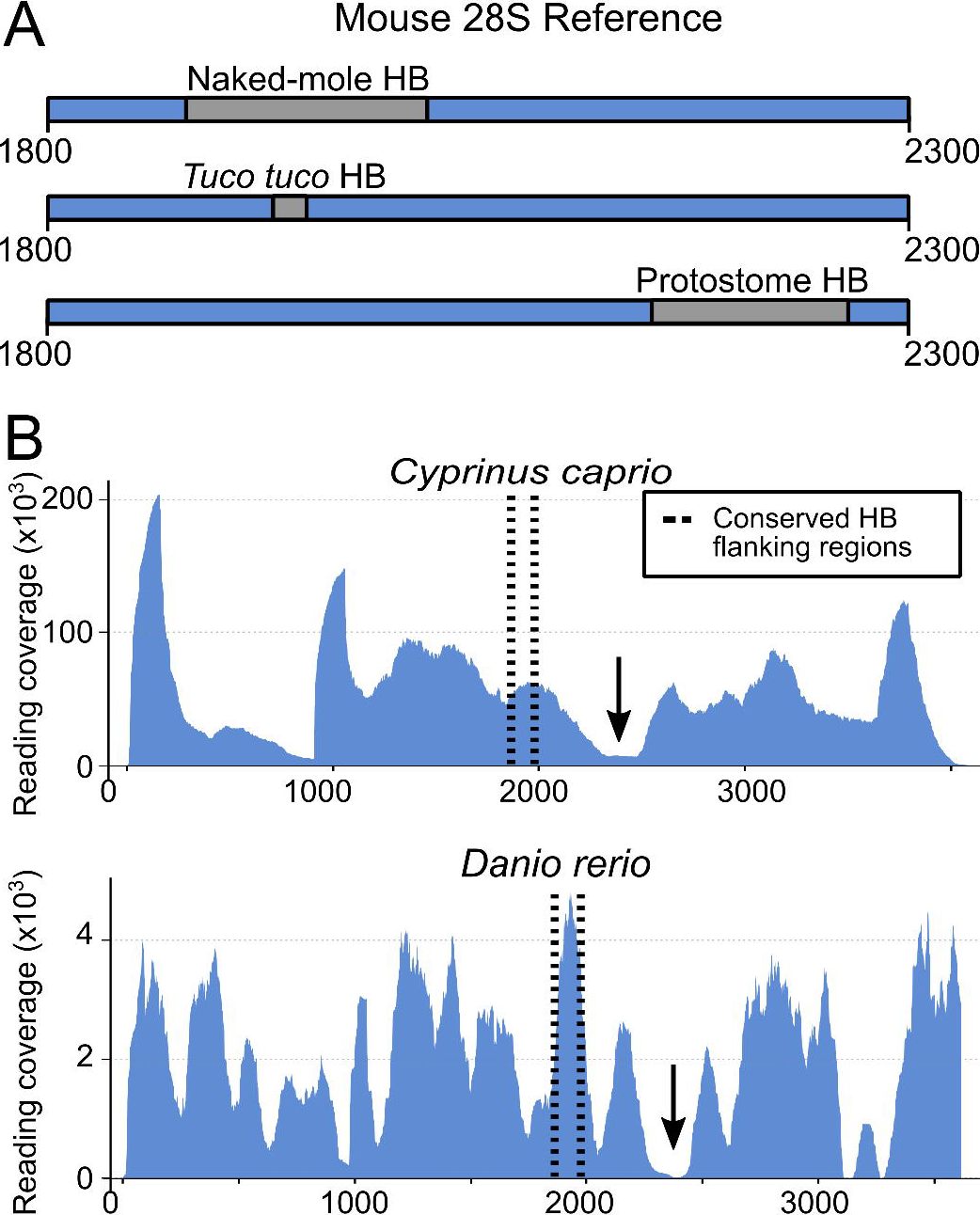
The hidden break is found in the protostome animals. A) Six examples of the application our method to different metazoan species. Evidence for a hidden break was found in most molluscs and annelids, including the orthonectid *Intoshia linei*. The fast-evolving rhombozoan *Dicyema japonica* is one of the few examples where the break was secondarily lost. The Nematoda also appear to be a special case, with the break present in the enoplean species, e.g. *Rmanomermis culicivorax*, but not in the Chromadorea, e.g. *Dictyoaulus viviparus* or the model *Caenorhabditis elegans* (not shown).B) The phylogenetic distribution of hidden break across metazoan phyla shows that it is a protostome synapomorphy, with occasional secondary losses.

Unlike previous findings [8], we have found evidence for the hidden break in a chaetognath; at the time of Ishikawa’s work members of this phylum were thought to be relatives of the deuterostomes but are now known to be protostomes [9]. Also noteworthy is the presence of this protostomian character in the orthonectid *Intoshia linei* although not in the dicyemid *Dicyema japonica*. While these two highly simplified animals were initially classified together within the phylum Mesozoa as a group intermediate between Protozoa and Metazoa, the dicyemids and orthonectids are now known to be separate lineages and both to be taxa within the Protostomia Lophotrochozoa [27]. The presence of the hidden break in *Intoshia* fits with this protostomian affiliation and the break has presumably been lost in *Dicyema*.

We also find evidence that the hidden break is absent in a number of other protostomian clades (Fig 2). We confirm, for example, Ishikawa’s work showing that there is no break in certain nematodes [7], or in the aphid *Acyrthosiphon pisum* [28]. We are also able to show for the first time that there is no break in the 28S rRNA of the priapulid *Priapulus caudatus* or in 2 species of tardigrades. Interestingly, in most of these cases, expanding our sampling to relatives of species without the break uncovers taxa that possess a hidden break. We find the break in the enoplean nematodes *Romanomermis* and *Trichinella* and also in the sister group of the nematodes - the Nematomorpha. The same is true of priapulids as we find the break in a second priapulid - *Halicryptus spinulosus*. While the pea aphid lacks a break [28], we have shown the break is present in 157 other species of arthropod. The break is also known to be absent in *Thrips tabaci* [29] yet we find evidence for a hidden break in four other closely related members of the thrips.

We find that almost every protostome species where we demonstrate the absence of the hidden break has relatives possessing it, showing that the lack of a break is a derived character - the break existed in an ancestor and has been lost. The exceptions to this rule are the tardigrades and dicyemids for which we currently have limited data. While absence of a break is infrequent in our sample of protostomes, losses have nevertheless occurred repeatedly suggesting that loss is easily achieved and relatively easily tolerated.

### Non Protostomian Metazoans

We next applied our method to the two clades of deuterostomes: the Xenambulacraria [30] (29 echinoderms, 2 hemichordates and 7 xenacoelomorphs) and the Chordata (2 urochordates and 29 vertebrates). We find that the break found in protostomes does not exist in any of the sampled species. There are, nevertheless, two special case previously described in vertebrates: hystricomorph rodents, where species of the genus *Ctenomys* (the south american Tuctuc) and the naked mole-rat (*Heterocephalus glaber*) are found to possess the hidden break [31, 32]. We show, however, that the break found in these two rodents is not homologous to the protostome hidden break. We were unable to retrieve 28S sequences for these two species, however, by using the mouse 28S rRNA sequence as reference, we managed to locate these two breaks, as well as the two conserved markers that enclose the protostome break. As it can be seen in fig. 3A, the two recorded rodent breaks do not fall between these markers.

**Figure 3.**
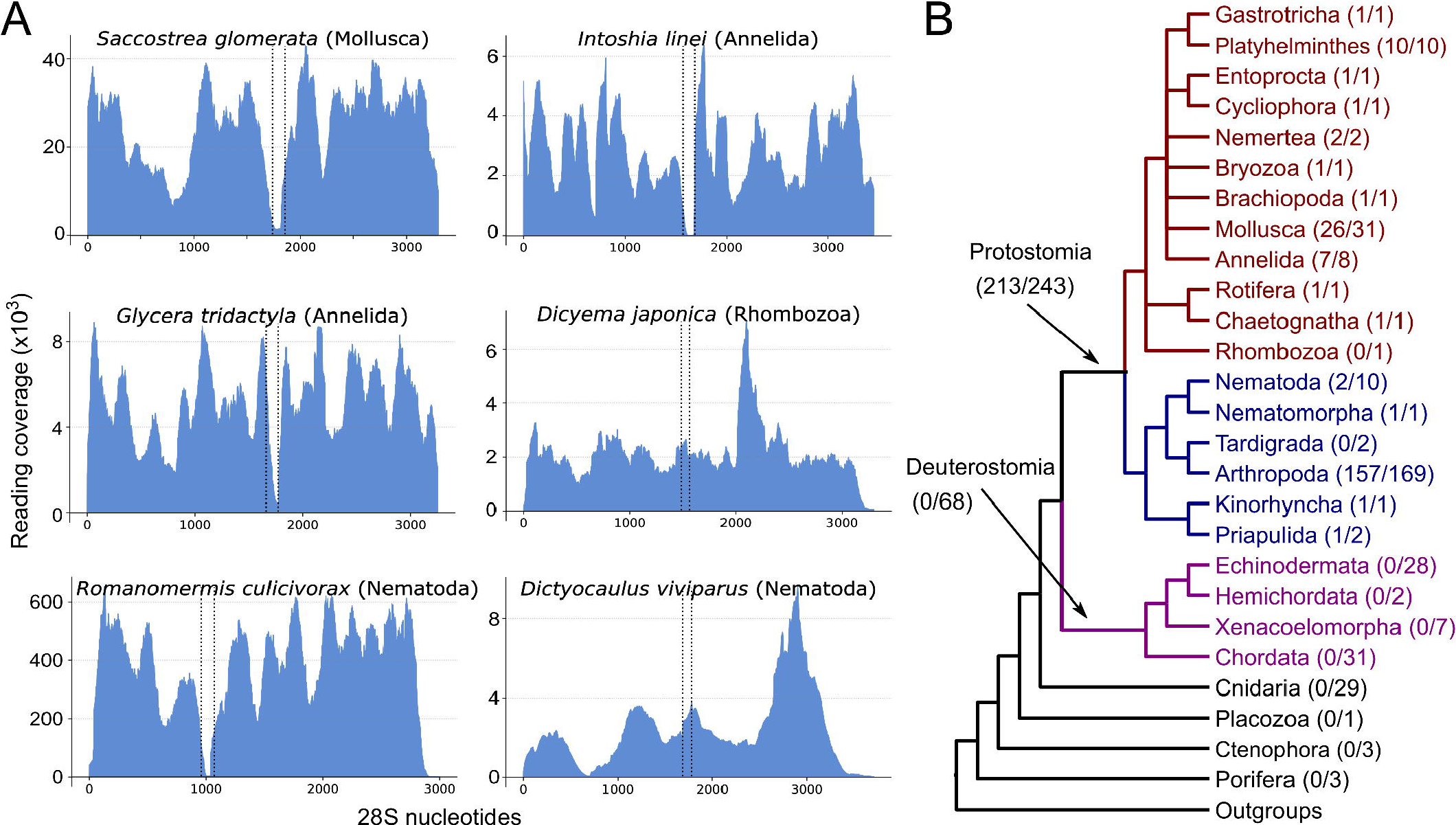
Vertebrate hidden breaks are not homologous to protostome breaks. A) Using the mouse 28S rRNA sequence as a reference, we show that the break regions recorded in rRNA extracted from naked mole-rat and Tuco-tuco testes are not homologous to the protostome hidden break. In both species the break is 5’ of the two conserved markers we found surrounding the break in protostomes. B) The break recorded in cyprinid fish is not homologous to the protostome break; here we found the break is (black arrows) 3’ of the two conserved markers we found surrounding the break in protostomes.

In more detail, the *Ctenomys* hidden break corresponds to nucleotides 1930-1950 in mouse 28S, while the naked mole-rat hidden break falls between nucleotides 1880-2020 of the mouse 28S. The two conserved satellites of the protostome hidden break lie in the positions 2149 and 2264 of the same mouse 28S sequence. The rodent hidden break site is located nearly 300bp upstream of the conserved protostome site. The hidden break has also been reported in cyprinid fish [33] and we have analysed two members of this clade. Our results show that the break in cyprinids is also in a different location that of protostomes and does not match the break described in rodents (Fig 3B). We next looked at non-bilaterian animals to determine whether they possess the hidden break. Ishikawa’s work had been inconclusive in this regard - at least in the Cnidaria [10] - leaving open the possibility that the break found across Protostomia is a primitive character that had been lost in the deuterostomes. In a total of 36 non-bilaterian species (3 sponges, 29 cnidarians, 3 ctenophores, and the placozoan *Trichoplax adhaerens*) we find no evidence for the existence of a hidden break (Fig 2B). The distribution of the break in the animal kingdom points to the hidden break having appeared in the lineage leading to the common ancestor of the protostomes.

#### Convergent evolution of the hidden break outside Metazoa?

Alongside the animals, there have been reports of a hidden 28S rRNA break in a small number of distantly related species of eukaryotes, the ciliate *Tetrahymena* [34], the amoebozoan *Acanthamoeba* [35] and the desert truffle *Tirmania nivea* [36]. We have investigated these taxa using our pipeline (Fig. 4A) to define the position of their breaks. We have greatly extended our analyses of non-animal eukaryotes to over 300 non-metazoans, including 119 plants, 64 fungi and 148 other taxa (Fig. 4B). The hidden break previously described in plant chloroplast 23S rRNAs (as opposed to nuclear 28S rRNA) does not appear to be located in the same region of the molecule although the evolutionary distance makes alignment difficult [11].

**Figure 4.**
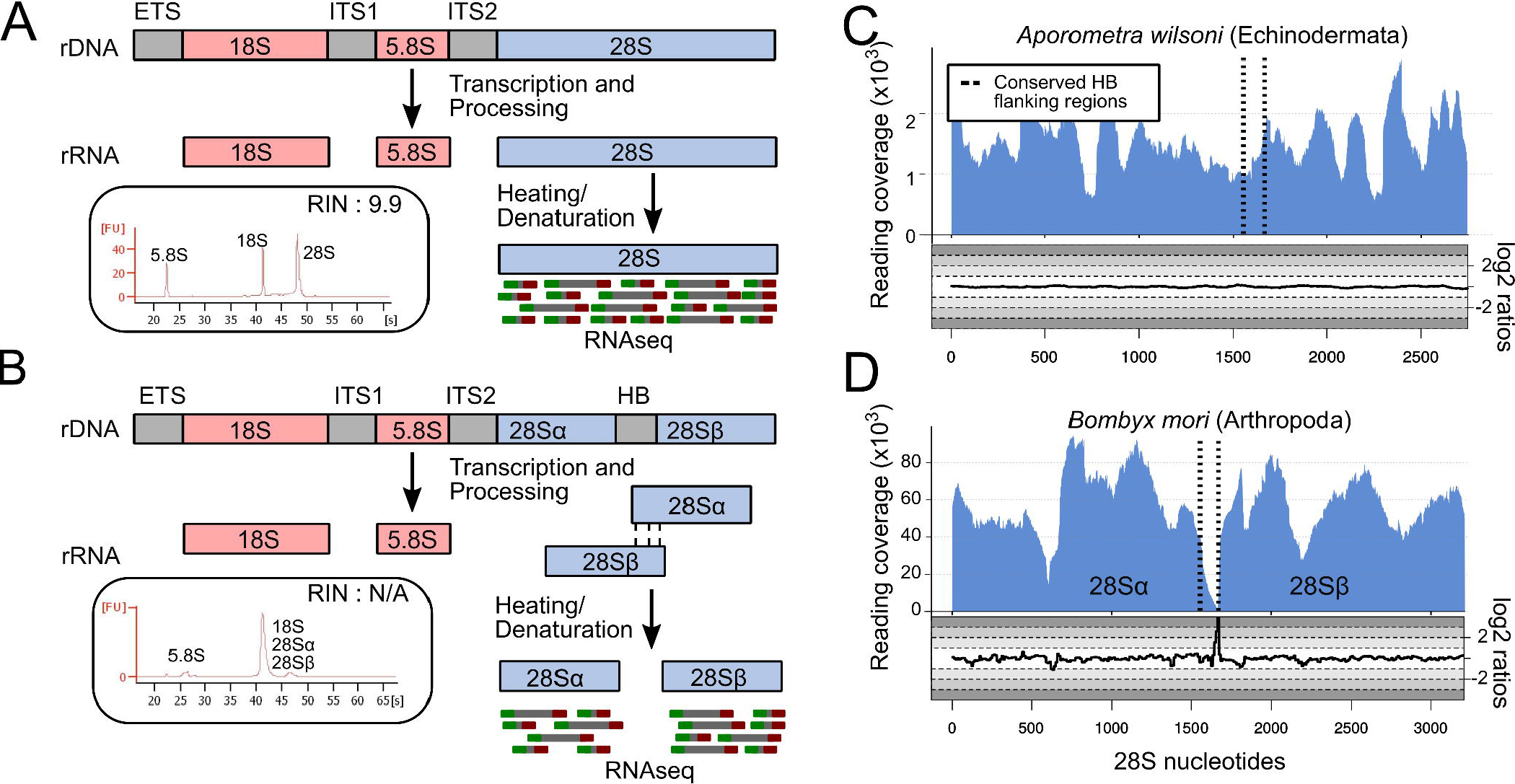
Distribution of the hidden break across non-metazoan eukaryotes. A) Four non-metazoan eukaryotes show evidence of a break. In three of these, (*Tirmania nivea*, *Capsaspora owczarzaki*, *Tetrahymena thermophila*) our analysis shows a hidden break that is in a homologous position to the protostome one. In *Acanthamoeba castellanii*, the drop in read coverage is observed 3’ of the two conserved markers we found surrounding the break in protostomes. This occurrence of a hidden break has previously been characterised experimentally (see main text). B) The phylogenetic distribution of hidden break across Eukaryota suggests that the trait evolved convergently in the protostome ancestor and in a few taxa outside the Metazoa.

In *Acanthamoeba*, gel electrophoresis of heat denatured LSU rRNA showed that it separates into fragments corresponding to molecular weights of 0.88 × 10^6 and 0.60 × 10^6 daltons [35]. Our results provide some support for this, as we find a break in the molecule that would divide it into fragments of 2320 and 1890 nucleotides (a ratio of 0.88:0.71). As suggested by the unequal sizes, the break is in a different position to the protostome 28S break. Surprisingly, the same is not true of the other two described instances of non-animal hidden breaks. We find a break as predicted in both *Tetrahymena* [34] and the desert truffle *Tirmania* [36] and in both cases the break is positioned in the same region of the 28S rRNA as the protostome hidden break. Our analysis of over 300 other eukaryotes (albeit heavily biased towards plants and fungi) found only one other clear instance of a break; this was in the unicellular opisthokont *Capsaspora owczarzaki* and, as we observed in *Tetrahymena* and *Tirmania*, the break was in the same position as the protostomian break.

## Conclusions

We present a computational approach that enabled us to perform a thorough and taxonomically broad examination of the 28S rRNA molecules of the animals. Our results strongly support Ishikawa’s observation of a hidden break that evolved in the common ancestor of the protostomes. We have searched for the hidden break in members of almost all protostomian phyla and have demonstrated its existence in a homologous region of the molecule in almost all cases - the exceptions being the tardigrades and dicyemids/rhombozoans. By expanding beyond the published observations of occasional absences of the break in protostomes we show that in almost all cases of a species lacking the break, we find its existence in sister taxa, implying that the absence is a derived state rather than a primitive absence. We interpret the observed lack of the hidden break in tardigrades and dicyemids as due to loss of the character in these lineages (at least in those few we have been able to sample).

We have also examined other instances of a break previously recorded in non-protostome animals (in two groups of rodents and cyprinid fish) and have been able to show in each case that, while experiments have suggested that they may share a mechanism with the introduction of protostomian break, they occur in non-homologous regions of the molecule and this, together with their phylogenetic distribution, shows that they are convergently evolved instances of a 28S break. The large evolutionary distance between the non-metazoan eukaryotes in which we have confirmed or discovered 28S rRNA hidden breaks as well as the lack of a break in most sampled eukaryotic taxa suggests these too are rare cases of convergent evolution. This conclusion makes the fact that the three, presumably convergently evolved breaks known in *Tetrahymena*, *Tirmania* and *Capsaspora* all fall within the same region of the molecule as the break in protostomes particularly notable and strongly suggestive both of a common mechanism and, probably, a common functional reasons for the evolution of these breaks. However, what the function of the break might be is still unknown. Our findings have important ramifications for the use of RNA Integrity Numbers or RIN [4]. The RIN, relying as it does on evidence for the integrity of the distinct 28S rRNA molecule, will tend to produce artificially low values for the great majority of protostomes in cases where denaturation of the RNA is possible. This source of error means that experimenters need to be careful in interpreting RIN when evaluating RNA samples from the more than 95% of animal species that are protostomes [37]. The scale of this potential problem suggests that an alternative to the standard RIN that takes into account the protostomian hidden break would be a valuable development for many researchers.

The emergence of a break between two regions of the large subunit ribosomal RNA is already known to have occurred previously in eukaryotic evolution. The 5.8S rRNA, in addition to the 28S rRNA, forms part of the large subunit of the ribosome [38]. As discussed, the eukaryotic 5.8S and 28S rRNAs are initially transcribed as a single molecule and are subsequently separated by the excision of the intervening Internal Transcribed Spacer (ITS2). A separate 5.8S rRNA does not, however, exist in bacteria where a sequence homologous to the eukaryotic 5.8S forms an uninterrupted part of the 23S rRNA (the homolog of the eukaryote 28S) [38]. The separation of 5.8S and 28S and the evolution of the ITS2 seem to be close counterparts of the protostomian separation of 28Sα and 28Sβ. We propose that the rapidly evolving, excised spacer sequence that lies between 28Sα and 28Sβ. should be considered as a third, protostome-specific, Internal Transcribed Spacer - the ITS3.

## METHODS

### General method

Our method to identify the hidden break rests on the assumption that, in species with the hidden break, RNA-Seq datasets will contain very few reads, in particular paired-end reads, in the hidden break region but will have normal levels of coverage on its flanking regions (28Sα and 28Sβ). We established a computational pipeline to determine the existence of the hidden break in representative species of as many phyla as possible in a semi-automated fashion.

### Identifying the hidden break region

In a first step we identified conserved sequences that could be used as flanking markers for the hidden break region described in protostomes [7], to help us determine potential homology of breaks in other organisms with the protostome break. We aligned 28S rRNA sequences from *Bombyx mori*, which has a characterised break region [16] and other species, with and without a described hidden break, using mafft [17]. We identified two highly conserved 20-mers (5’-AGUGGAGAAGGGUUCCAUGU-3’ and 5’-CGAAAGGGAATCGGGTTTAA-3’) flanking the hidden break region. These two 20-mers can be used as markers to identify the hidden break region characterised in protostomes when looking at other species.

### Establishing a pipeline to search for the hidden break

We used Python v3.7.0 to establish a semi-automated pipeline that proceeds from read mapping to results visualization (as graphs of read coverage). Firstly, our pipeline employs kallisto v0.44 [18] to map paired-end RNA-Seq reads against the respective 28S rRNA sequence from the species of interest. To ensure that kallisto pseudoalignments are not influencing the analysis [19], we also tested the slower STAR [20] and bwa [21] aligners and found no notable differences in the results (data not shown). Next, we use these mapped reads to calculate read coverage for each position in the 28S sequence using BEDtools v2.27.1 [22]. Finally, the pipeline uses Python’s ‘matplotlib’ library to produce plots of the depth of read mapping along the 28S rRNA molecule. We then inspected these plots. A species was considered to have the hidden break if there was an obvious drop in the read coverage between the two conserved flanking 20-mers. The method was tested using data from *Bombyx mori*, which has an experimentally identified and characterised hidden break.

### Data collection and large-scale application of the pipeline

We retrieved all available entries (as of May 2019) from the SILVA database [23] in a single FASTA file comprising 633,348 eukaryotic 28S sequences. We filtered these data to obtain a set of sequences that could be analysed with our method by removing all duplicates, (i.e. entries coming from the same species) and discarding all sequences shorter than 2,000bp. After these filtering steps we retained 28S RNA sequences from 12,460 species.

We next searched for RNA-Seq data for each of these species on the SRA database using eutils. We set a threshold requirement of at least 1 Gigabase of RNA-Seq data and imposed an upper limit of 4 Gigabases for our analysis. We retrieved data within these size boundaries from 1,024 species. A list of the species that produced an interpretable plot can be found in Supplementary Table 1, and the corresponding coverage plots can be seen in Supplementary File 1.

To achieve maximum representation of animal phyla in our results, we manually added 80 species (highlighted in bold, Supplementary Table 1) to those emerging from the semi-automated pipeline. The 28S rRNA sequences for these species were retrieved from NCBI and RNAcentral databases [24] and the paired-end RNA-Seq data from the SRA database [25]. For 40 species only we also calculated the proportion of read pairs spanning each residue of 28S using SAMtools v1.9 (option ‘view -F 12’, [26]). This metric was highly correlated to the read coverage and was not applied to the rest of the species. We also calculated the number of forward and reverse reads mapped in the interval between 300 nucleotides before the break and 300 nucleotides after using SAMtools (options ‘view -f 67’ for forward and ‘view -f 131’ for reverse reads). We then calculated the ratios of forward/reverse and reverse/forward reads for each position and visualised the result, using logarithms of ratios to make the fluctuations symmetric. This analysis was run for 17 species (10 with and 7 without the hidden break) and the results can be seen in Supplementary File 2.

## ACKNOWLEDGEMENTS

We thank Kate Rawlinson for the electropherograms shown in Fig 1. We thank members of the Telford lab for useful feedback. This work was supported by the European Union’s Horizon 2020 research and innovation programme under the Marie Skłodowska-Curie grant agreement No 764840 to M.T.; the European Research Council (ERC-2012-AdG 322790) to M.T. and Human Frontier Science Program (HFSP RGP0002/2016) to M.T.

## AUTHOR CONTRIBUTIONS

MJT proposed original idea. MJT, ISM, PHS developed idea and guided research. PN conducted research with help from ISM, PHS. MJT wrote initial draft. MJT, PN, PHS, ISM contributed to final draft. PN and ISM prepared figures and table. All authors reviewed the manuscript.

## ADDITIONAL INFORMATION

The authors declare no competing interests. A list of all retrieved species and the corresponding data sets and can be found under: https://github.com/pnatsi/hiddenbreak/tree/master/data/suitable_species.tsv. The code that performs the analysis and plots the result can be found under: https://github.com/pnatsi/HBinspector.

