## Supplementary figures and images for "Computational discovery of hidden breaks in 28S ribosomal RNAs across eukaryotes and consequences for RNA Integrity Numbers"

### Acyrthosiphon_pisum_coverage.png

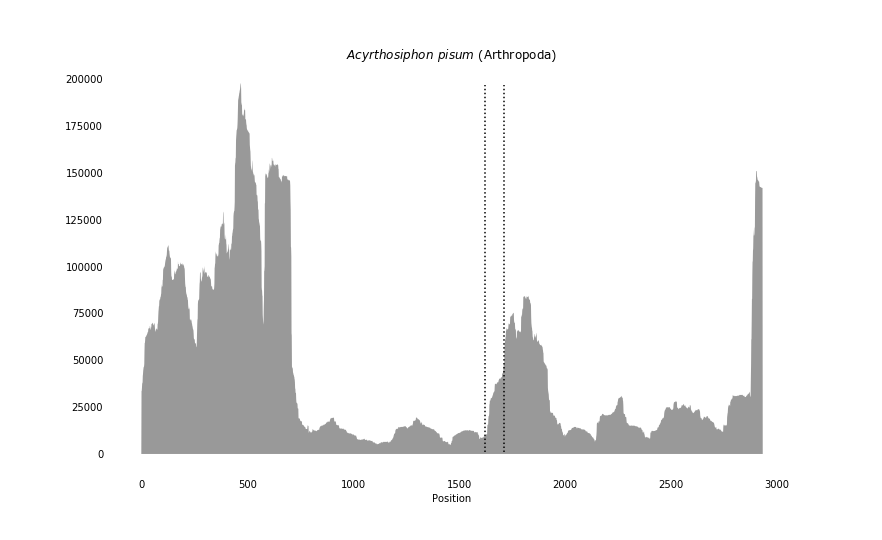

### Anadara_trapezia_coverage.png

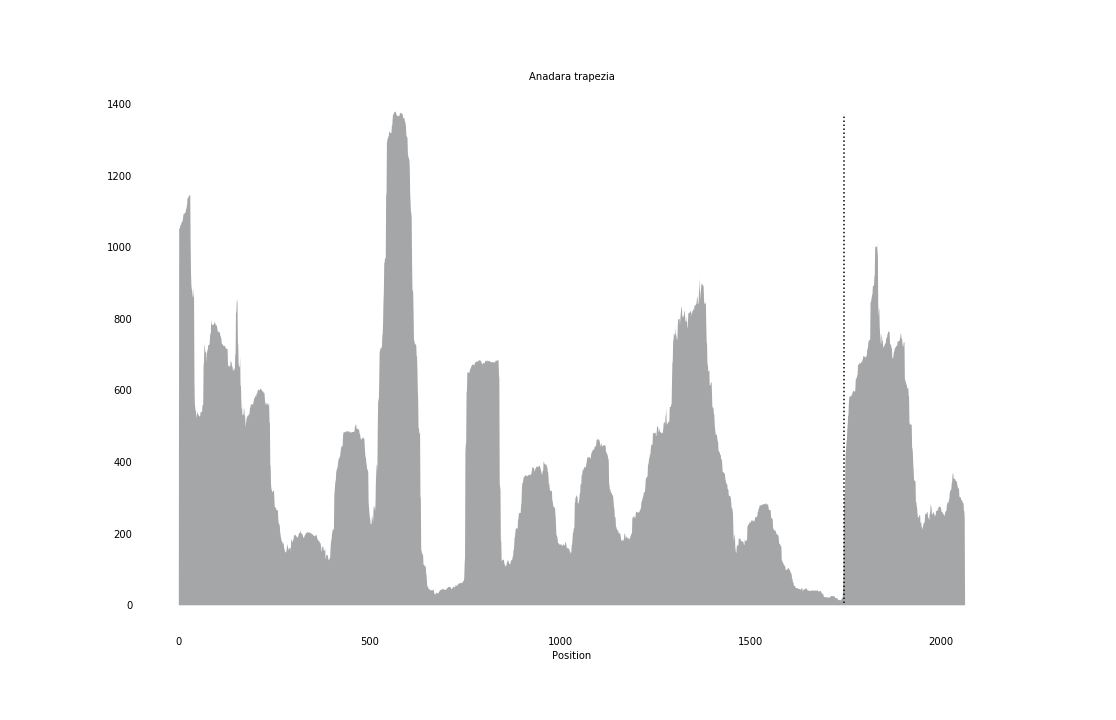

### Antheraea_pernyi_coverage_correct.png

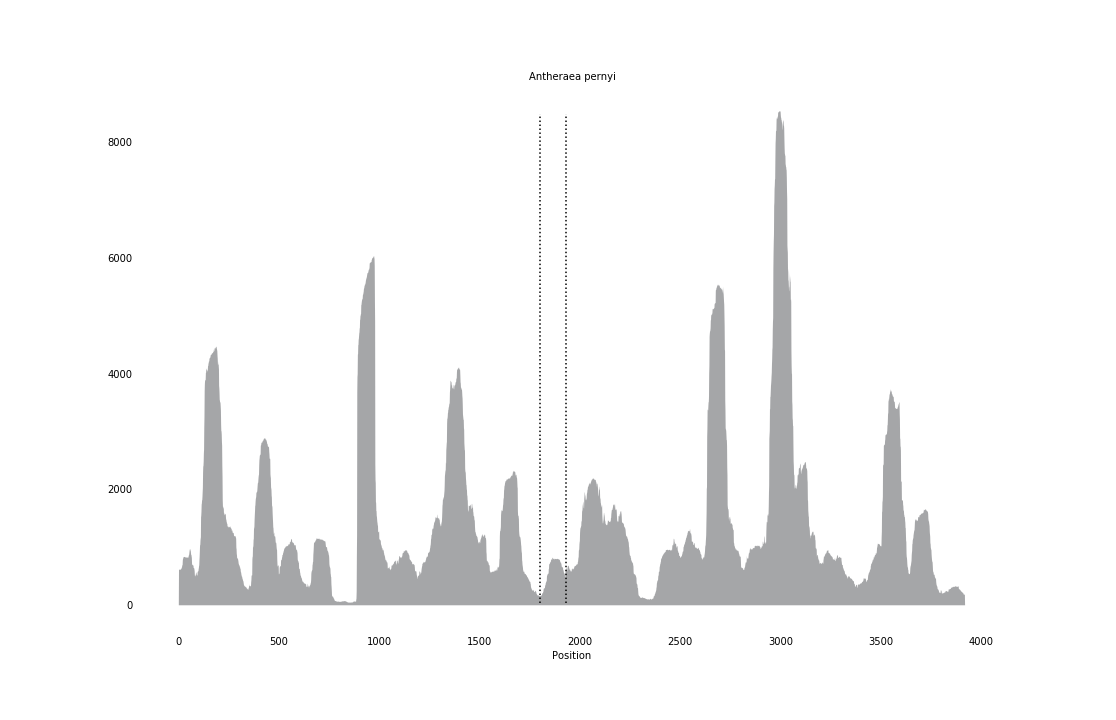

### Anurida_maritima_coverage_correct.png

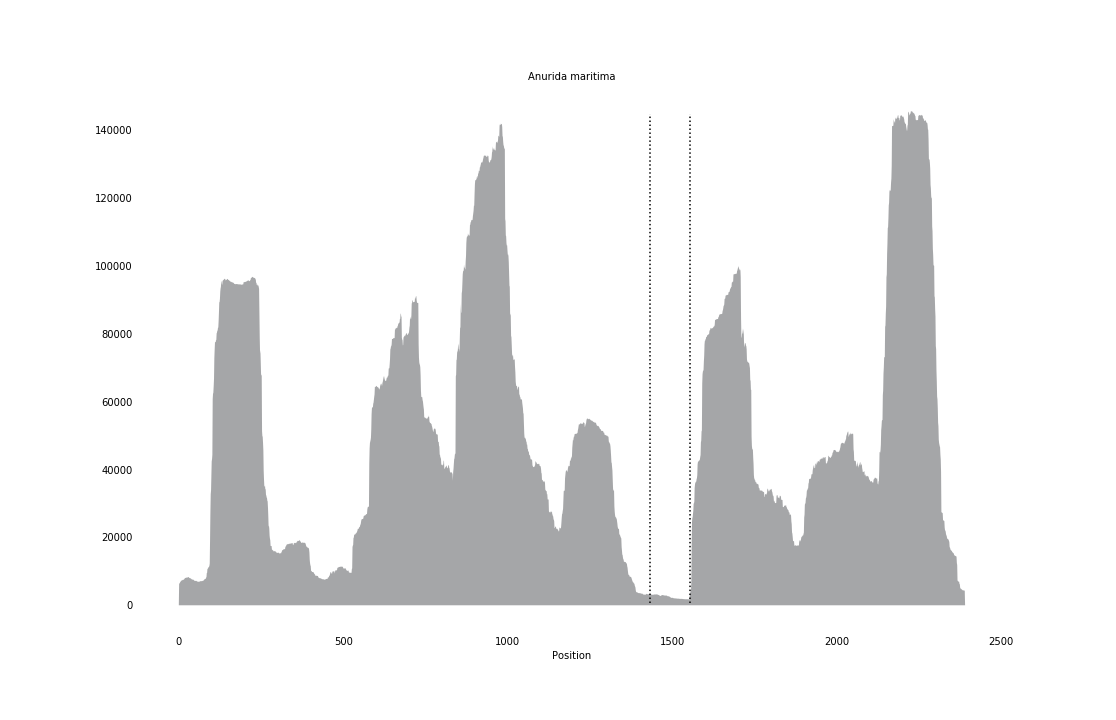

### Aphidius_colemani_coverage_correct.png

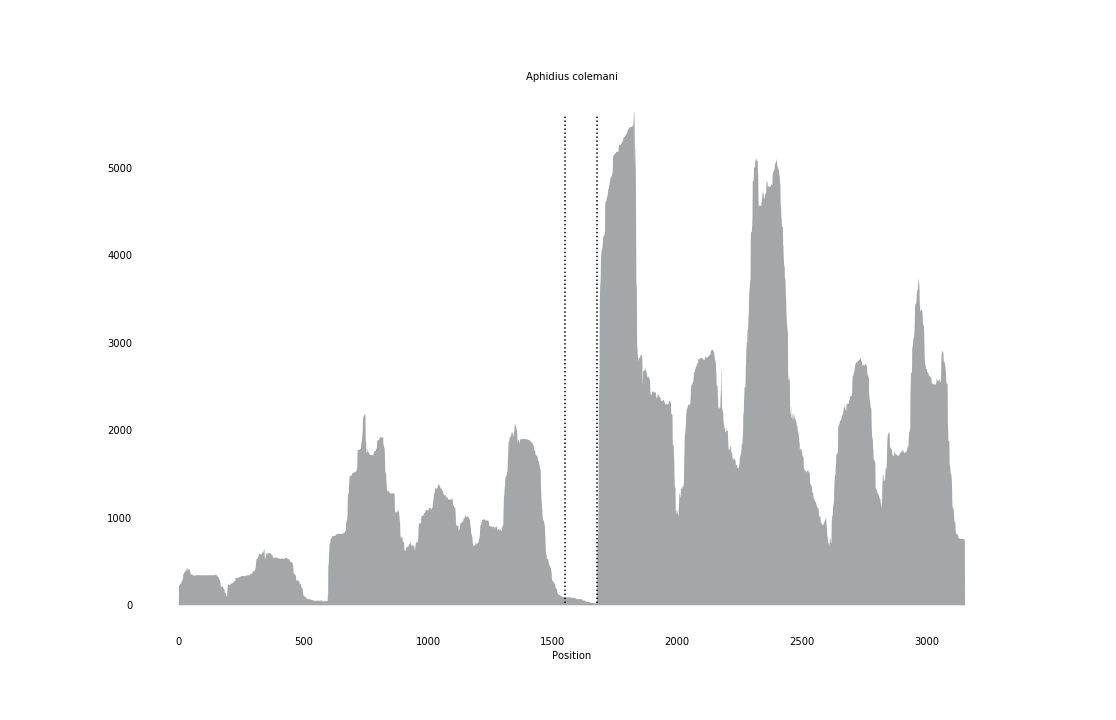

### Argopecten_irradians_coverage_correct.png

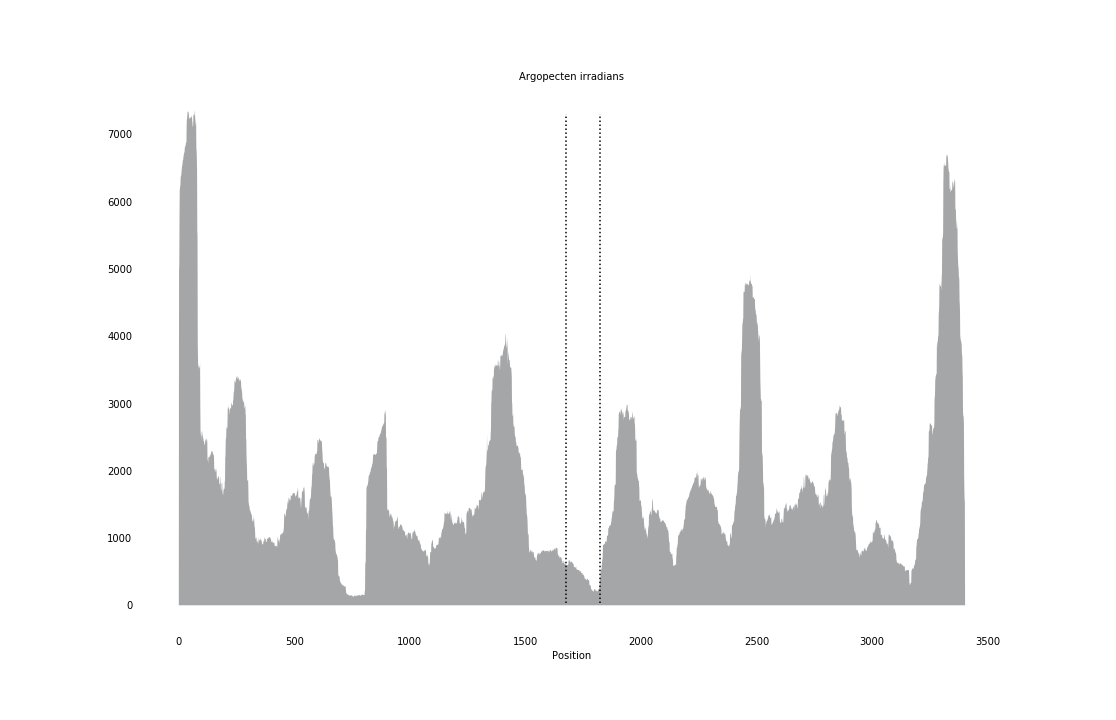

### Atelura_formicaria_coverage_correct.png

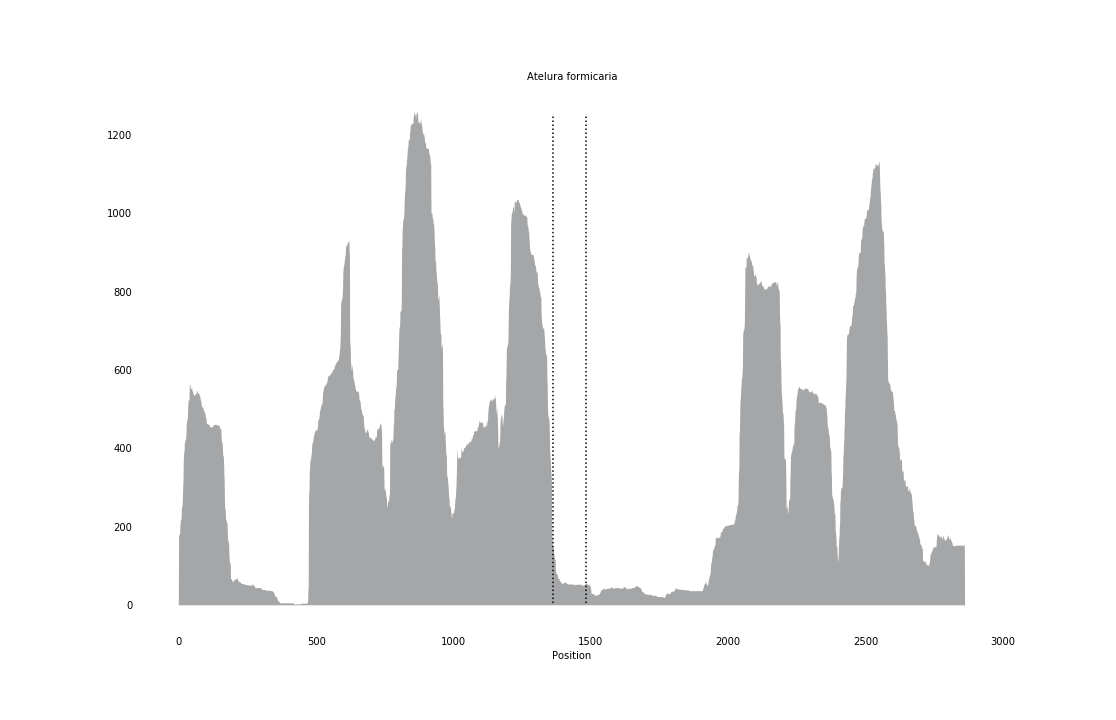

### Athalia_rosae_coverage_correct.png

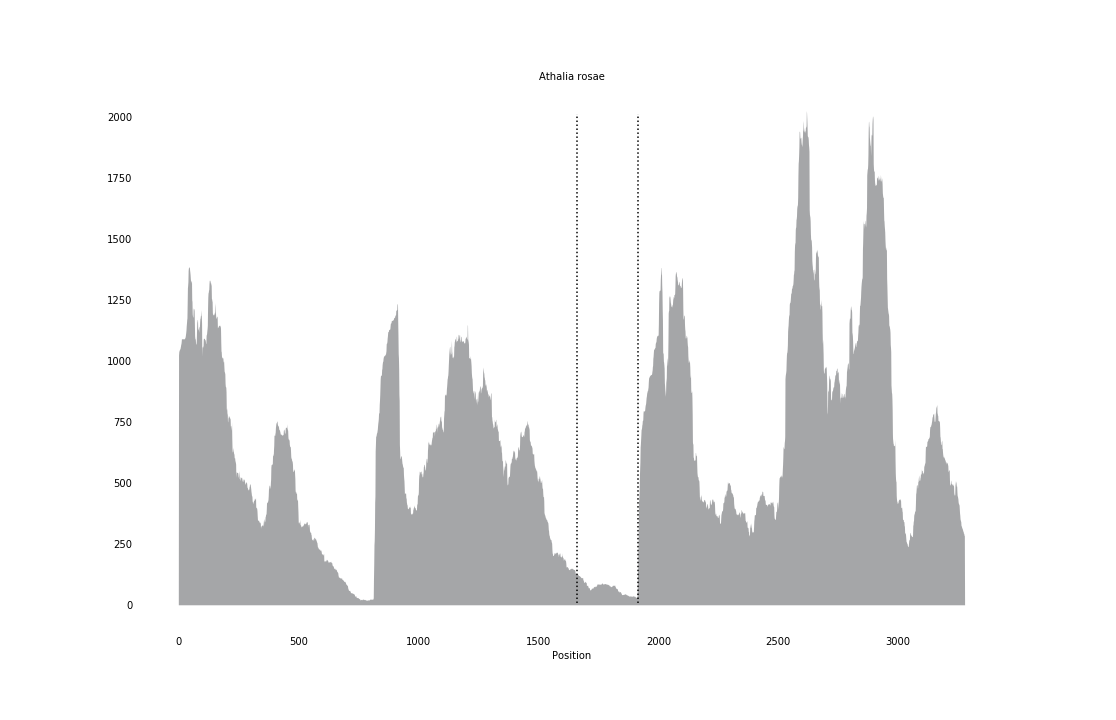

### Atrina_rigida_coverage.png

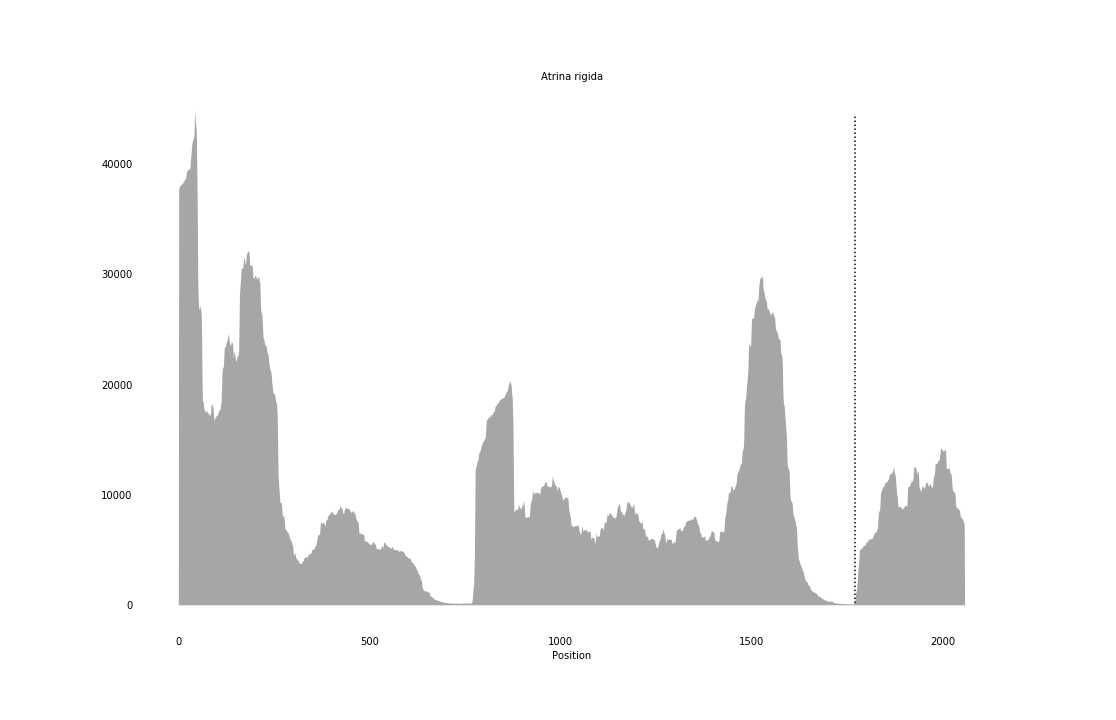

### Bembix_rostrata_coverage_correct.png

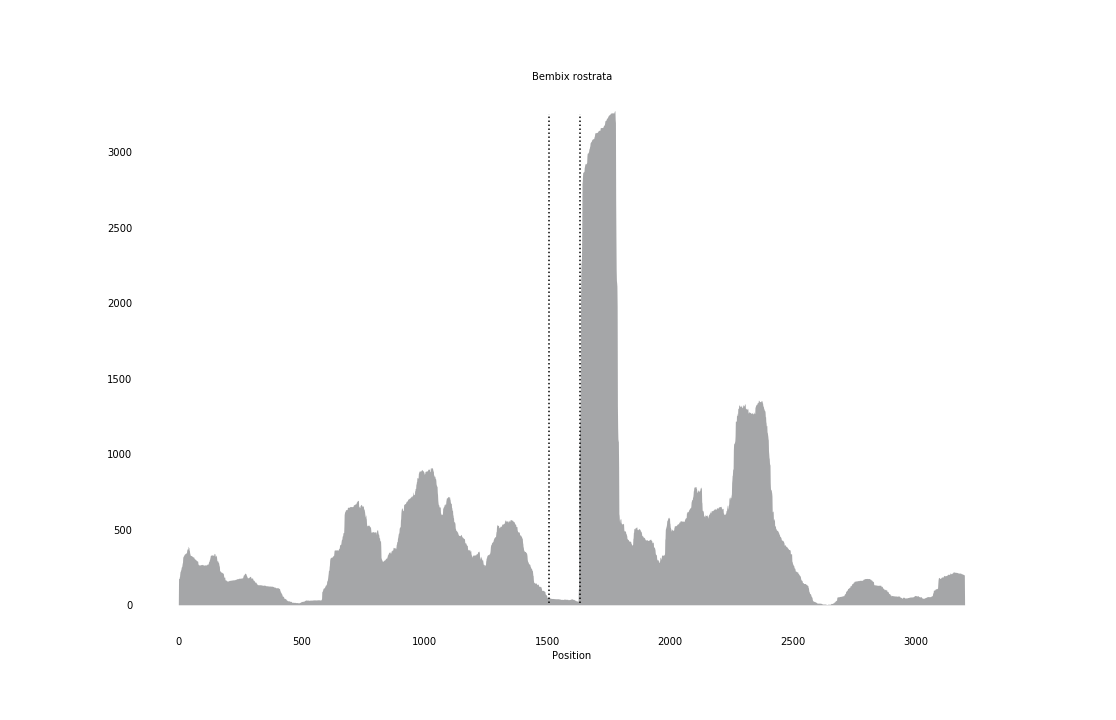

### Blattella_germanica_overage.png

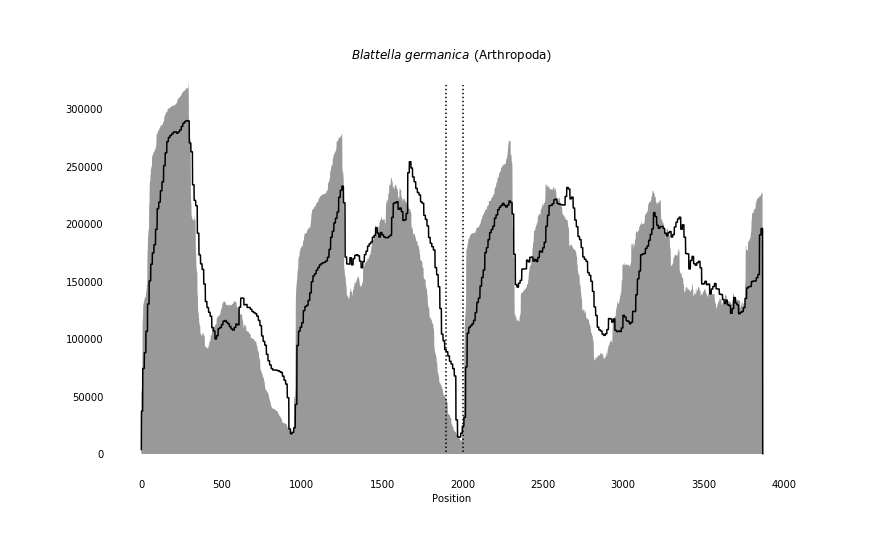

### Bombylius_major_coverage_correct.png

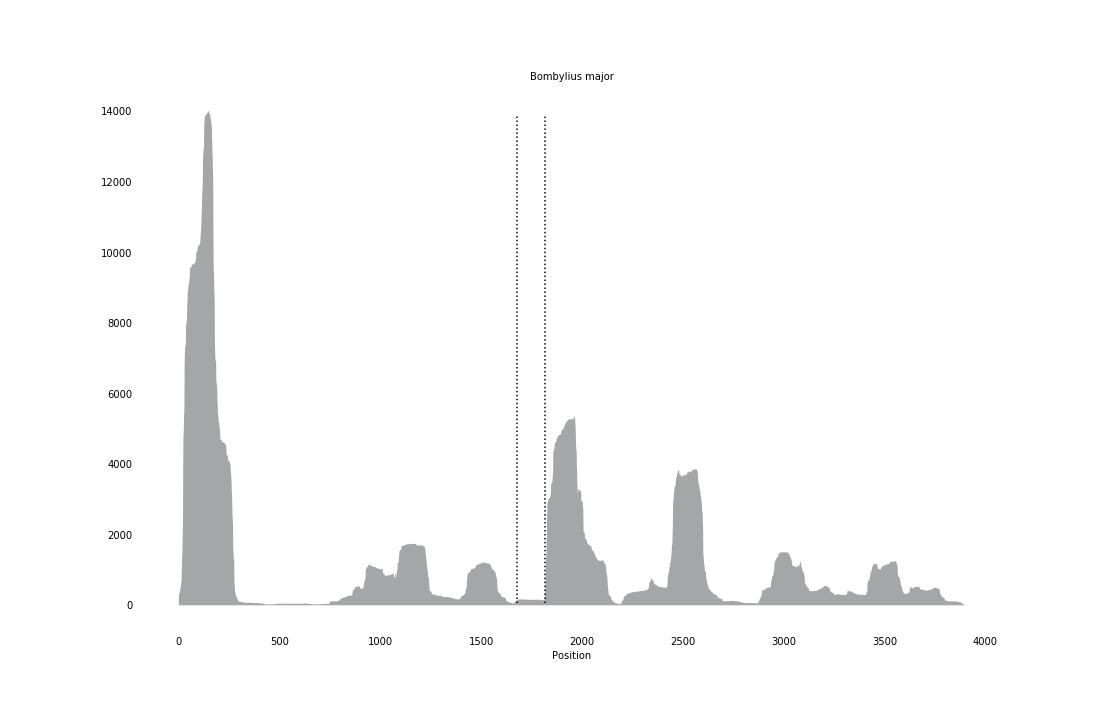

### Bombyx_mori_overage.png

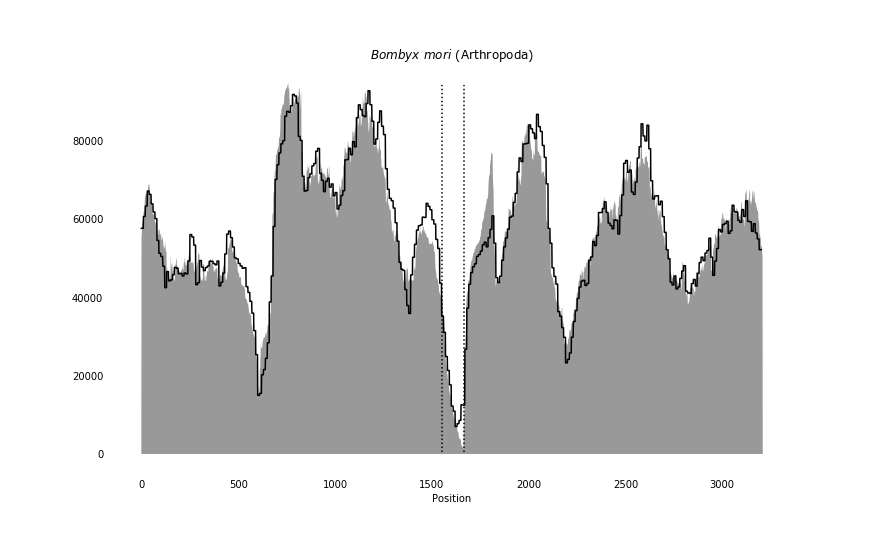

### Brachygaster_minutus_coverage_correct.png

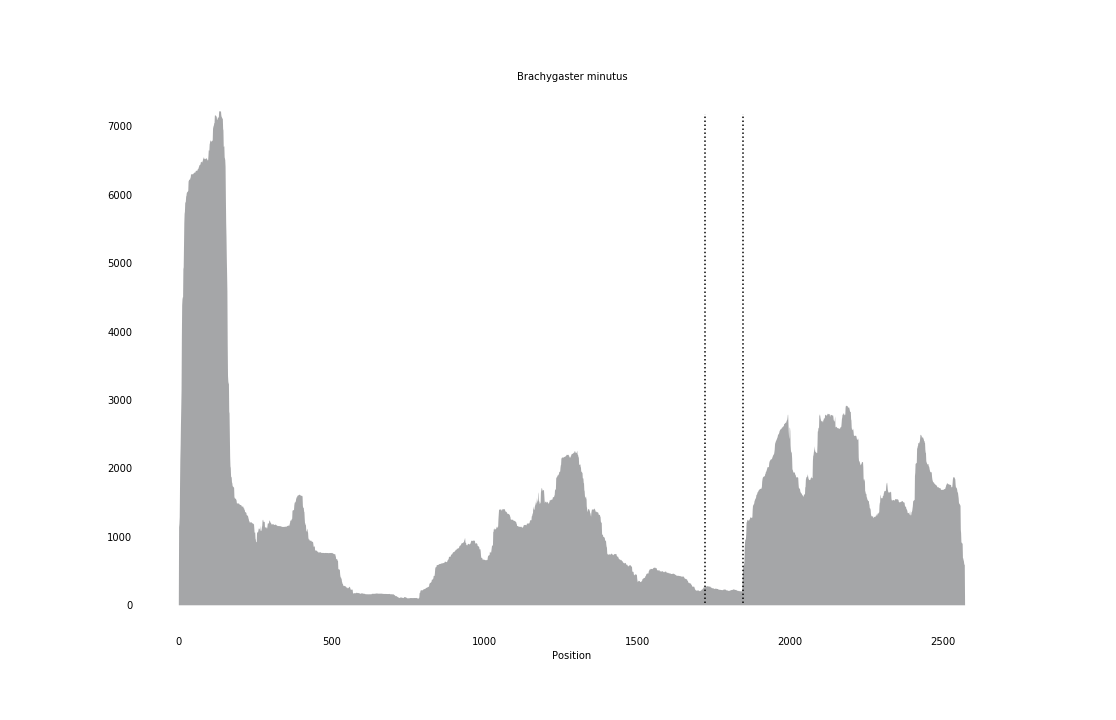

### Brachymeria_minuta_coverage_correct.png

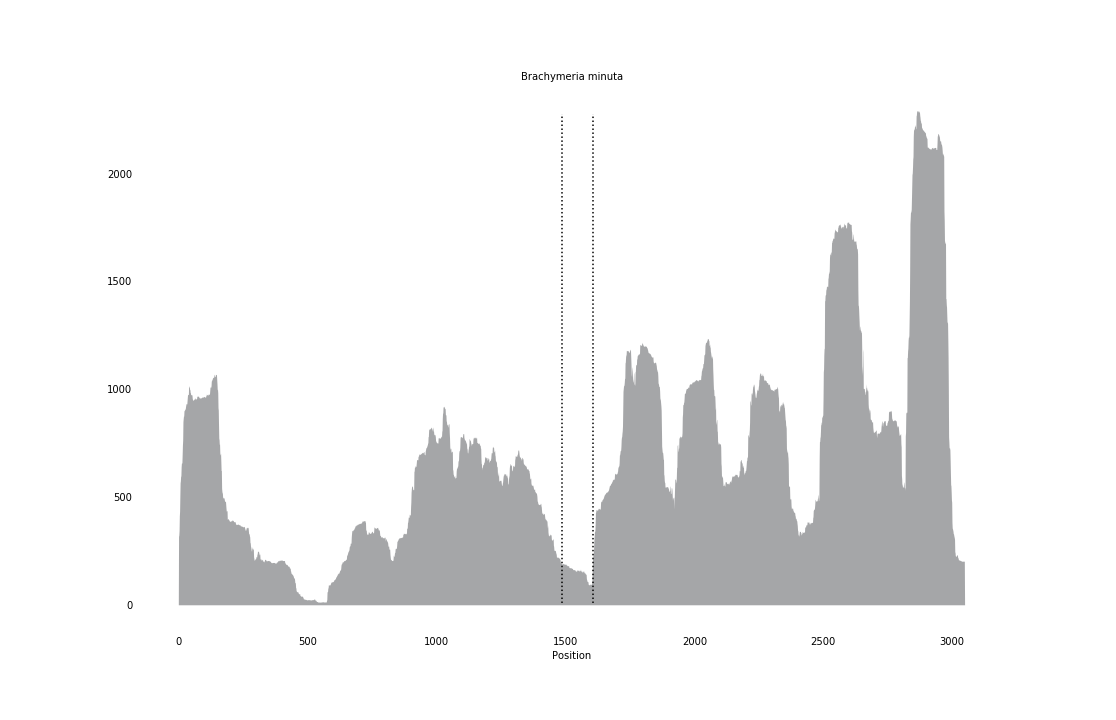

### Carausius_morosus_coverage_correct.png

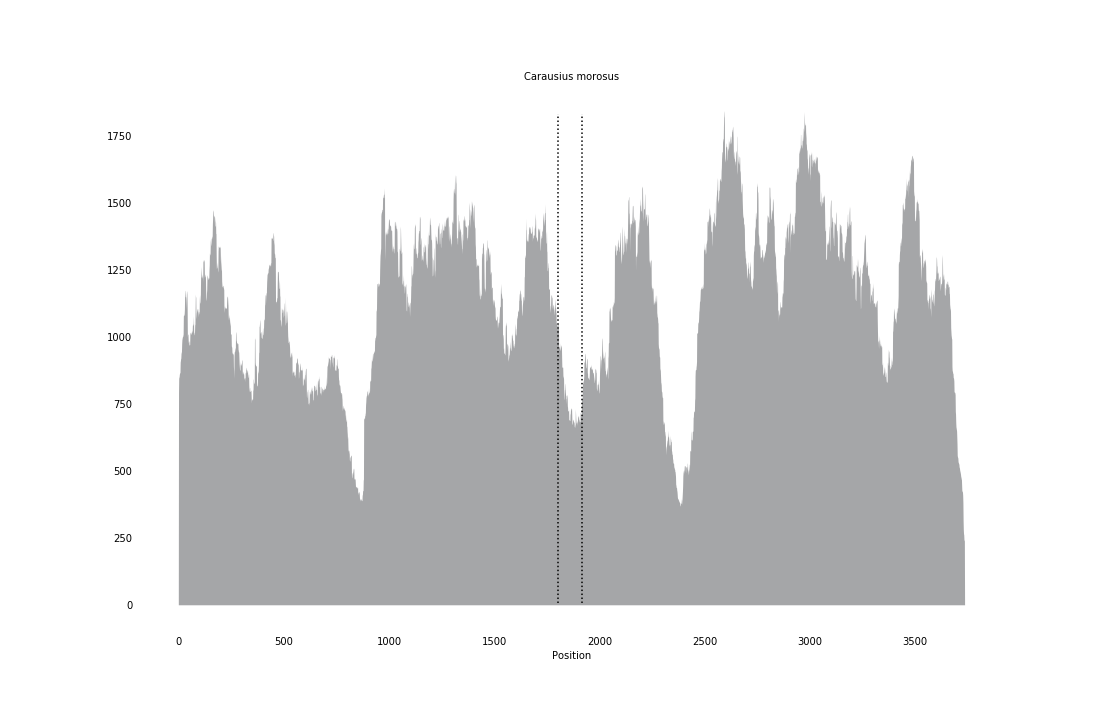

### Cardites_antiquatus_coverage.png

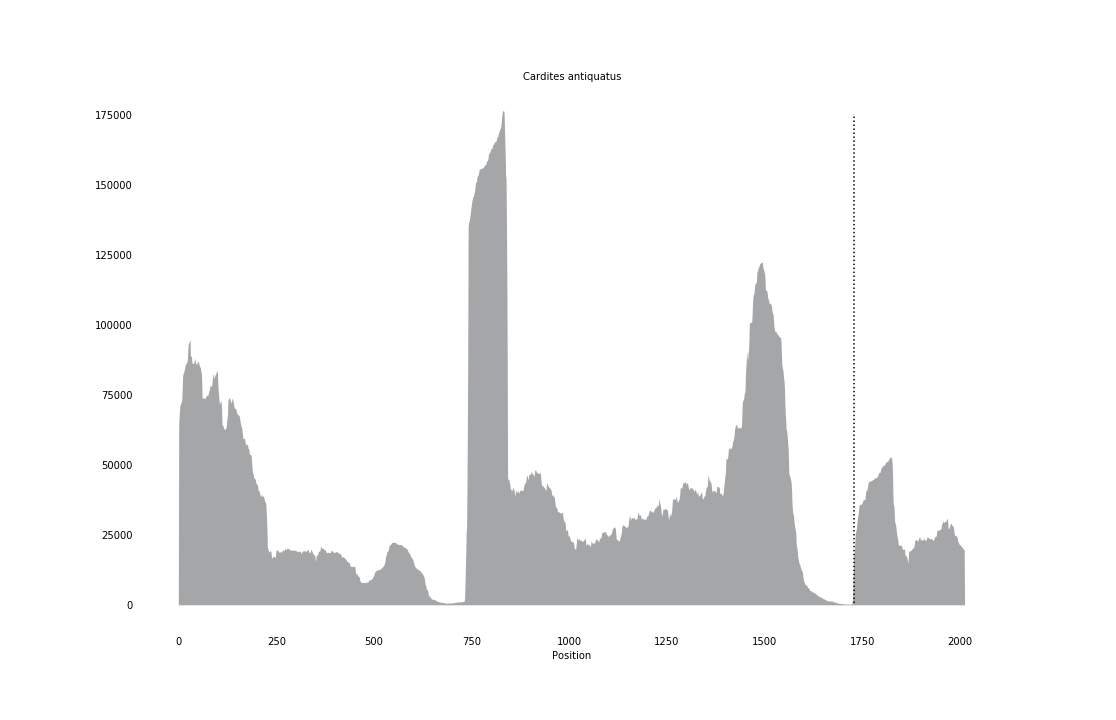

### Cerastoderma_edule_coverage.png

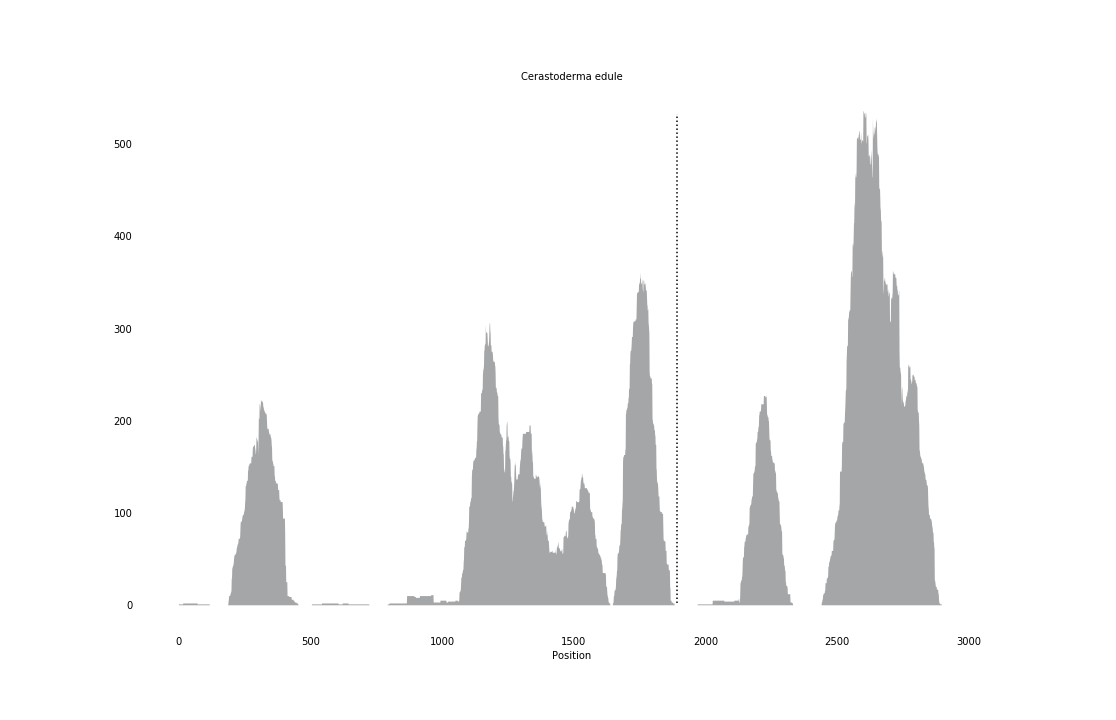

### Cerceris_arenaria_coverage_correct.png

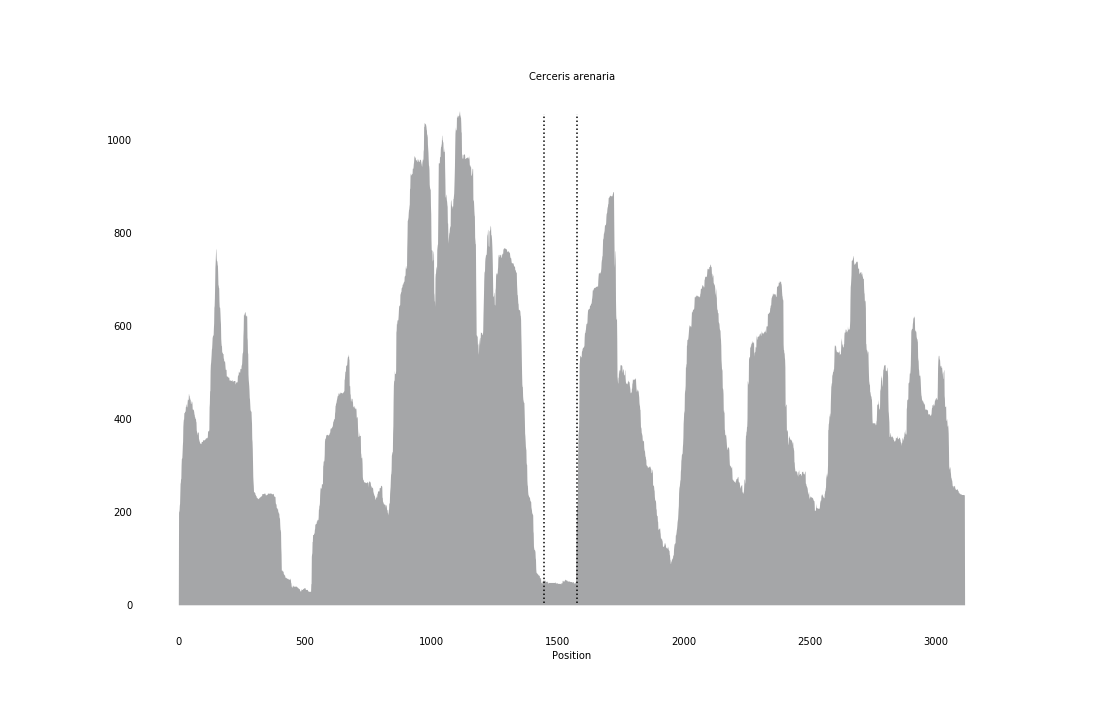

### Chrysis_viridula_coverage_correct.png

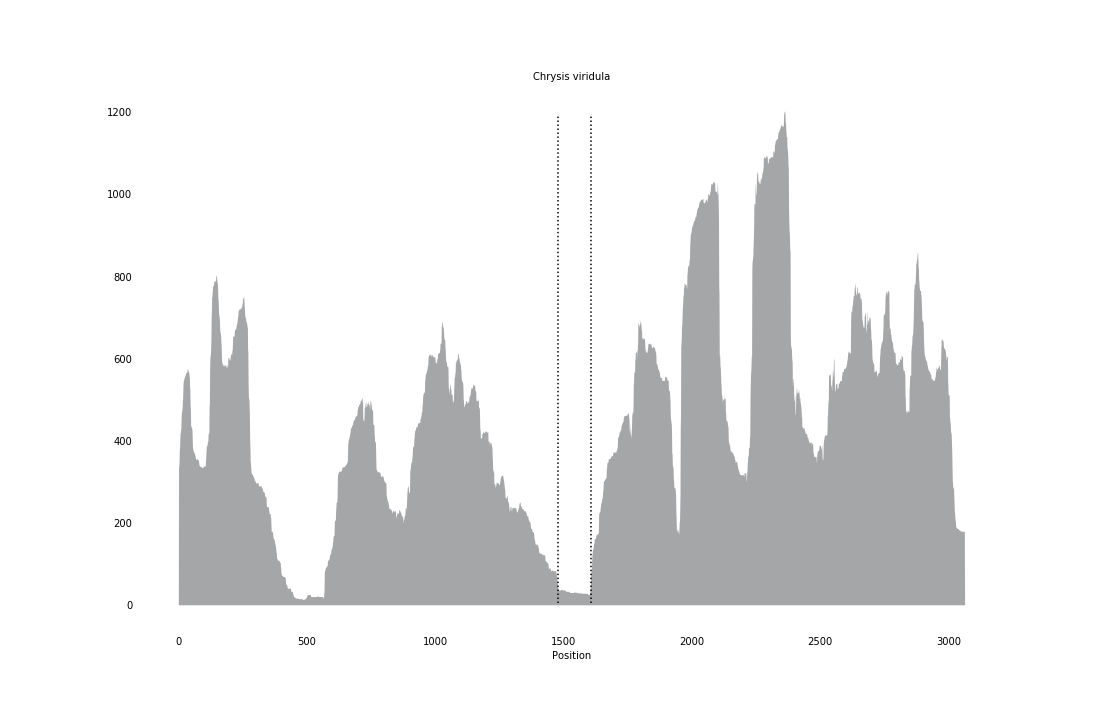

### Clogmia_albipunctata_coverage_correct.png

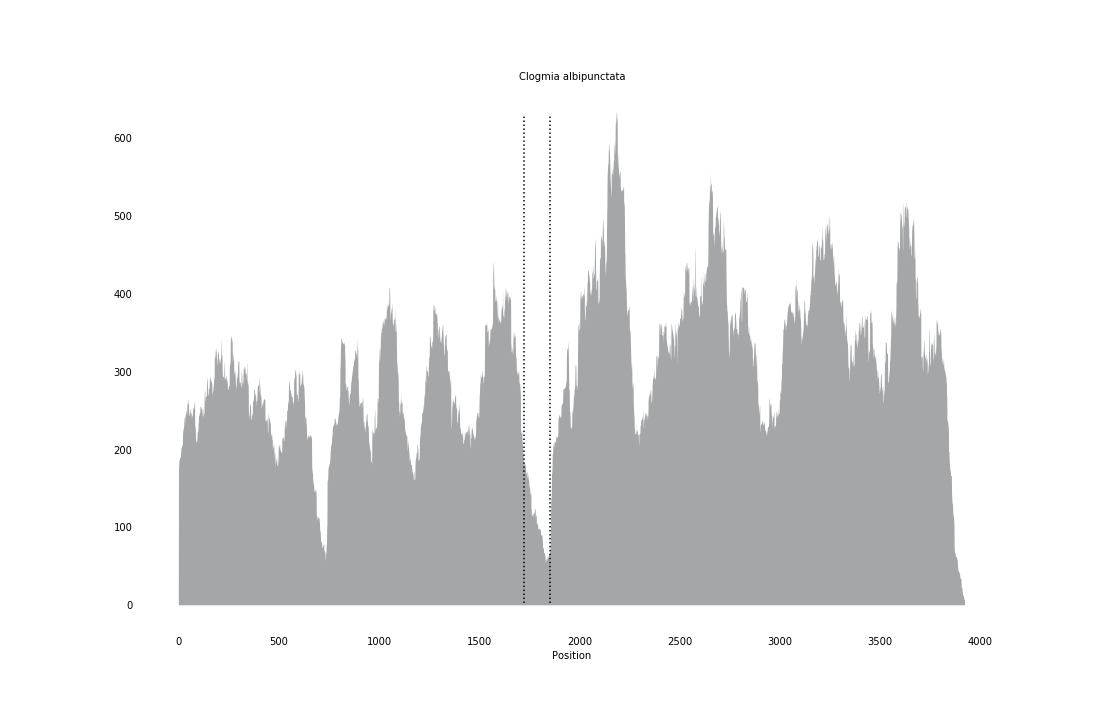

### Cochliomyia_hominivorax_coverage_correct.png

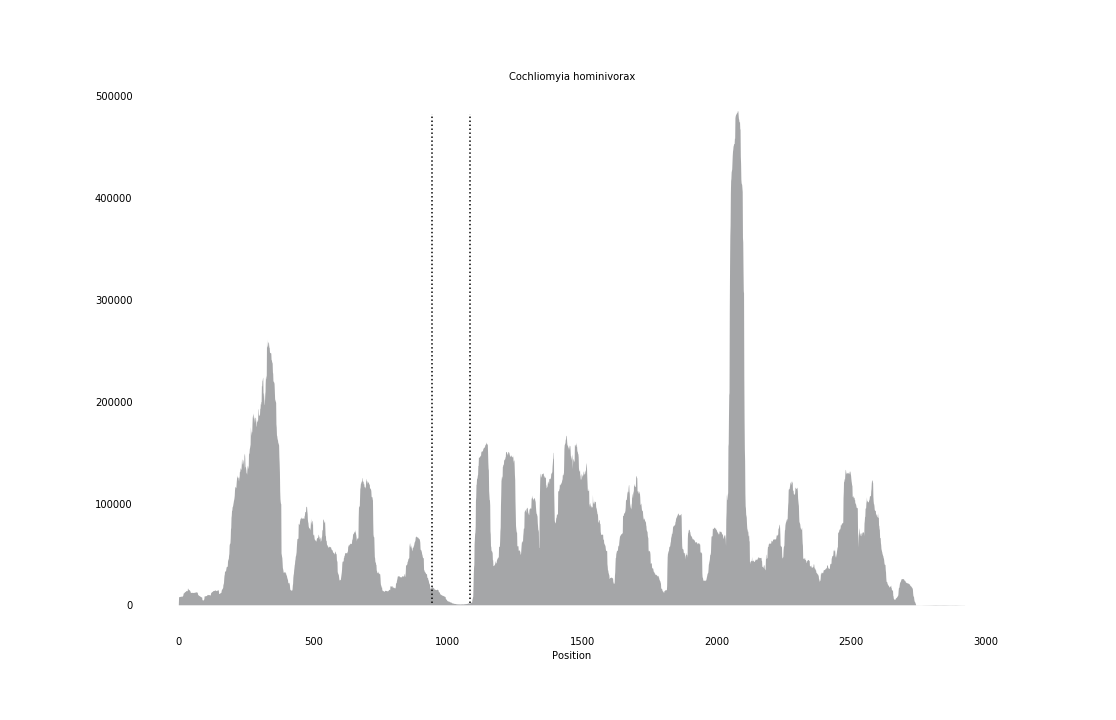

### Corbicula_fluminea_coverage.png

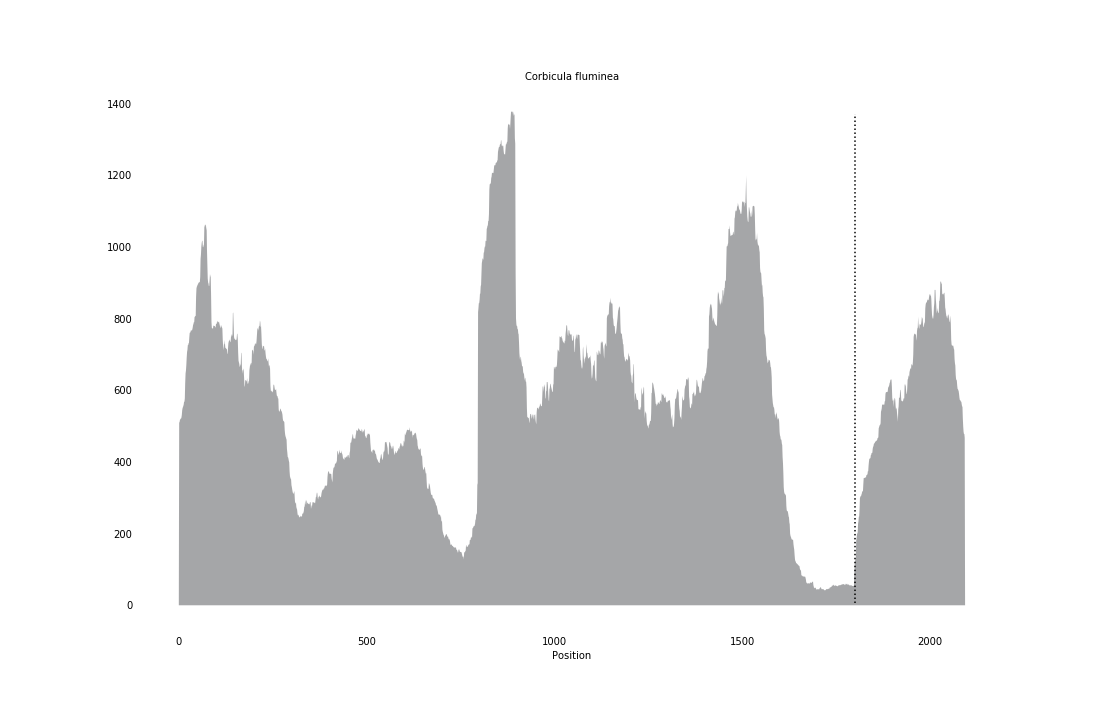

### Crassostrea_gigas_coverage_correct.png

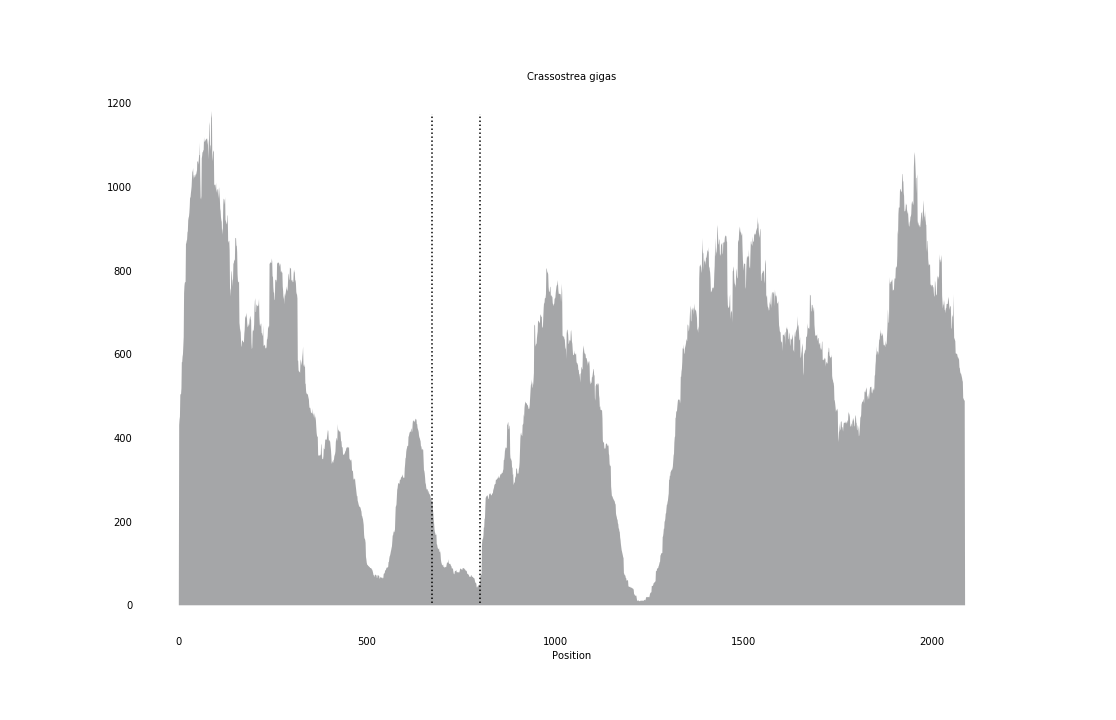

### Ctenoides_scaber_coverage.png

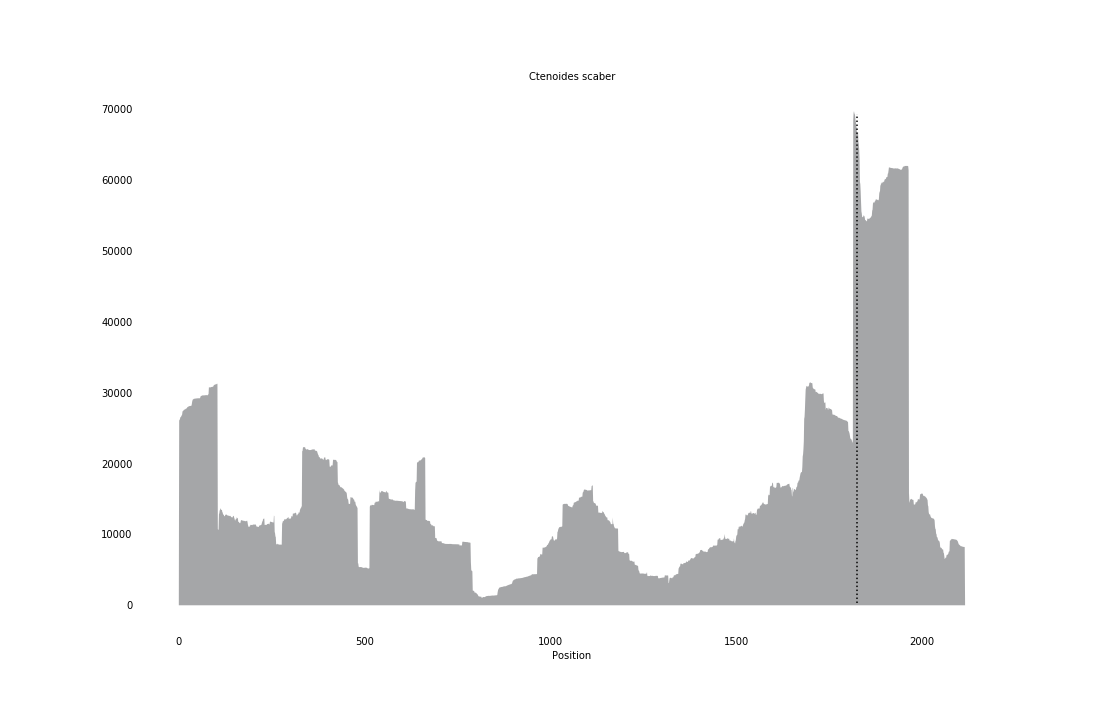

### Cycadothrips_chadwicki_coverage.png

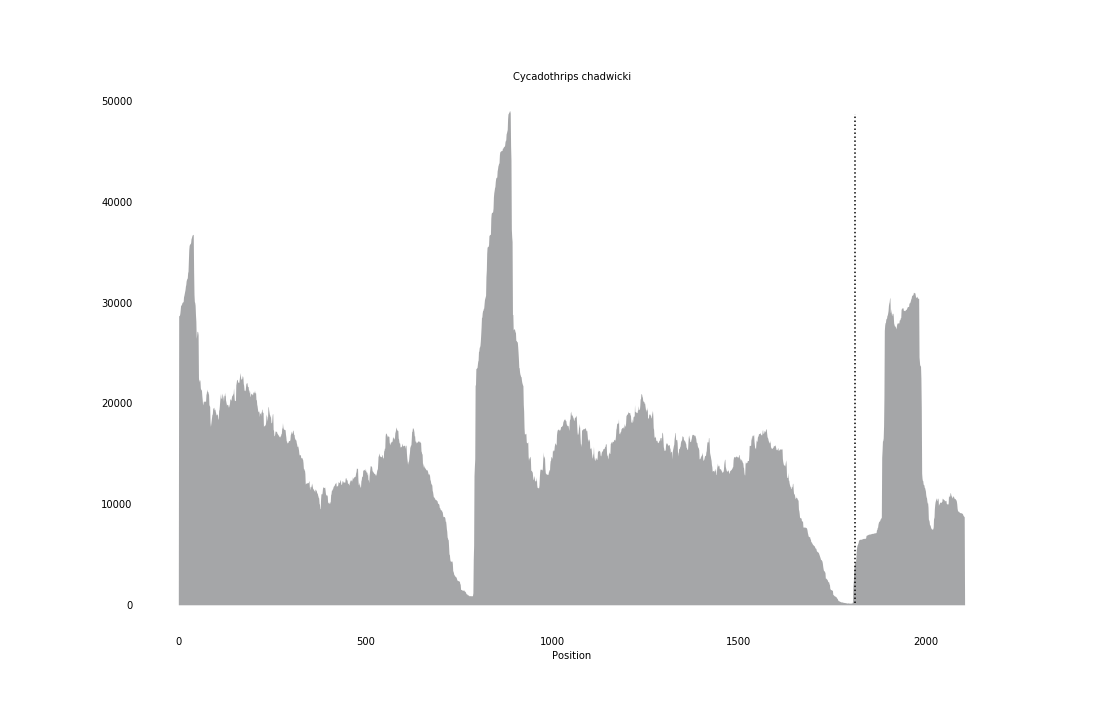

### Cyrenoida_floridana_coverage.png

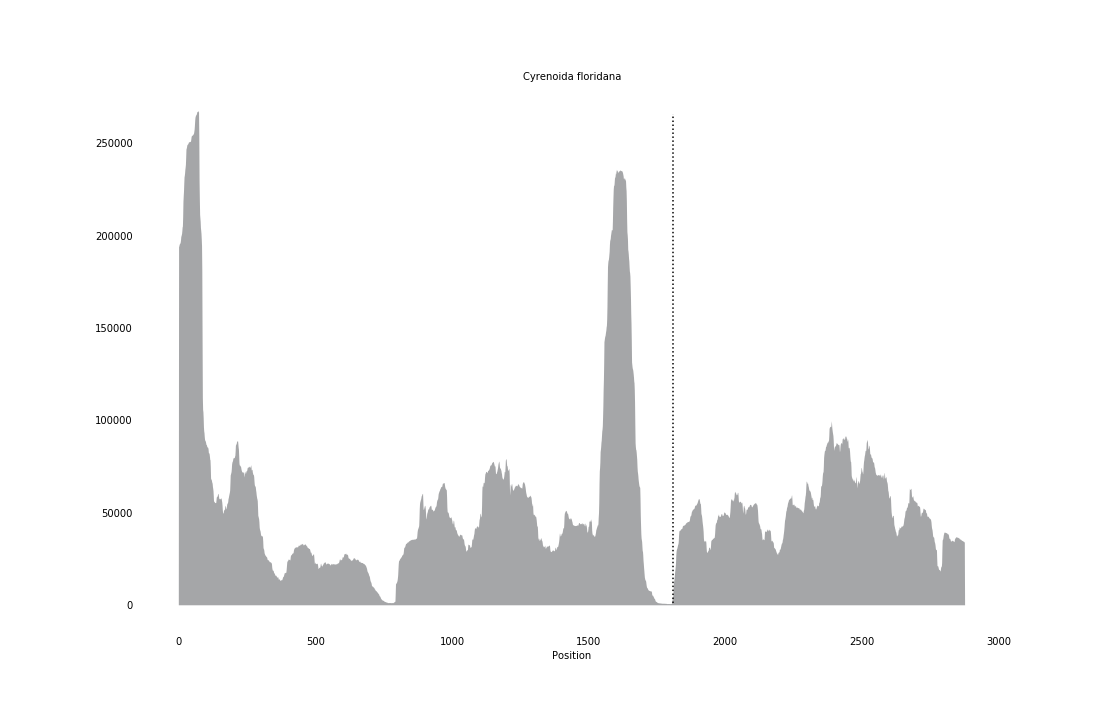

### Diaeretus_essigellae_coverage_correct.png

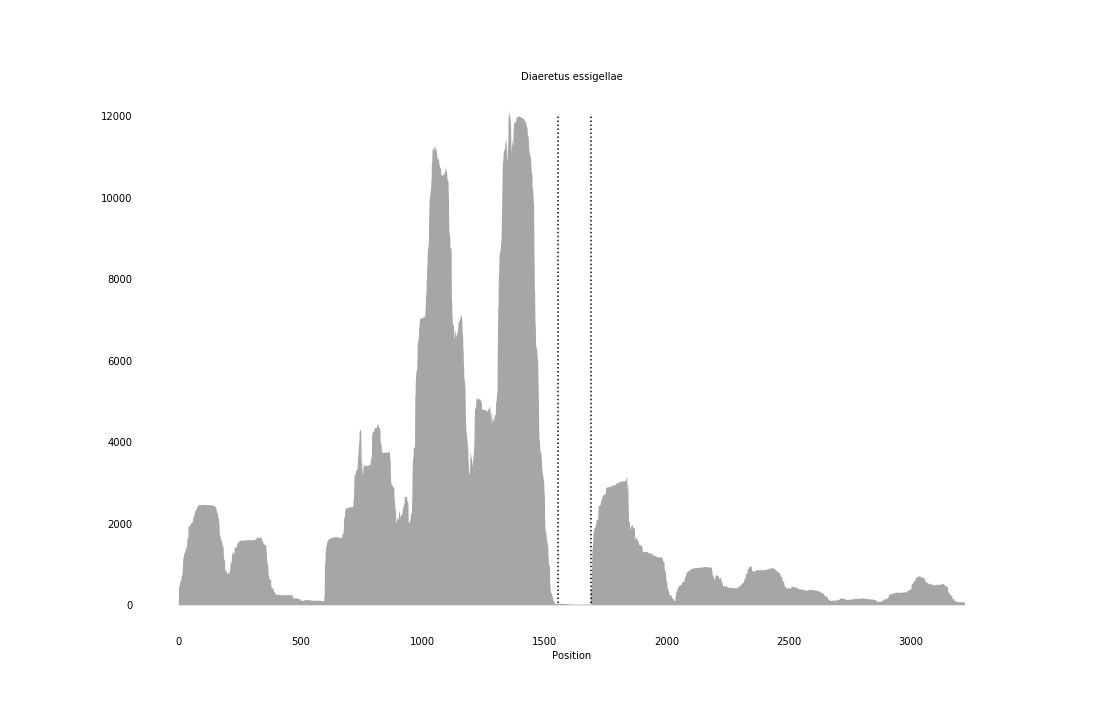

### Diglyphus_isaea_coverage_correct.png

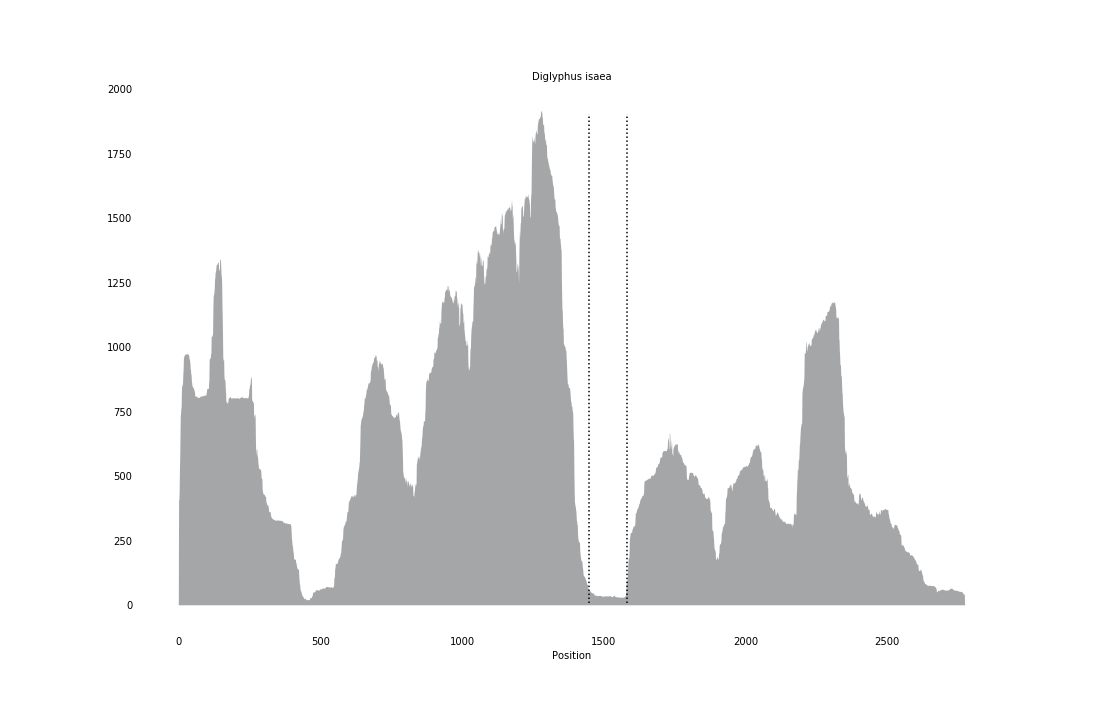

### Dinetus_pictus_coverage_correct.png

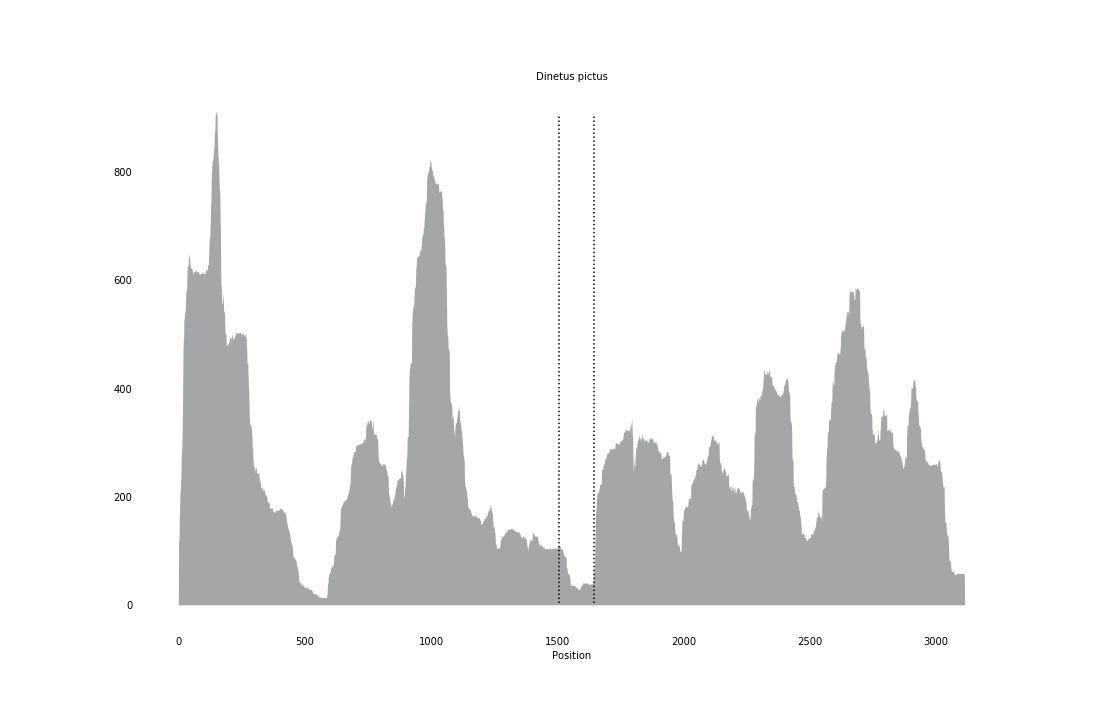

### Ducetia_japonica_coverage.png

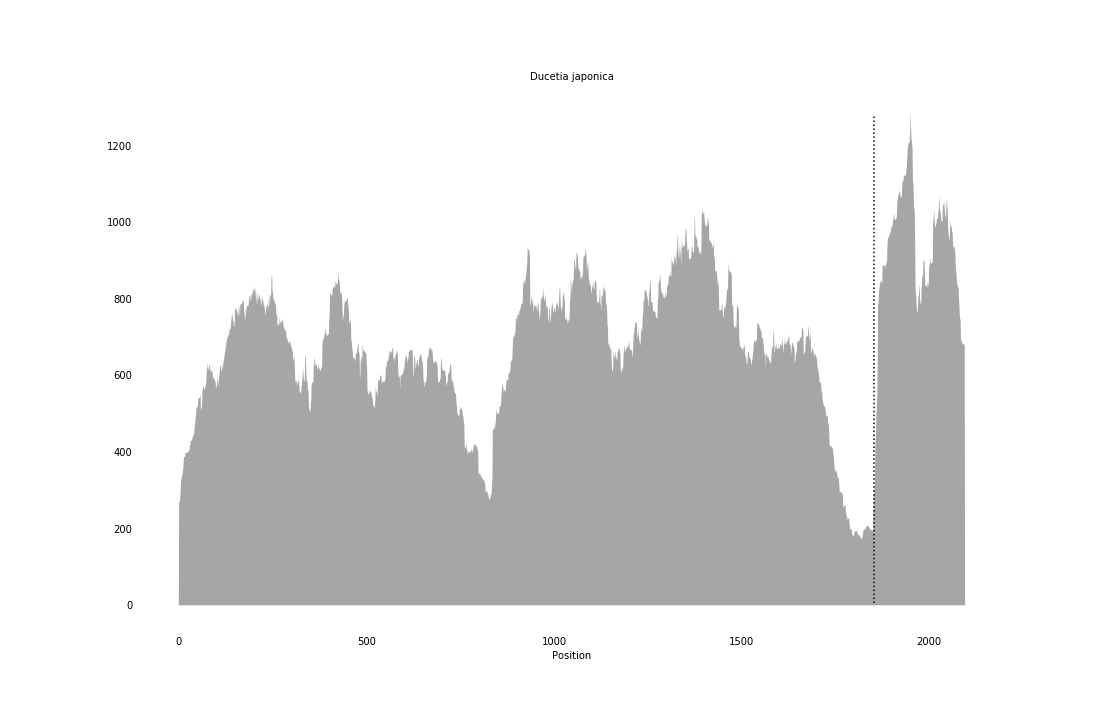

### Elampus_spina_coverage_correct.png

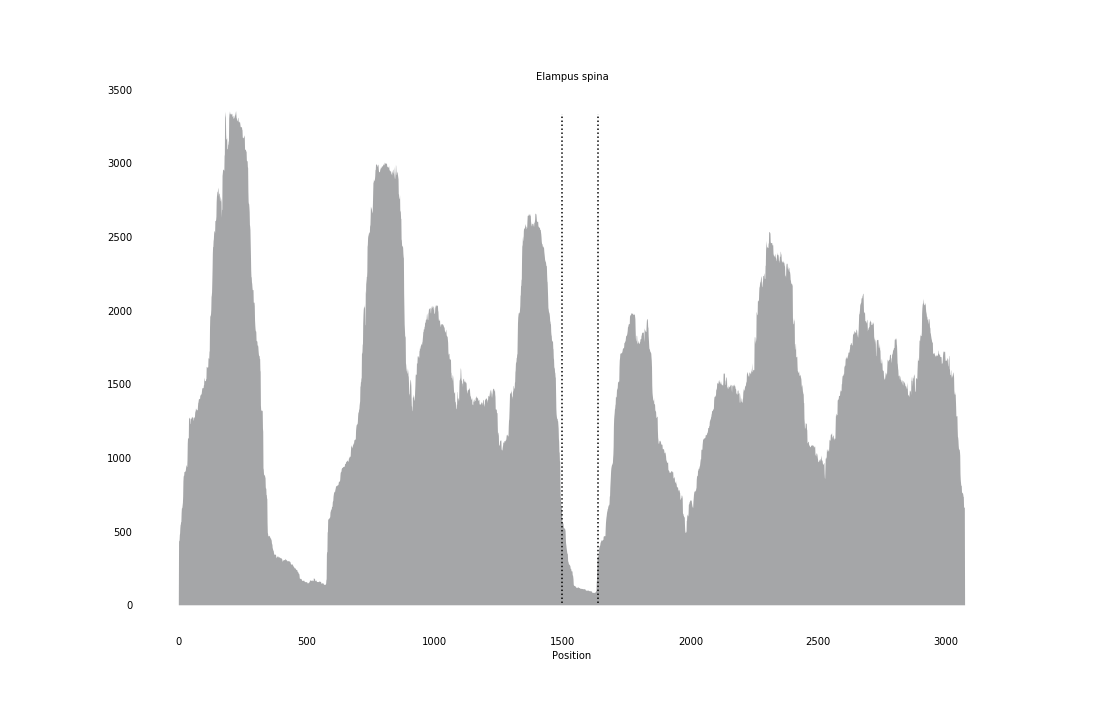

### Empoasca_fabae_coverage_correct.png

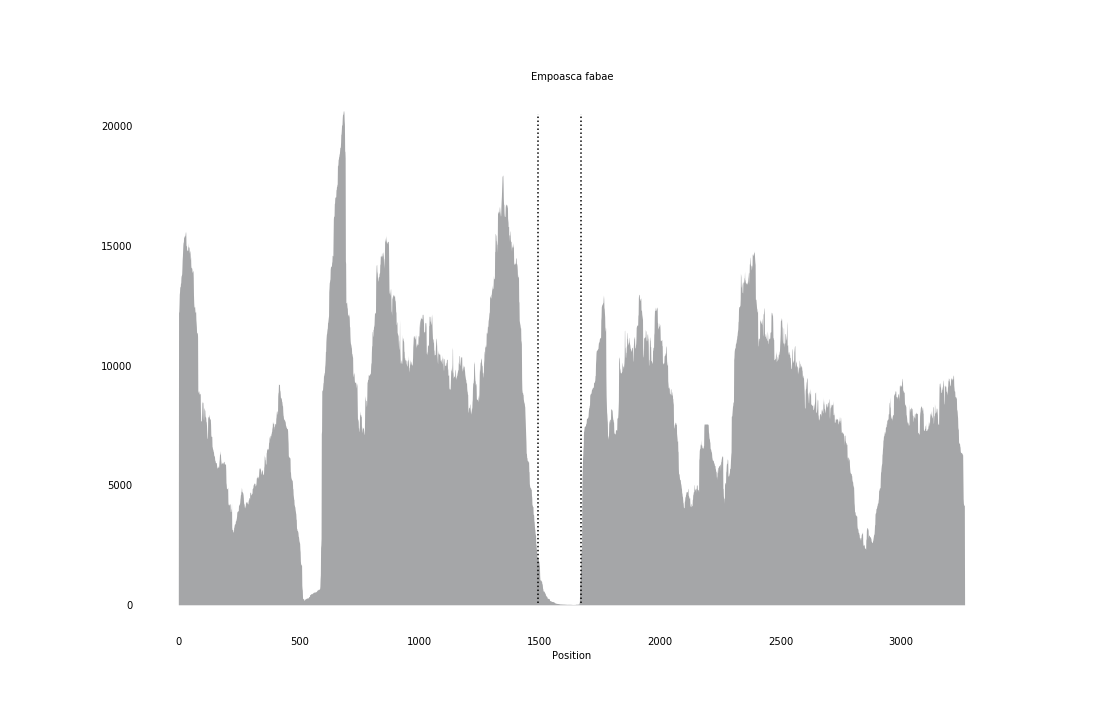

### Episyrphus_balteatus_coverage.png

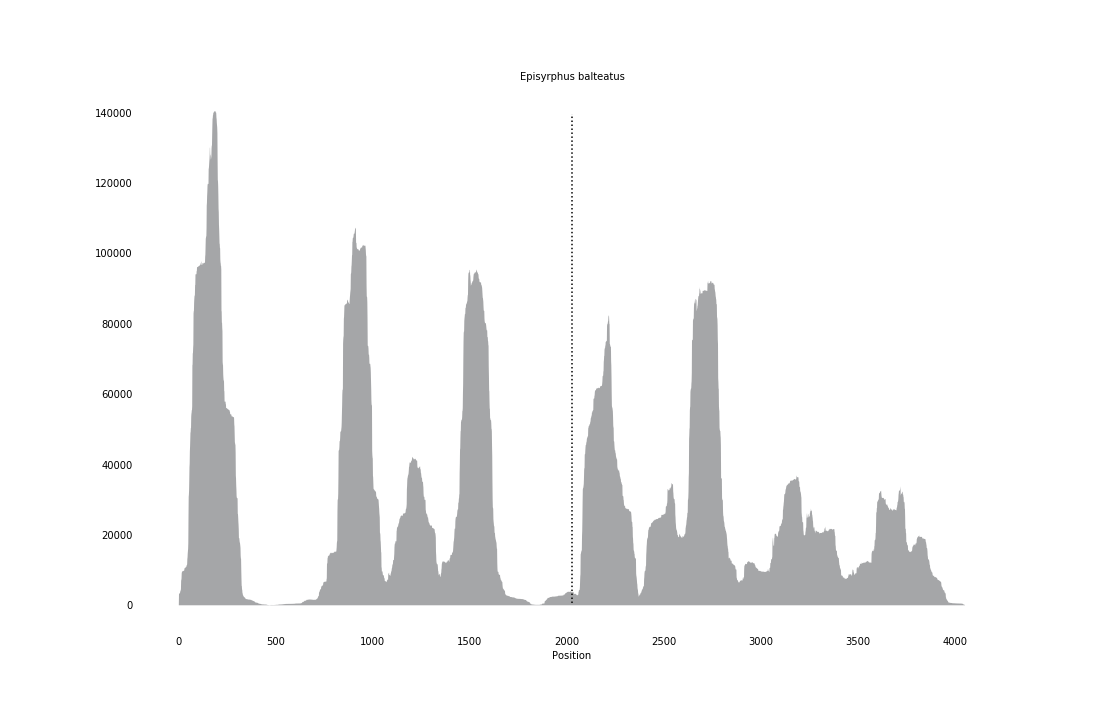

### Eucera_syriaca_coverage_correct.png

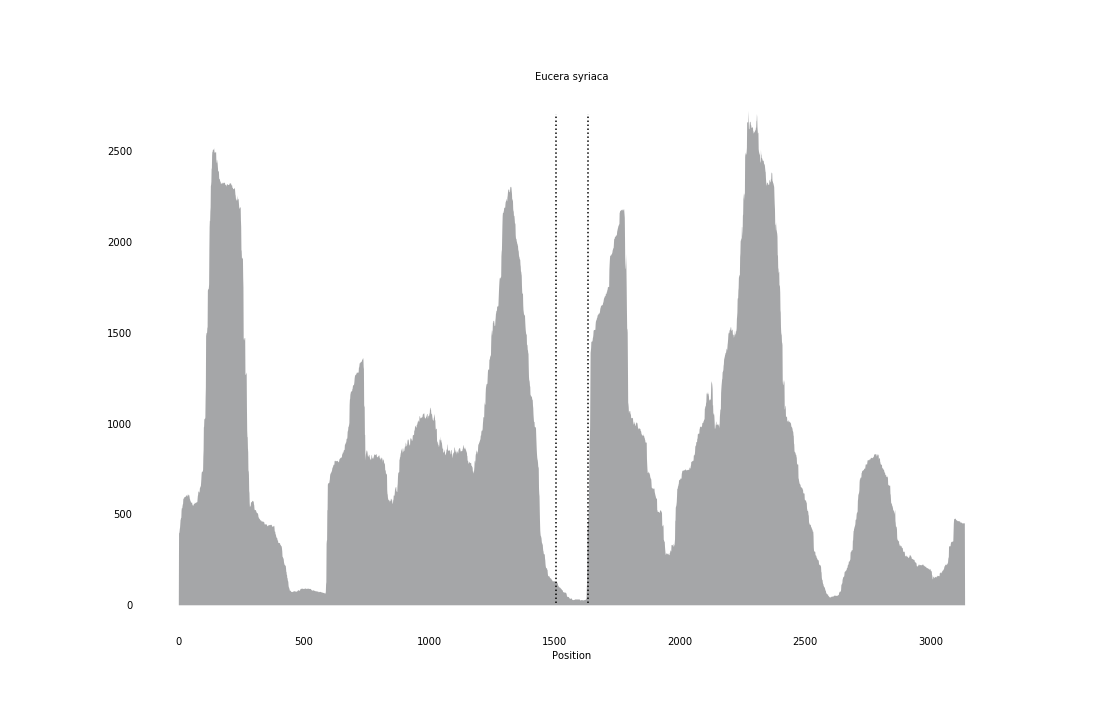

### Fannia_canicularis_coverage_correct.png

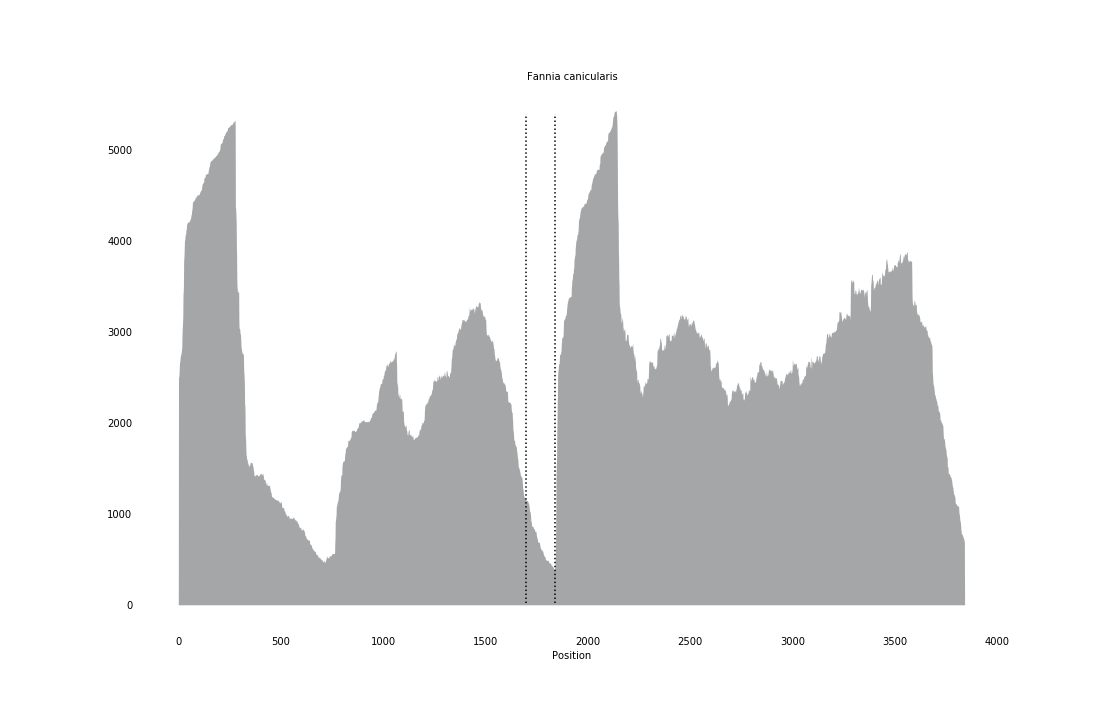

### Forficula_auricularia_coverage_correct.png

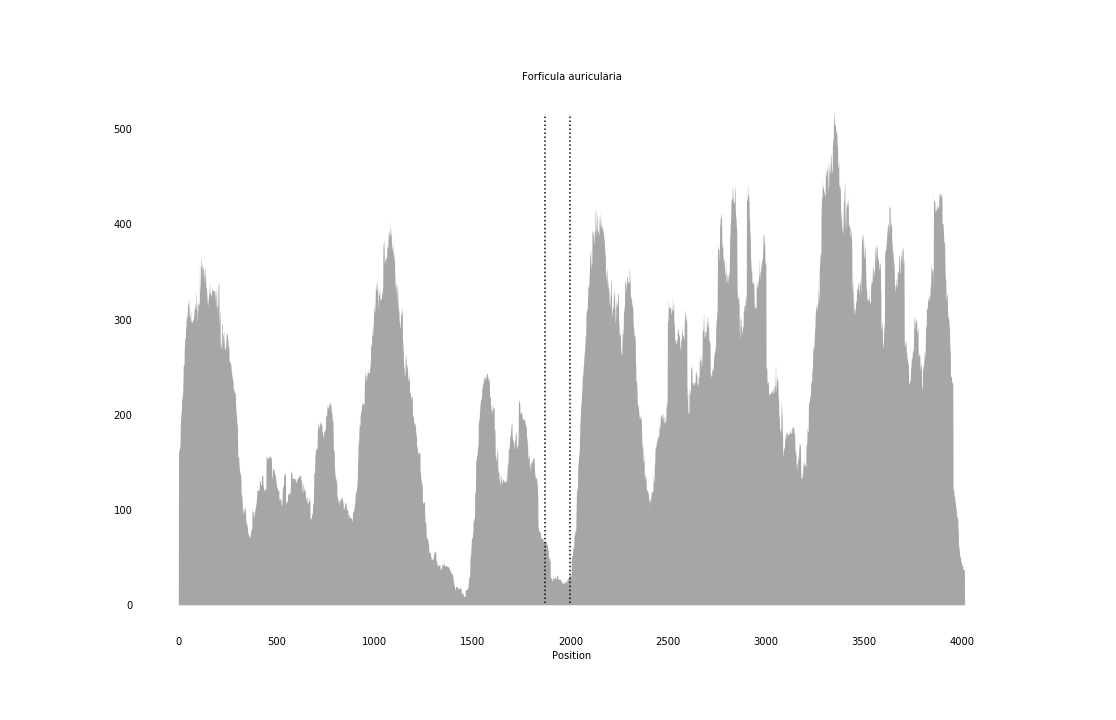

### Galeomma_turtoni_coverage.png

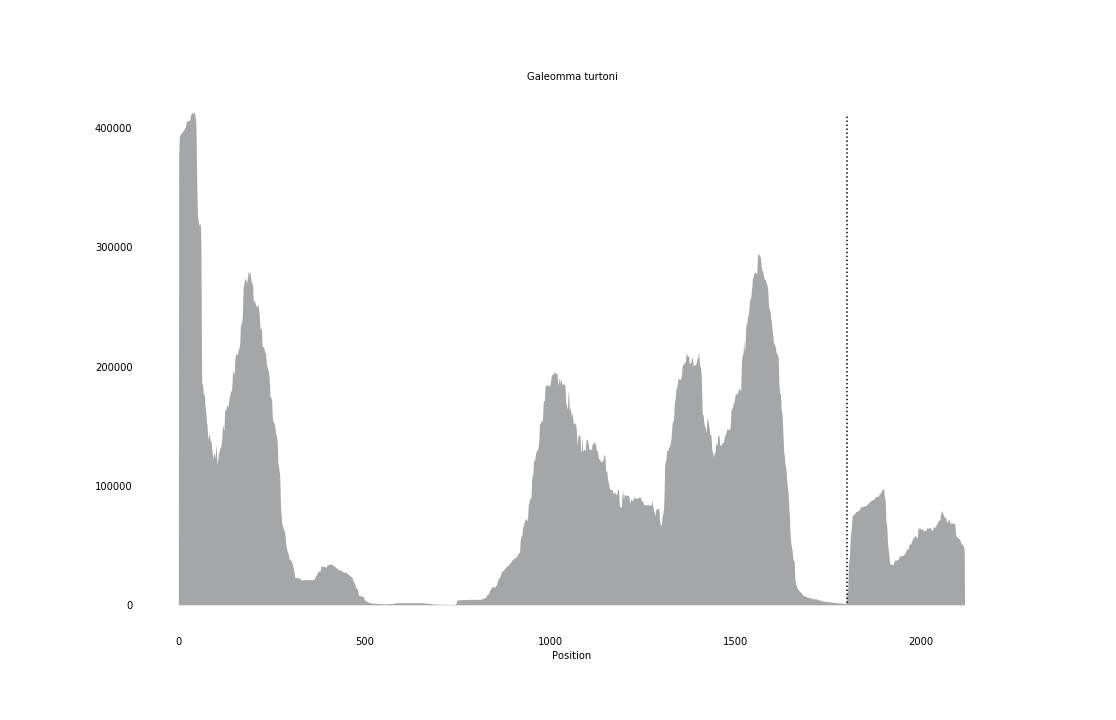

### Galloisiana_yuasai_coverage_correct.png

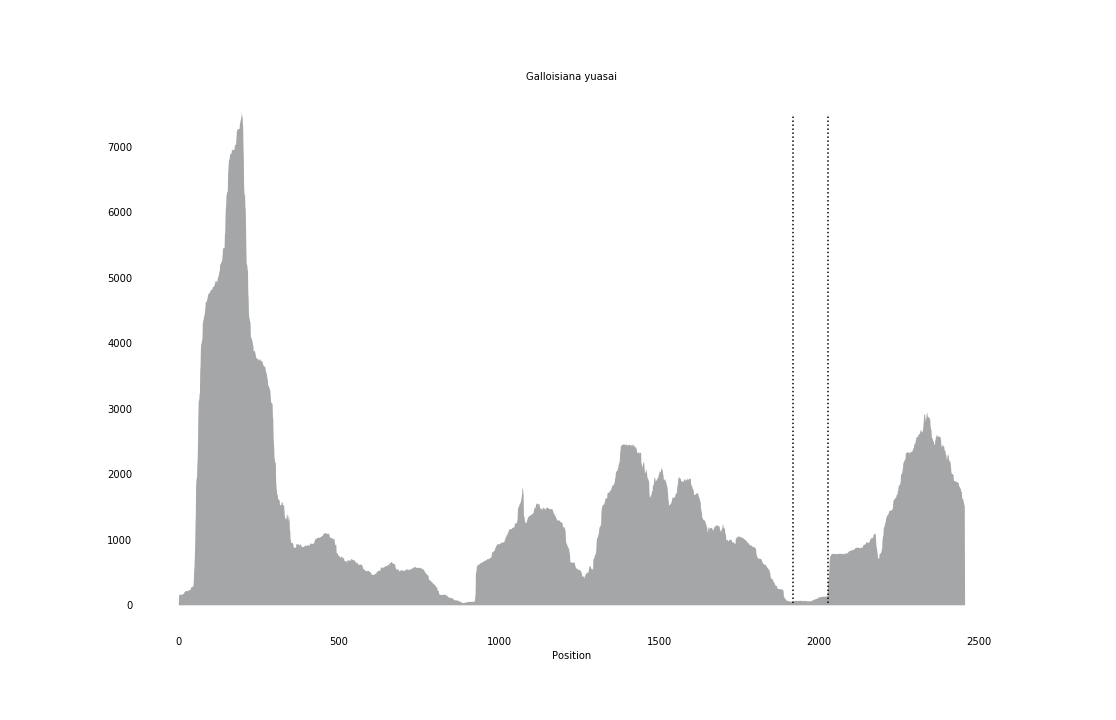

### Glossus_humanus_coverage.png

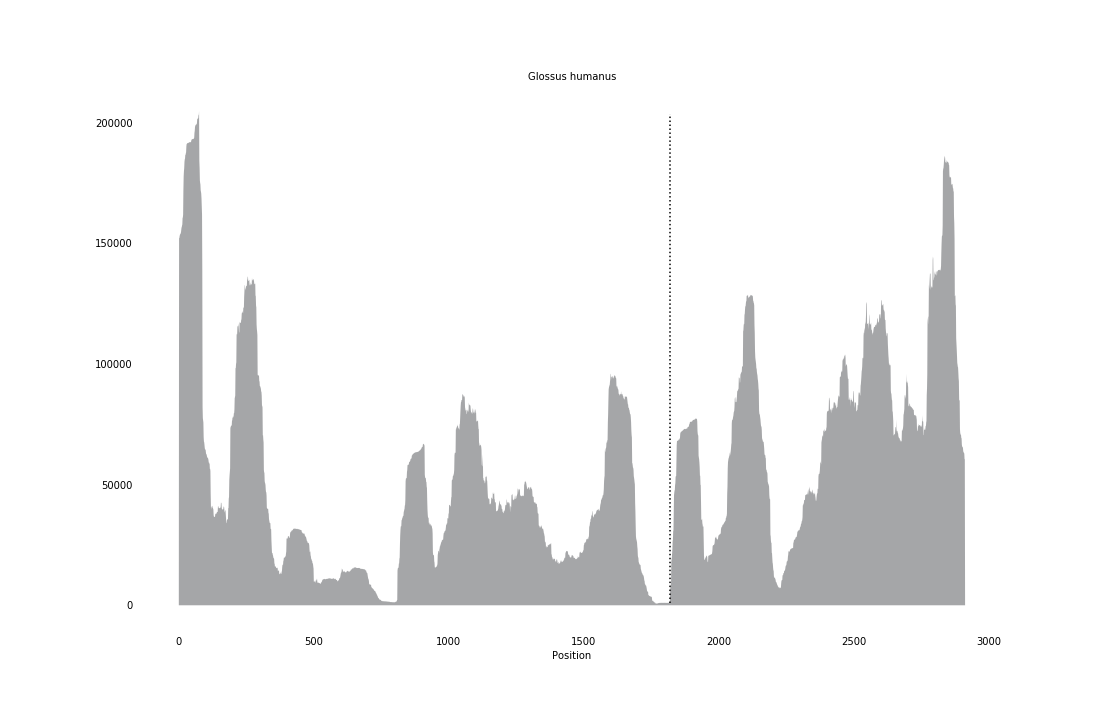

### Gorytes_laticinctus_coverage_correct.png

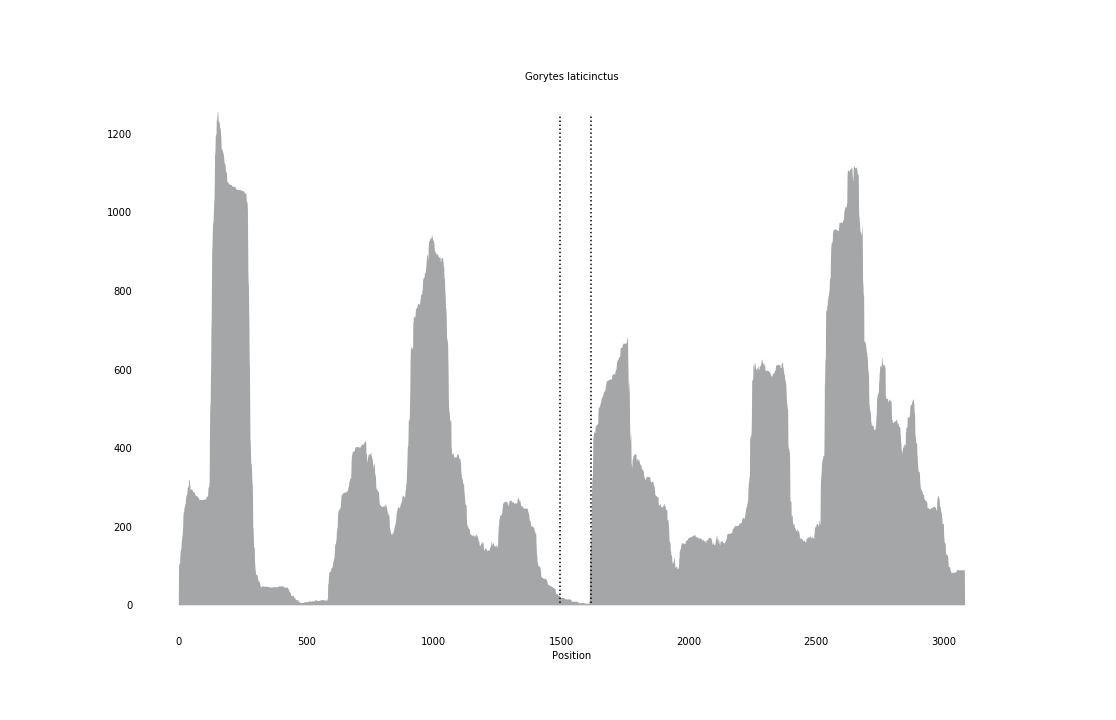

### Gynaikothrips_ficorum_coverage_correct.png

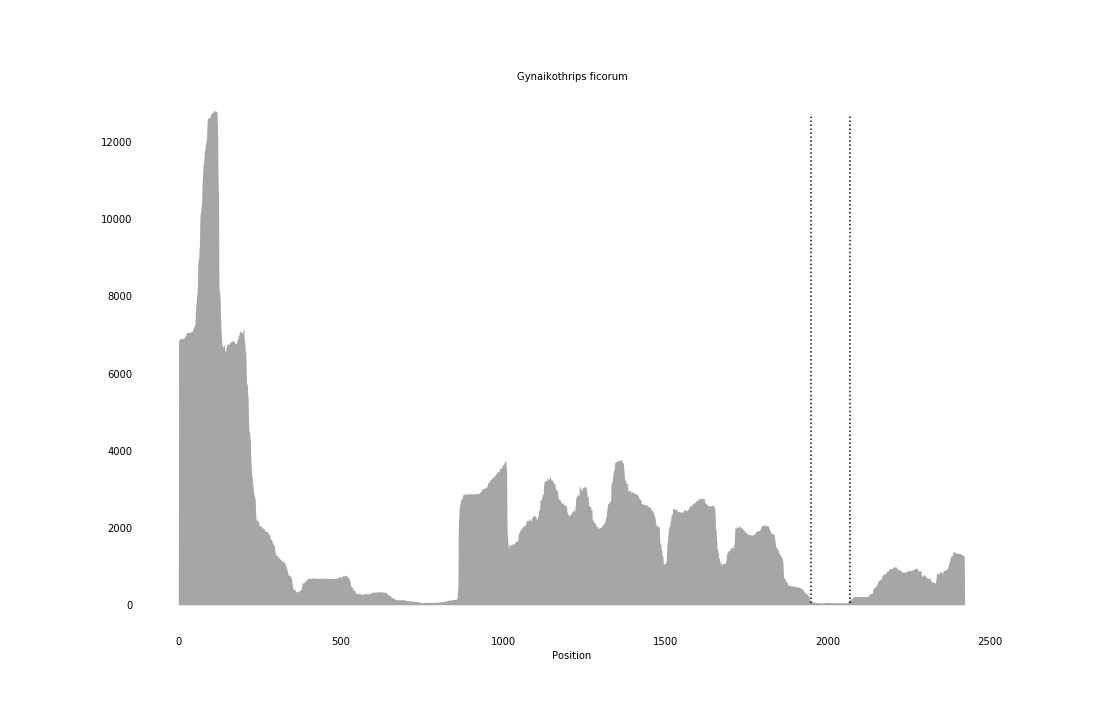

### Heteropelma_amictum_coverage_correct.png

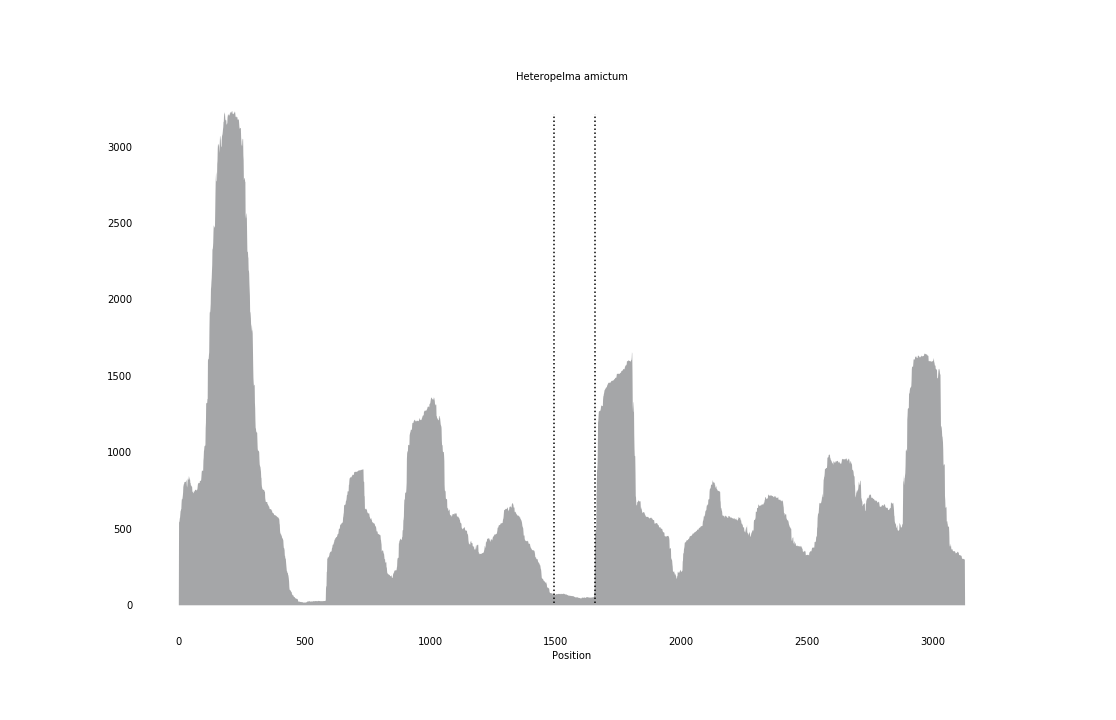

### Hiatella_arctica_coverage.png

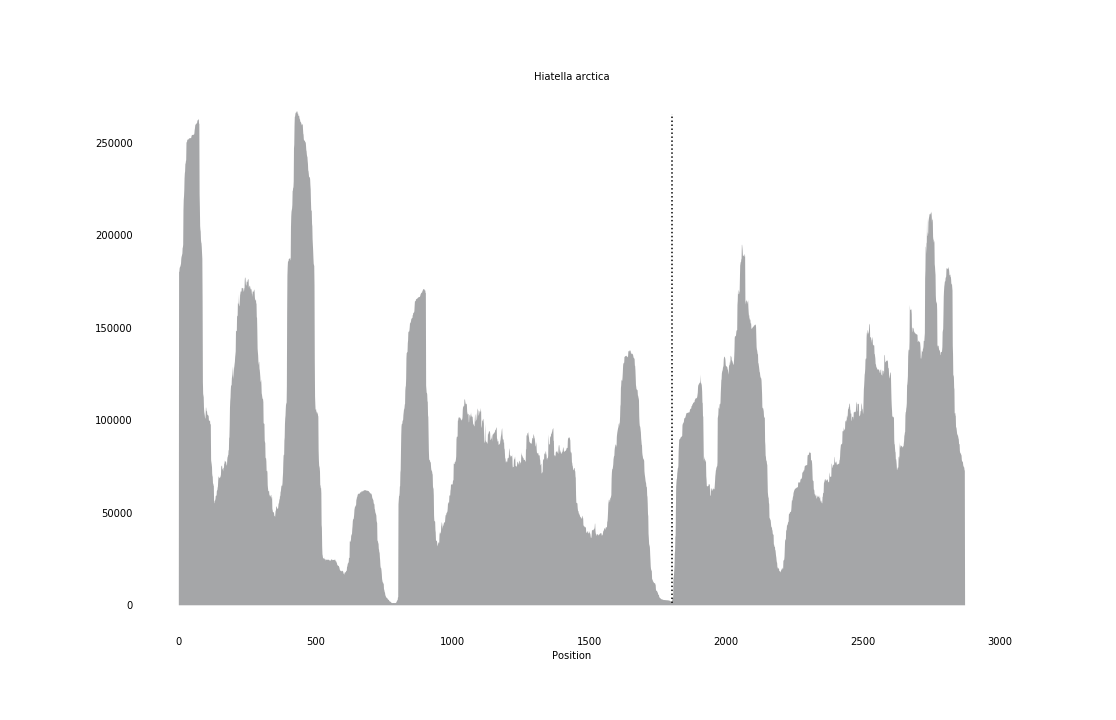

### Isognomon_alatus_coverage.png

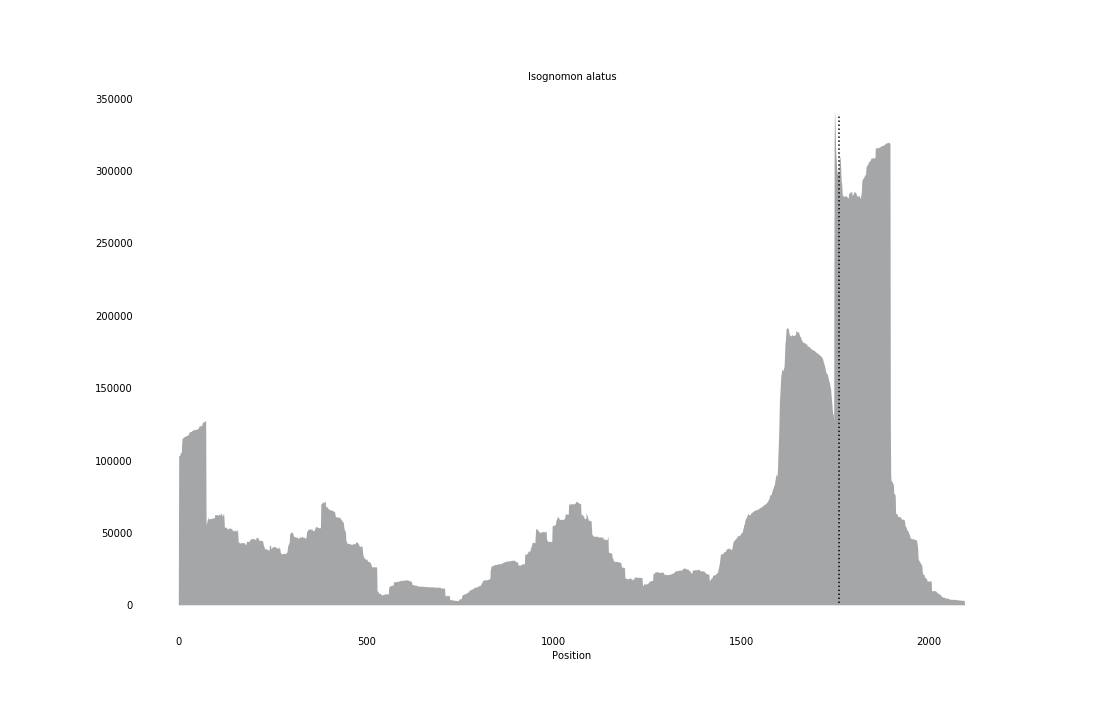

### Lampsilis_cardium_coverage.png

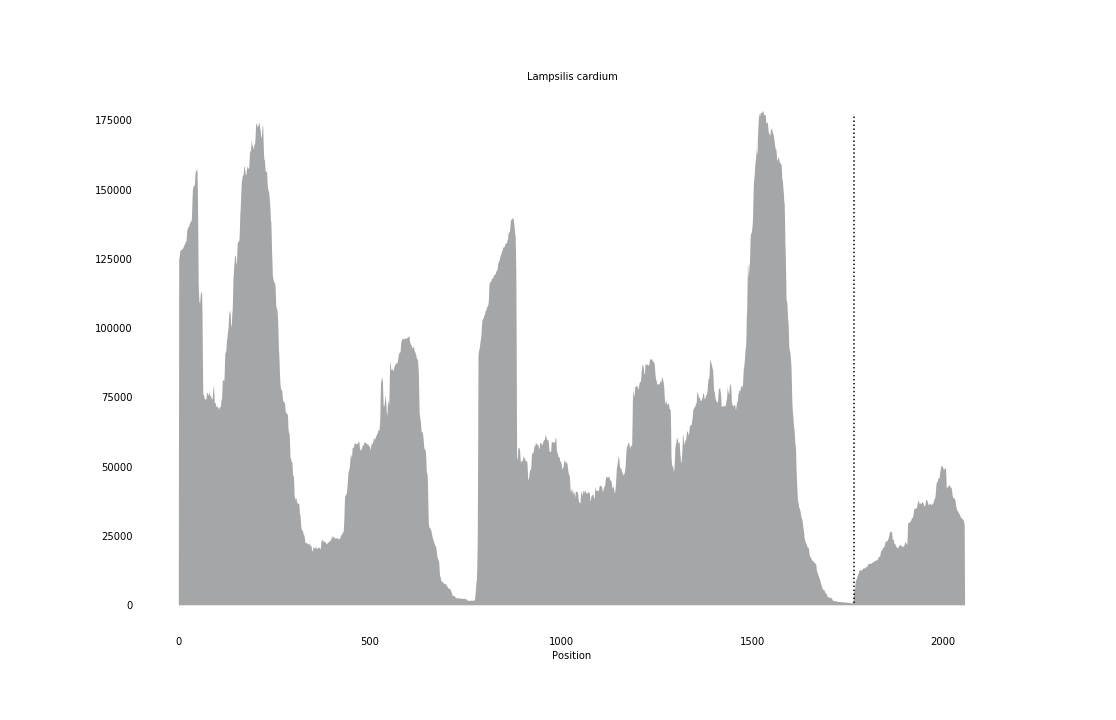

### Lasaea_adansoni_coverage_correct.png

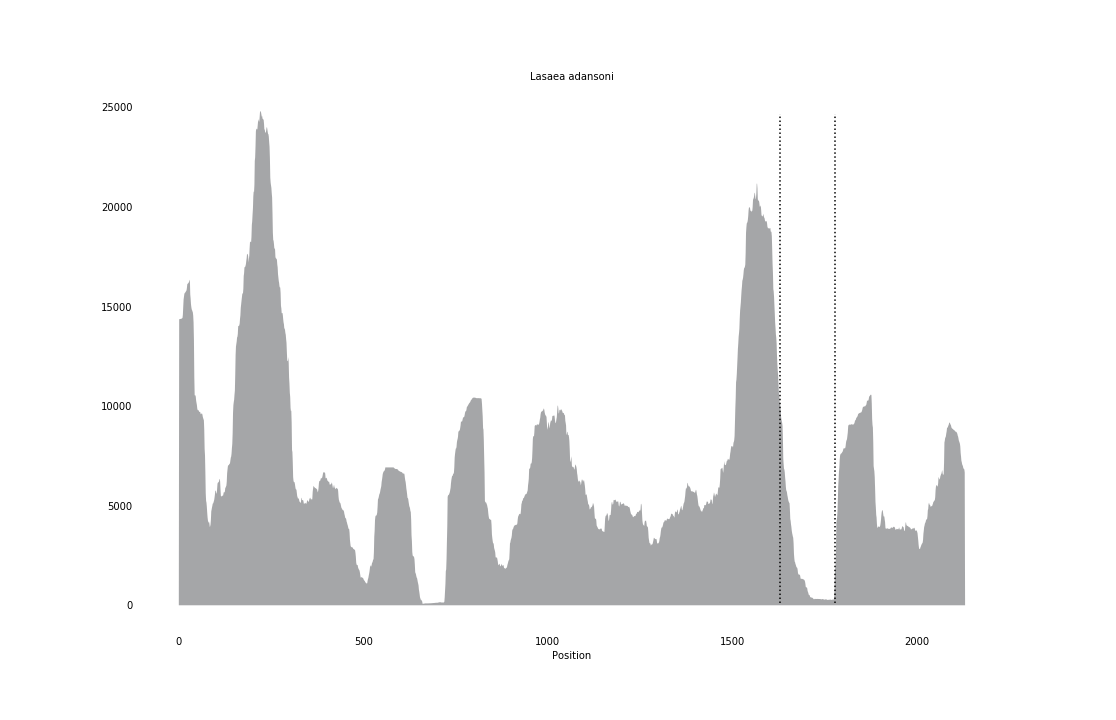

### Lasioglossum_xanthopus_coverage_correct.png

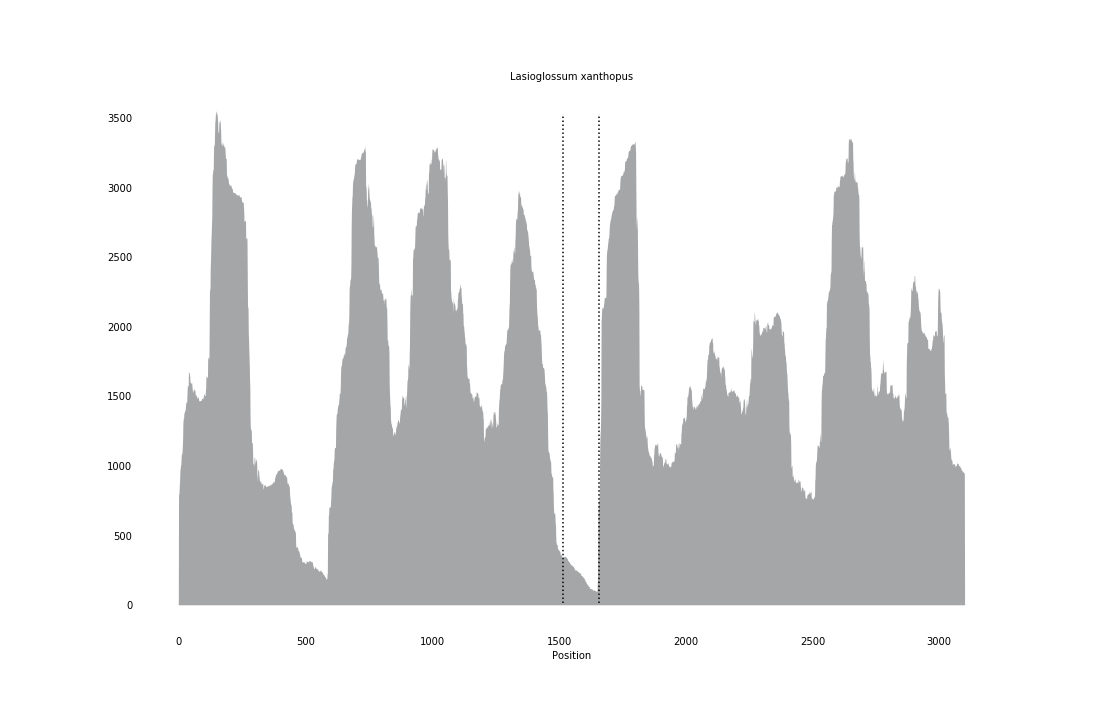

### Laternula_elliptica_coverage.png

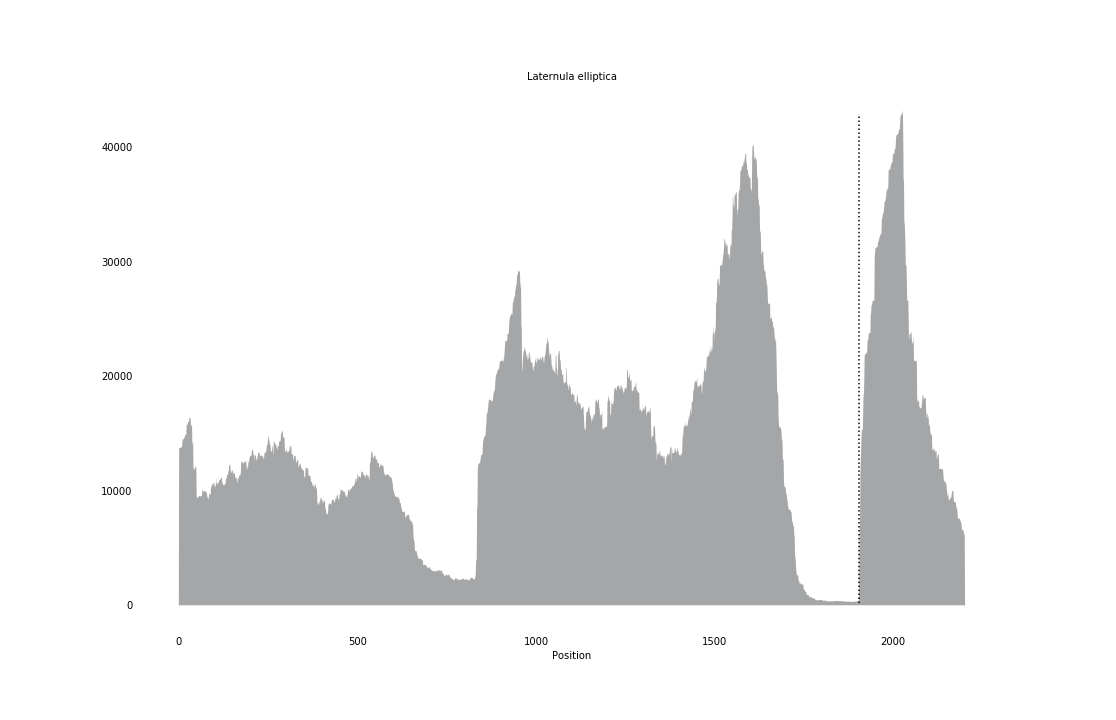

### Leptopilina_clavipes_coverage_correct.png

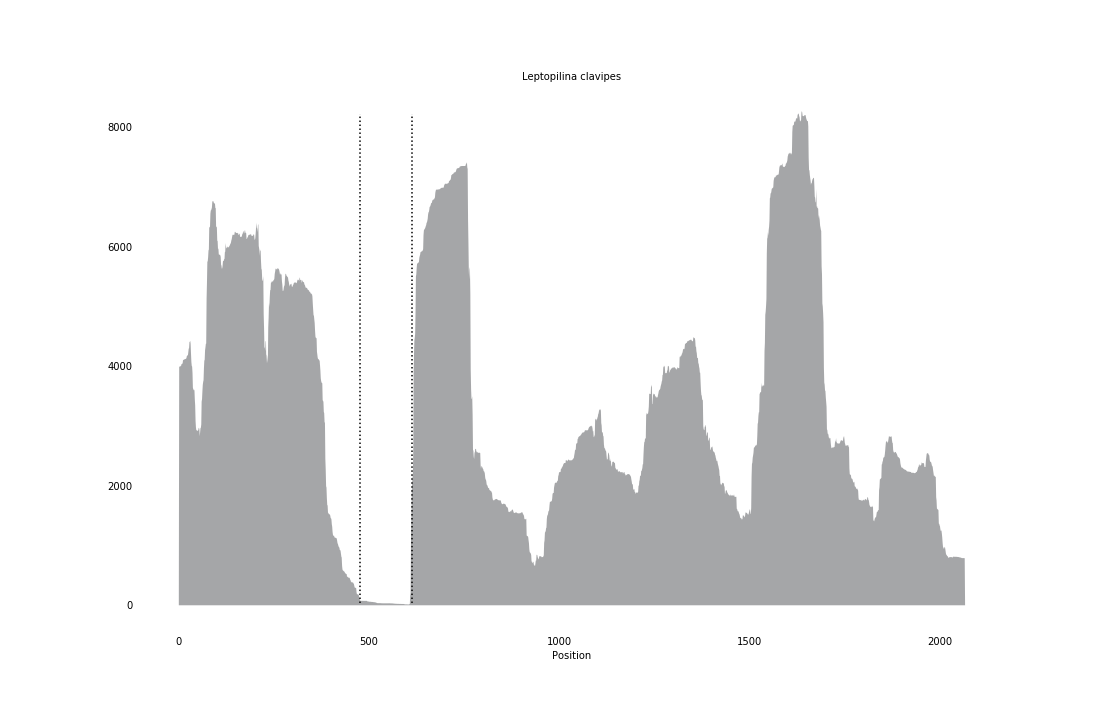

### Macrostomum_lignano_coverage_correct.png

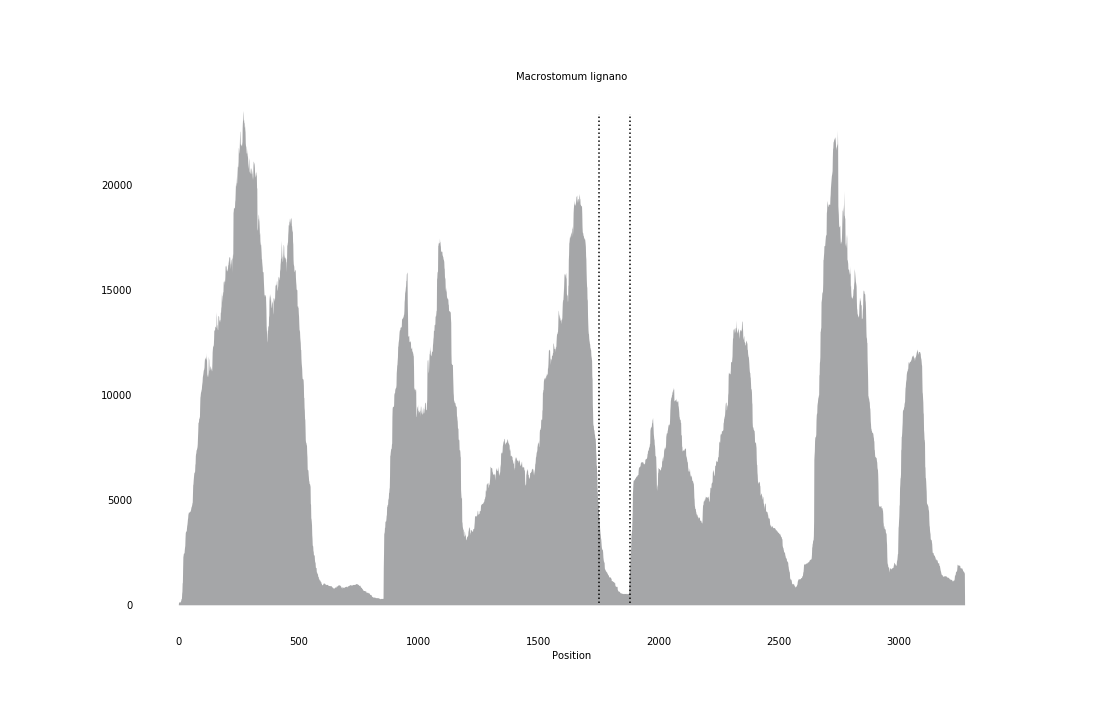

### Macrostomum_tuba_coverage_correct.png

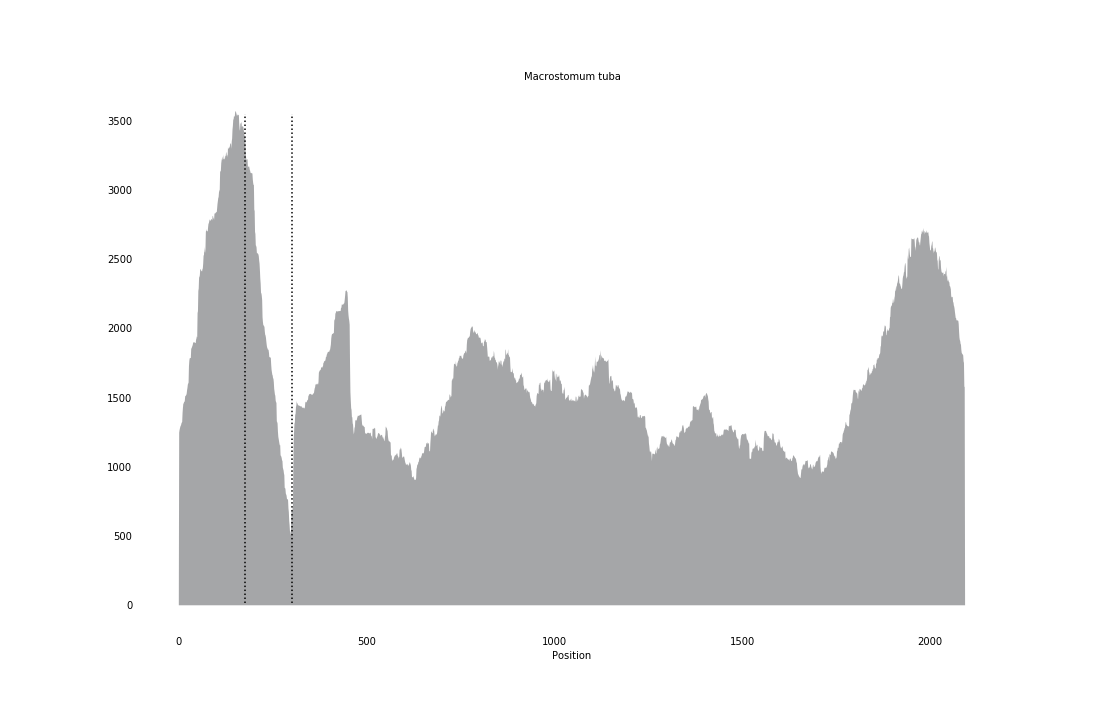

### Mantis_religiosa_coverage_correct.png

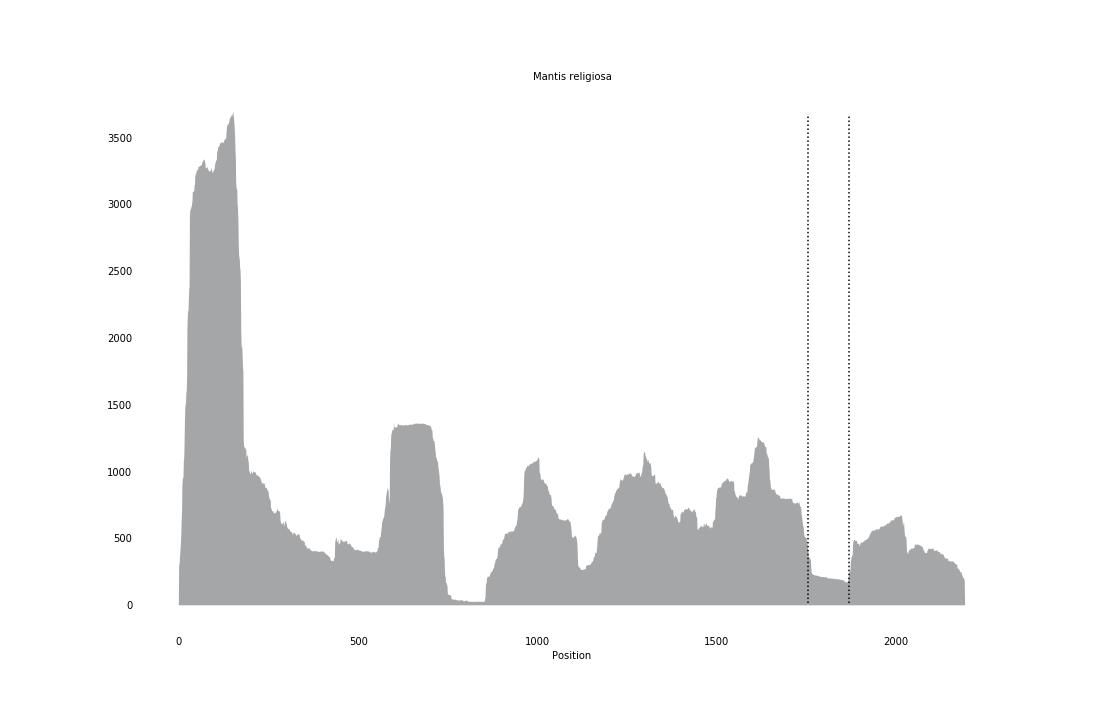

### Melipona_quadrifasciata_coverage_correct.png

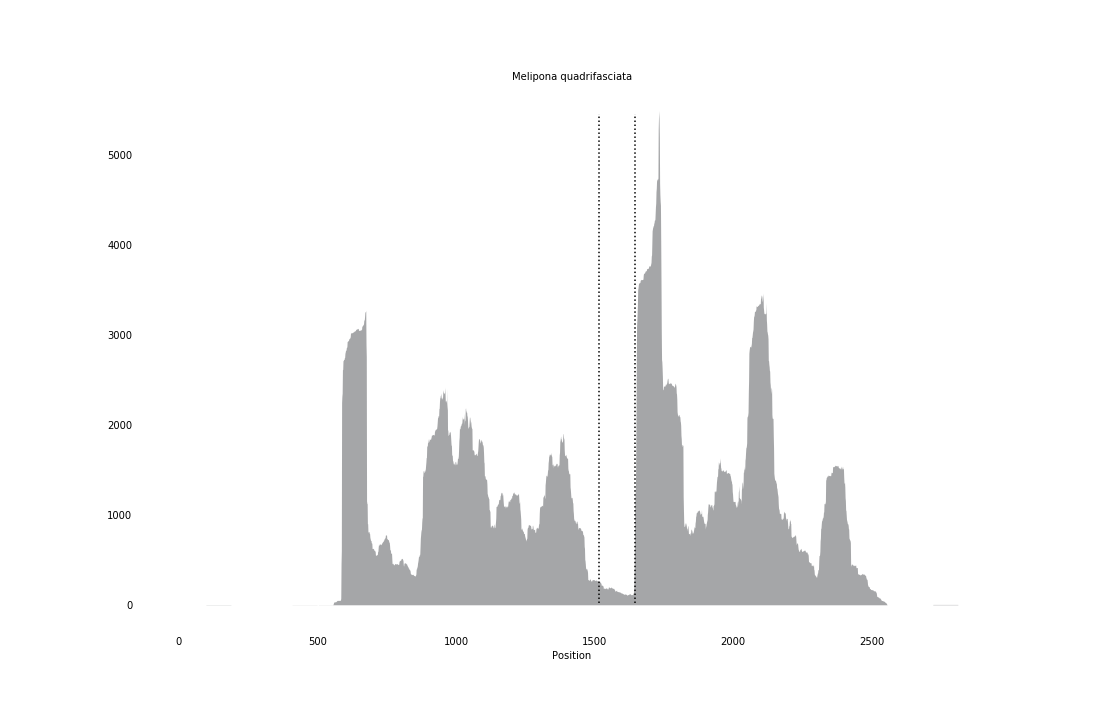

### Melitta_haemorrhoidalis_coverage_correct.png

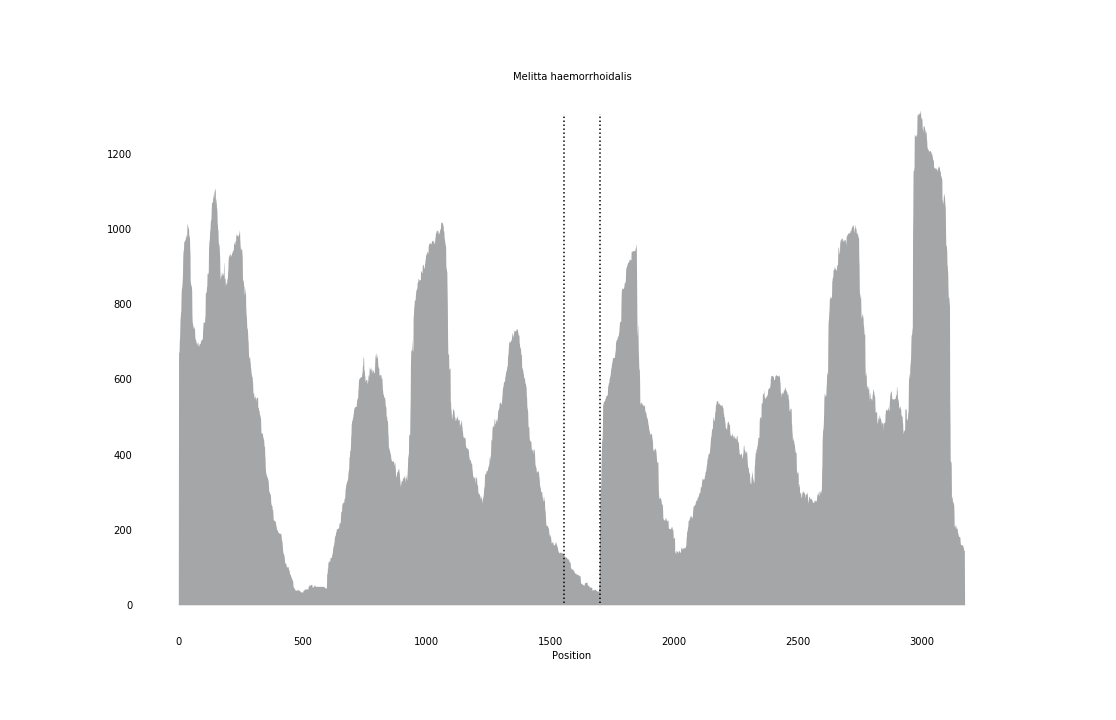

### Microstomum_lineare_coverage.png

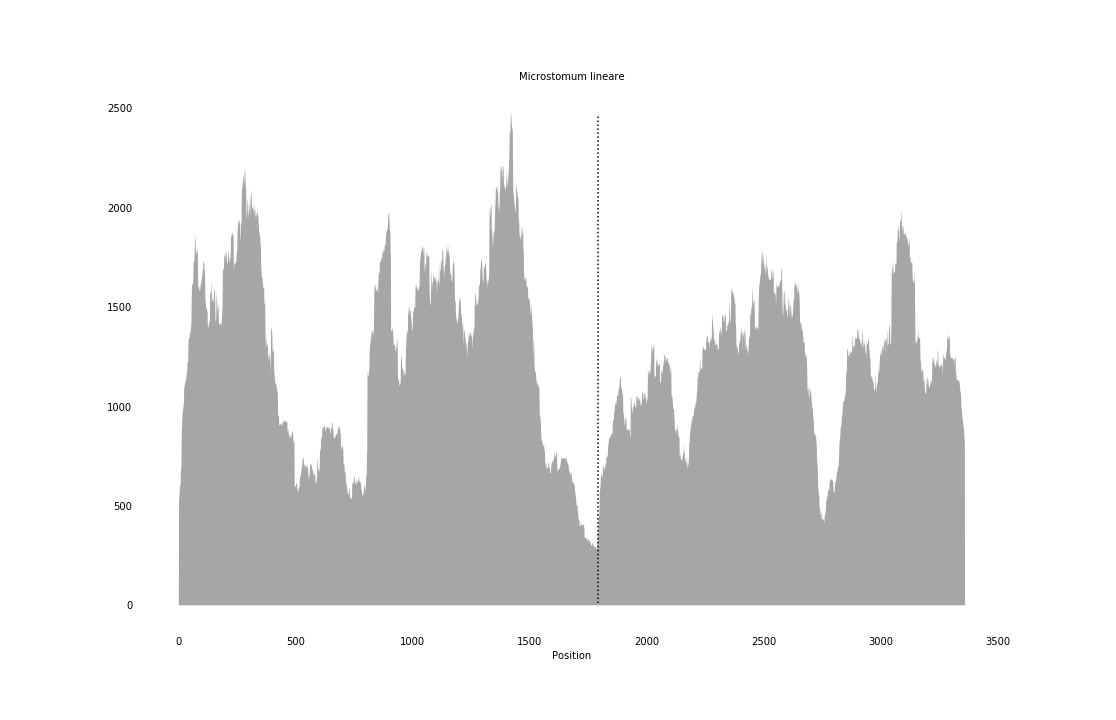

### Monocelis_fusca_coverage.png

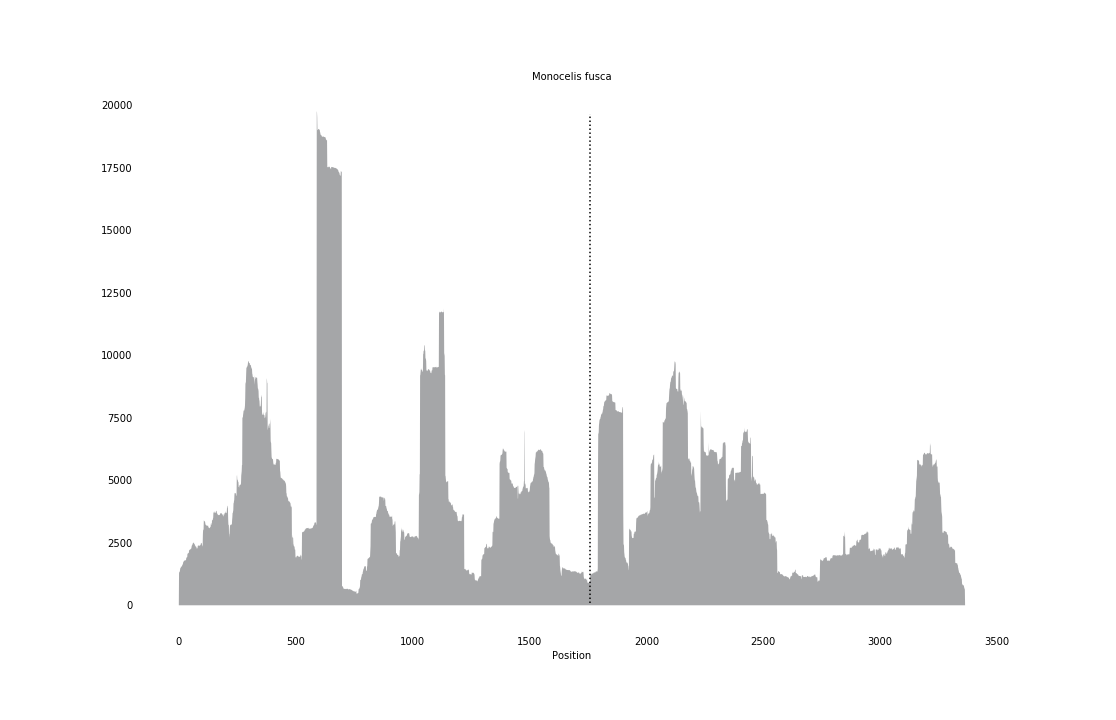

### Monosapyga_clavicornis_coverage_correct.png

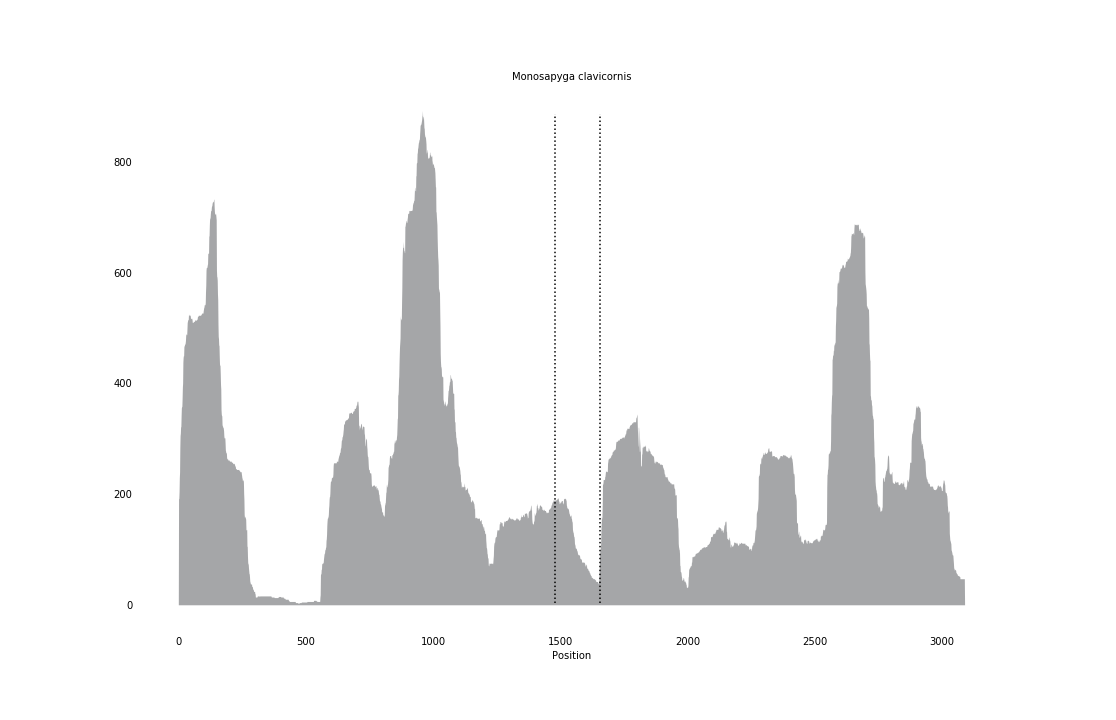

### Muscina_stabulans_coverage_correct.png

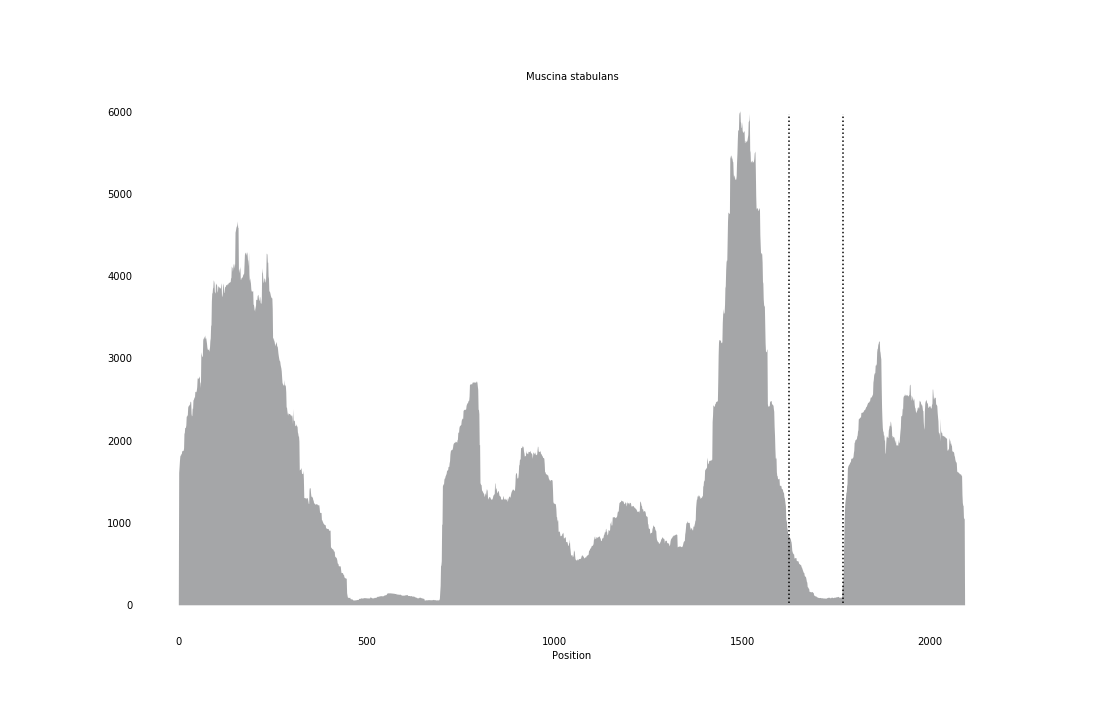

### Nasonia_vitripennis_coverage_correct.png

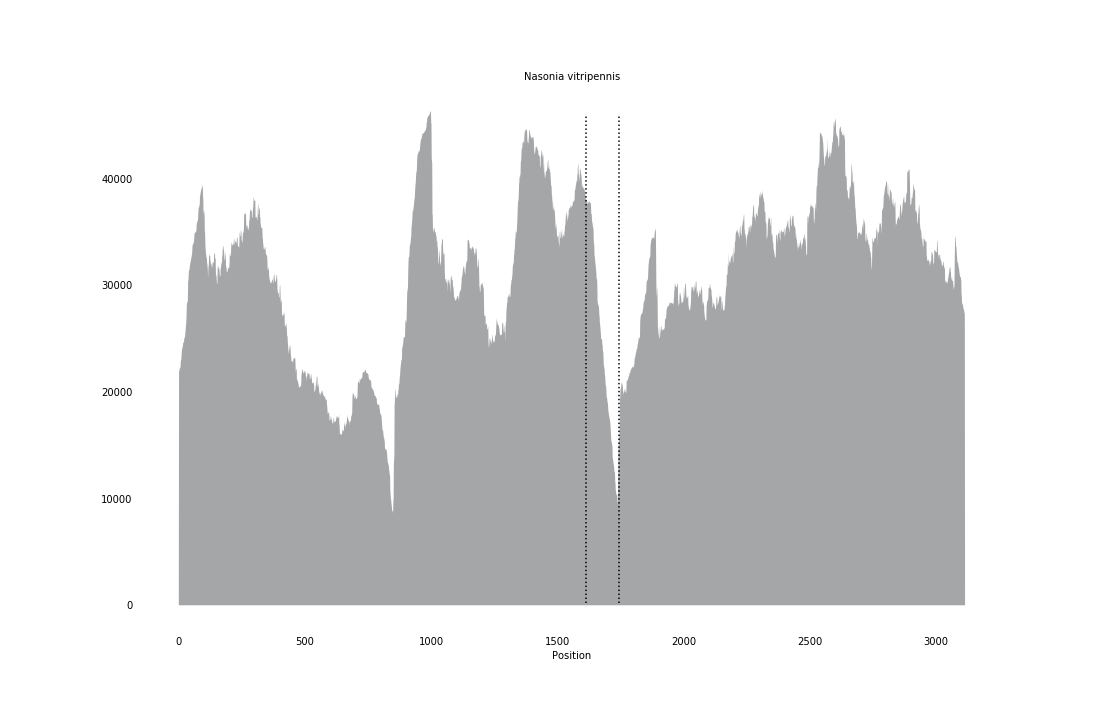

### Neobellieria_bullata_coverage.png

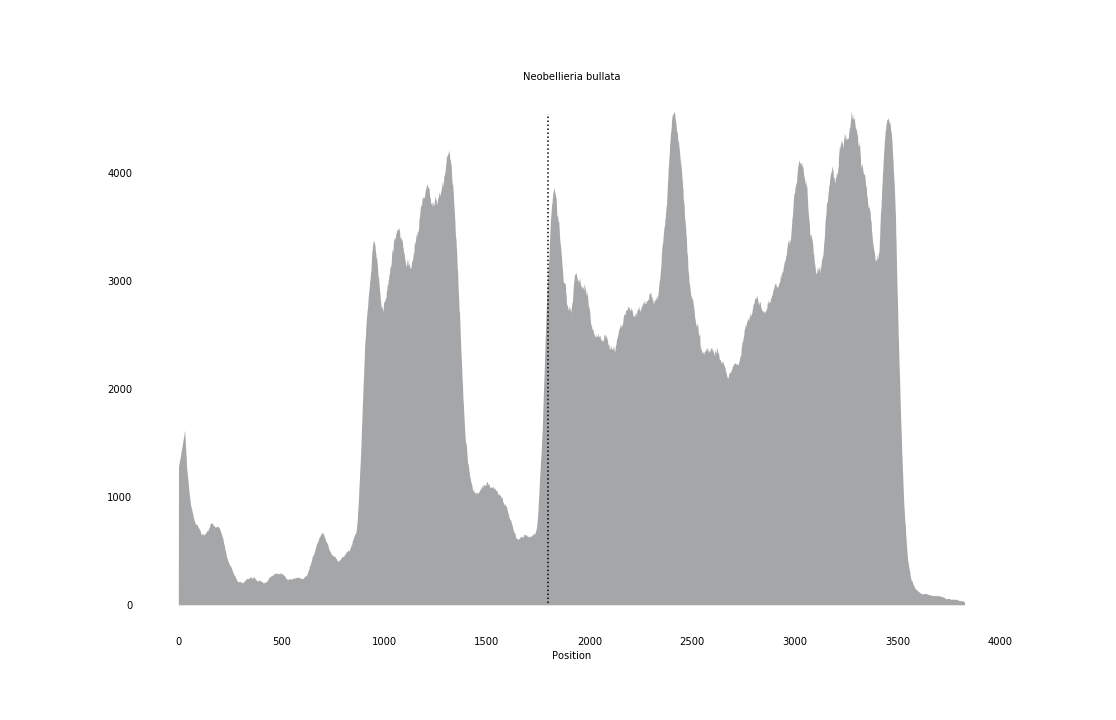

### Neotrigonia_margaritacea_coverage_correct.png

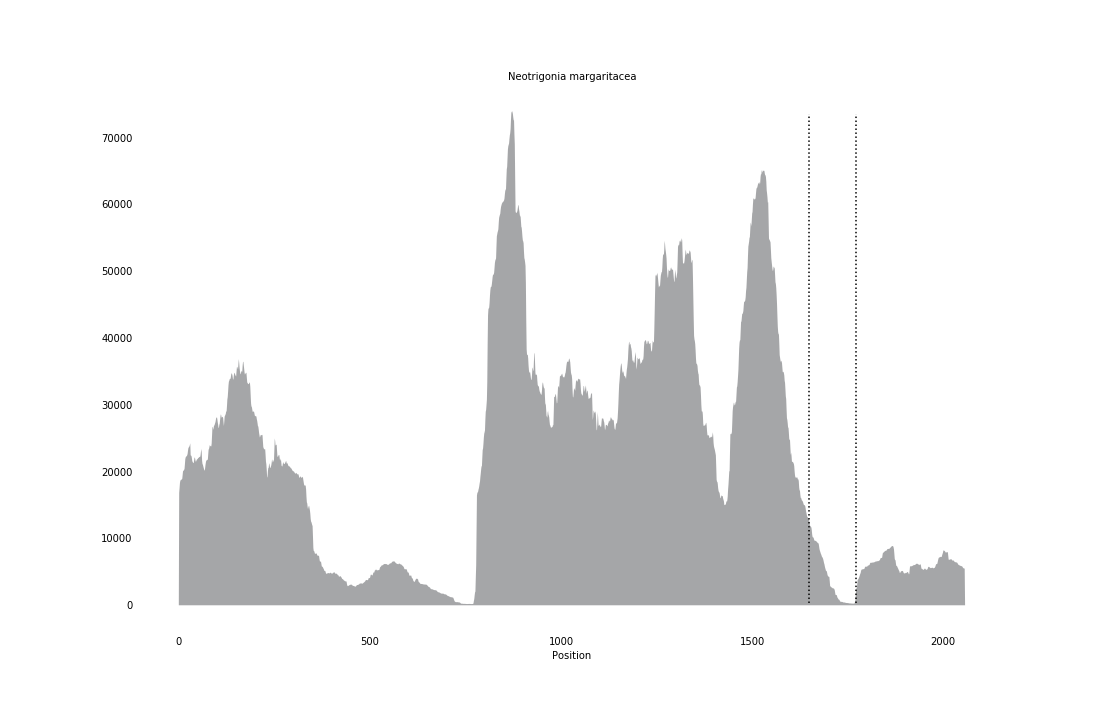

### Nerita_peloronta_coverage_correct.png

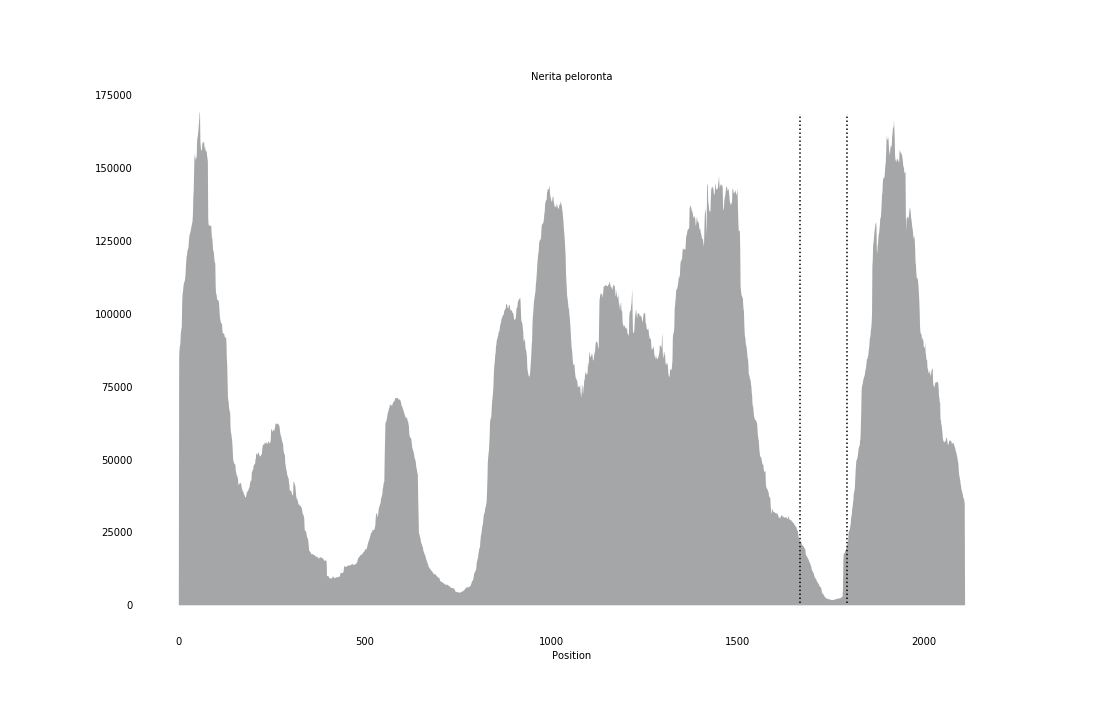

### Nysson_niger_coverage_correct.png

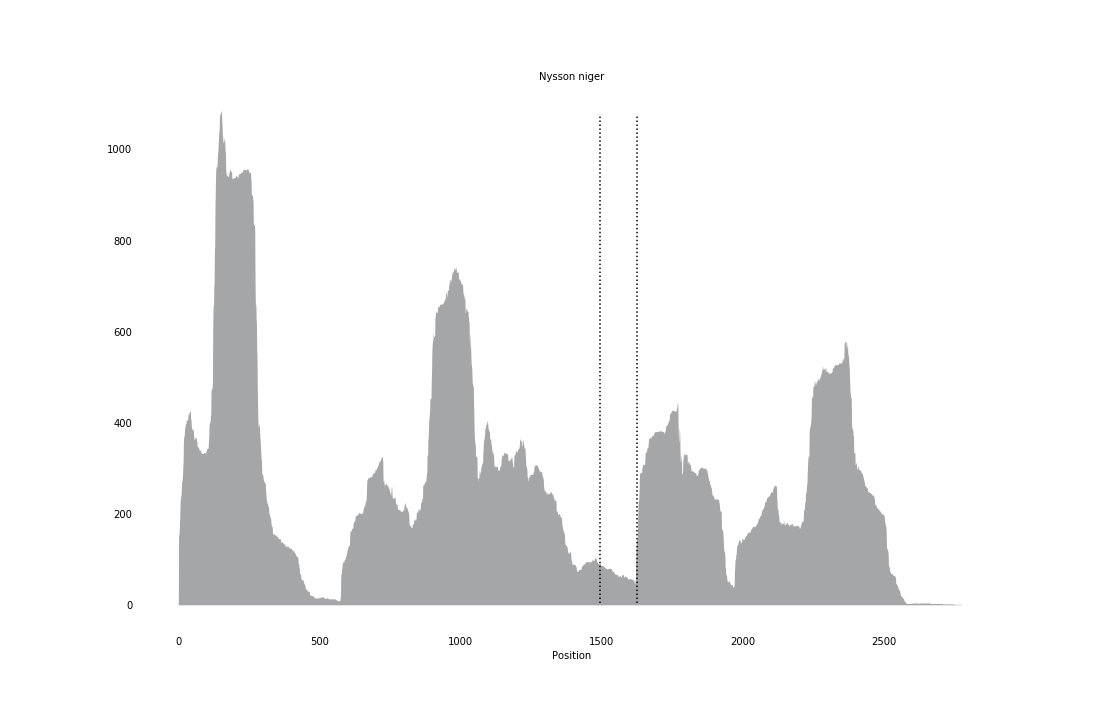

### Panopea_globosa_coverage.png

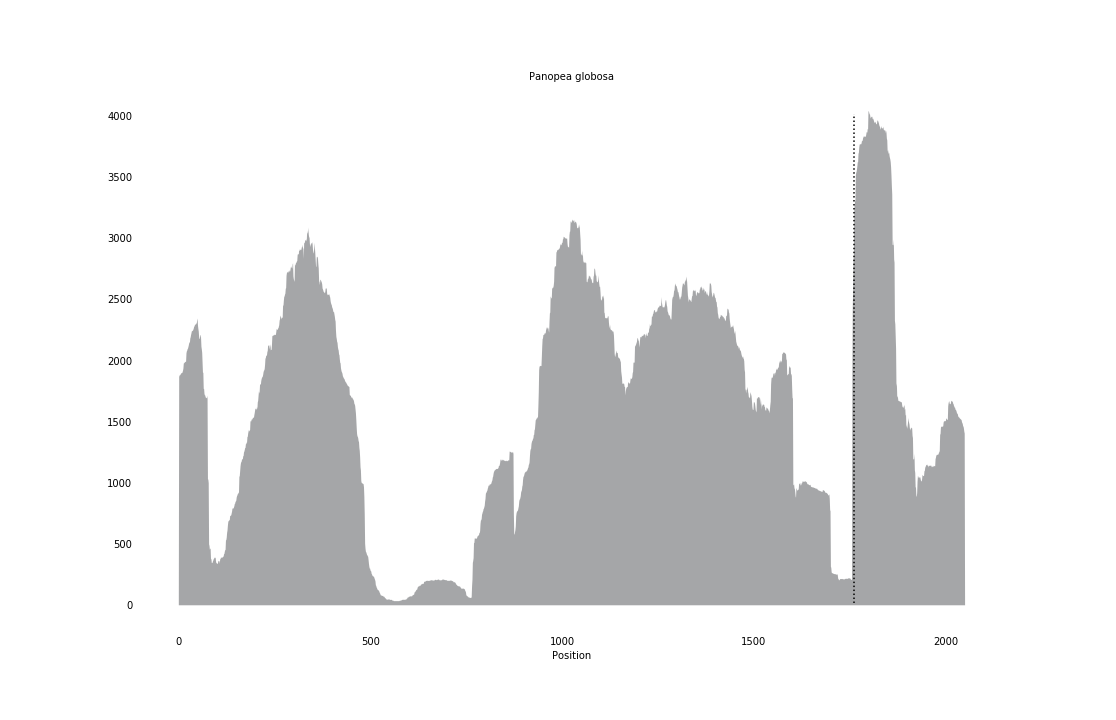

### Pelecinus_polyturator_coverage_correct.png

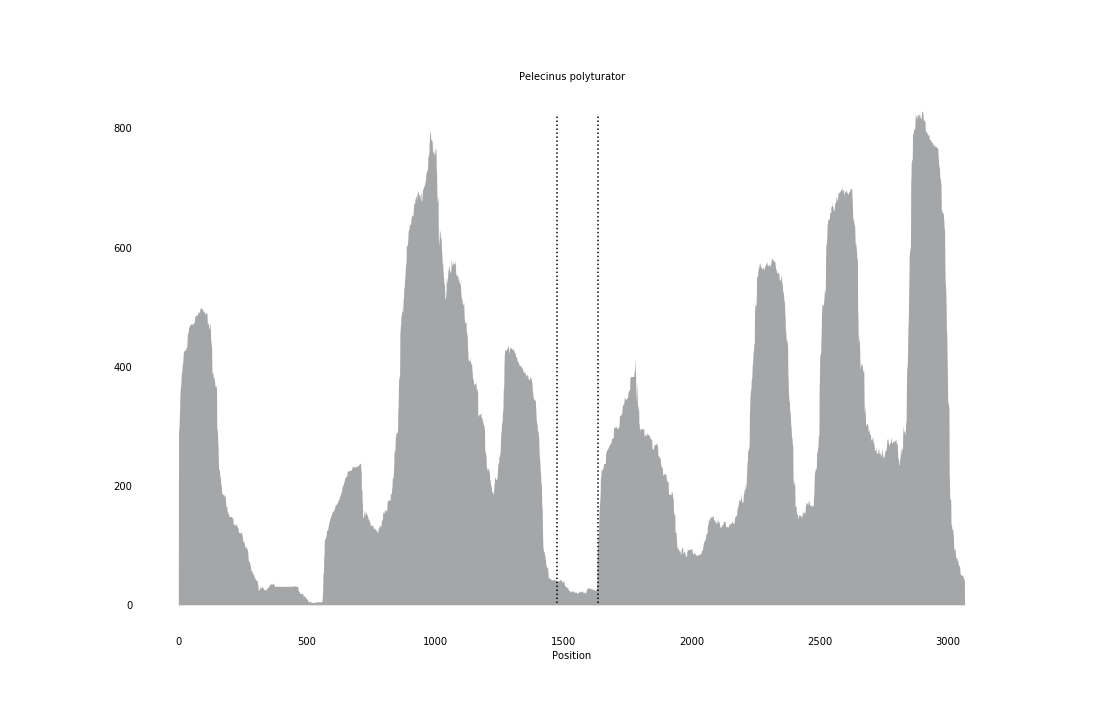

### Phagocata_velata_coverage_correct.png

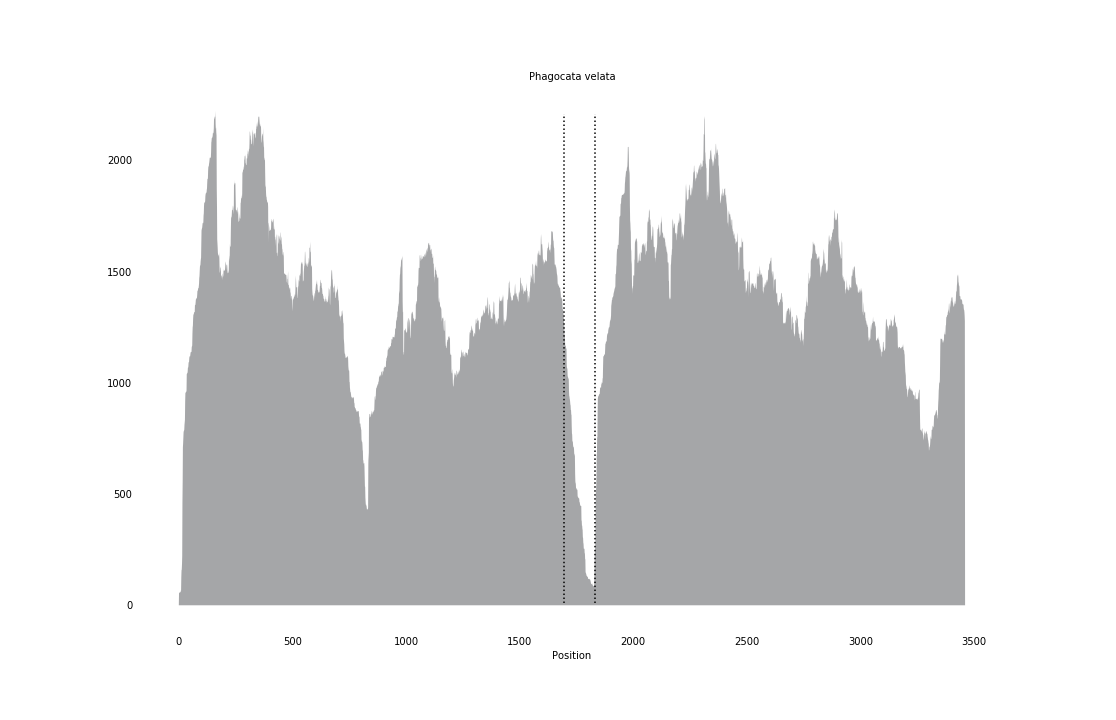

### Phallomedusa_solida_coverage_correct.png

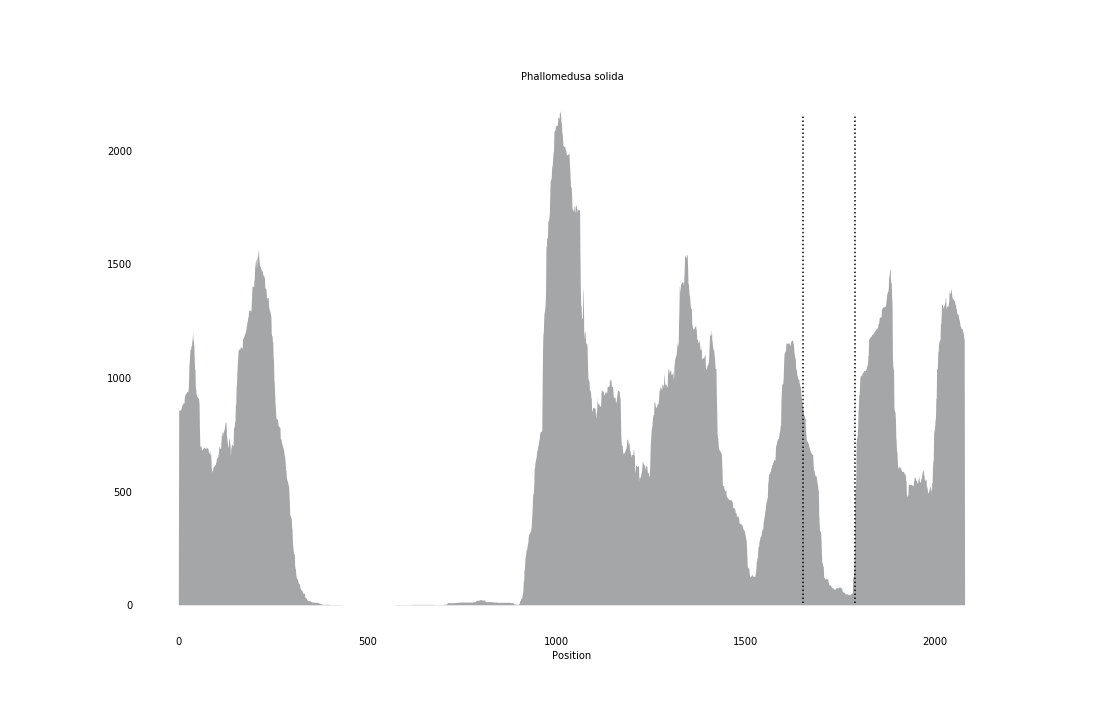

### Pinctada_fucata_coverage_correct.png

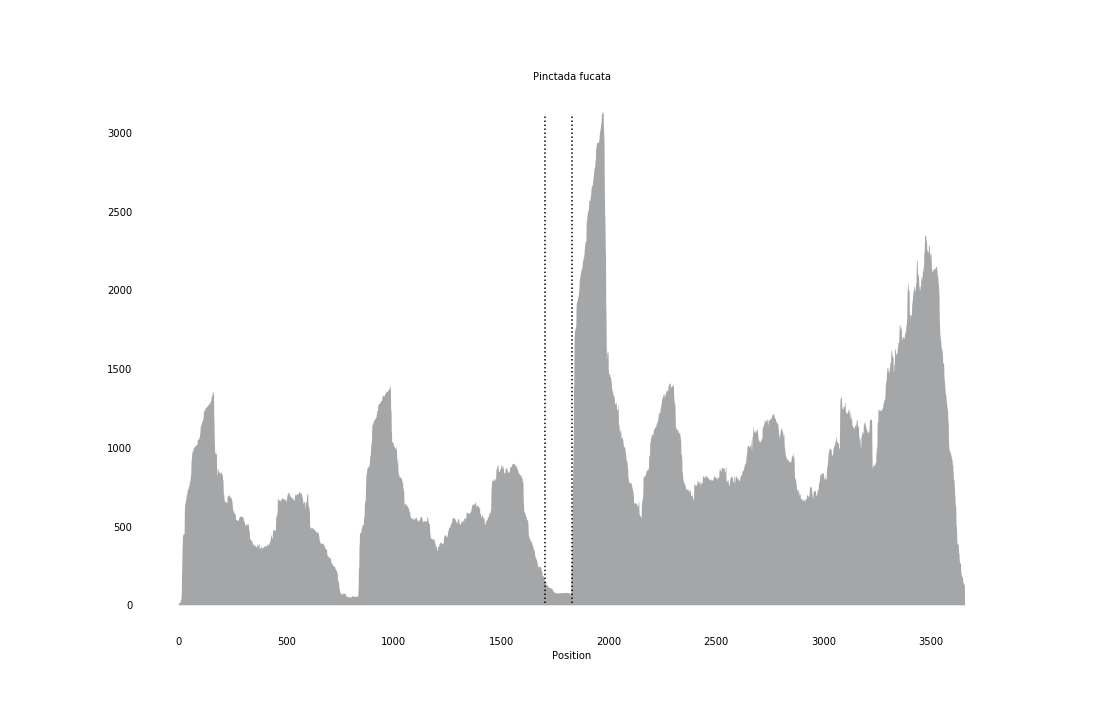

### Pinna_atropurpurea_coverage.png

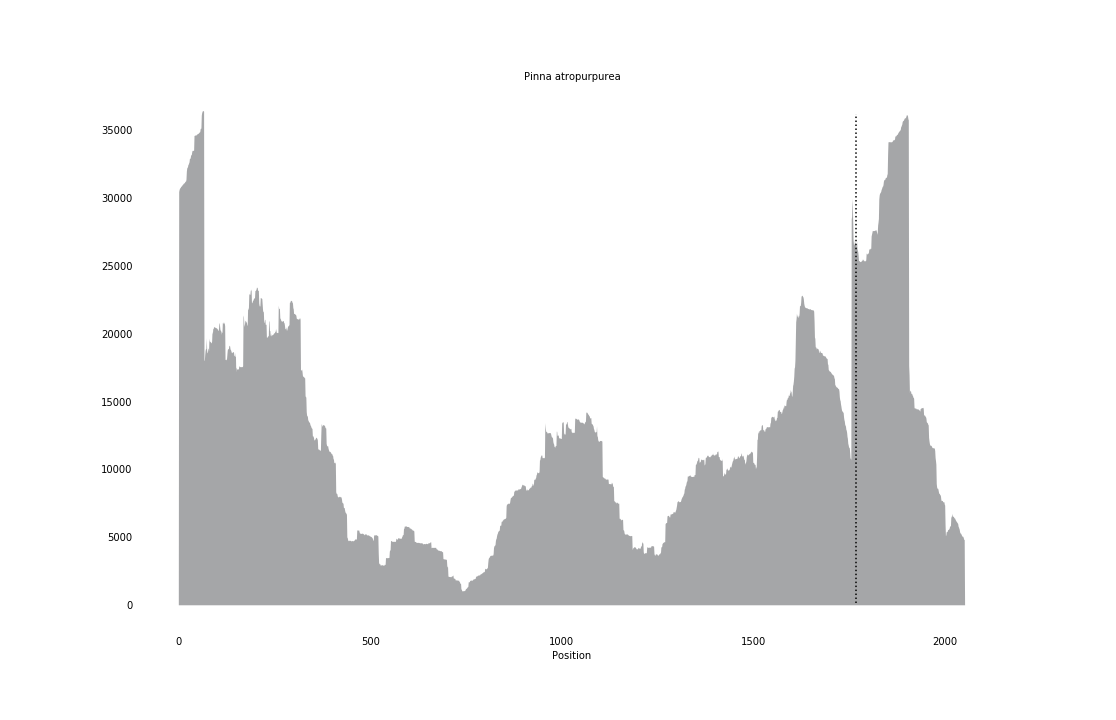

### Pinna_saccata_coverage_correct.png

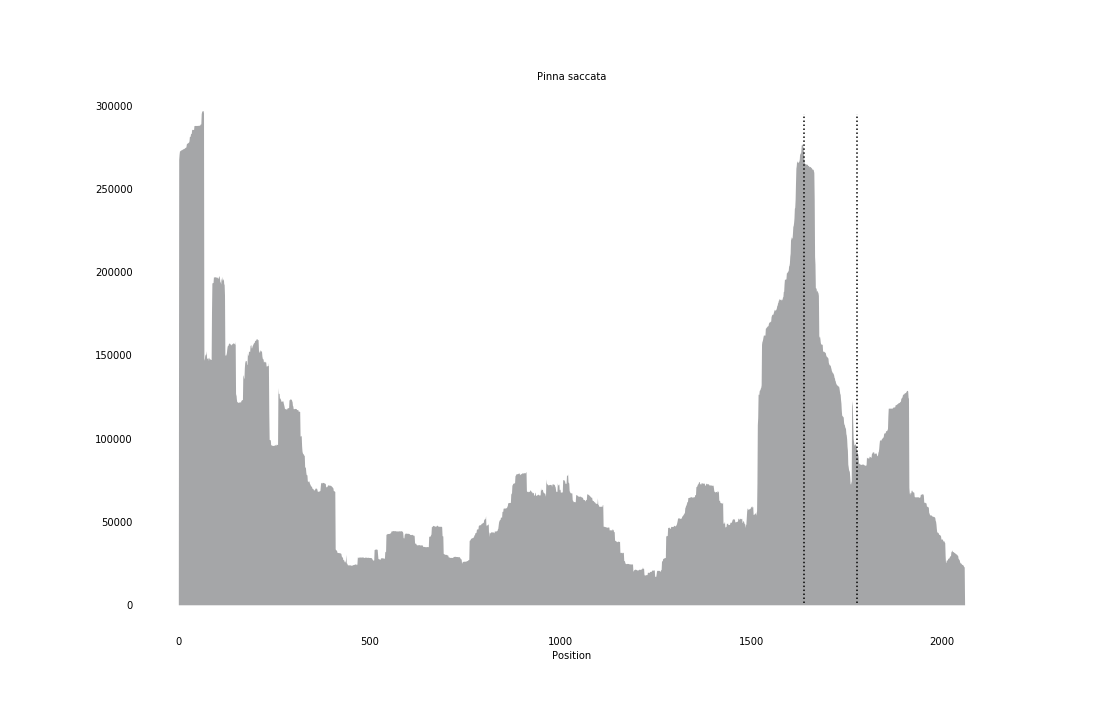

### Placopecten_magellanicus_coverage_correct.png

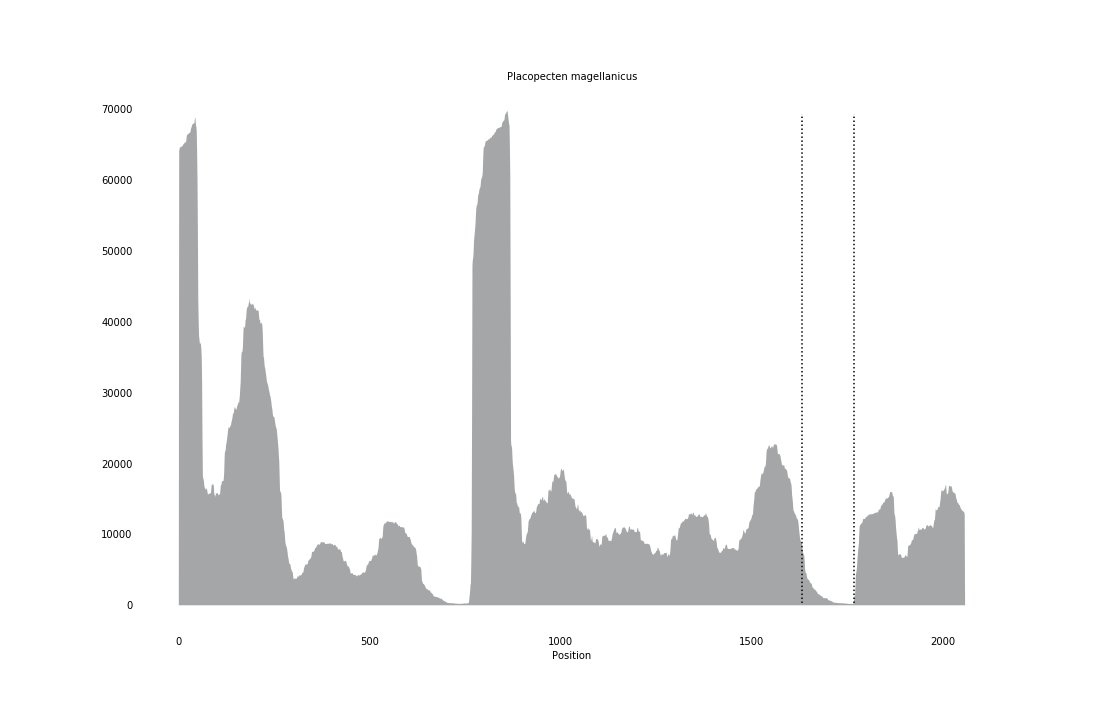

### Prorhinotermes_simplex_coverage_correct.png

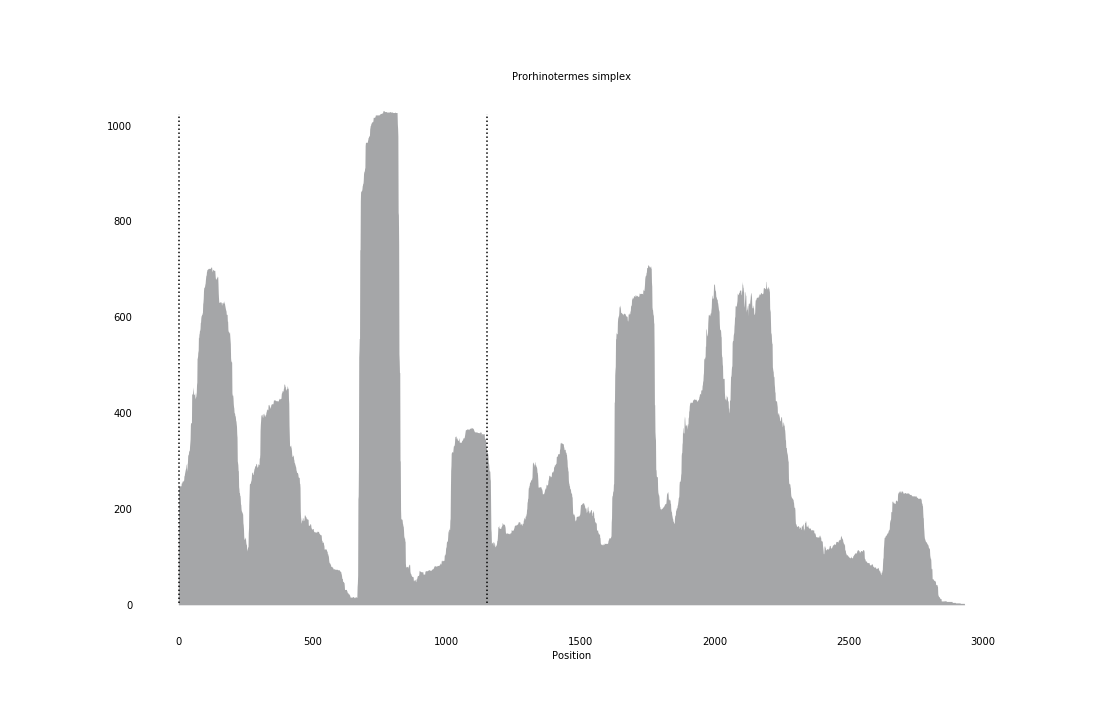

### Prorhynchus_alpinus_coverage.png

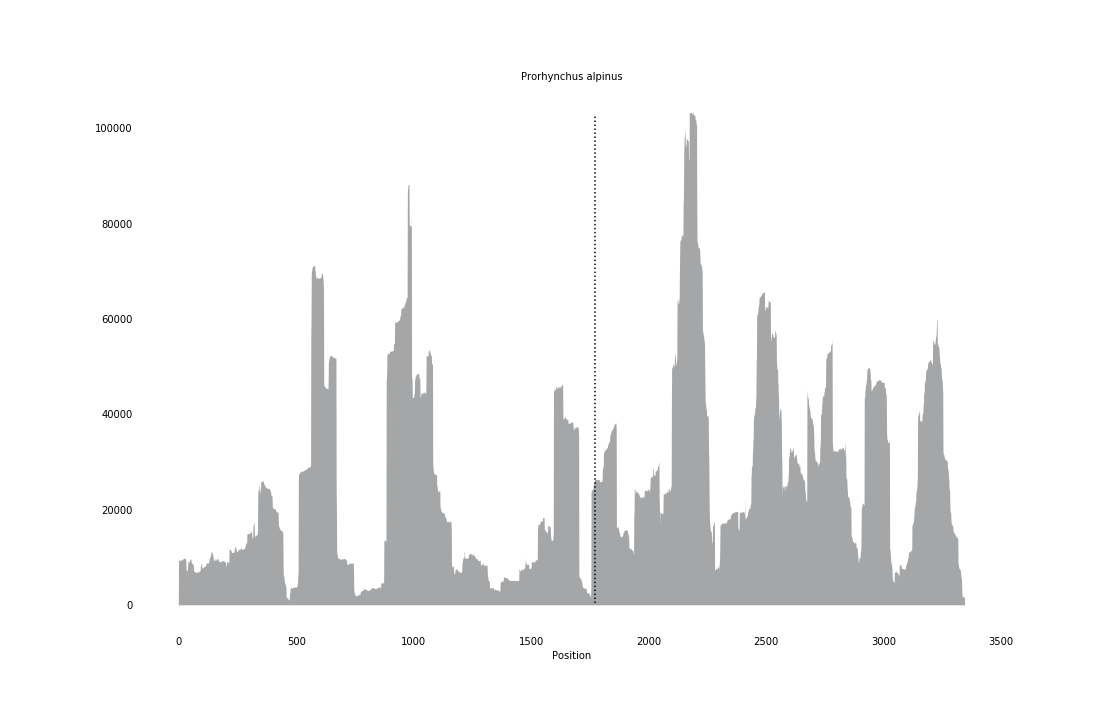

### Prorhynchus_stagnalis_coverage.png

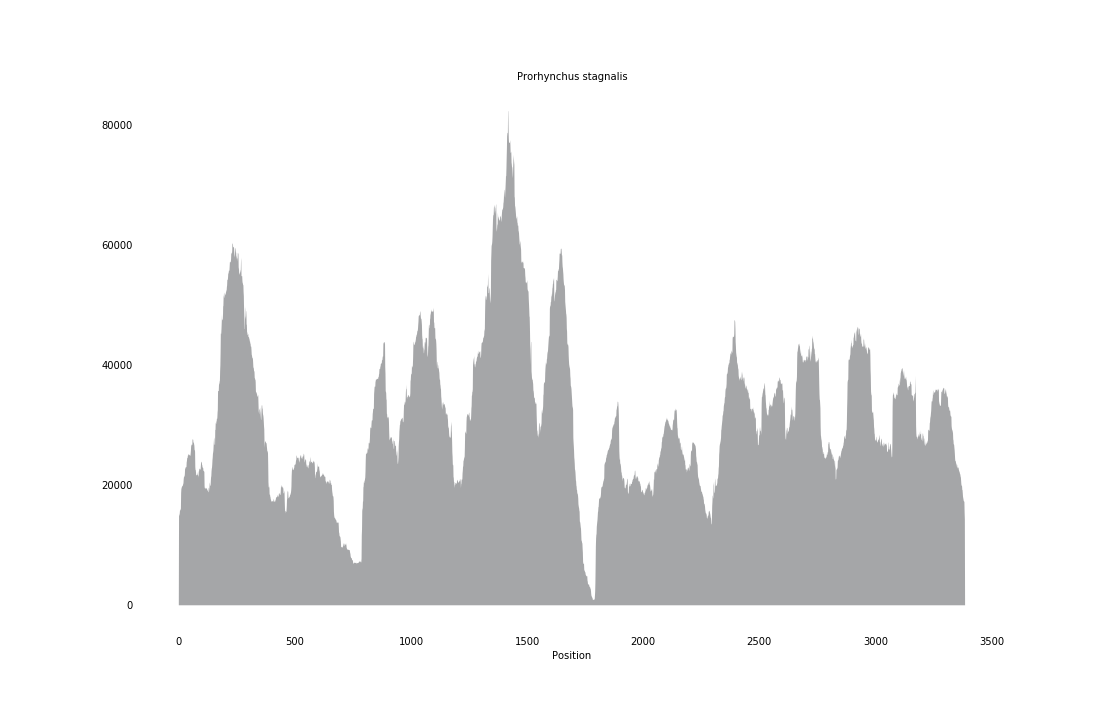

### Prosapia_bicincta_coverage_correct.png

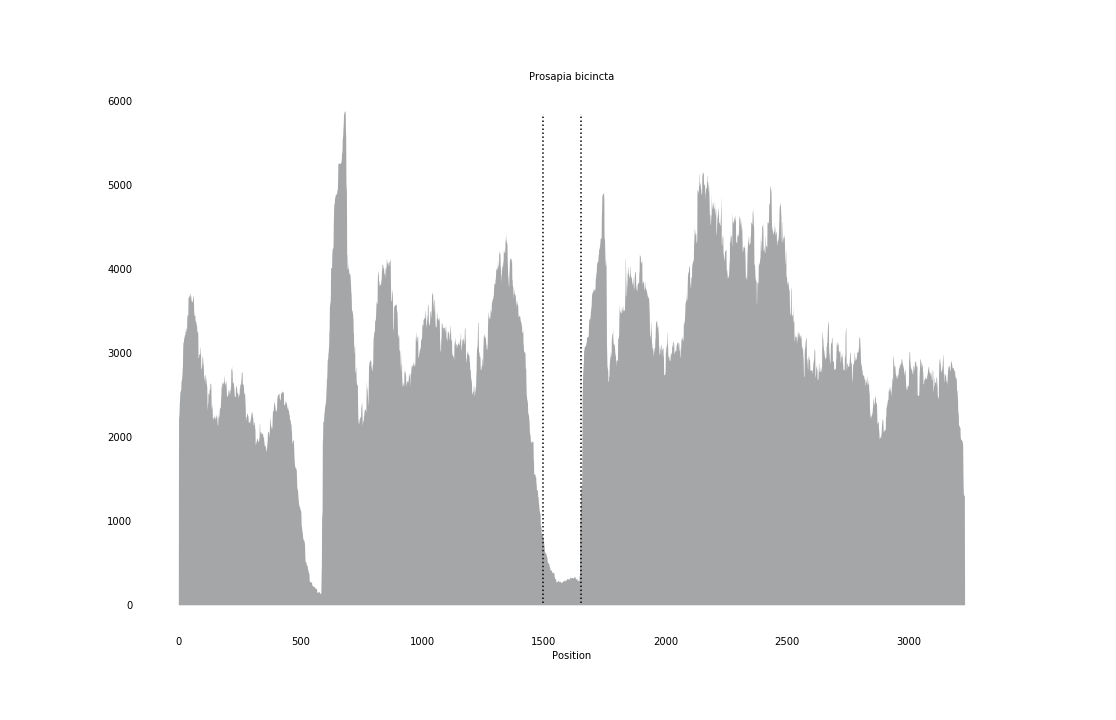

### Pseudomasaris_vespoides_coverage_correct.png

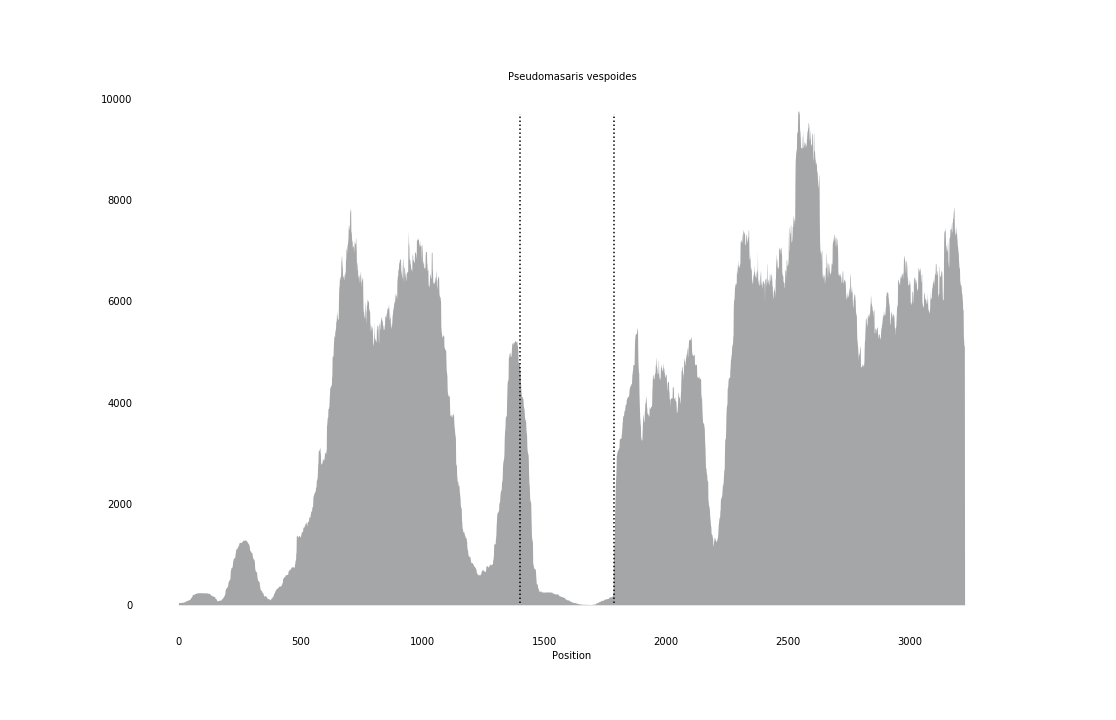

### Ramphogordius_sanguineus_coverage.png

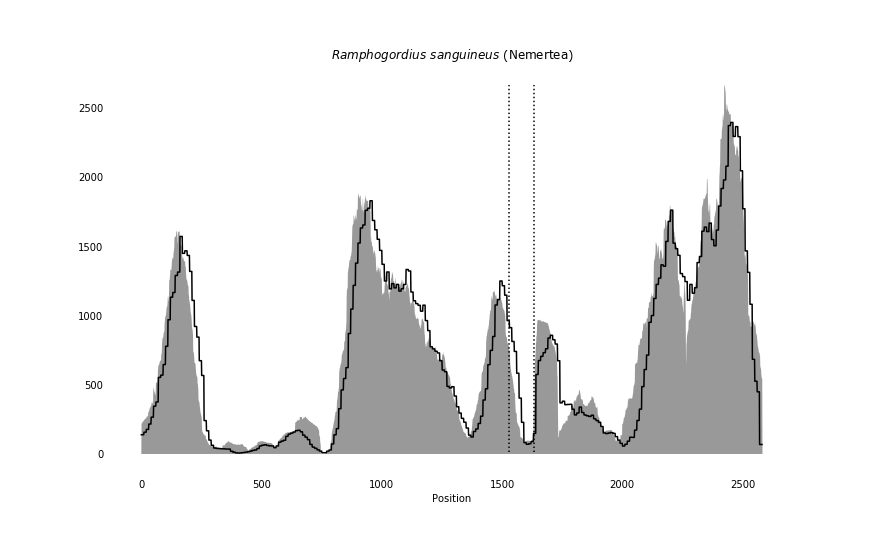

### Ruditapes_philippinarum_coverage_correct.png

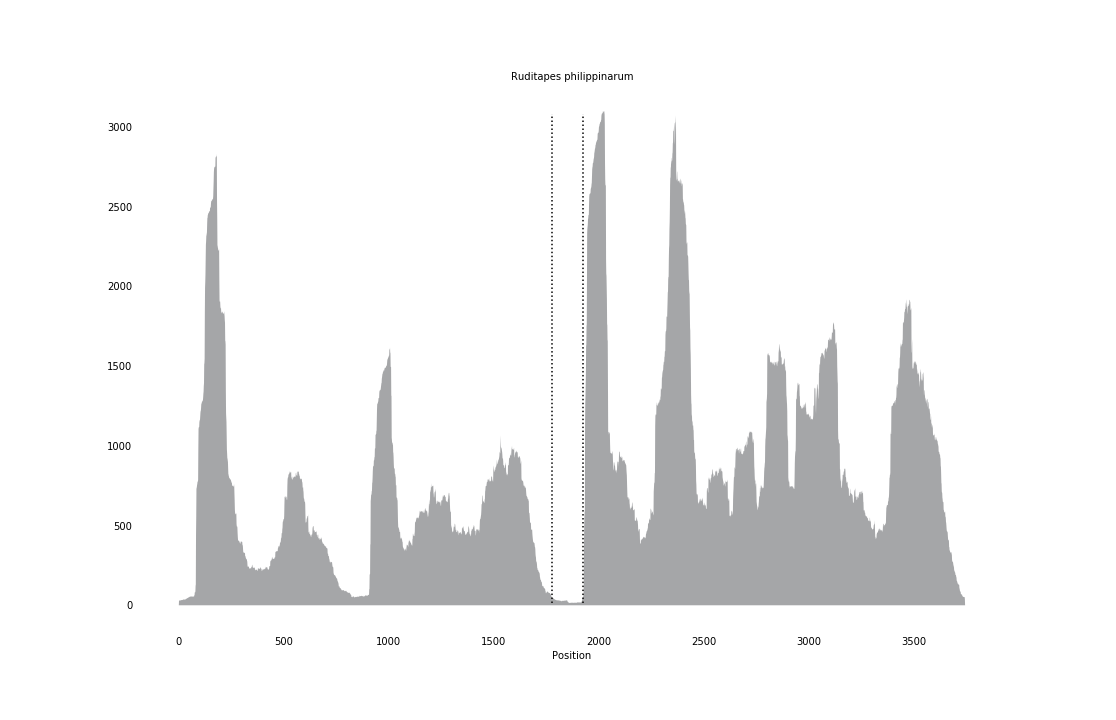

### Sapyga_quinquepunctata_coverage_correct.png

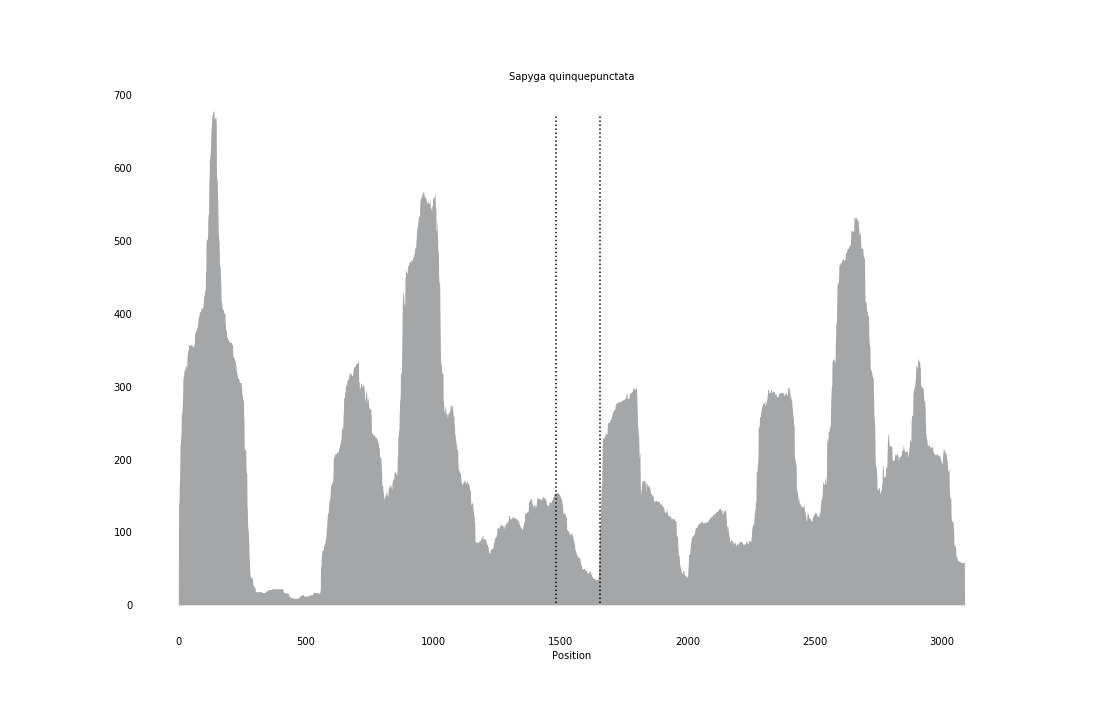

### Sapygina_decemguttata_coverage_correct.png

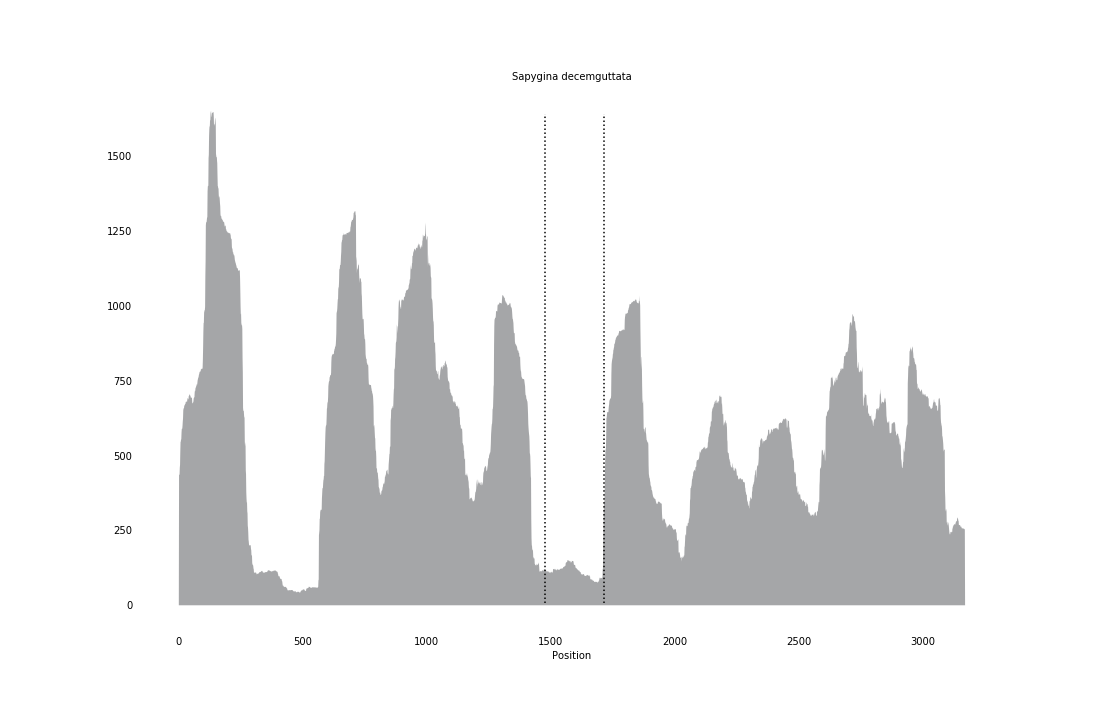

### Sceliphron_curvatum_coverage_correct.png

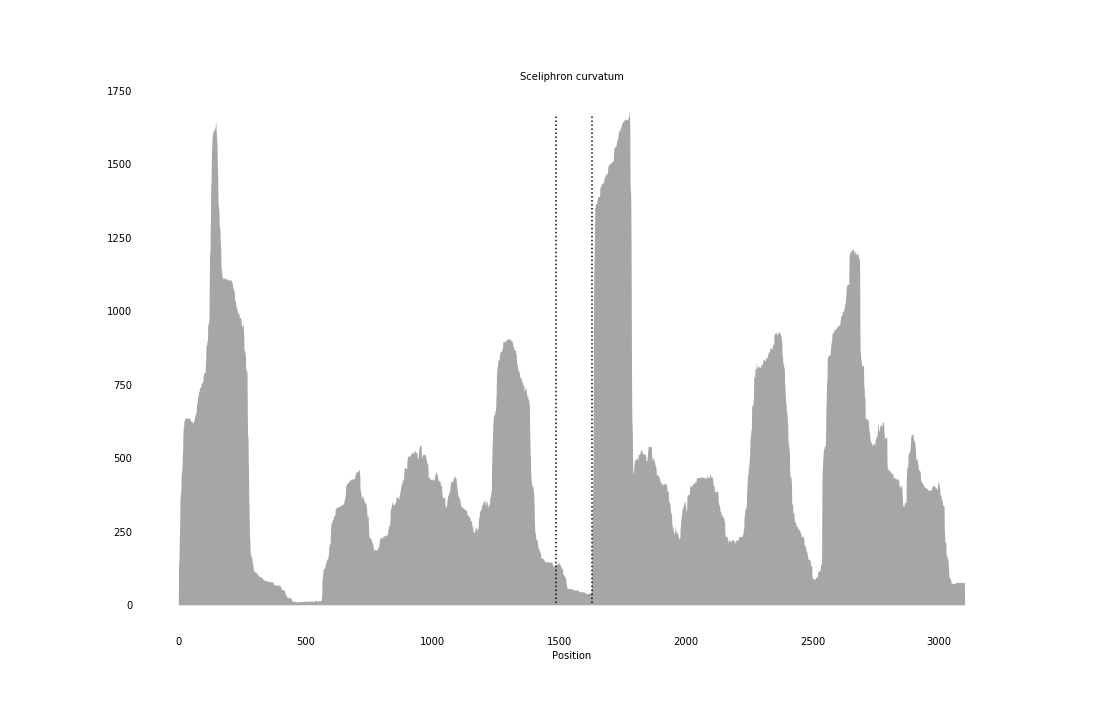

### Septifer_virgatus_coverage_correct.png

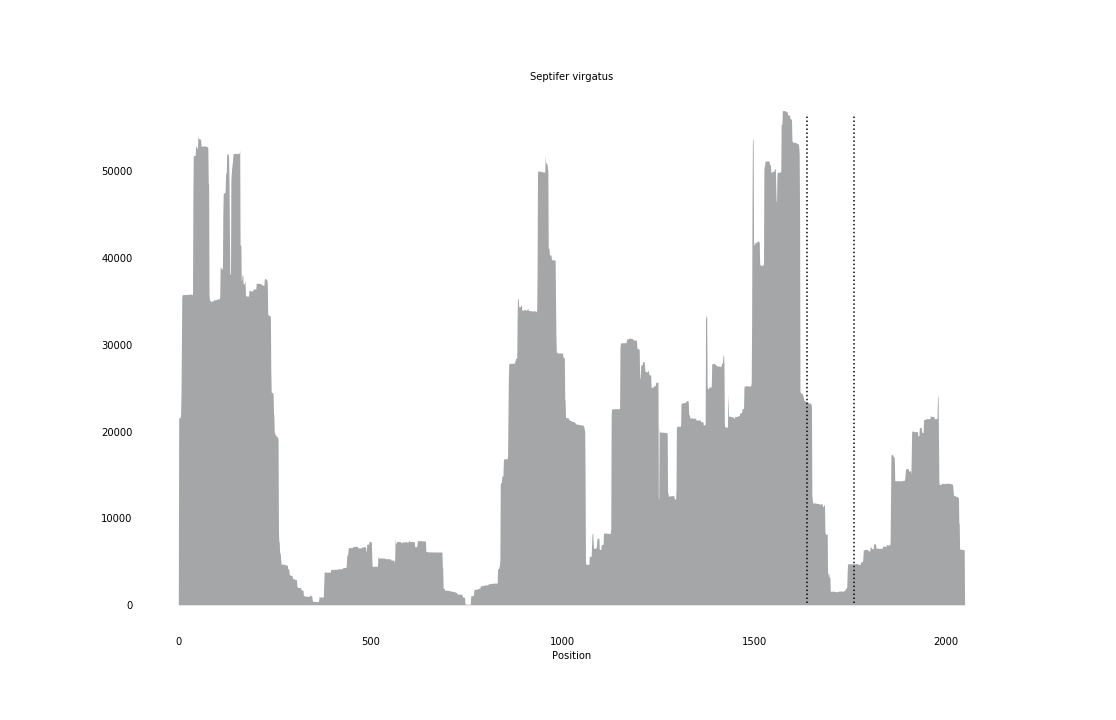

### Smicromyrme_rufipes_coverage_correct.png

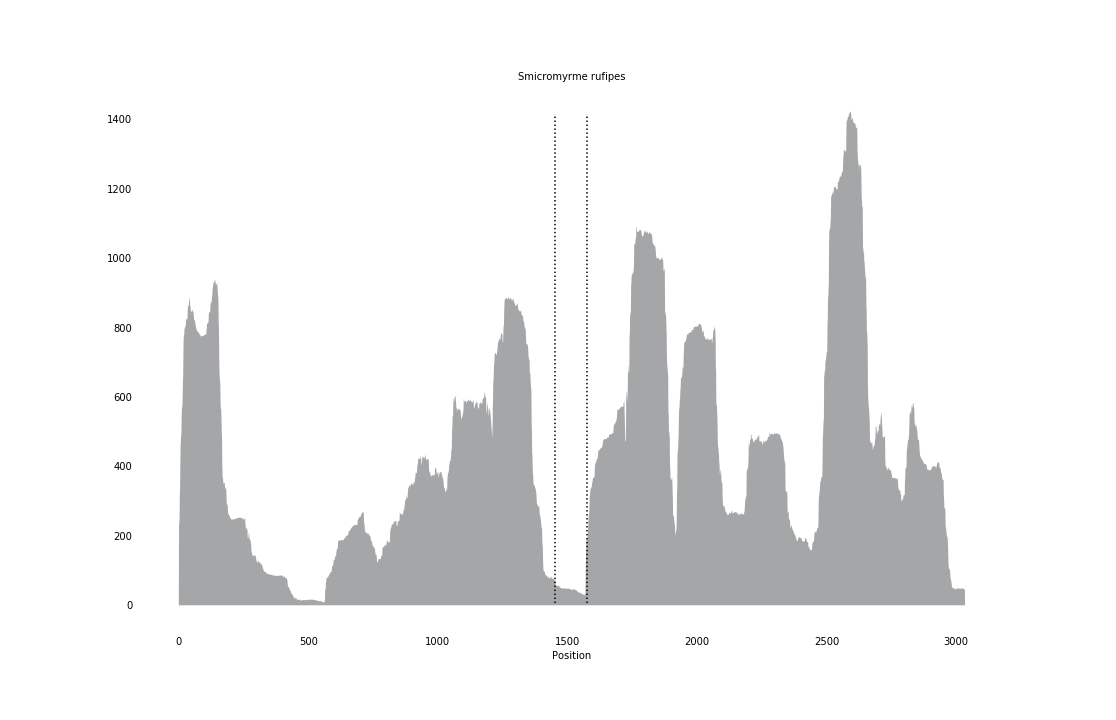

### Sminthurus_viridis_coverage_correct.png

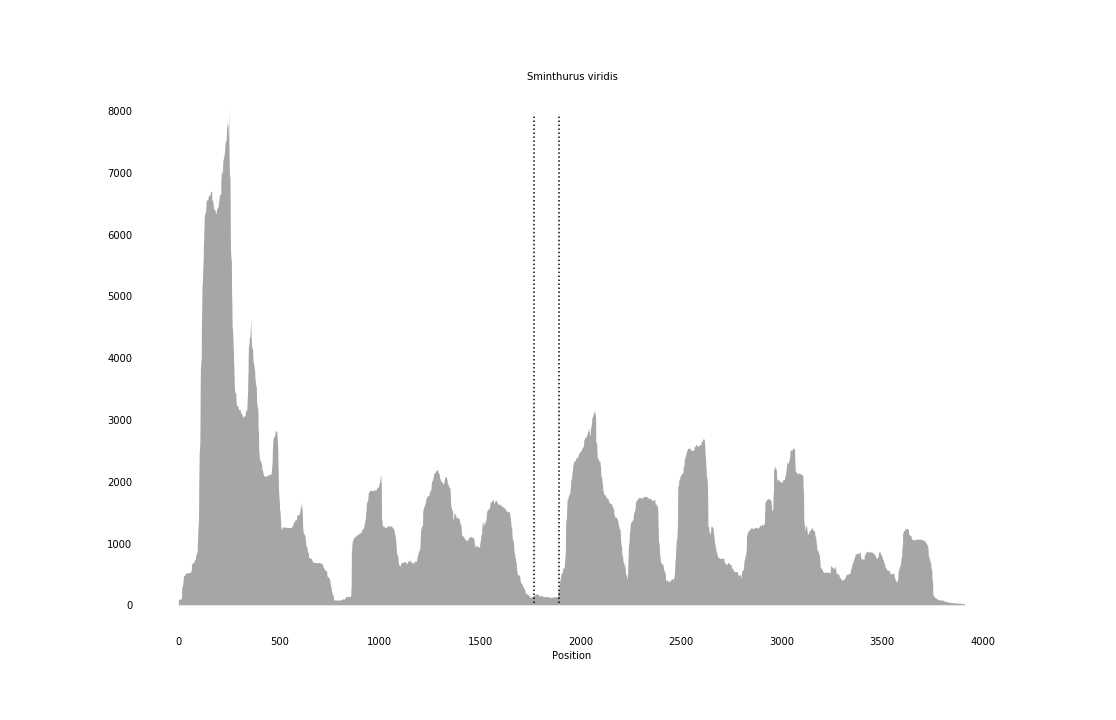

### Solemya_velum_coverage_correct.png

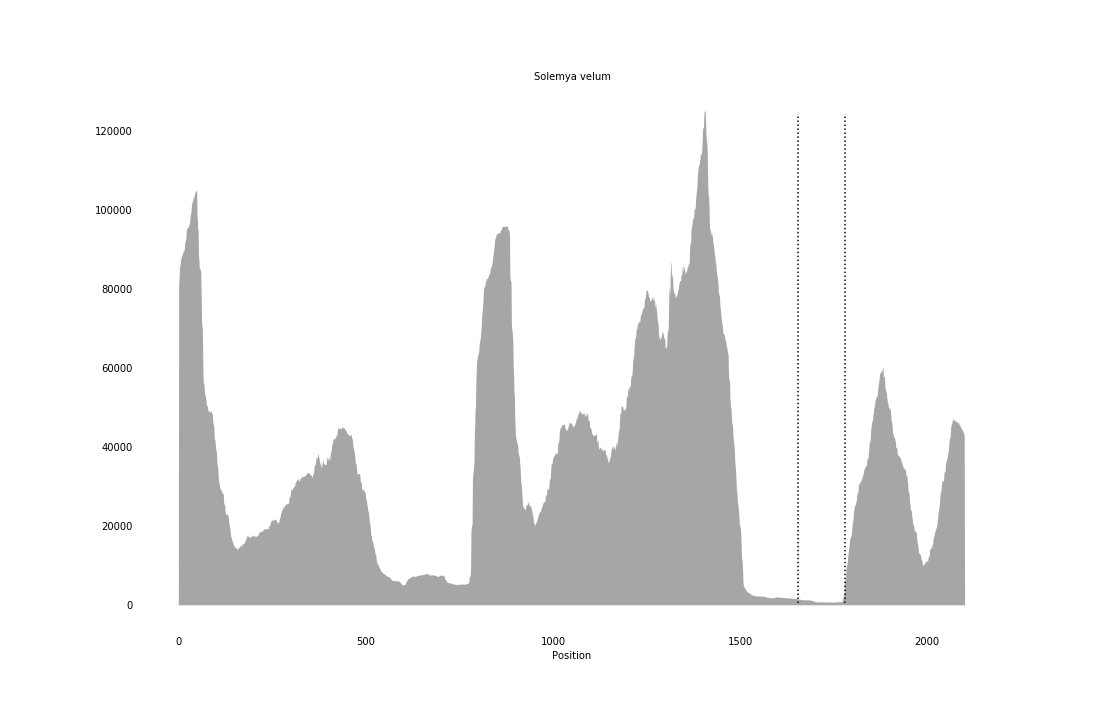

### Sphaeropthalma_orestes_coverage_correct.png

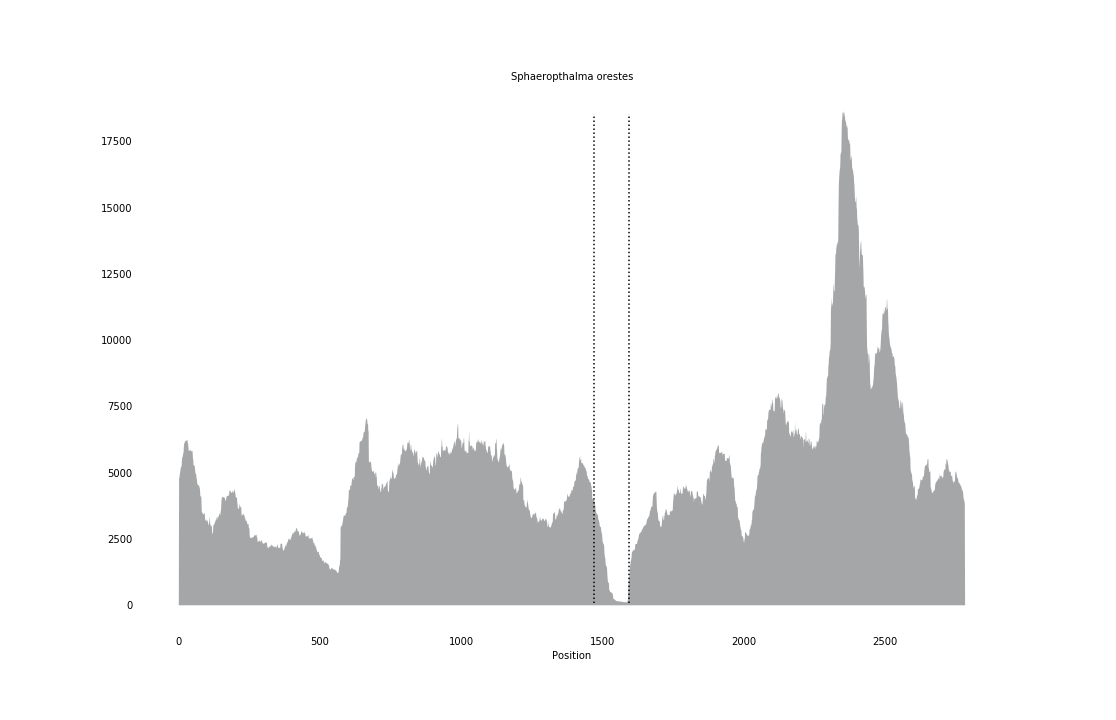

### Stenostomum_leucops_coverage_correct.png

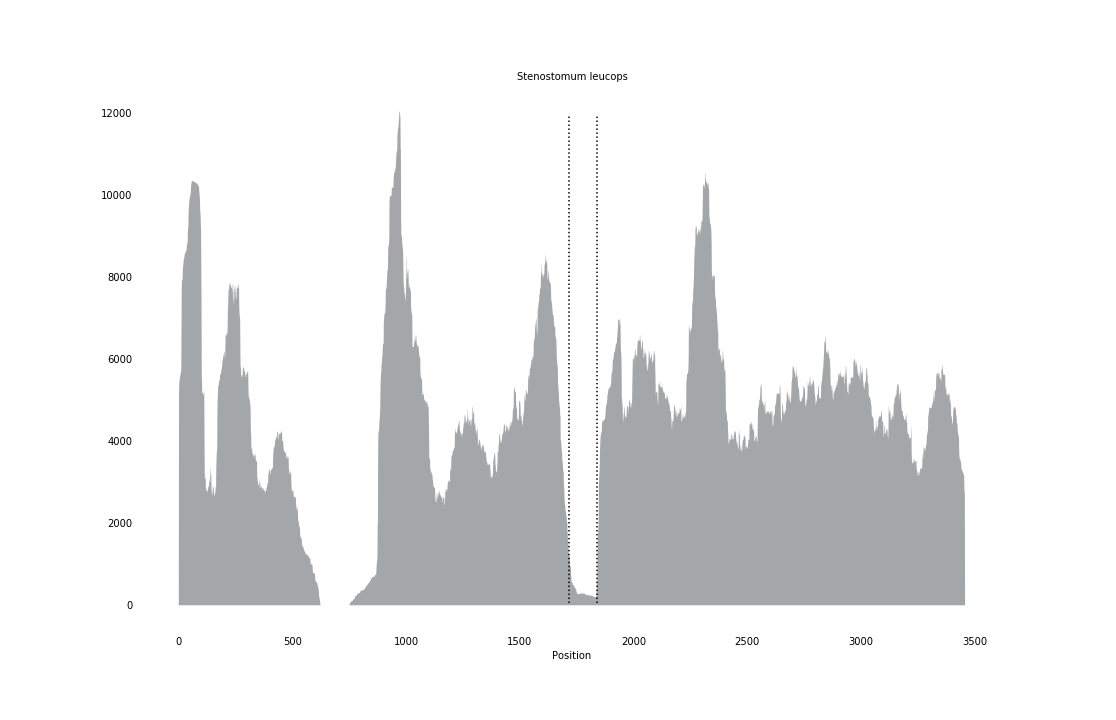

### Stephanus_serrator_coverage_correct.png

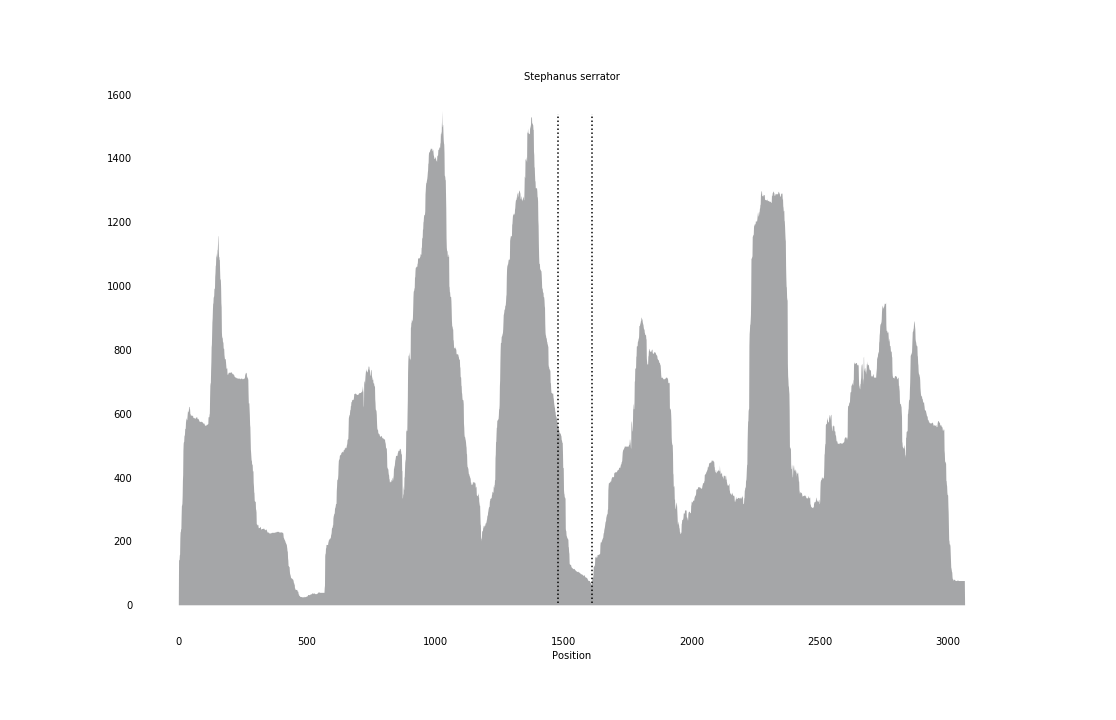

### Tenthredo_koehleri_coverage_correct.png

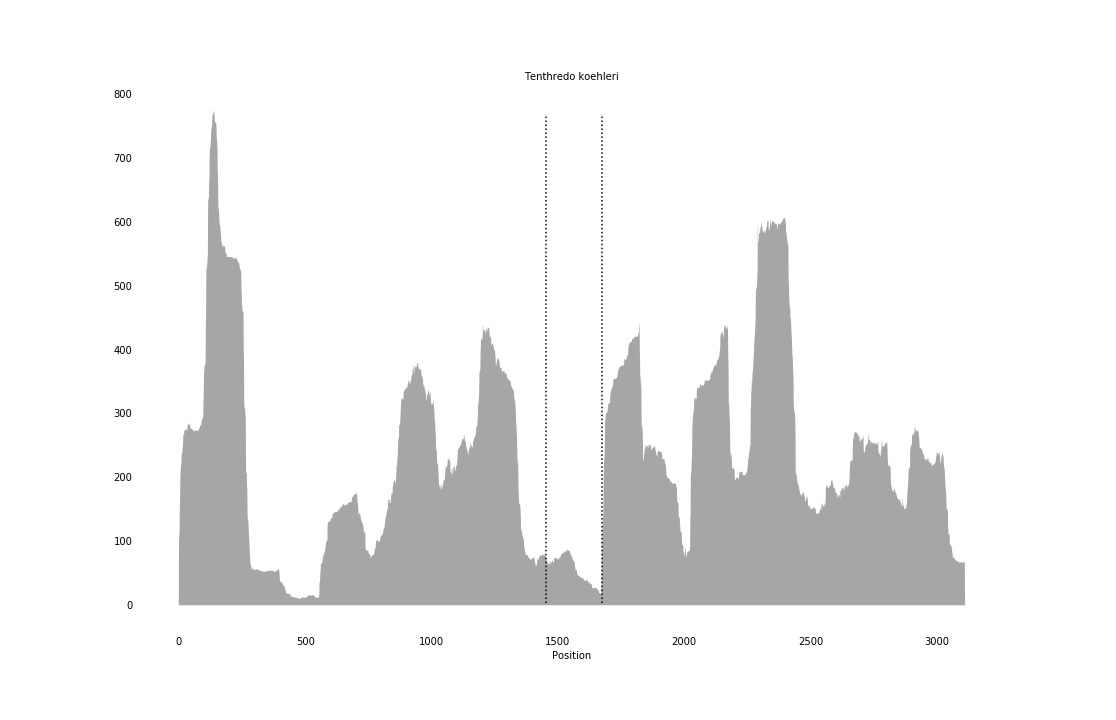

### Trialeurodes_vaporariorum_coverage_correct.png

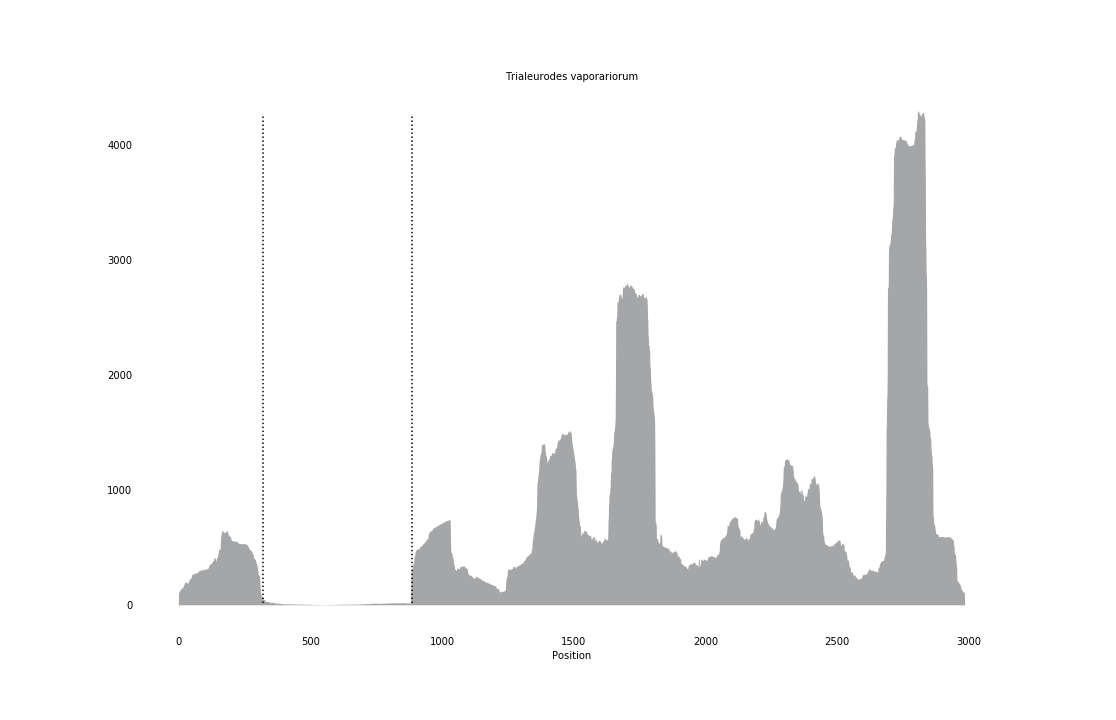

### Trichobilharzia_regenti_coverage_correct.png

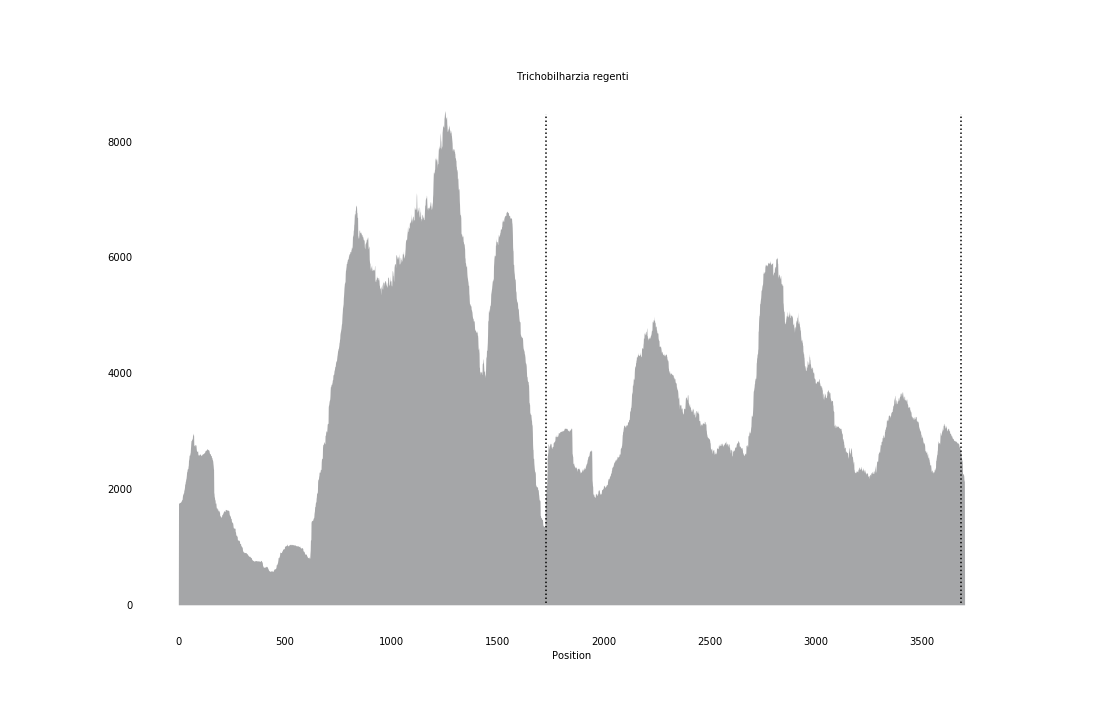

### Trichopria_drosophilae_coverage_correct.png

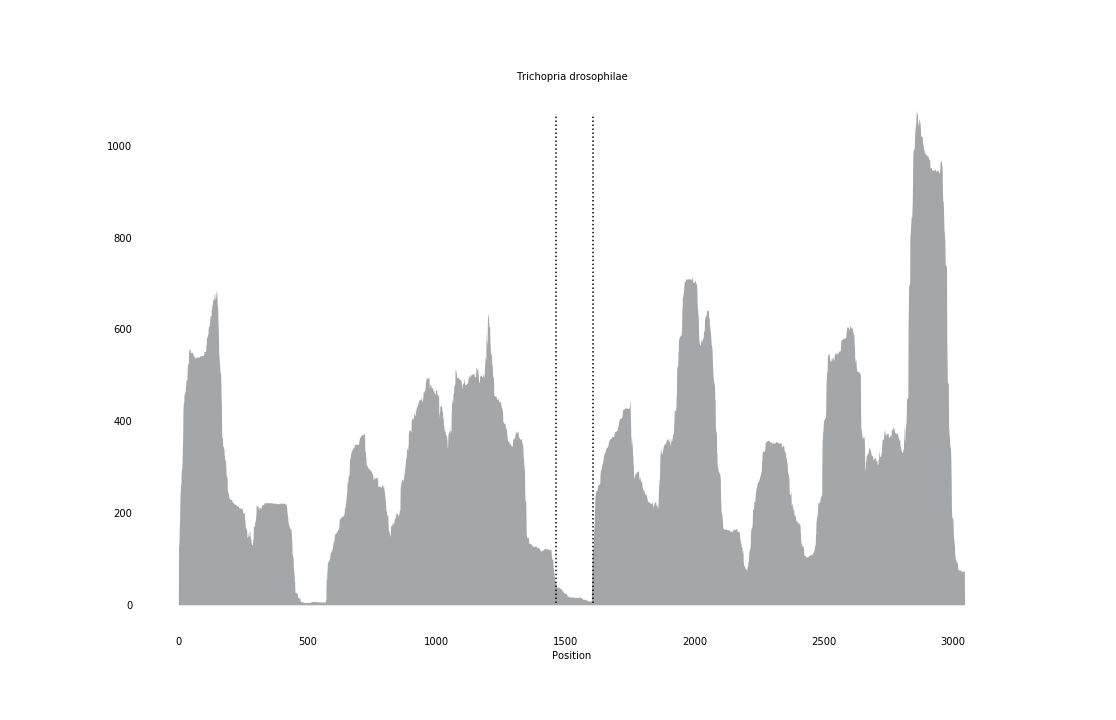

### Trypoxylon_figulus_coverage_correct.png

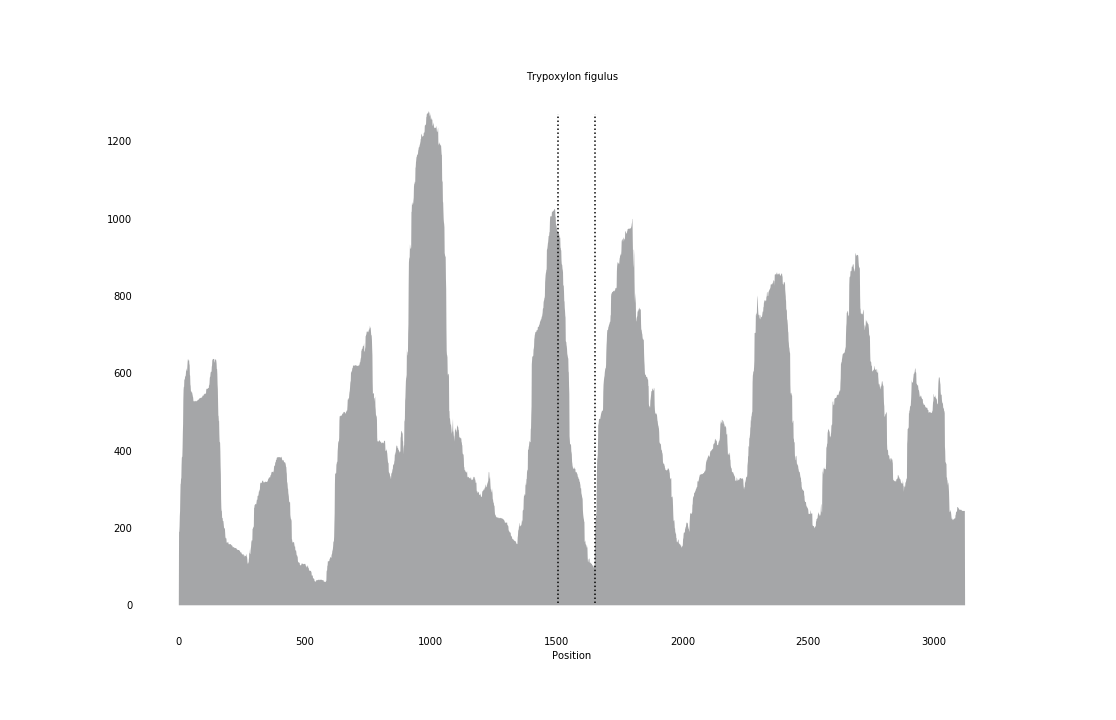

### Tupiocoris_notatus_coverage_correct.png

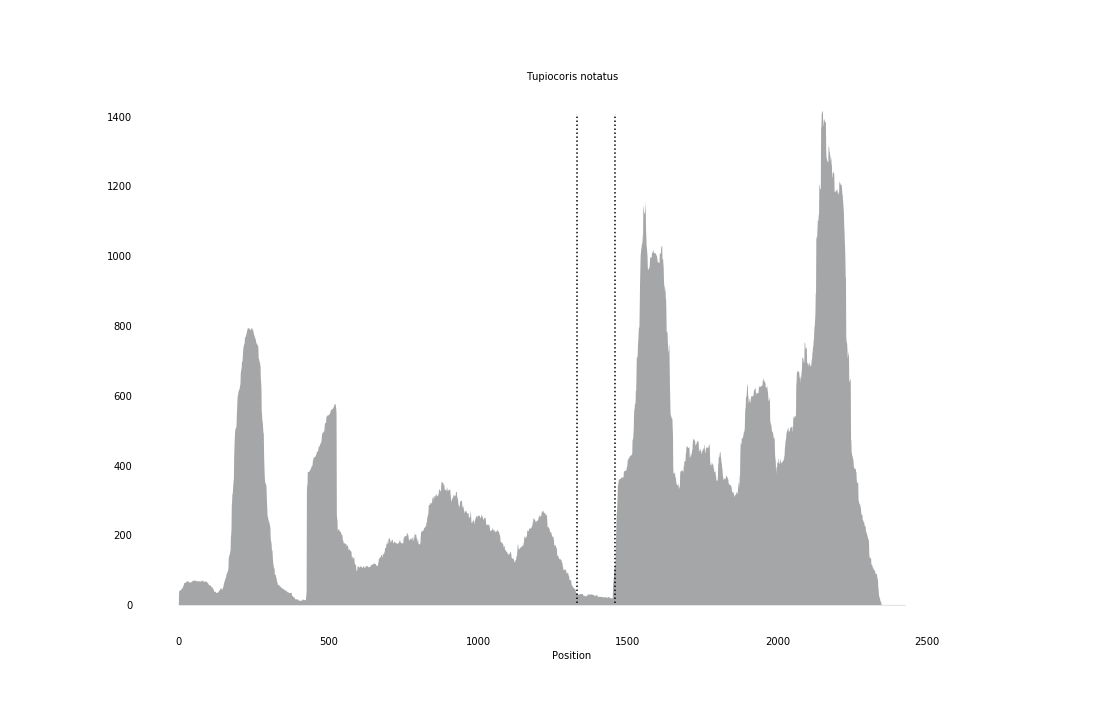

### Tytthus_chinensis_coverage_correct.png

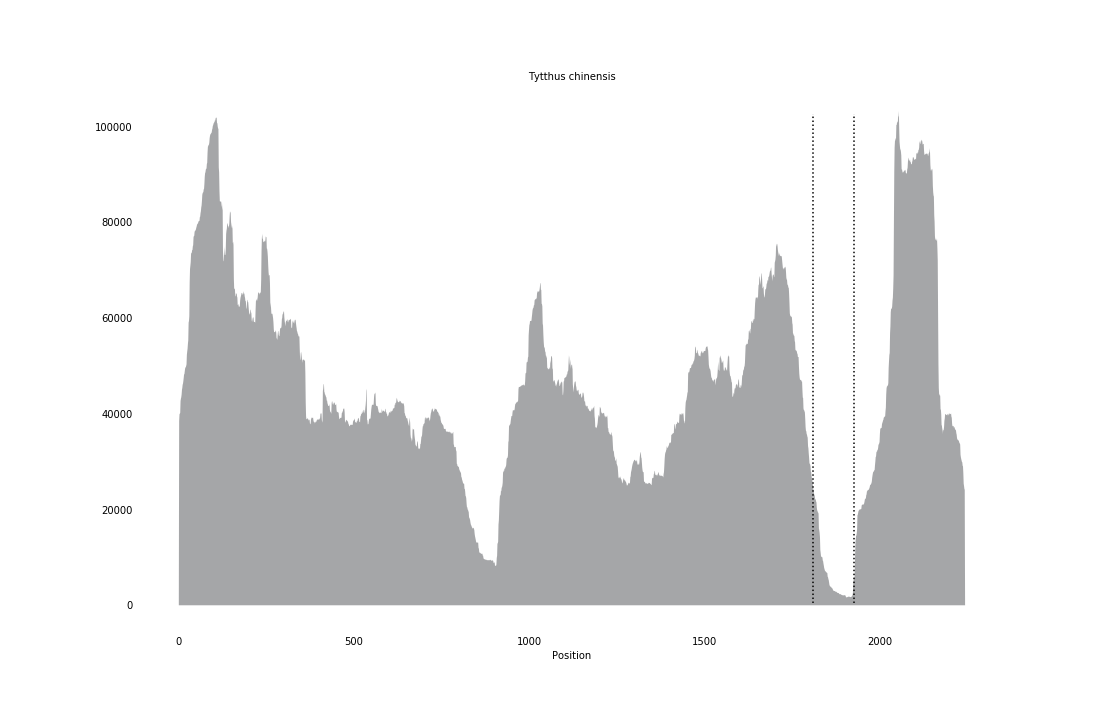

### Urocerus_augur_coverage_correct.png

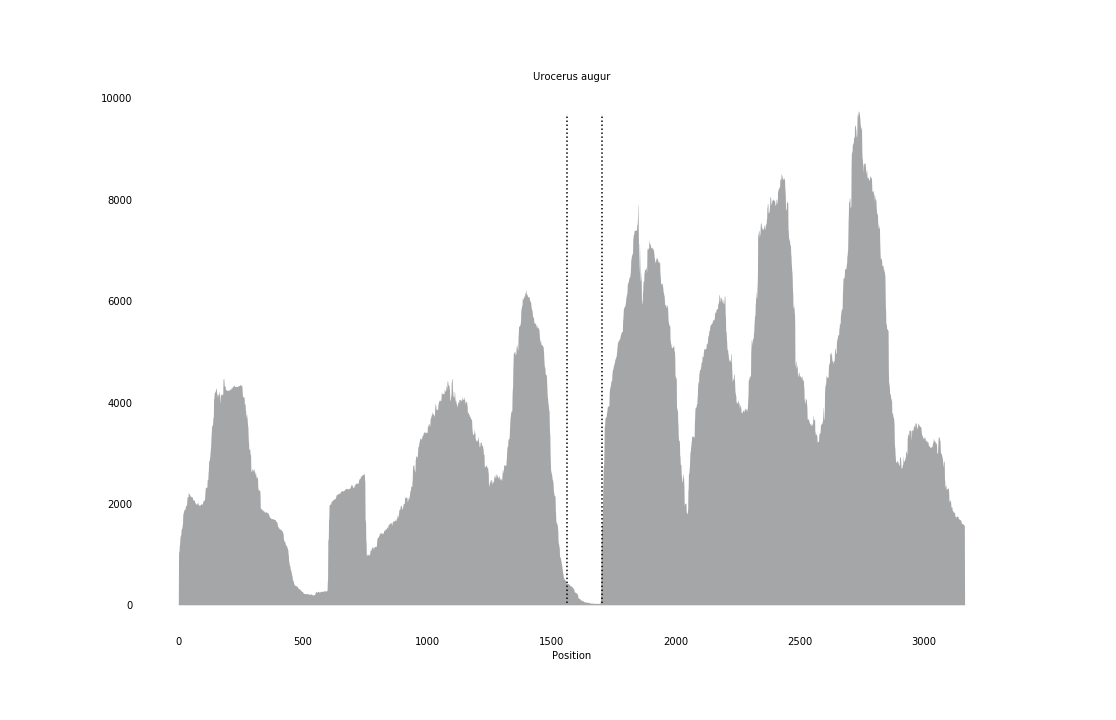

### Vespa_crabro_coverage.png

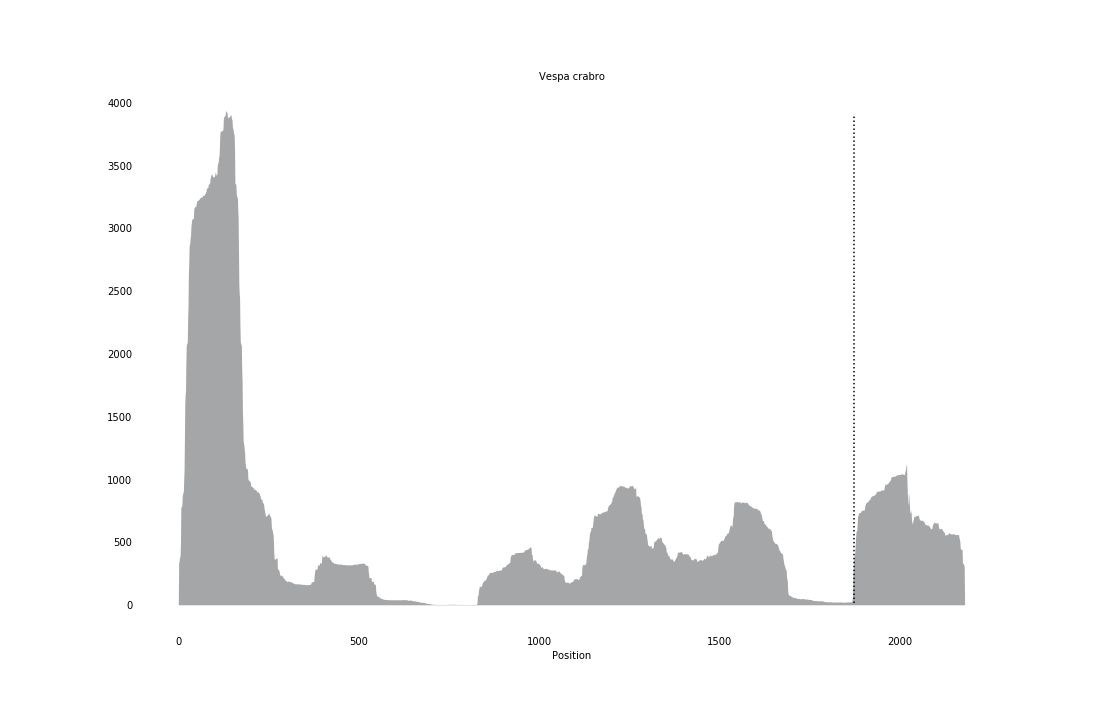

### Watasenia_scintillans_coverage_correct.png

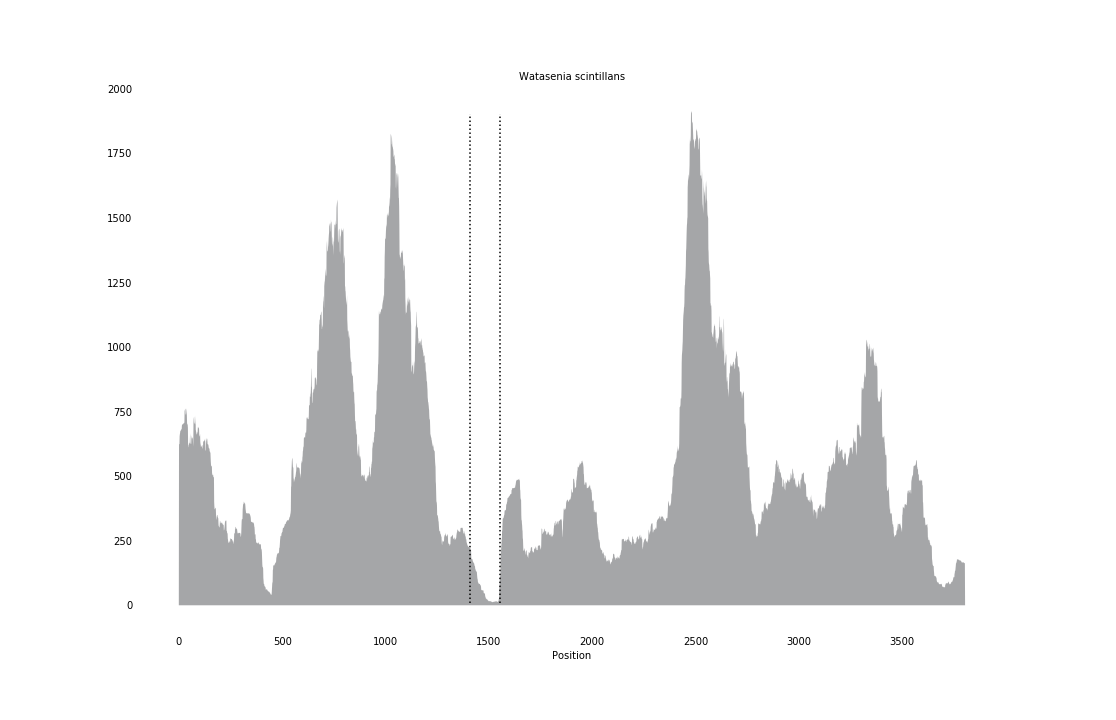

### Xylocopa_violacea_coverage_correct.png

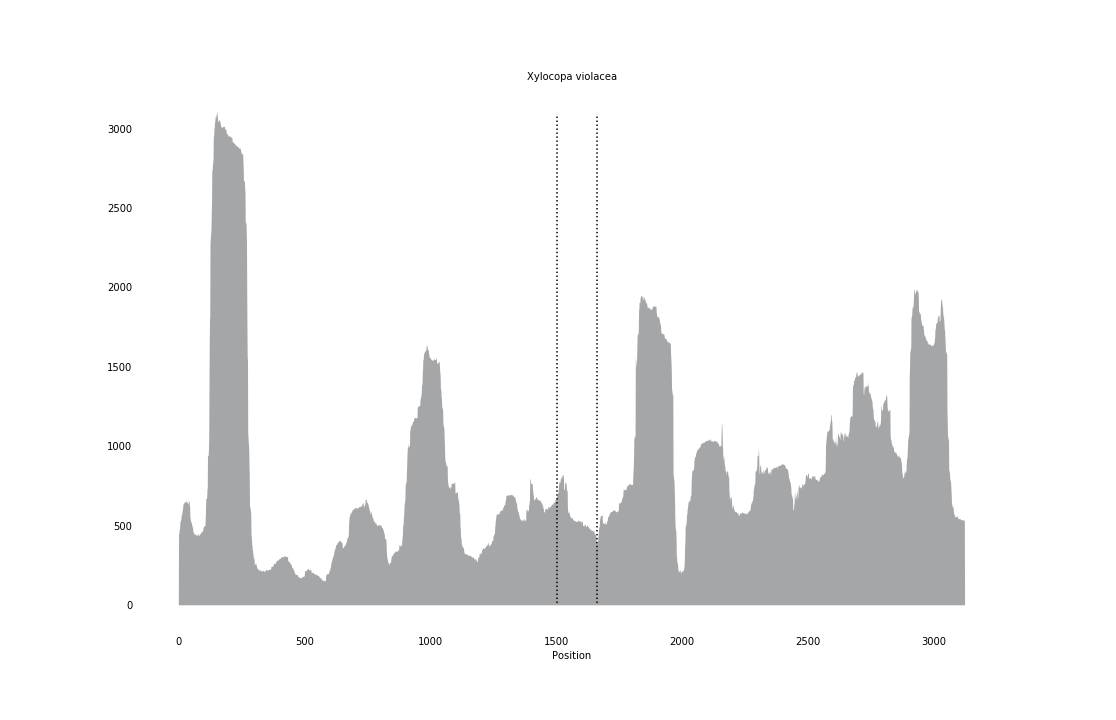
