## Supplementary figures and images for "Computational discovery of hidden breaks in 28S ribosomal RNAs across eukaryotes and consequences for RNA Integrity Numbers"

### Abrothrix_longipilis_coverage_correct.png

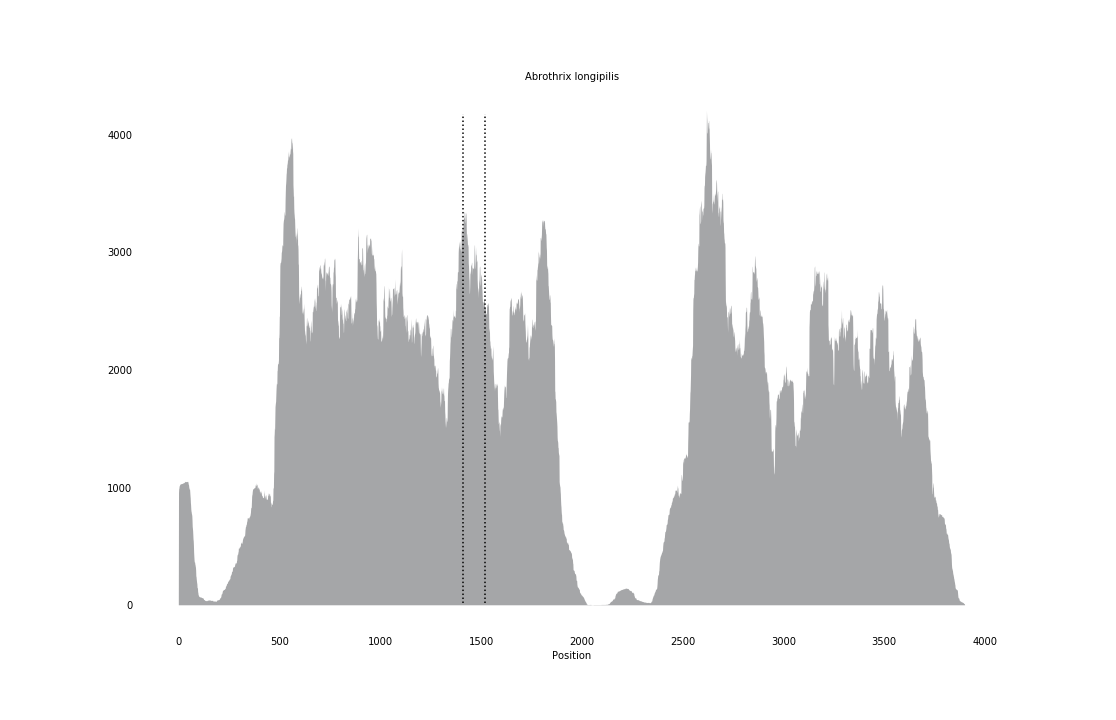

### Amia_calva_coverage_correct.png

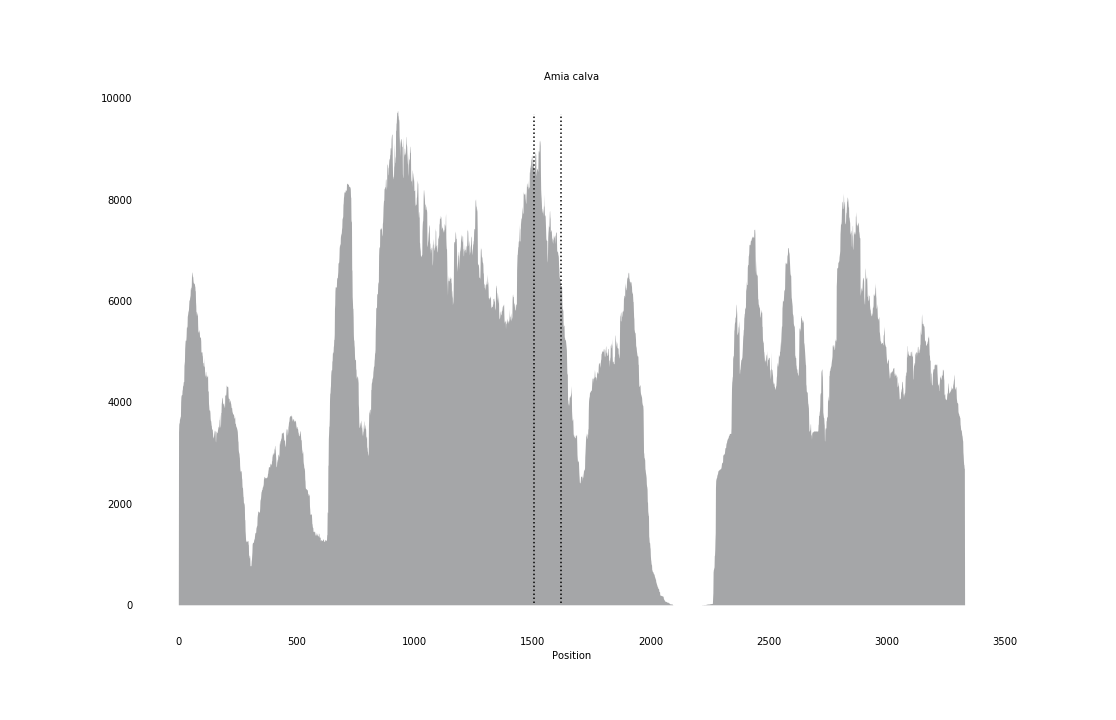

### Amphiophiura_laudata_coverage.png

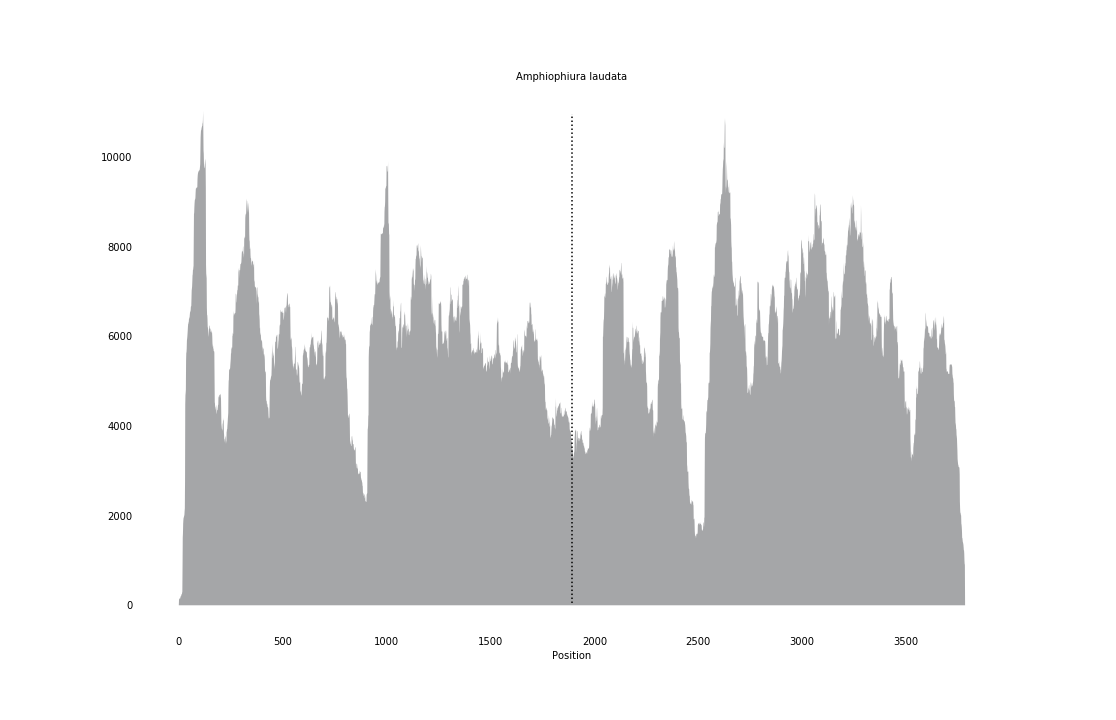

### Amphiura_constricta_coverage.png

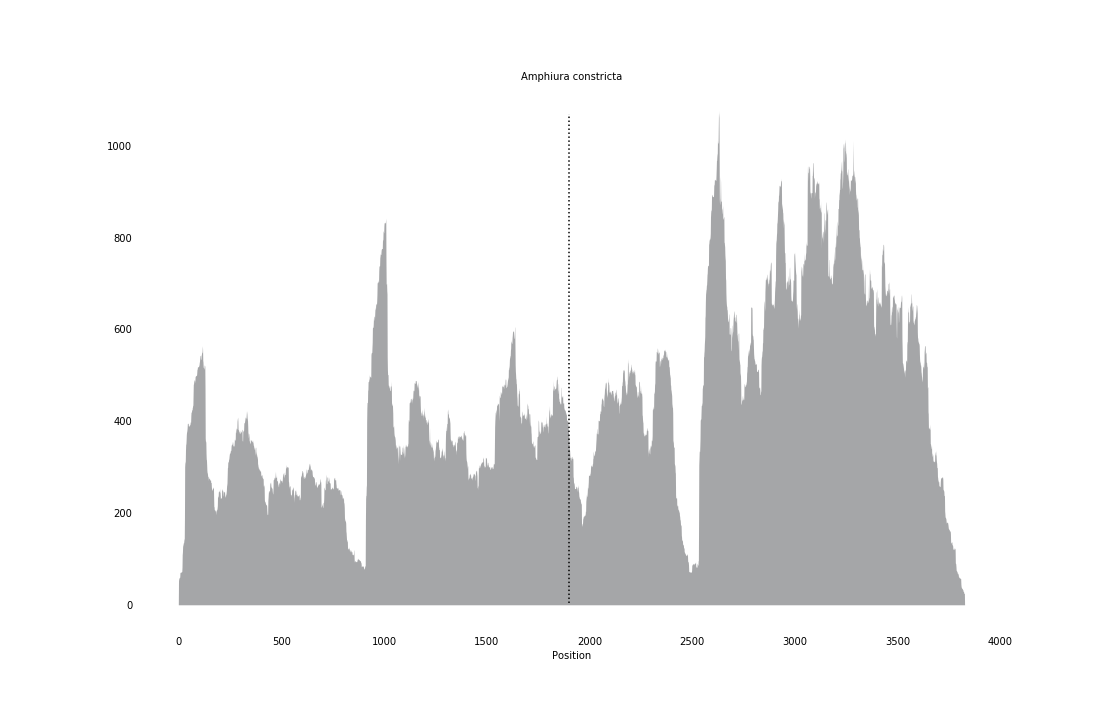

### Anguilla_anguilla_coverage_correct.png

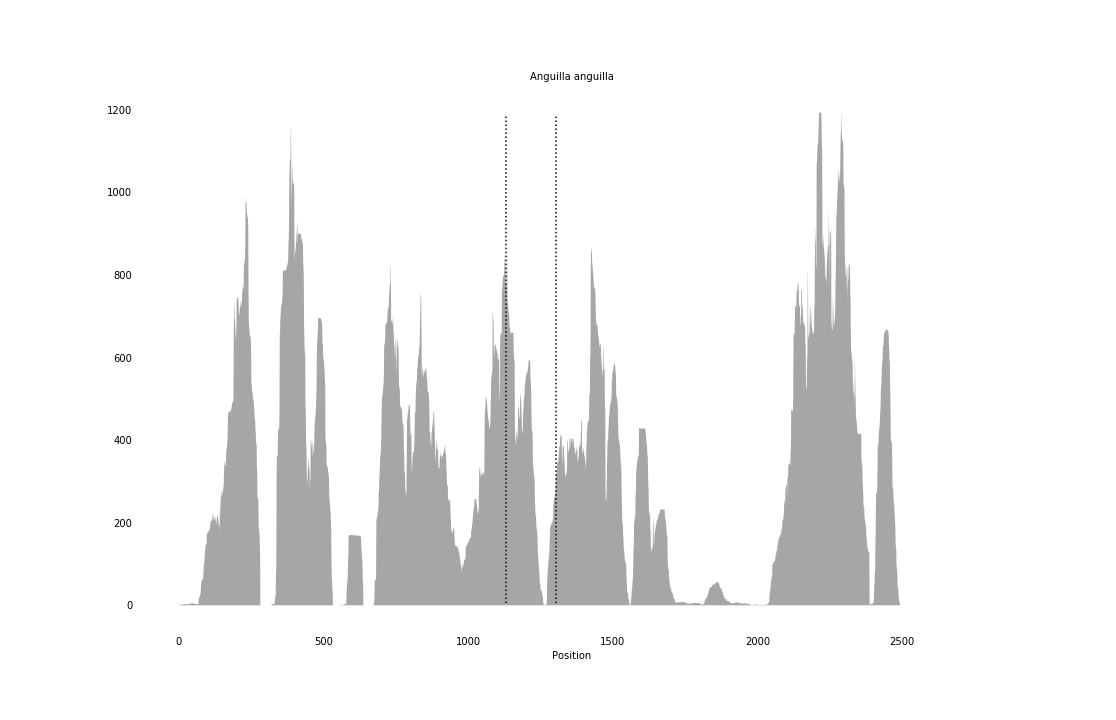

### Aporometra_wilsoni_coverage.png

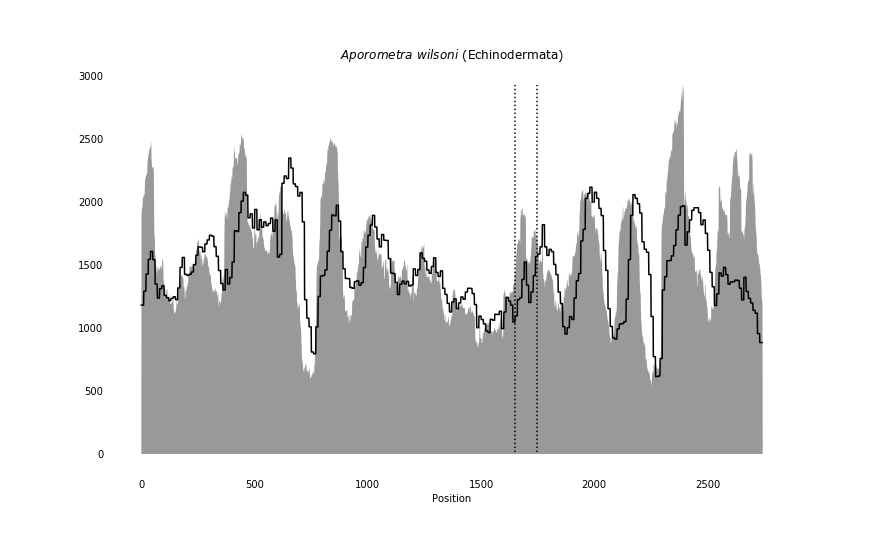

### Aspidosiphon_parvulus_coverage_correct.png

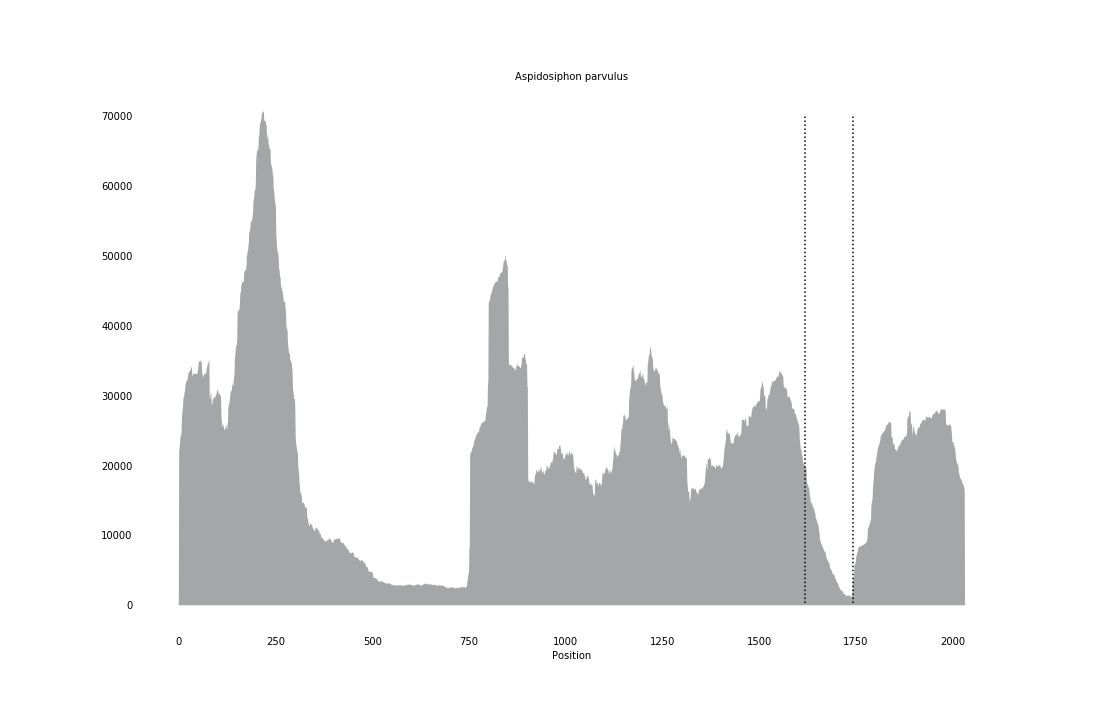

### Asterias_forbesii_coverage.png

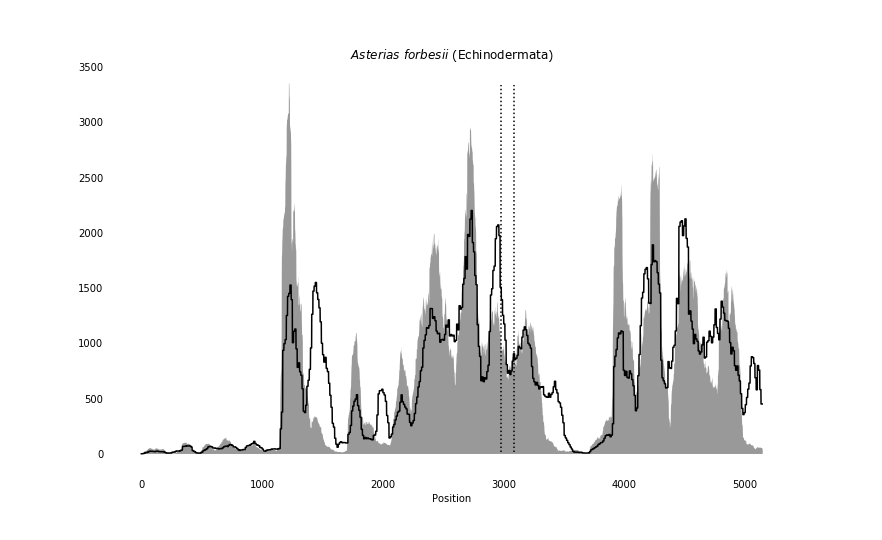

### Asteronyx_loveni_coverage.png

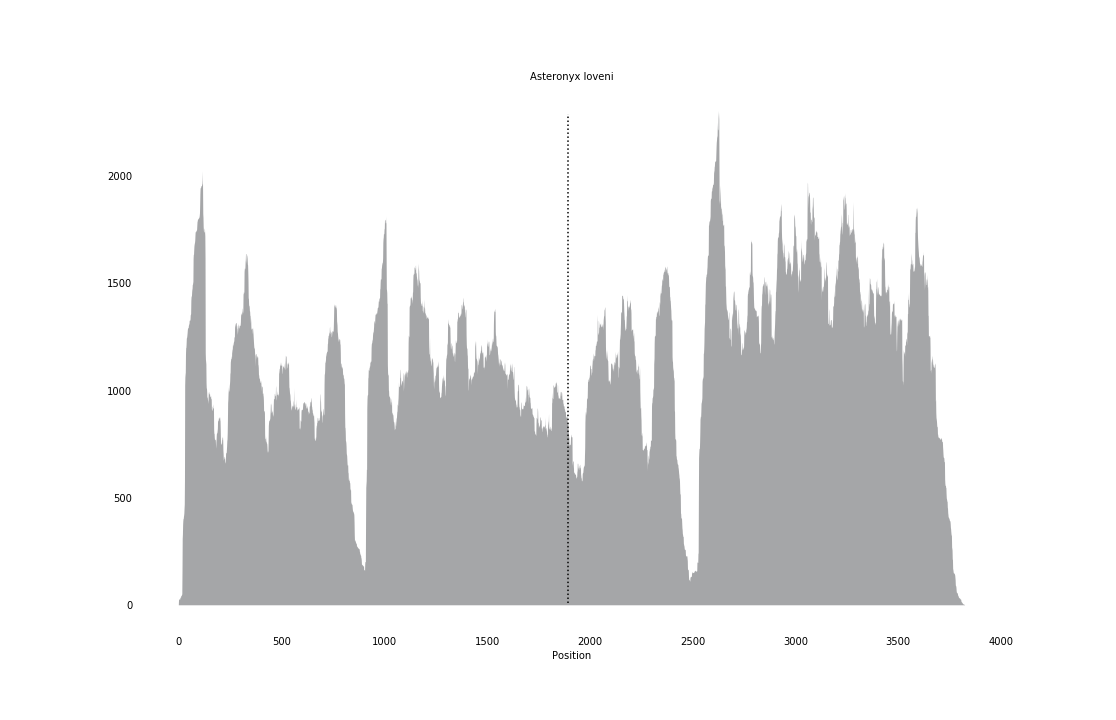

### Asteroschema_bidwillae_coverage.png

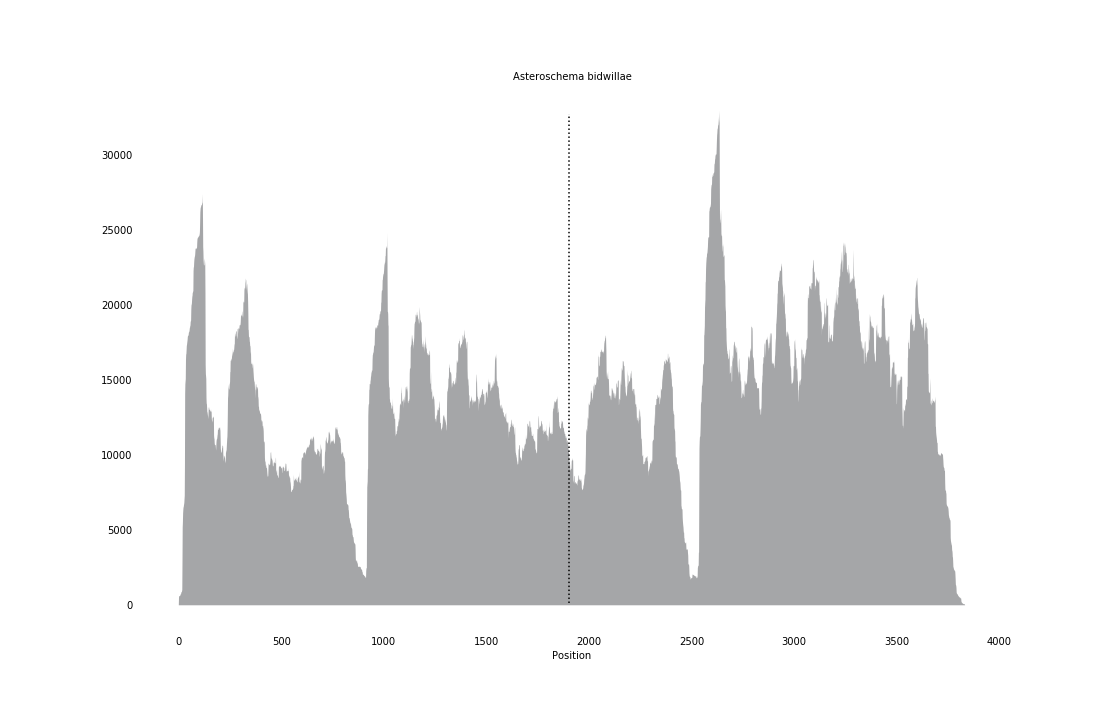

### Austrofundulus_limnaeus_coverage_correct.png

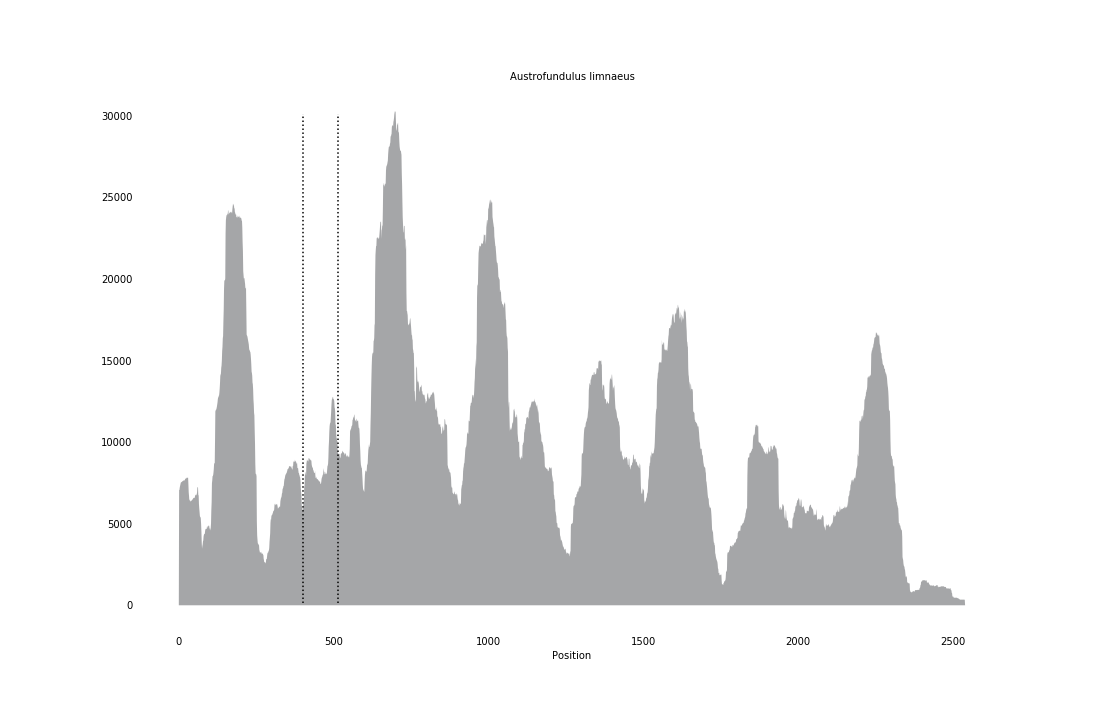

### Barentsia_gracilis_overage.png

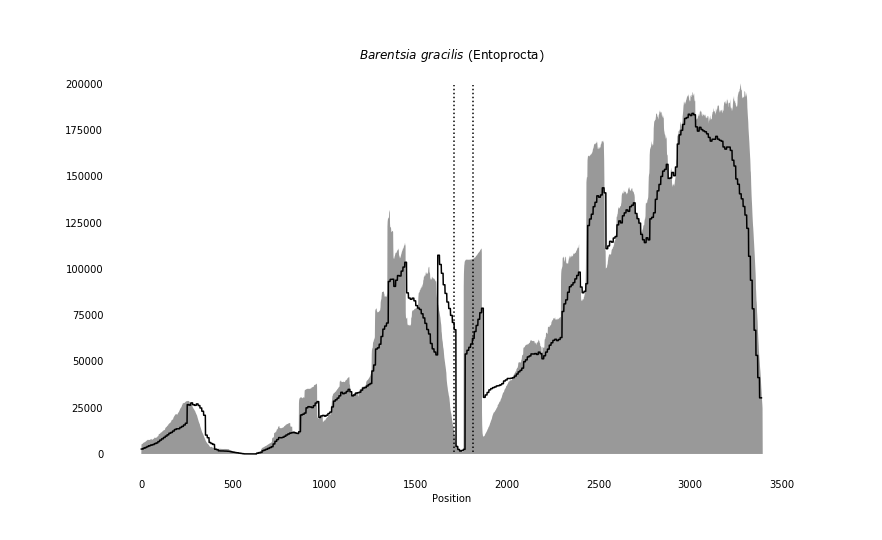

### Bathypectinura_heros_coverage.png

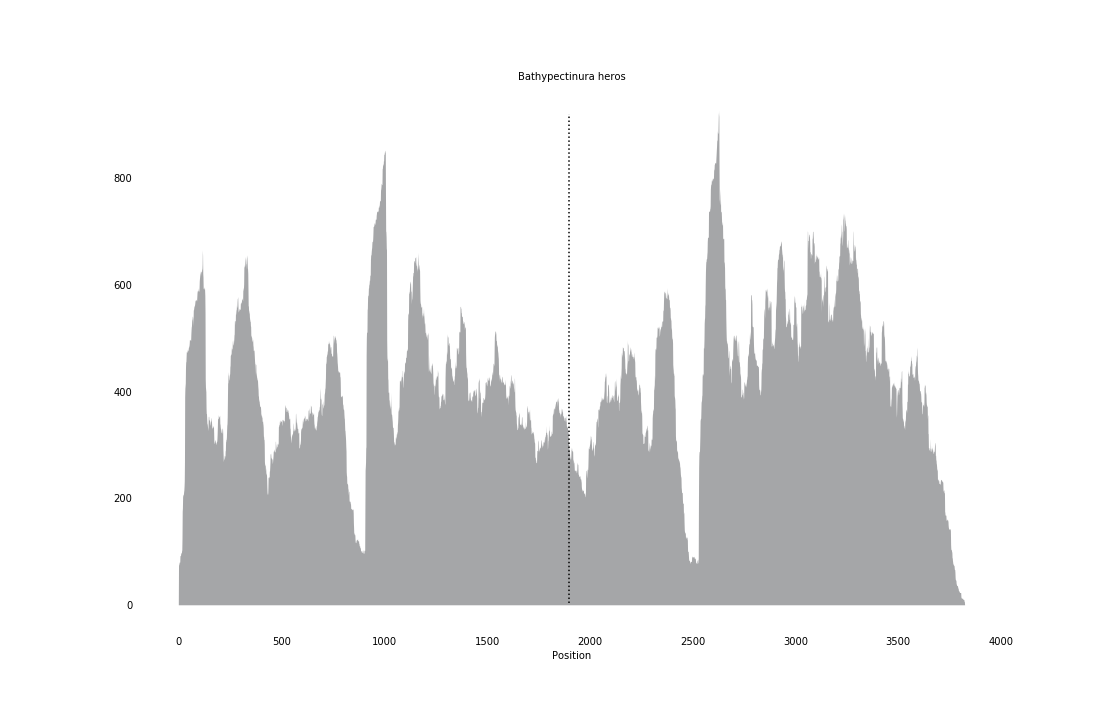

### Caenorhabditis_briggsae_coverage_correct.png

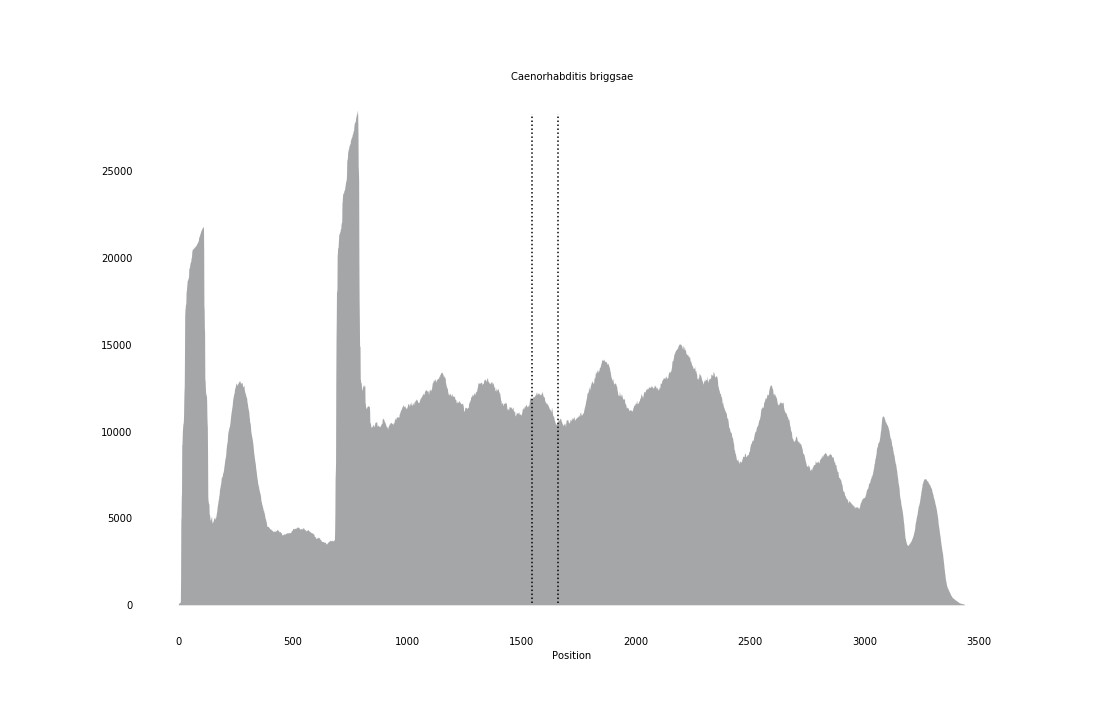

### Cerebratulus_lacteus_coverage.png

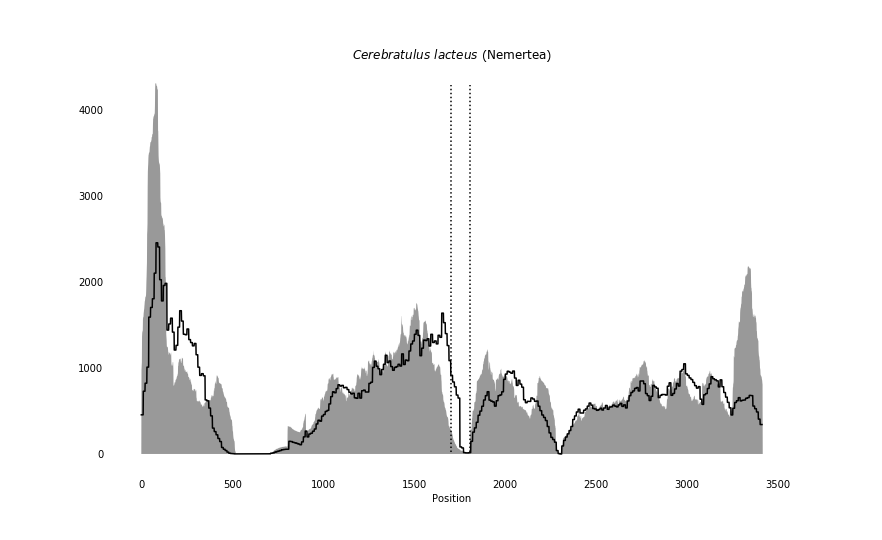

### Channa_argus_coverage_correct.png

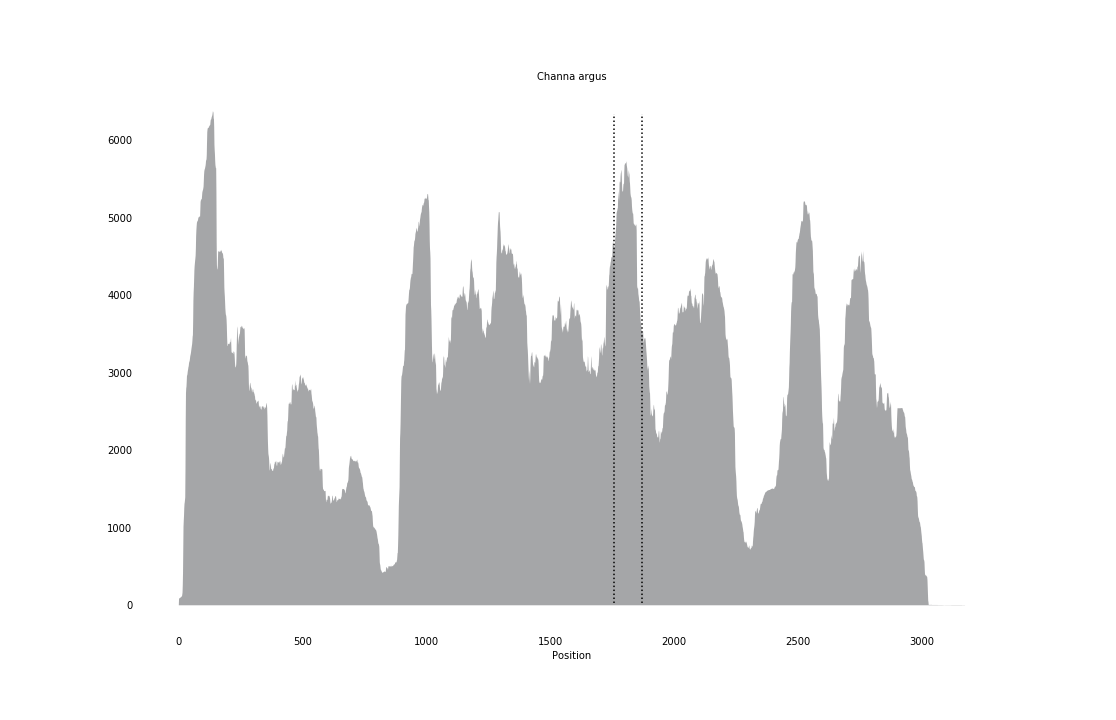

### Chanos_chanos_coverage_correct.png

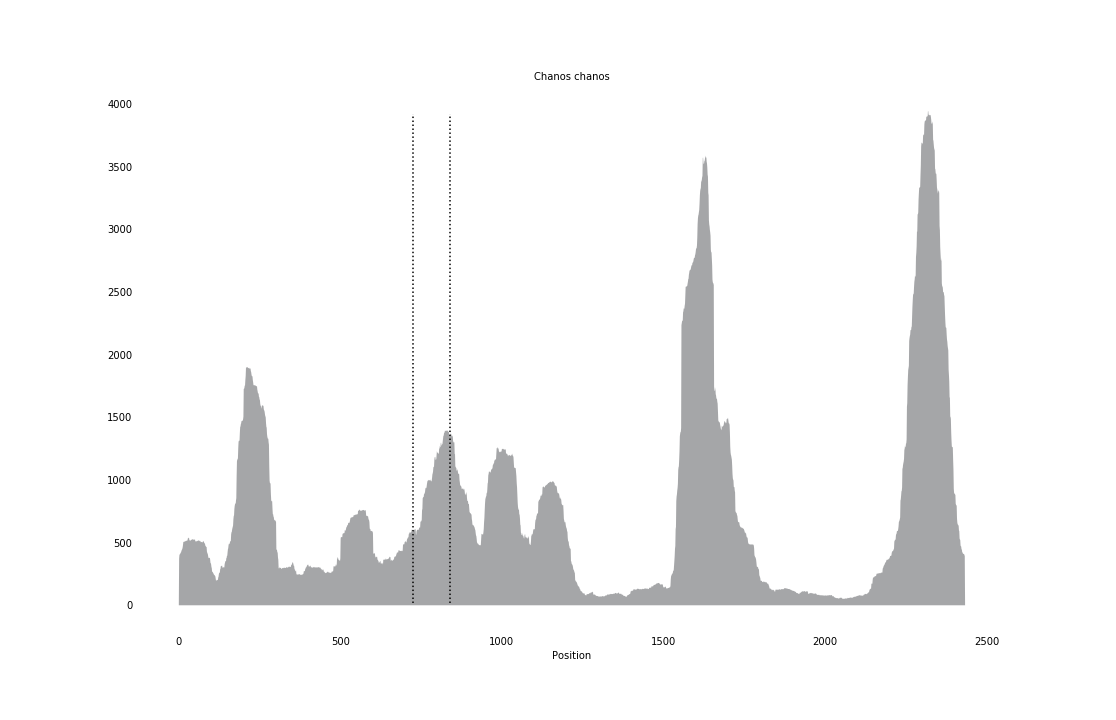

### Childia_submaculatum_coverage_correct.png

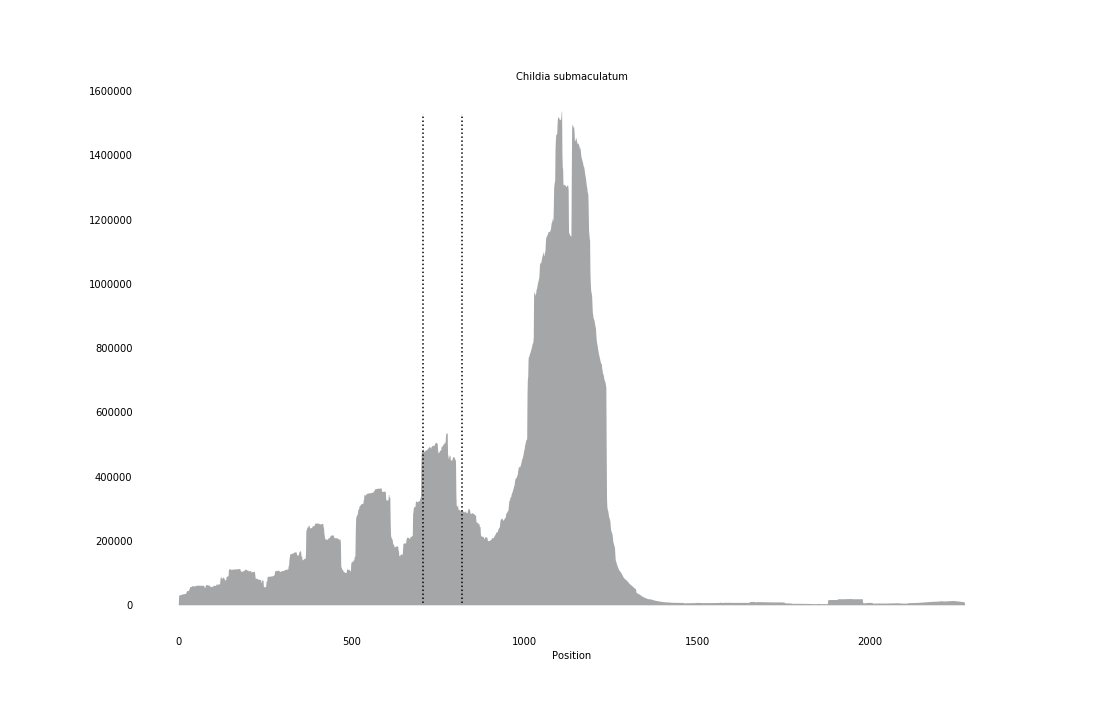

### Ciona_intestinalis_coverage.png

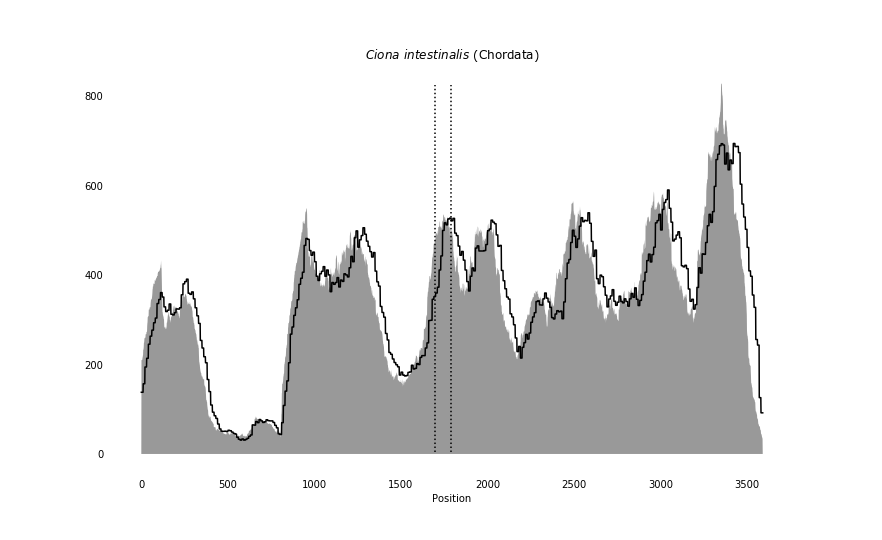

### Ciona_savignyi_coverage_correct.png

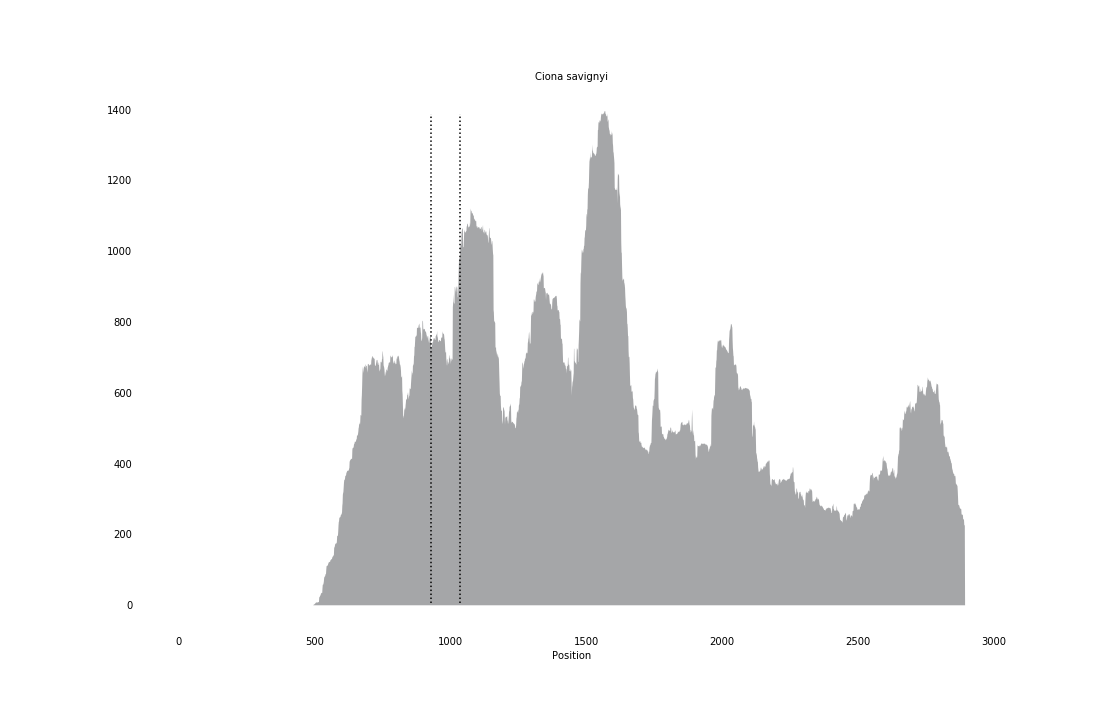

### Clarkcoma_canaliculata_coverage.png

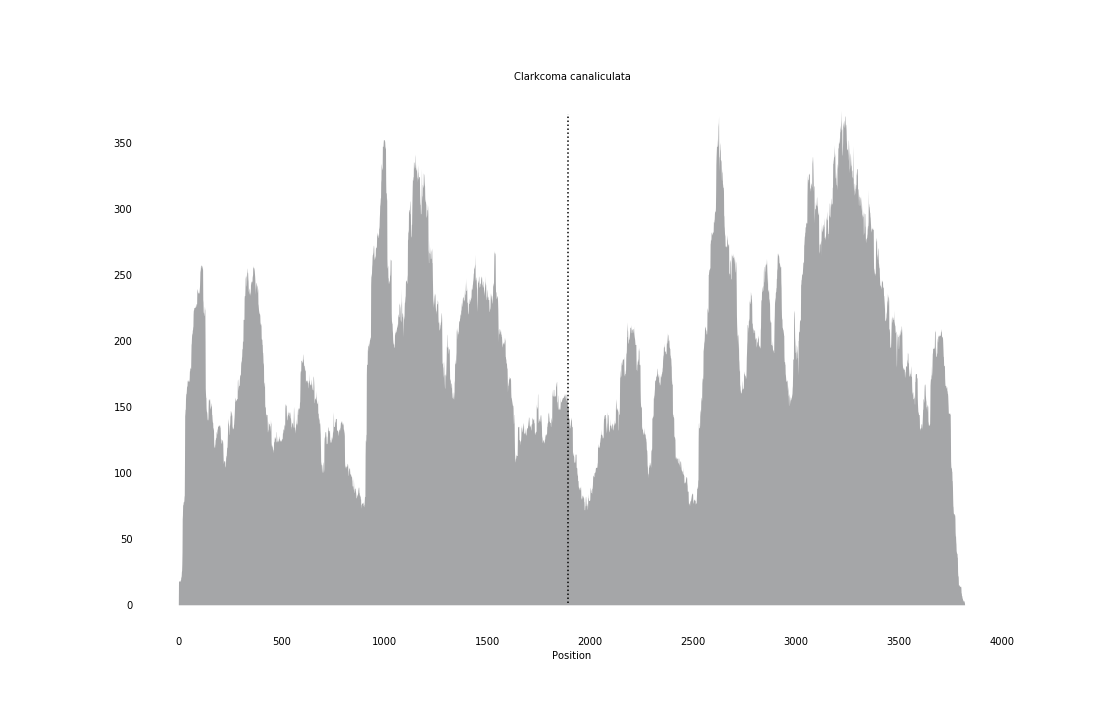

### Cyprinus_carpio_coverage.png

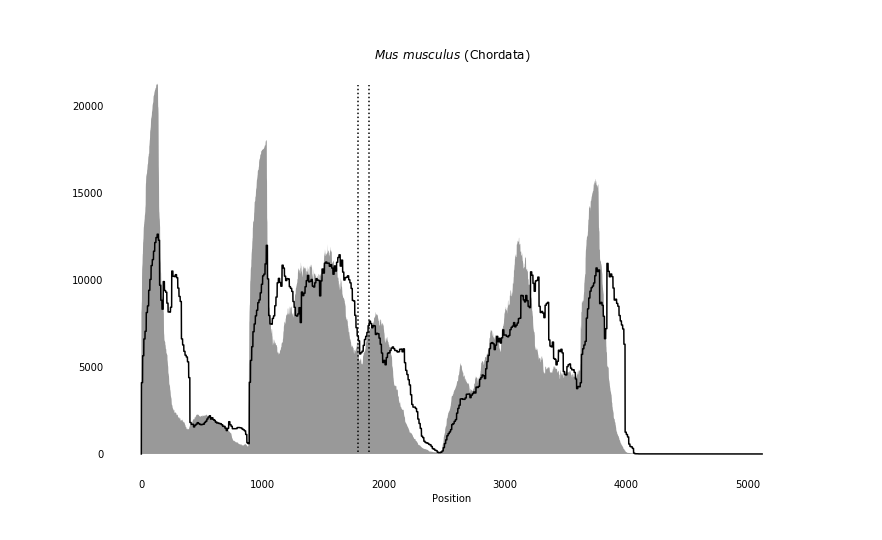

### Danio_rerio_coverage_correct.png

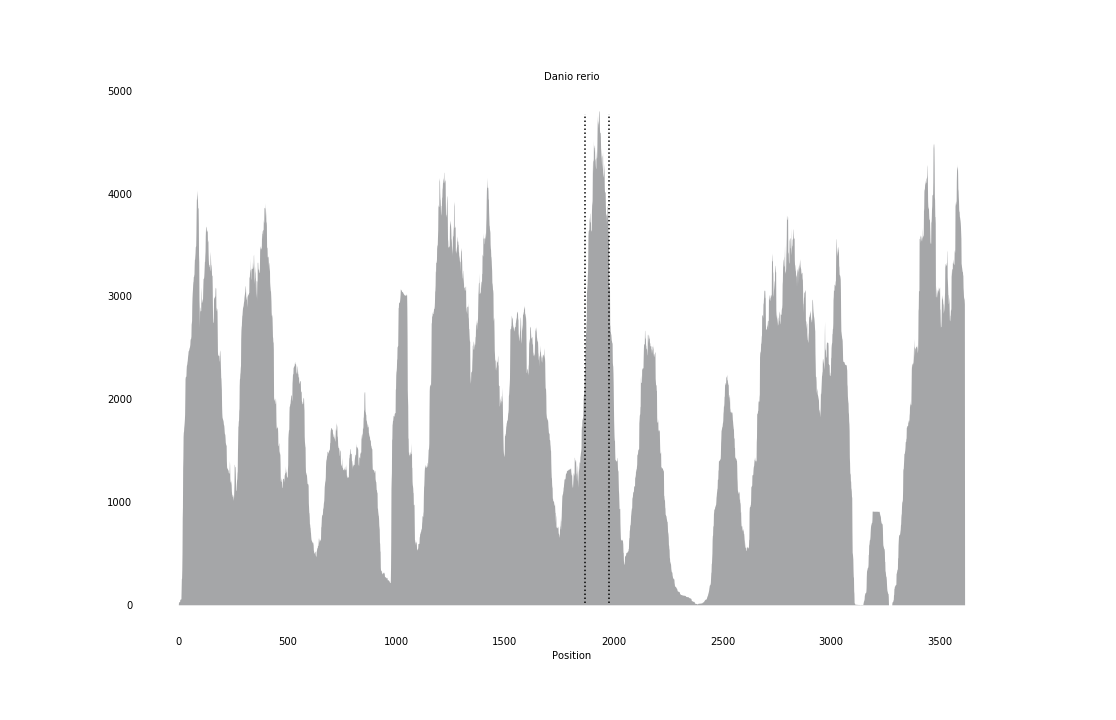

### Dicentrarchus_labrax_coverage_correct.png

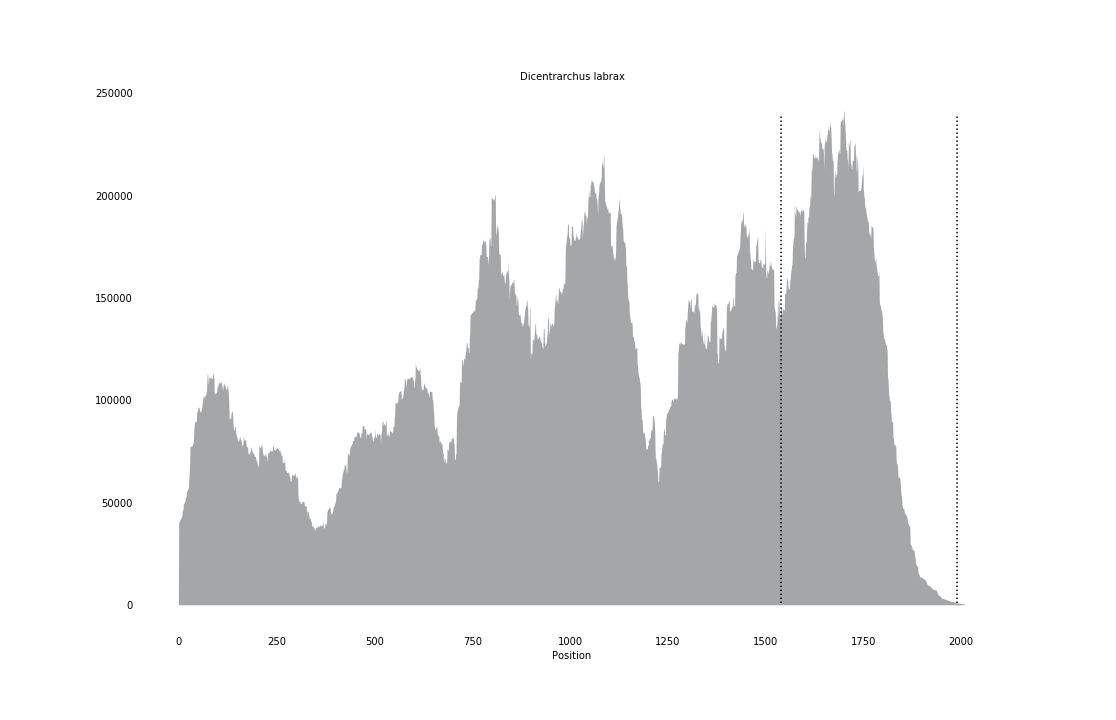

### Dictyocaulus_viviparus_coverage.png

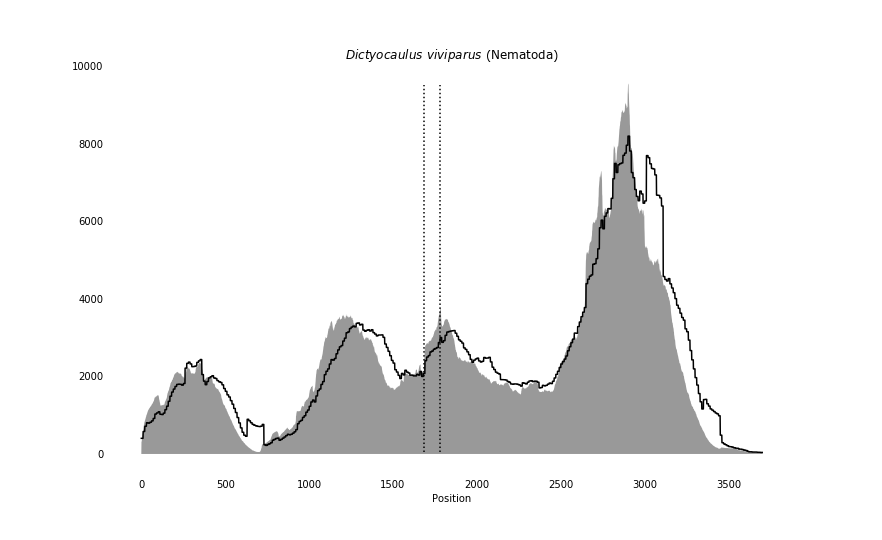

### Dicyema_japonica_coverage.png

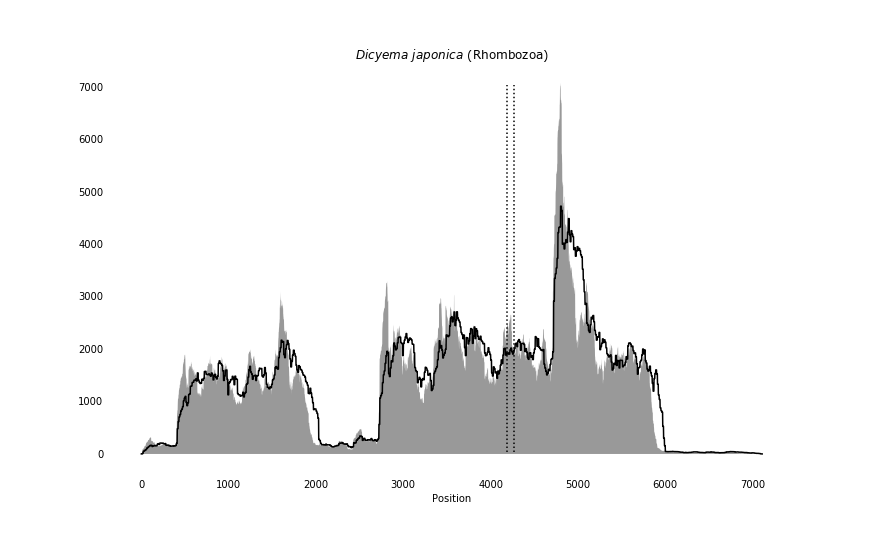

### Diopisthoporus_longitubus_coverage_correct.png

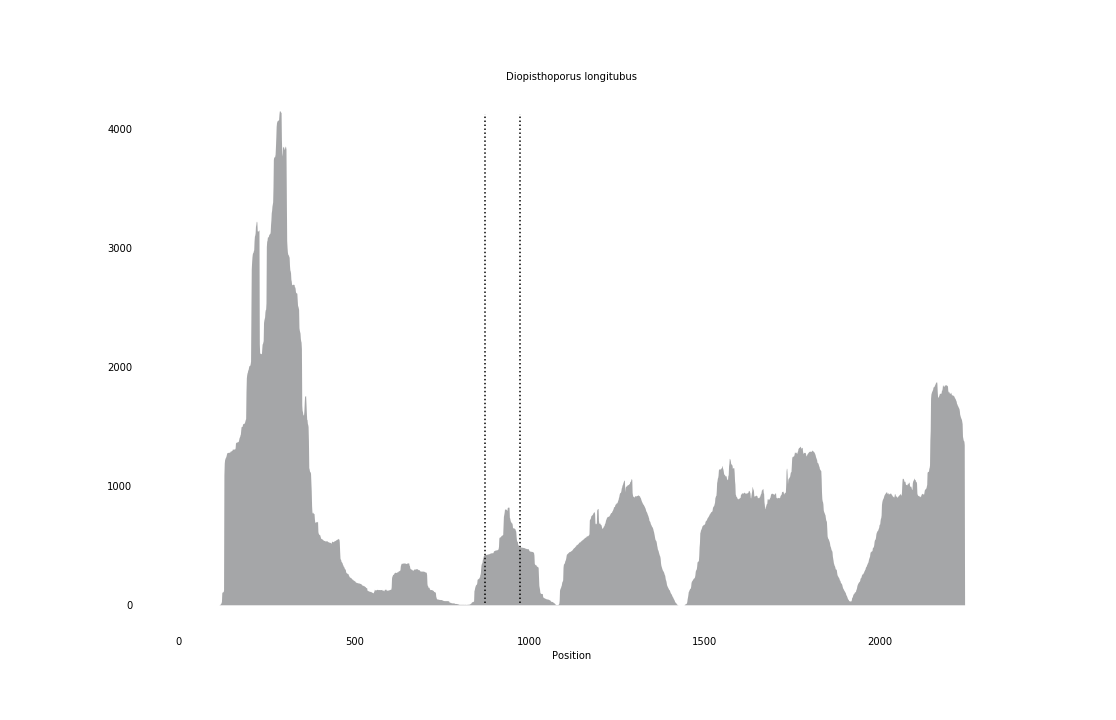

### Dirofilaria_immitis_coverage.png

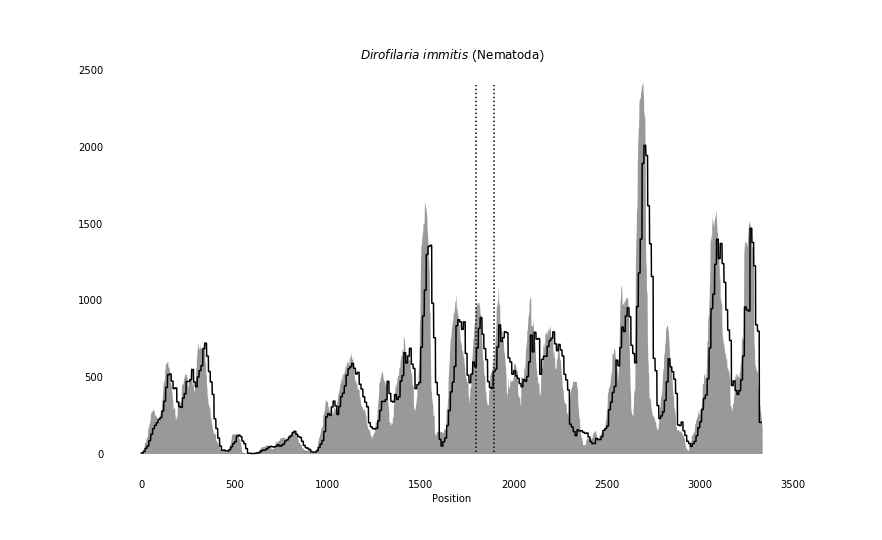

### Erinaceus_europaeus_coverage_correct.png

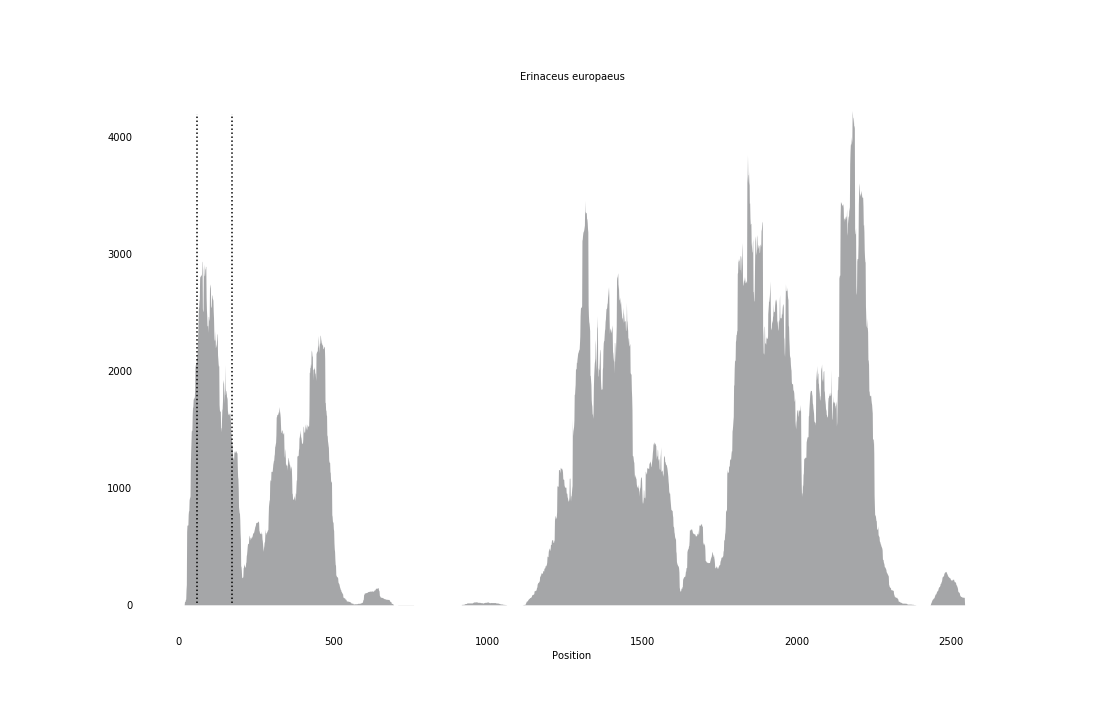

### Gadus_morhua_coverage_correct.png

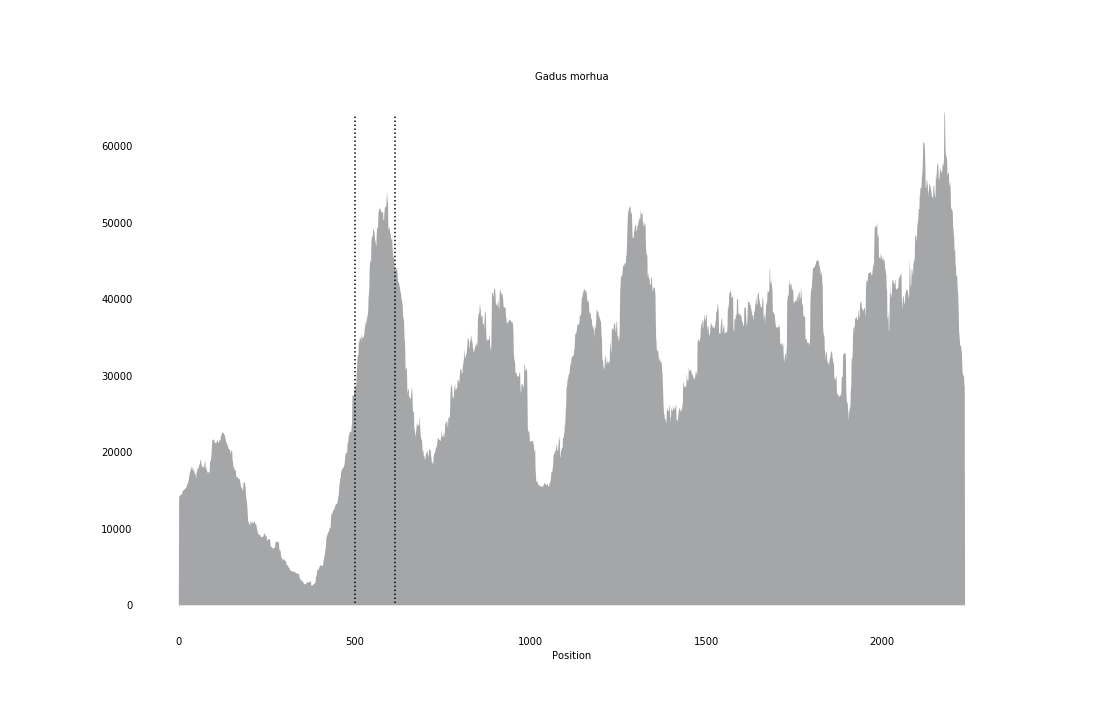

### Gallus_gallus_coverage_correct.png

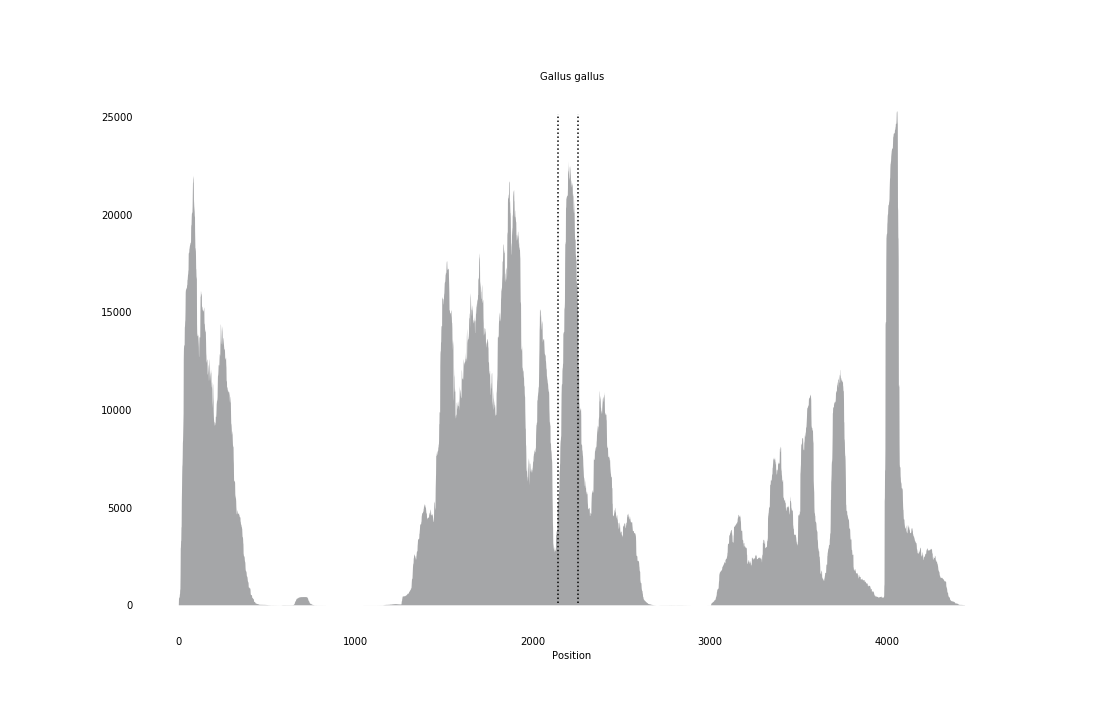

### Glycera_tridactyla_coverage.png

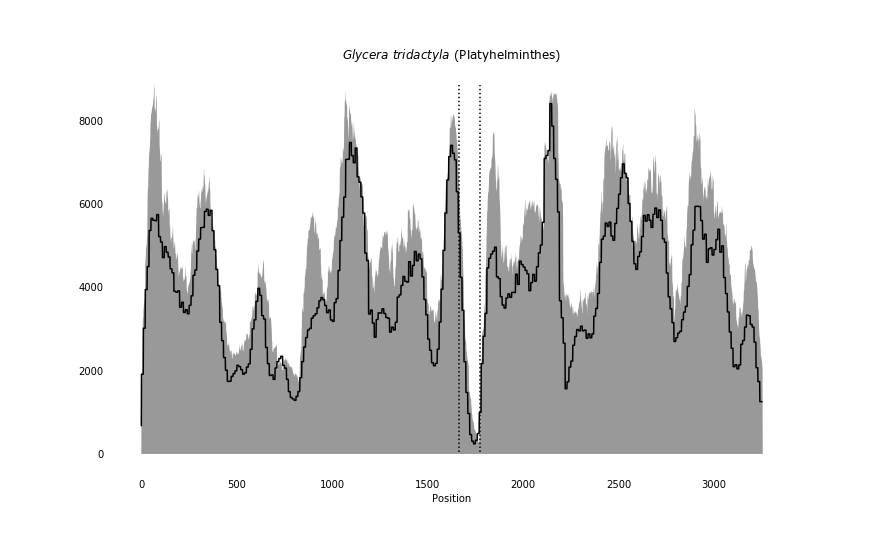

### Gorgonocephalus_pustulatum_coverage.png

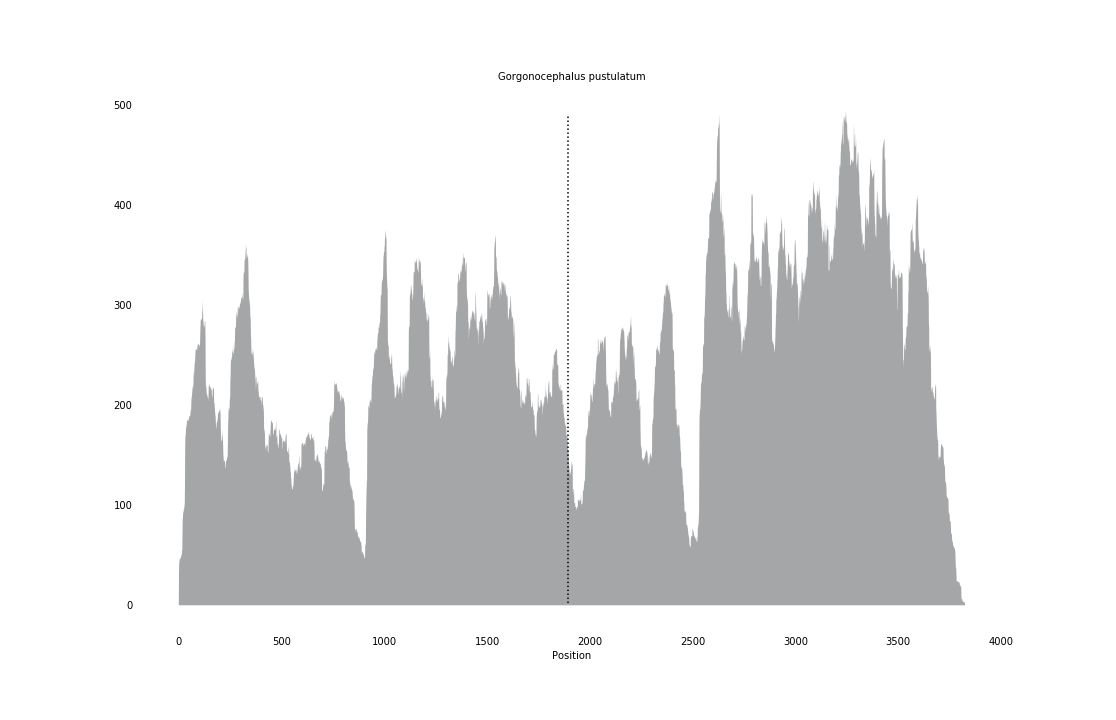

### Gorilla_gorilla_coverage_correct.png

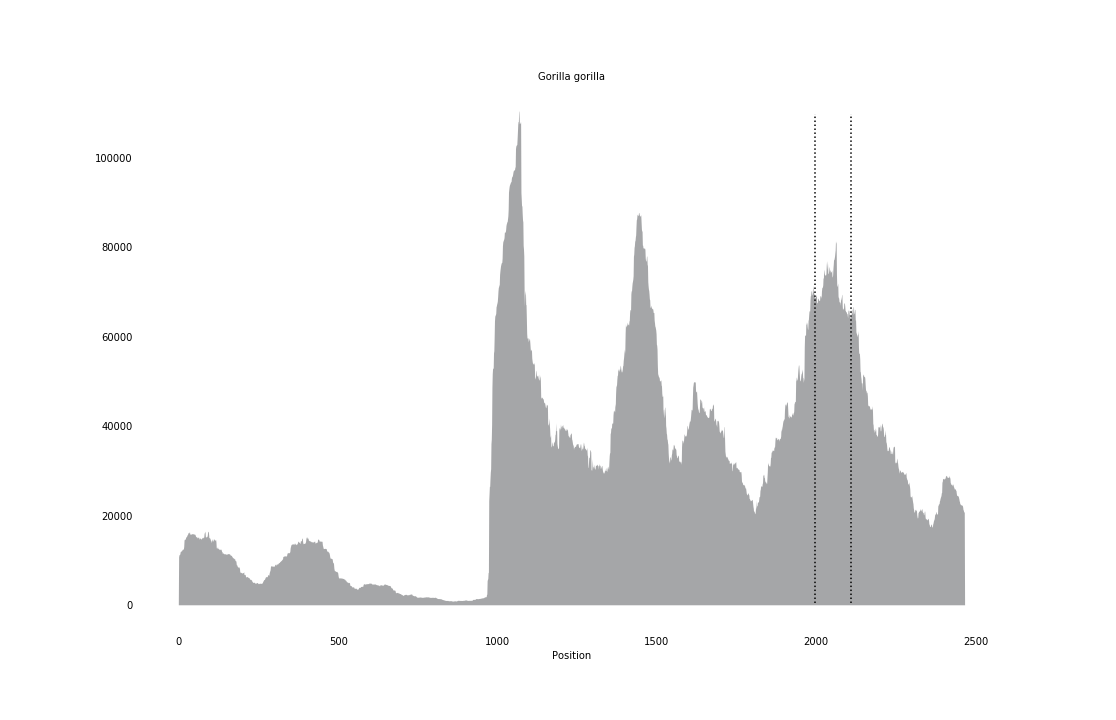

### Haemadipsa_cavatuses_coverage_correct.png

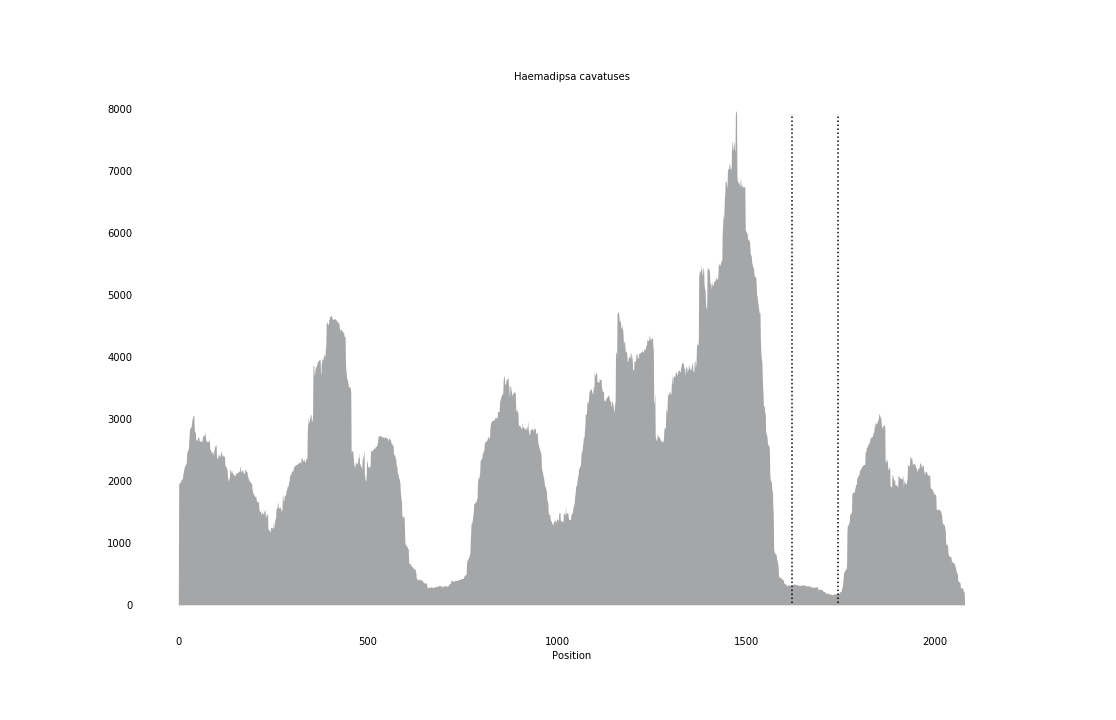

### Halicryptus_spinulosus_coverage.png

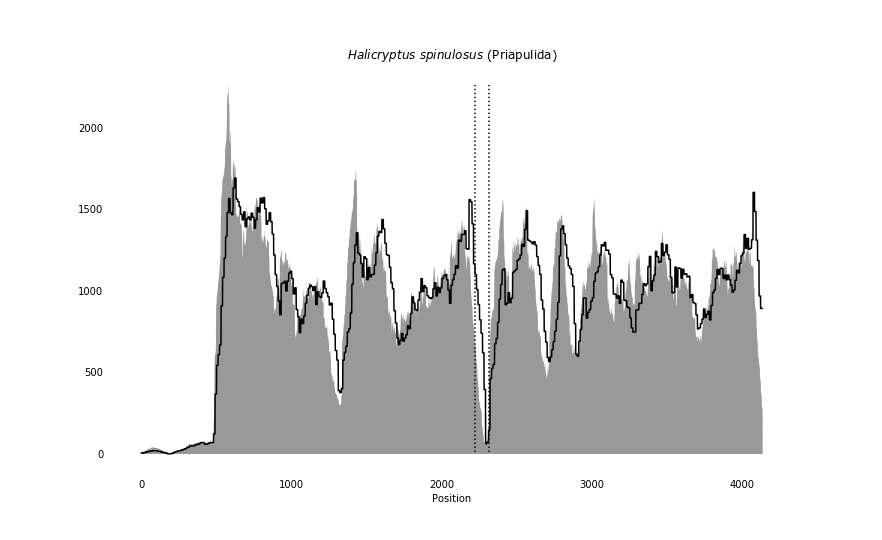

### Heligmosomoides_polygyrus_coverage_correct.png

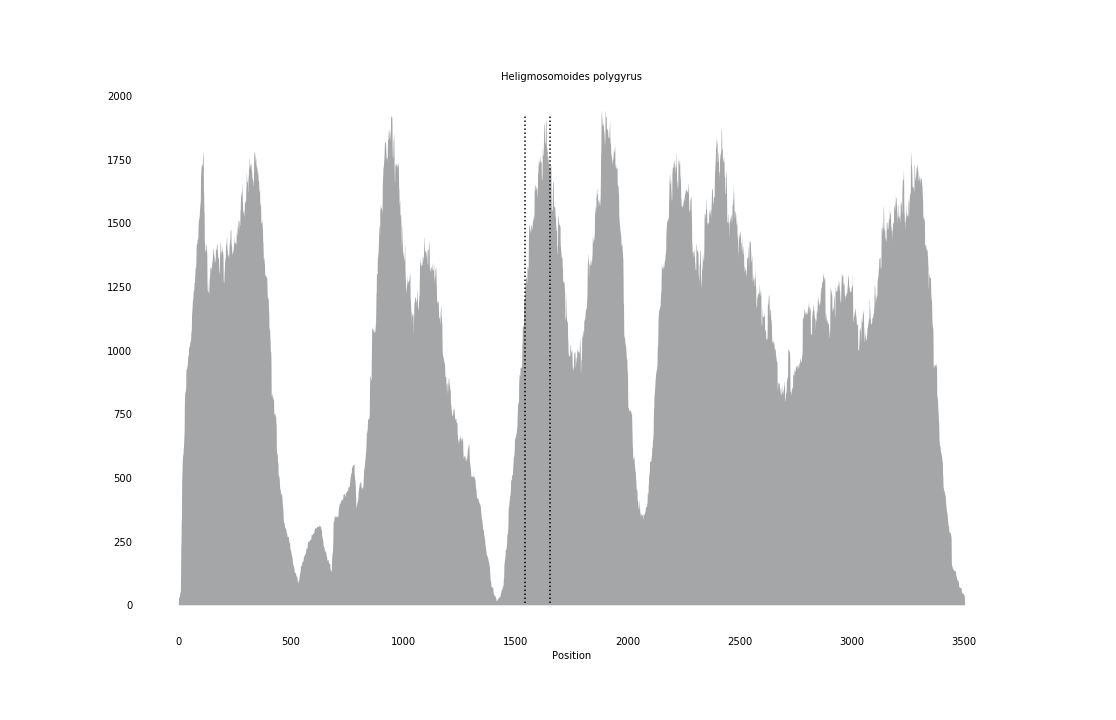

### Hypsibius_dujardini_coverage.png

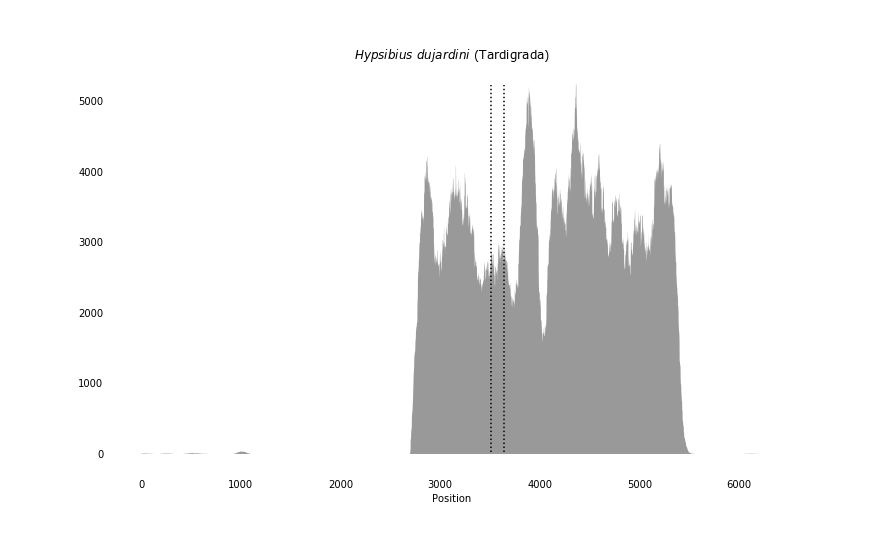

### Intoshia_linei_coverage.png

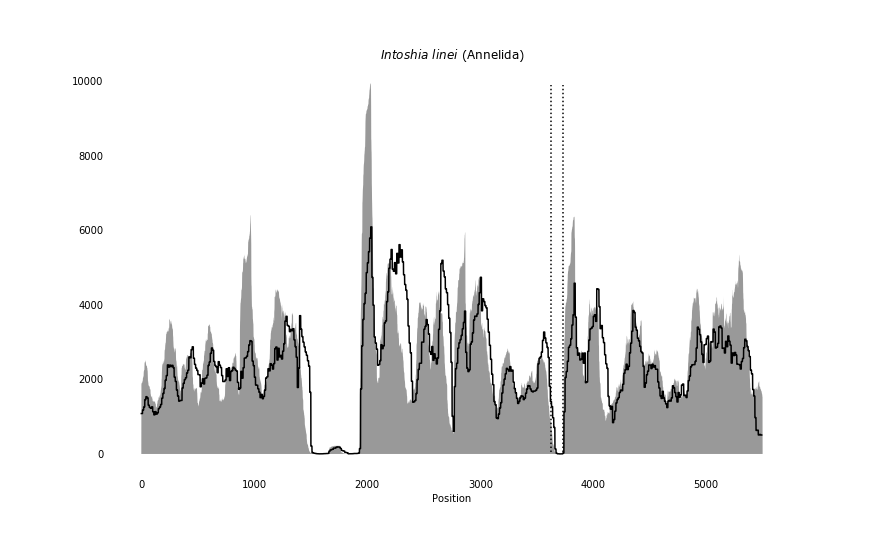

### Lumbricus_terrestris_coverage_correct.png

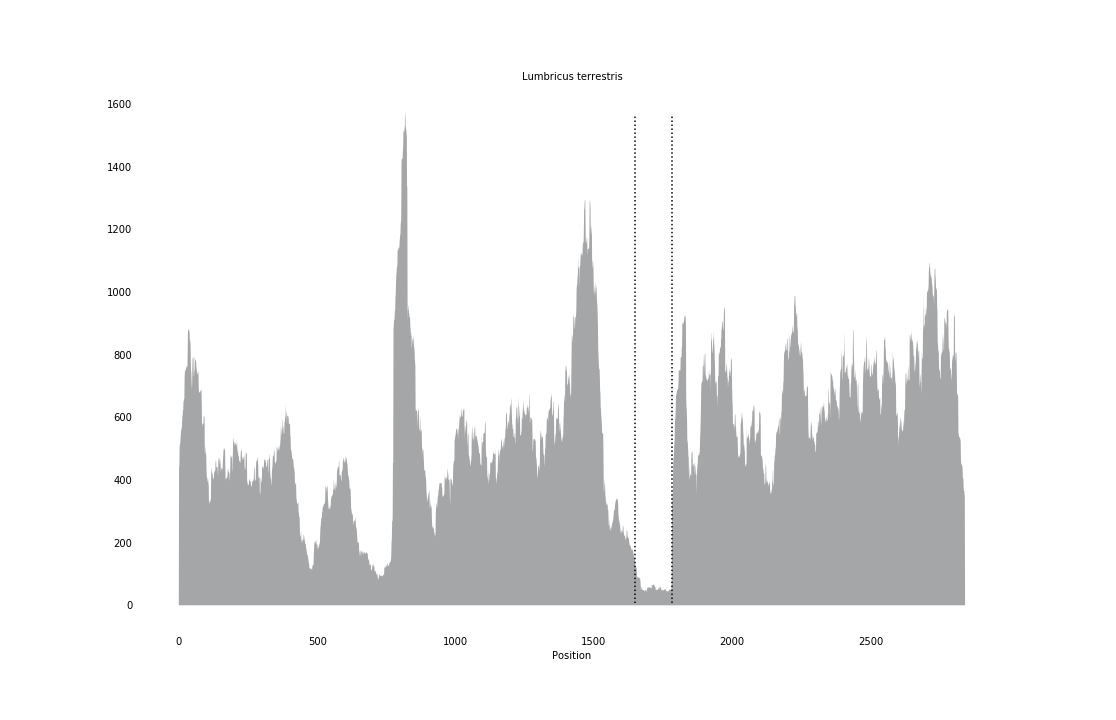

### Meara_stichopi_coverage.png

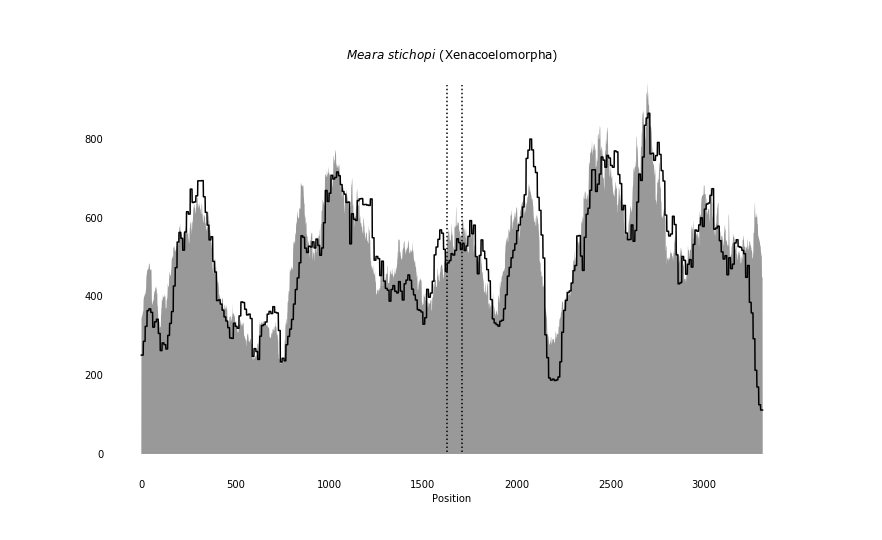

### Meloidogyne_incognita_coverage_correct.png

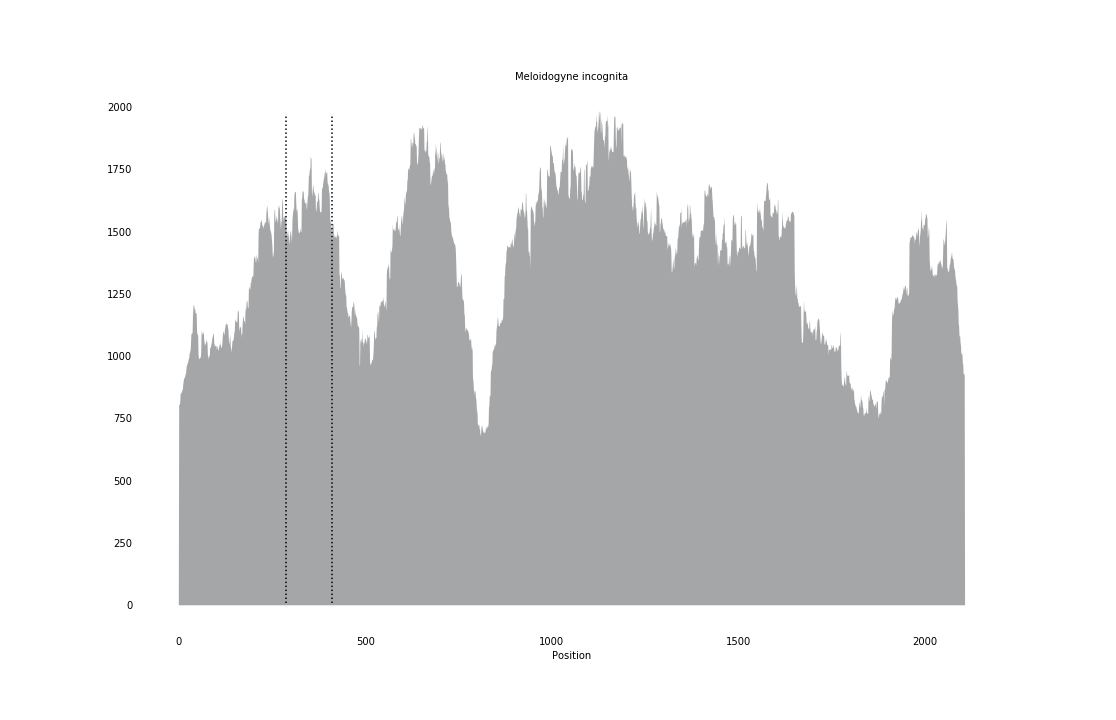

### Membranipora_membranacea_coverage.png

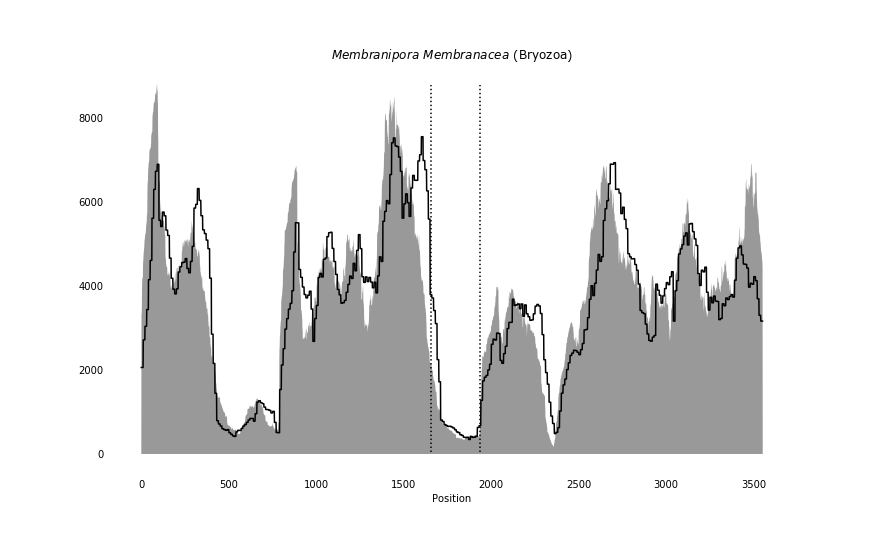

### Menidia_menidia_coverage_correct.png

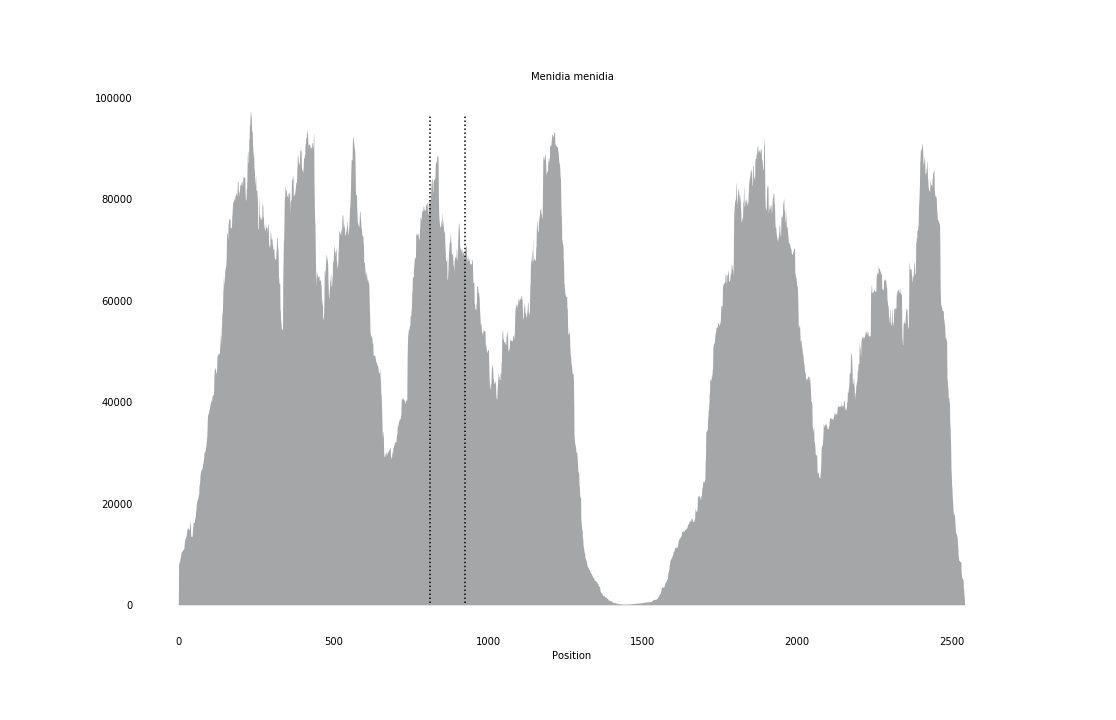

### Mesodasys_laticaudatus_coverage.png

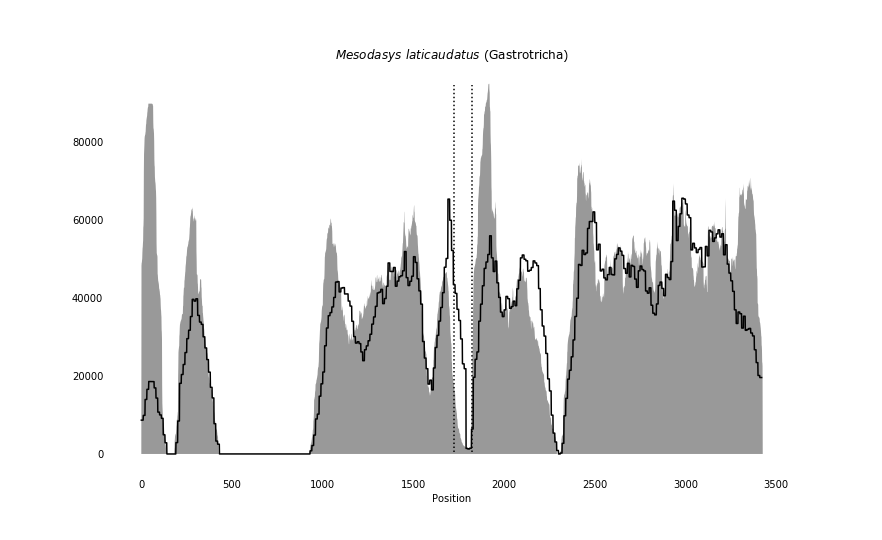

### Misgurnus_anguillicaudatus_coverage_correct.png

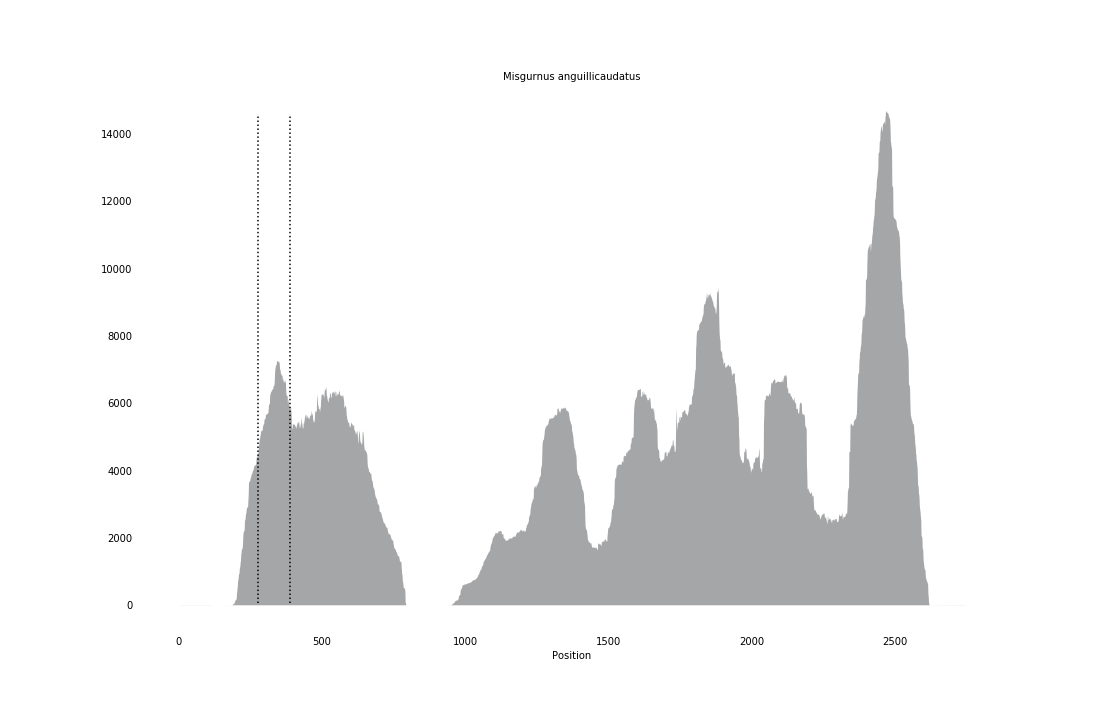

### Mus_musculus_coverage.png

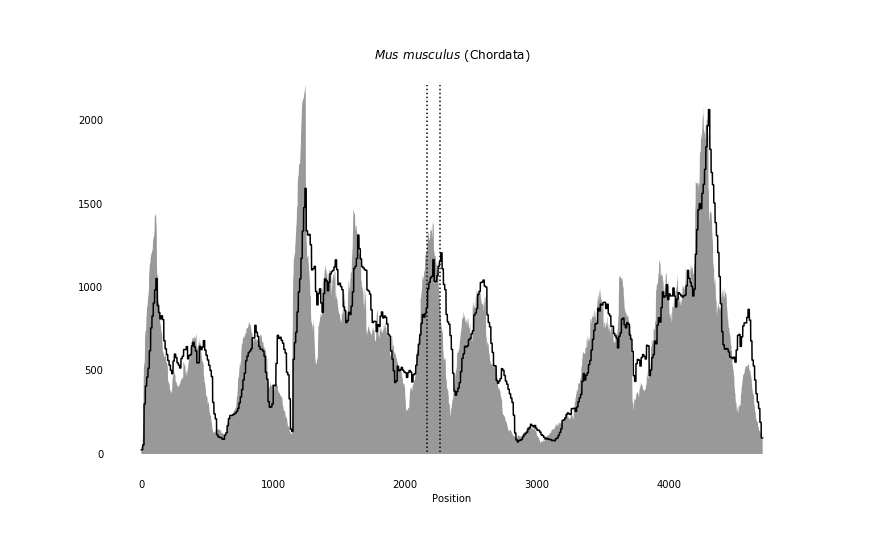

### Mya_arenaria_coverage_correct.png

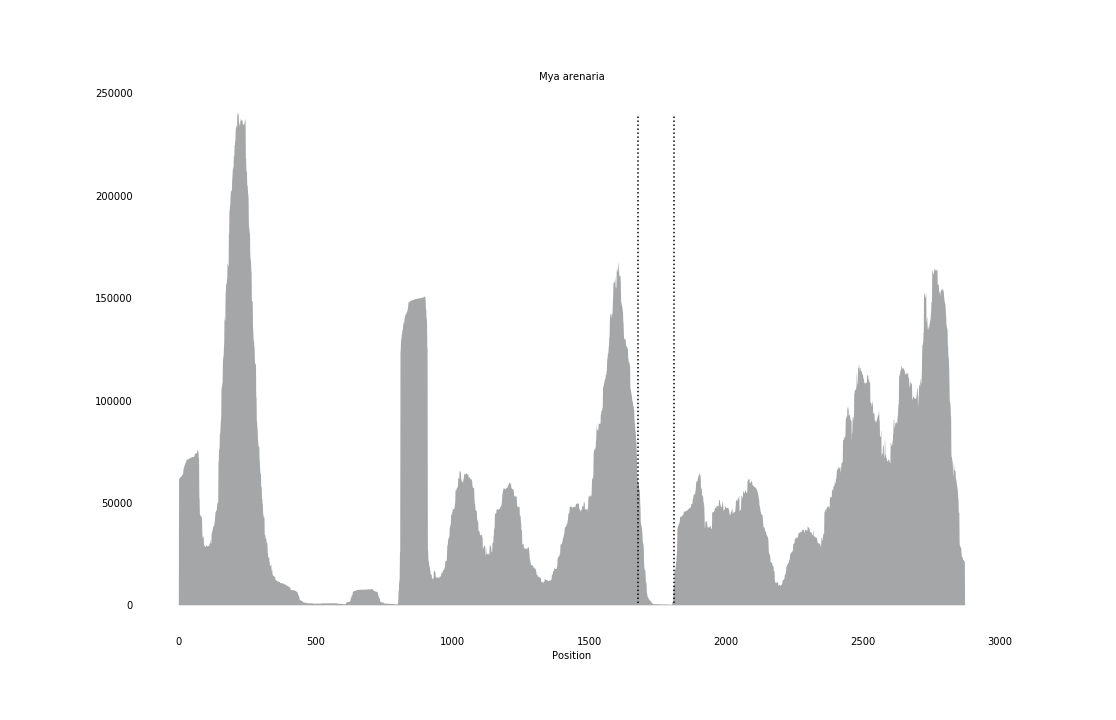

### Necator_americanus_coverage_correct.png

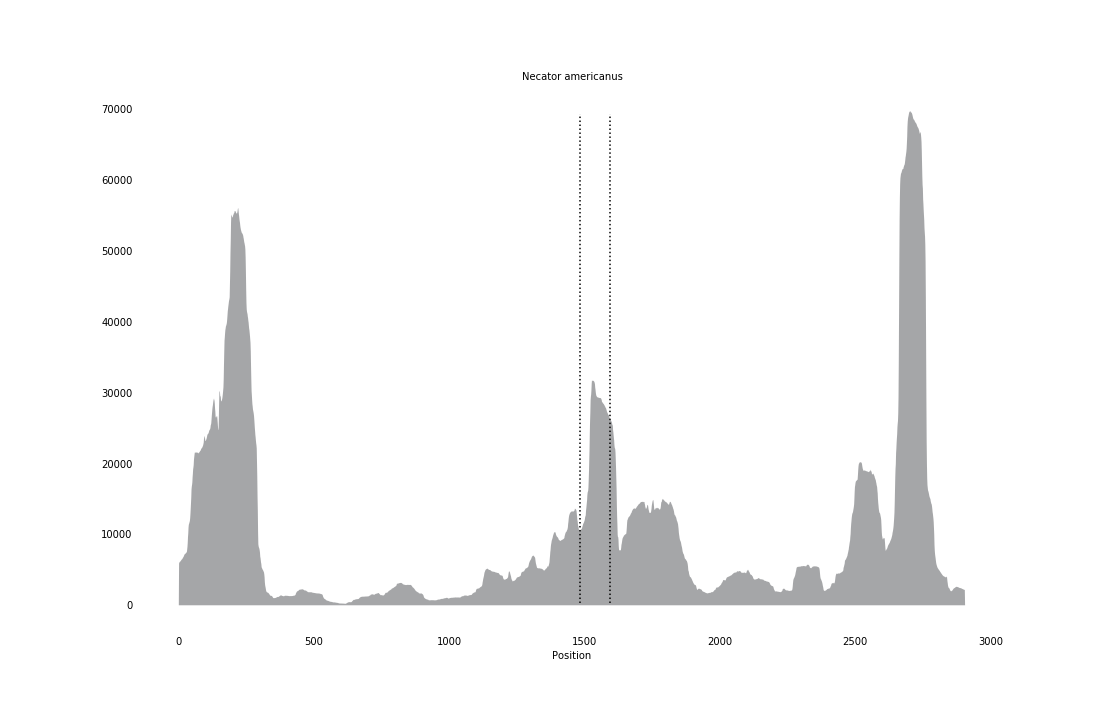

### Nemertoderma_westbladi_coverage.png

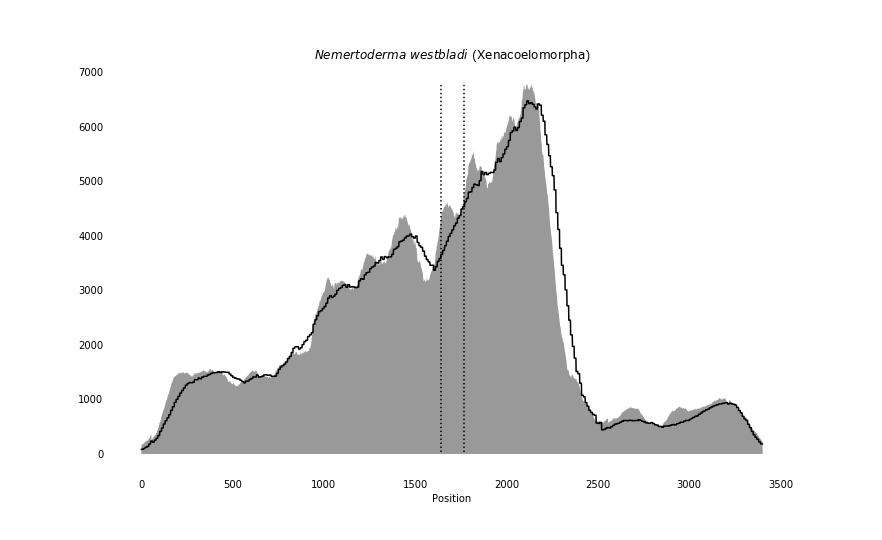

### Ophiactis_abyssicola_coverage.png

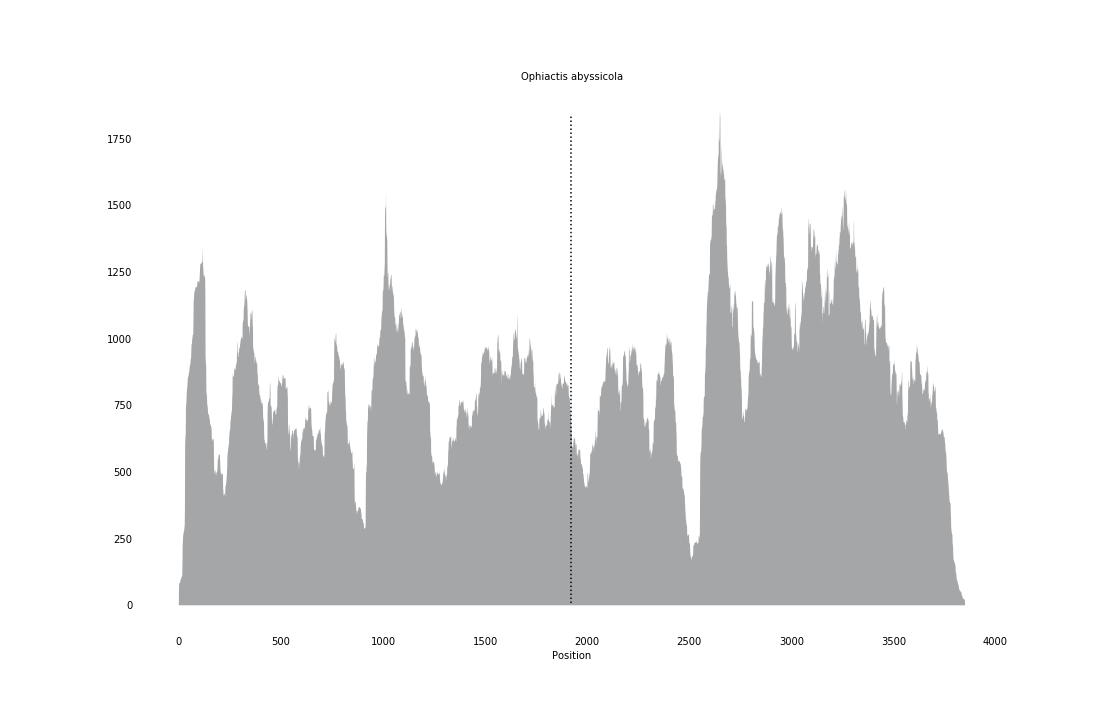

### Ophiactis_resiliens_coverage.png

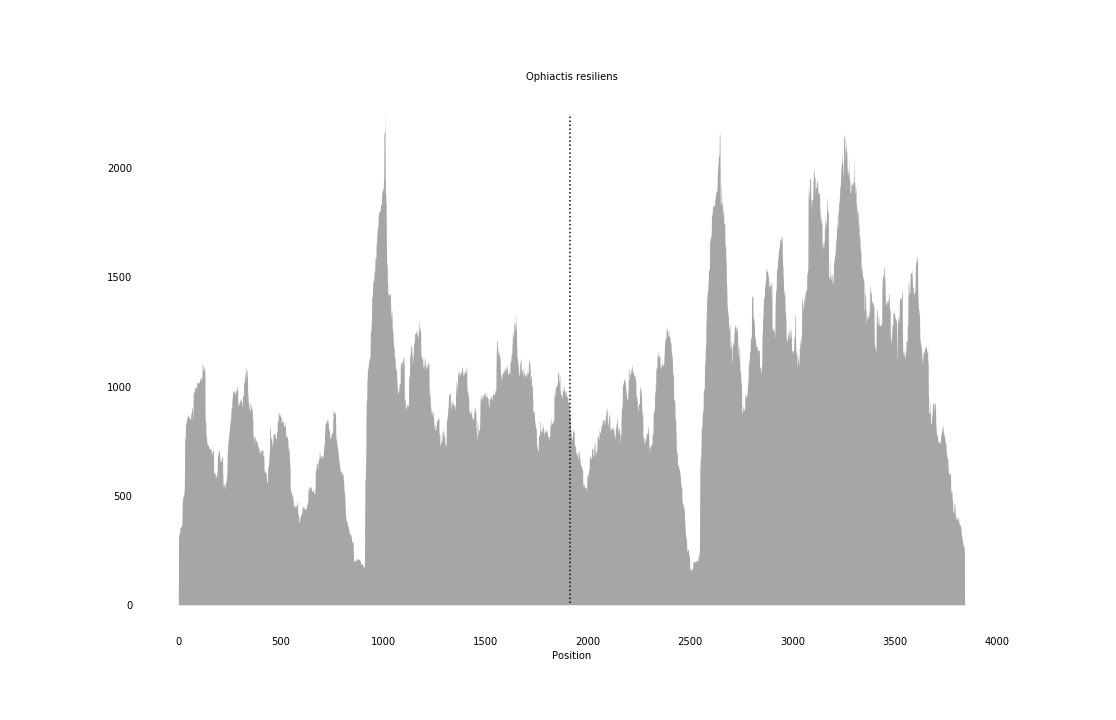

### Ophiernus_vallincola_coverage.png

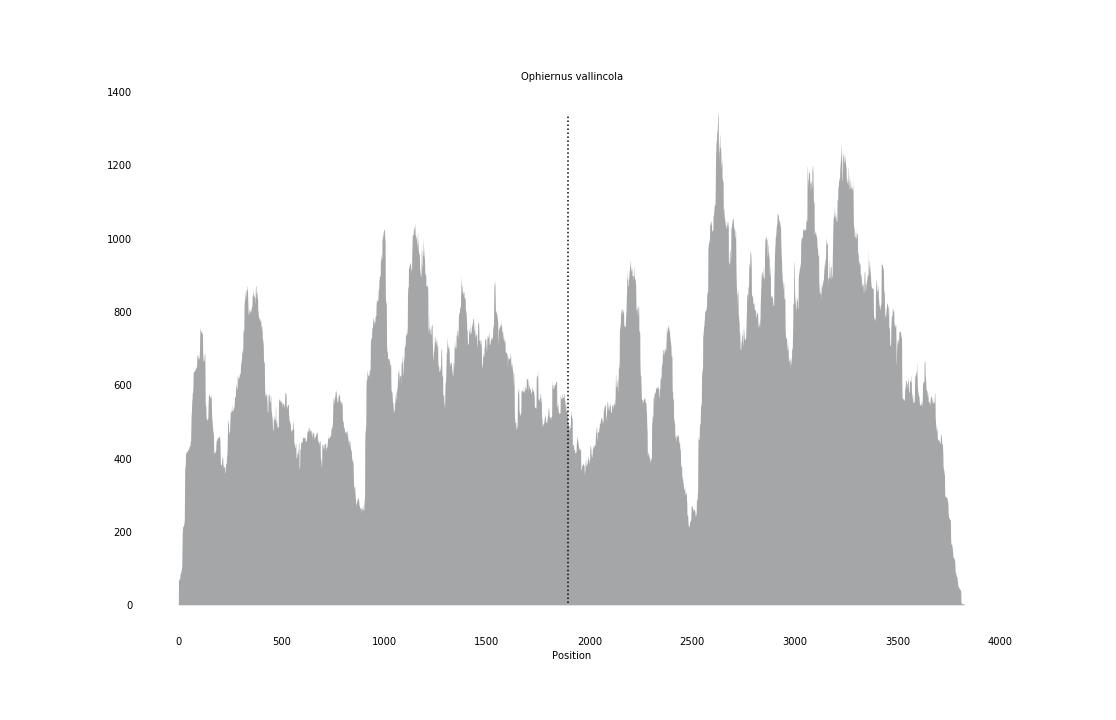

### Ophioceres_bispinosus_coverage.png

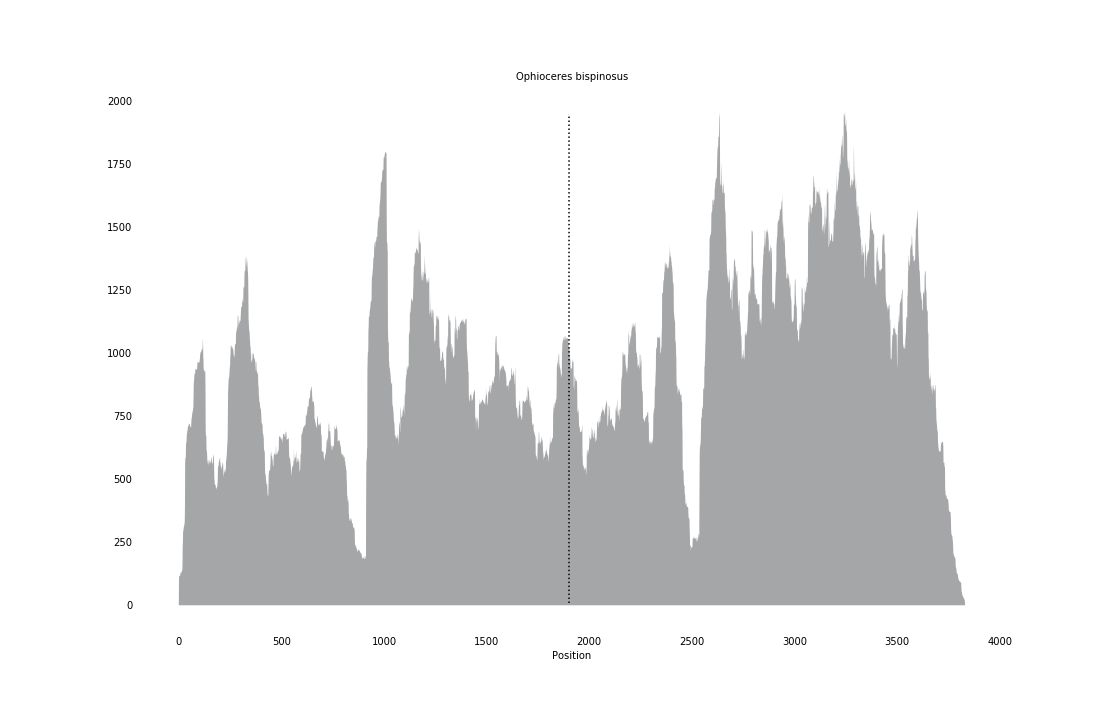

### Ophiolepis_impressa_coverage.png

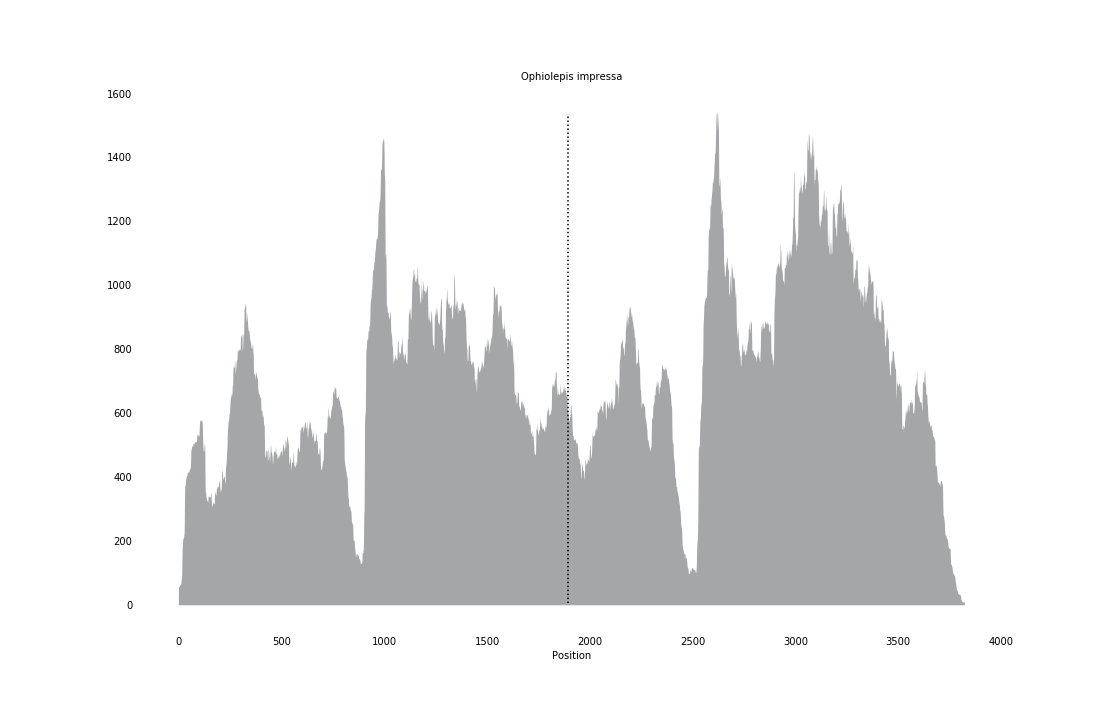

### Ophioleuce_brevispinum_coverage.png

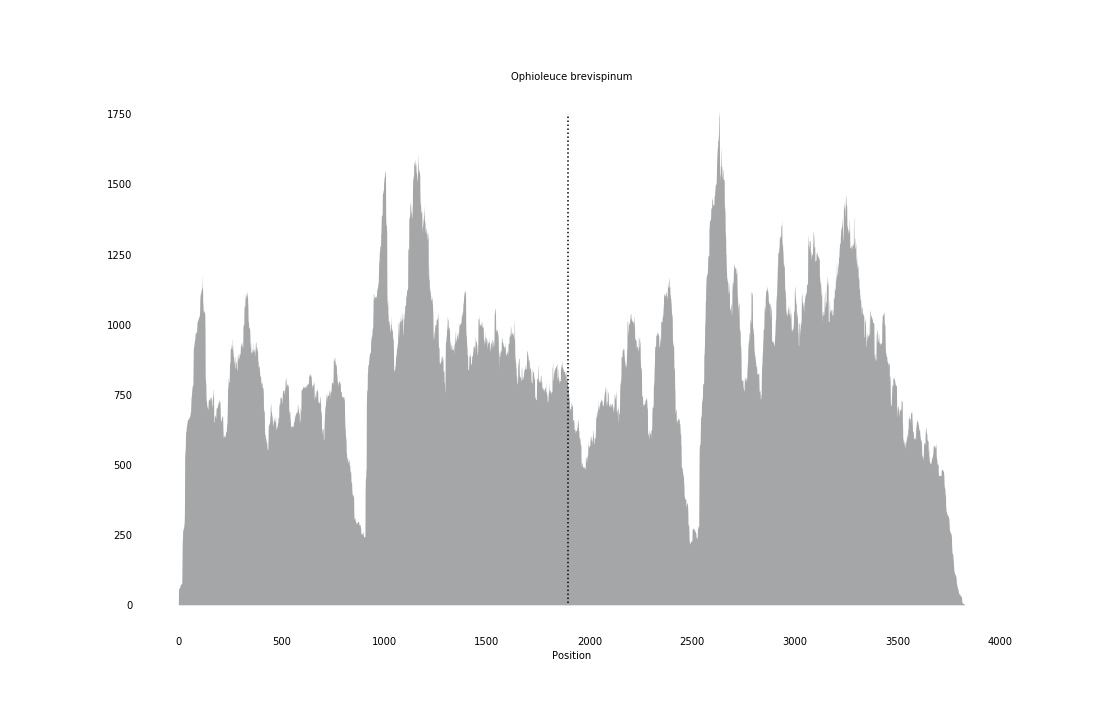

### Ophiolimna_perfida_coverage.png

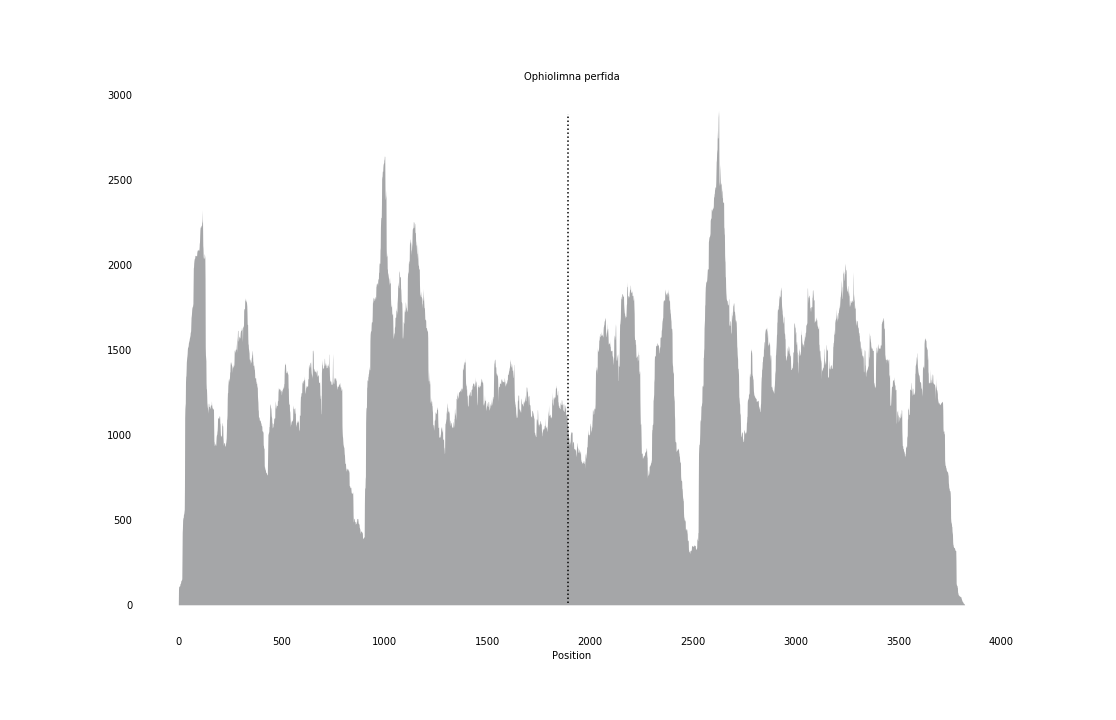

### Ophiologimus_prolifer_coverage.png

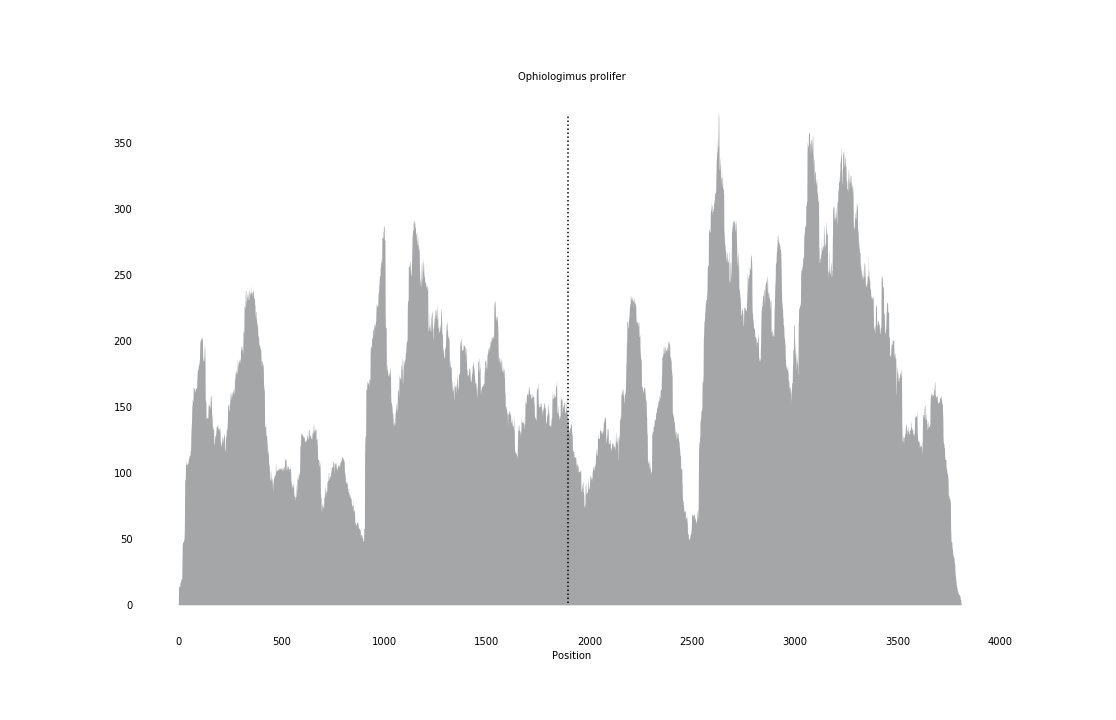

### Ophiomoeris_obstricta_coverage.png

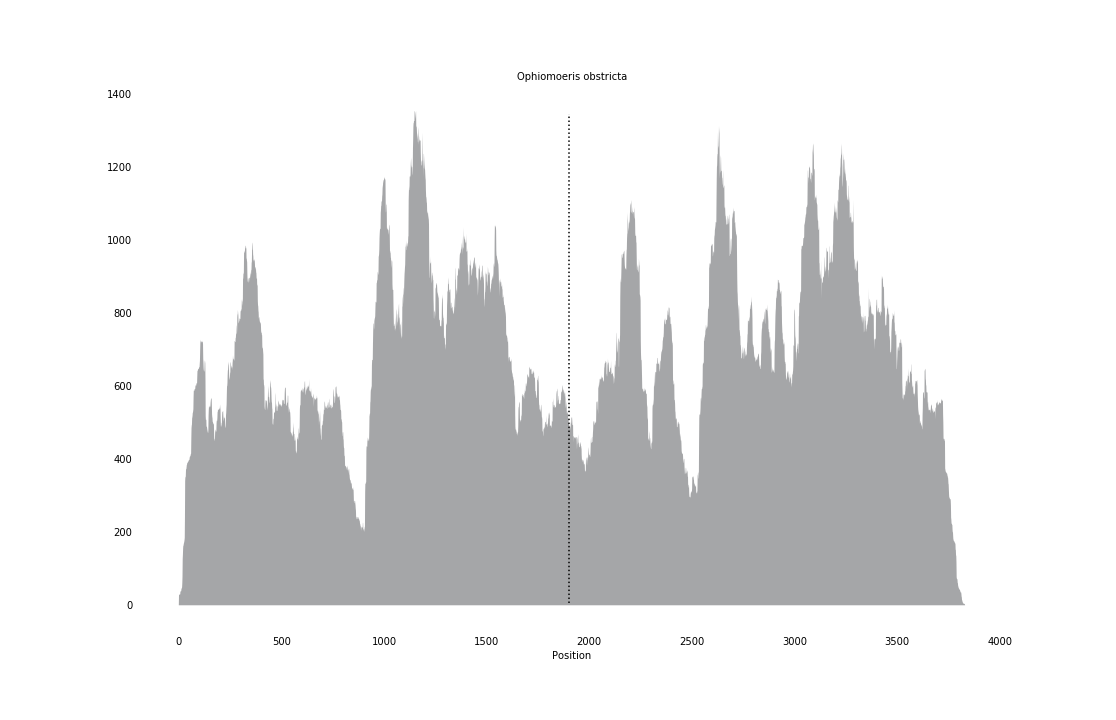

### Ophiomusium_lymani_coverage.png

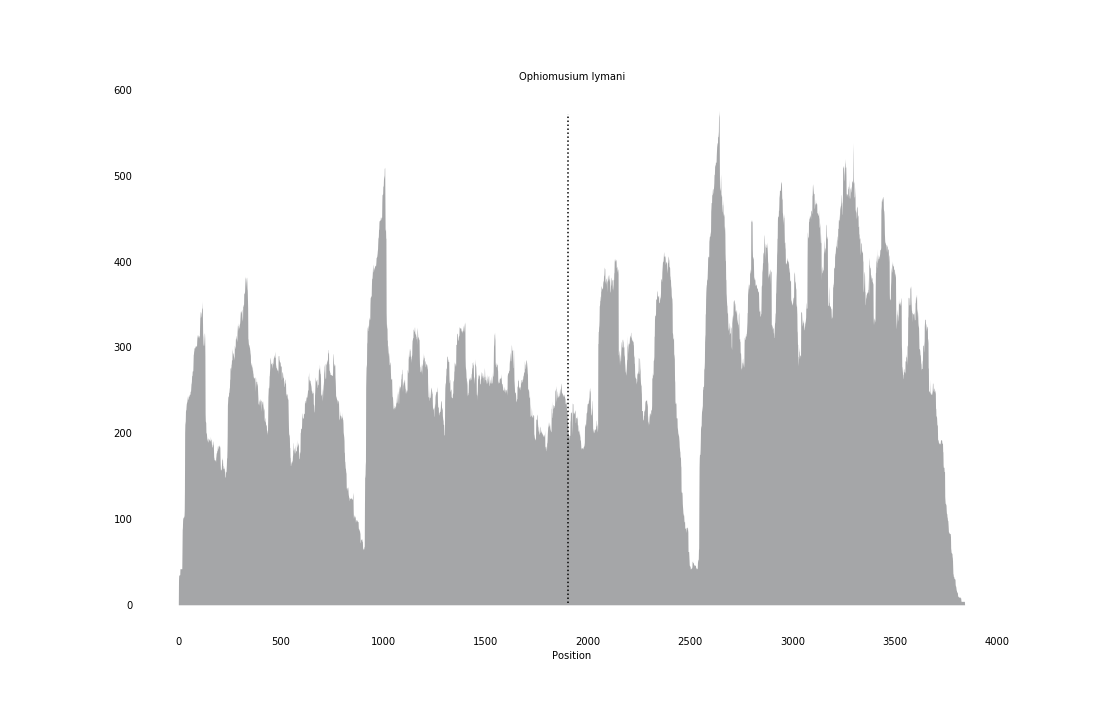

### Ophionereis_schayeri_coverage.png

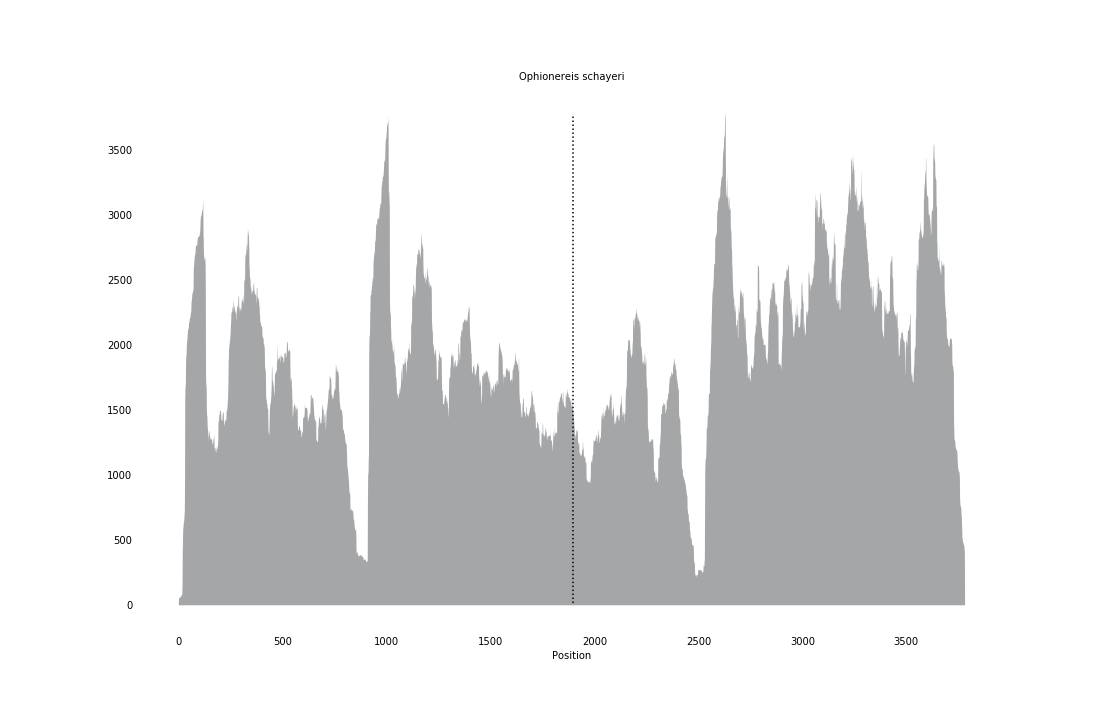

### Ophiopeza_cylindrica_coverage.png

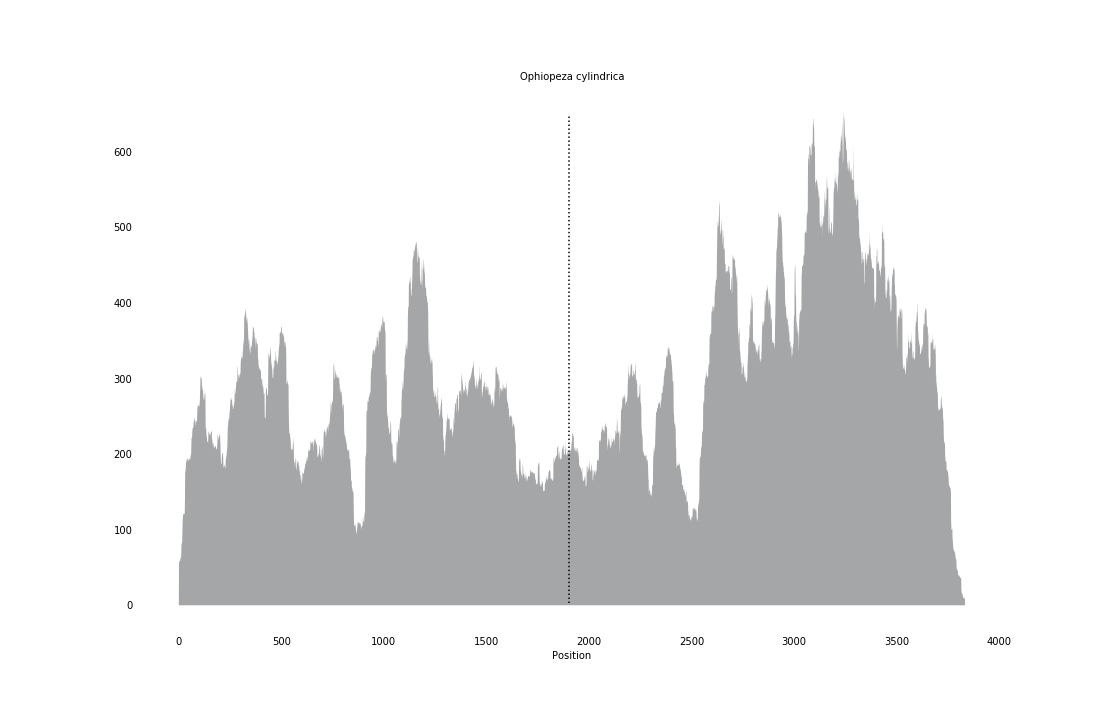

### Ophiophragmus_wurdemanii_coverage.png

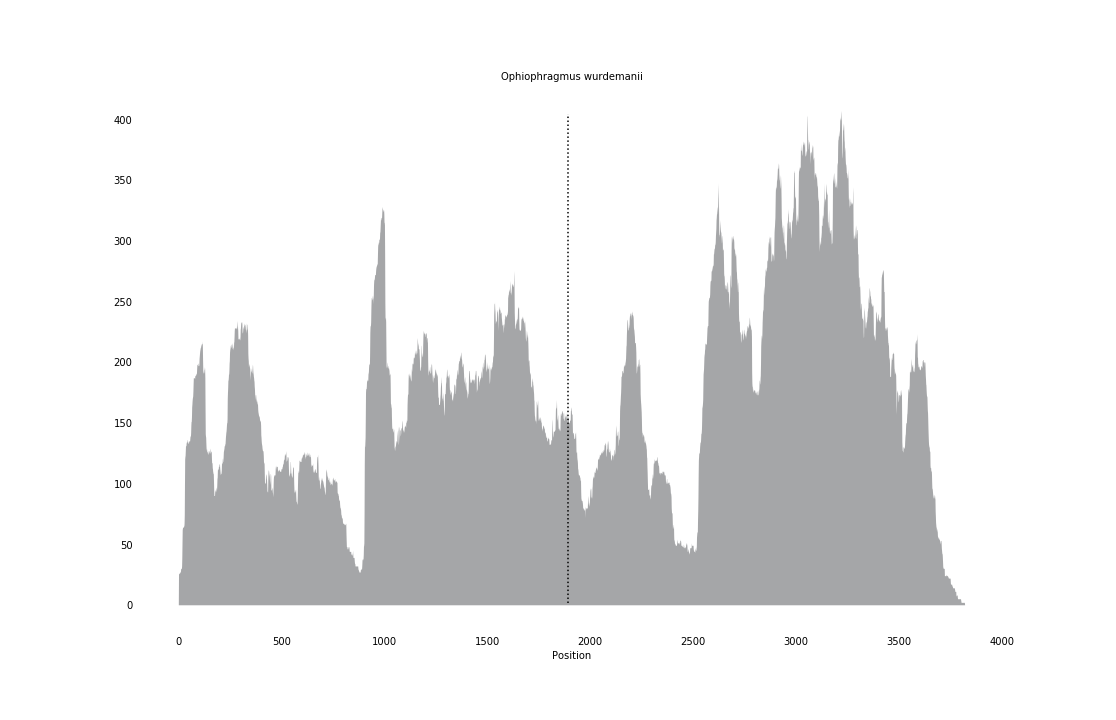

### Ophiophrura_liodisca_coverage.png

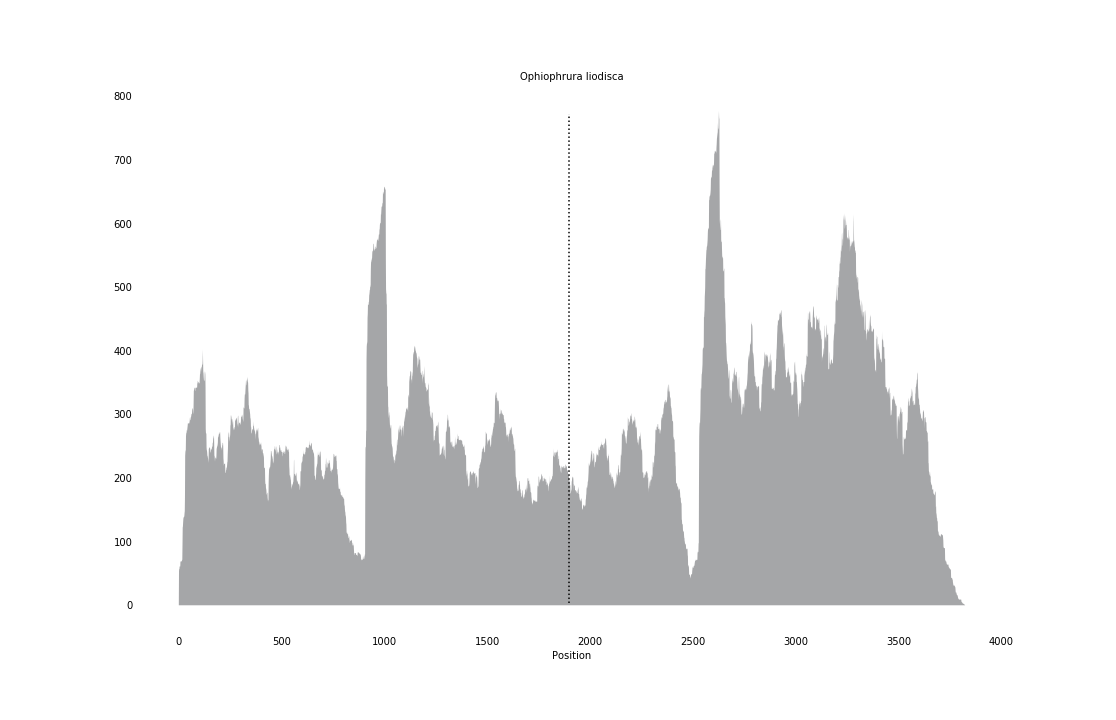

### Ophiophycis_johni_coverage.png

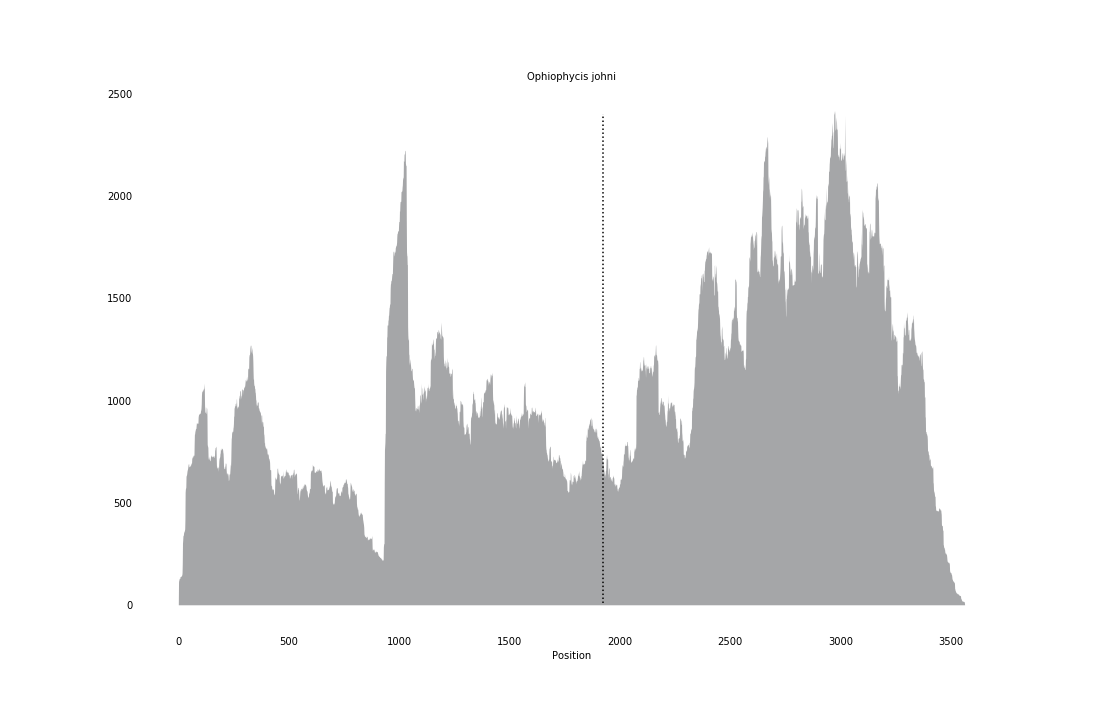

### Ophiothrix_angulata_coverage.png

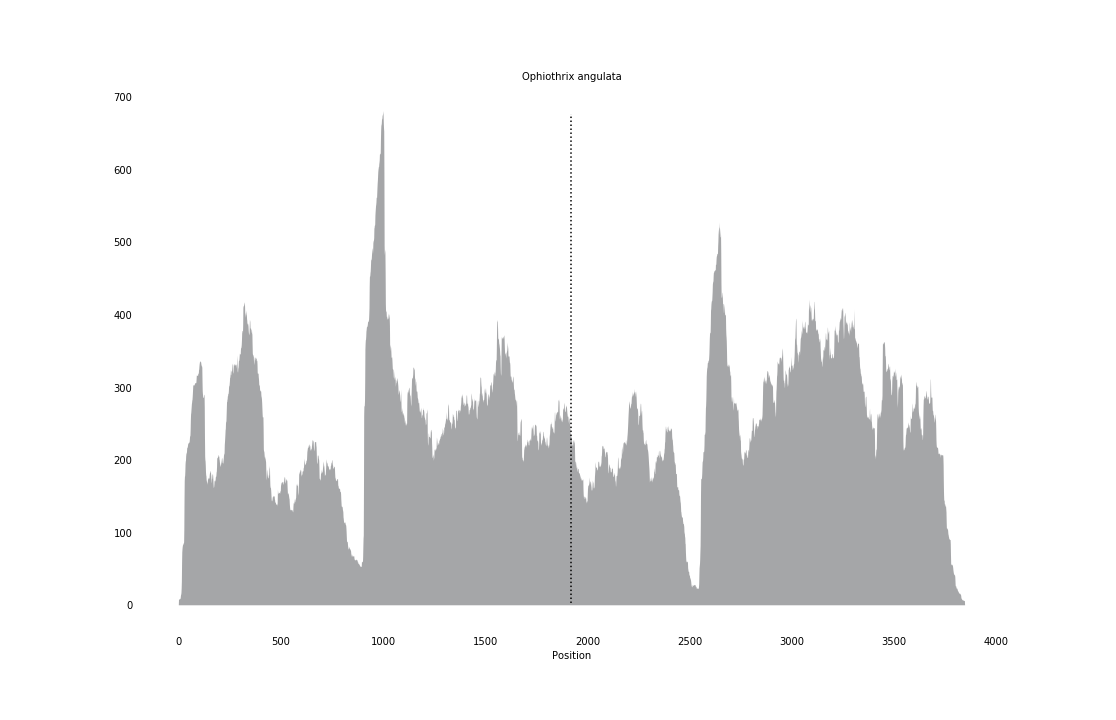

### Ophiothrix_caespitosa_coverage.png

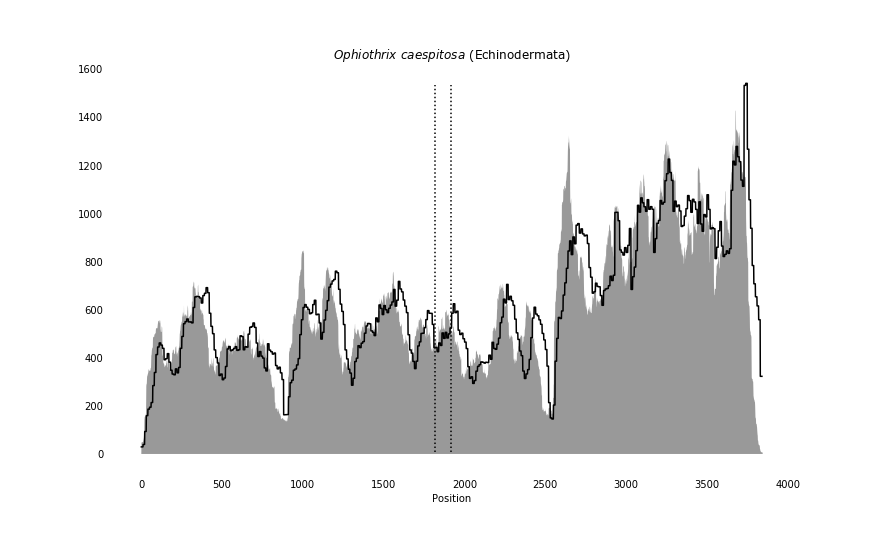

### Oreochromis_aureus_coverage.png

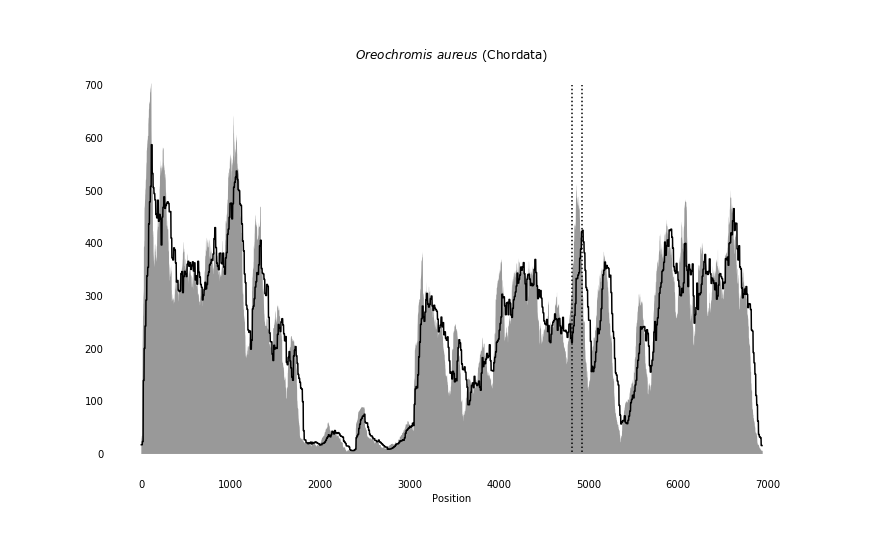

### Oreochromis_niloticus_coverage_correct.png

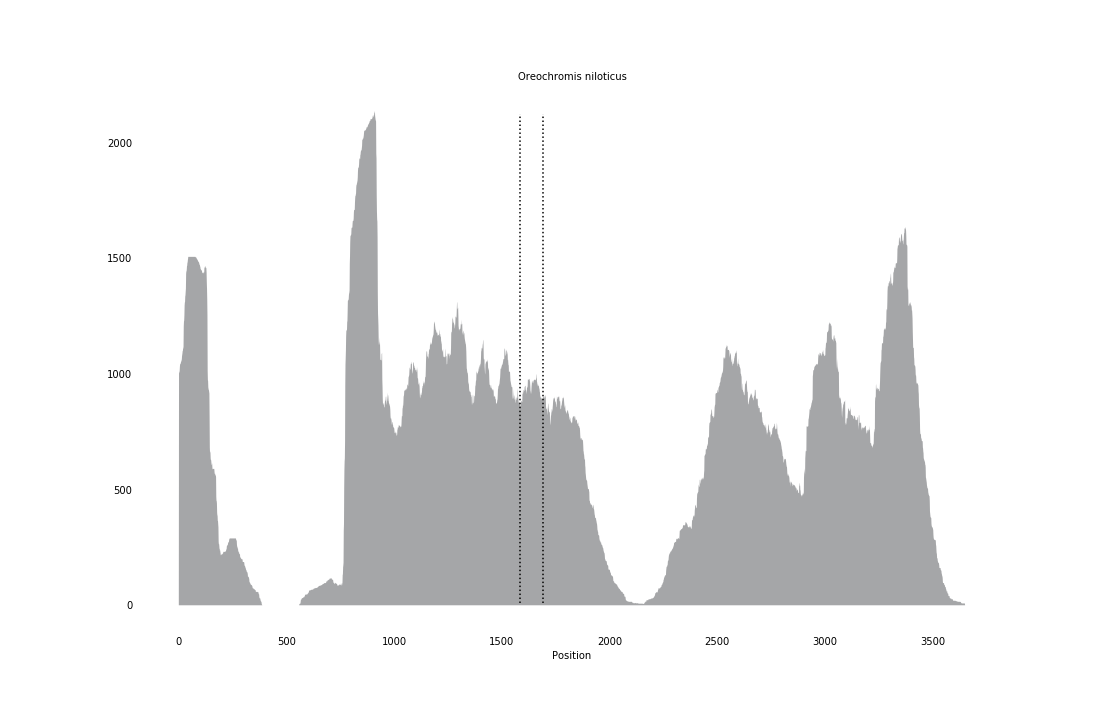

### Oryzias_melastigma_coverage_correct.png

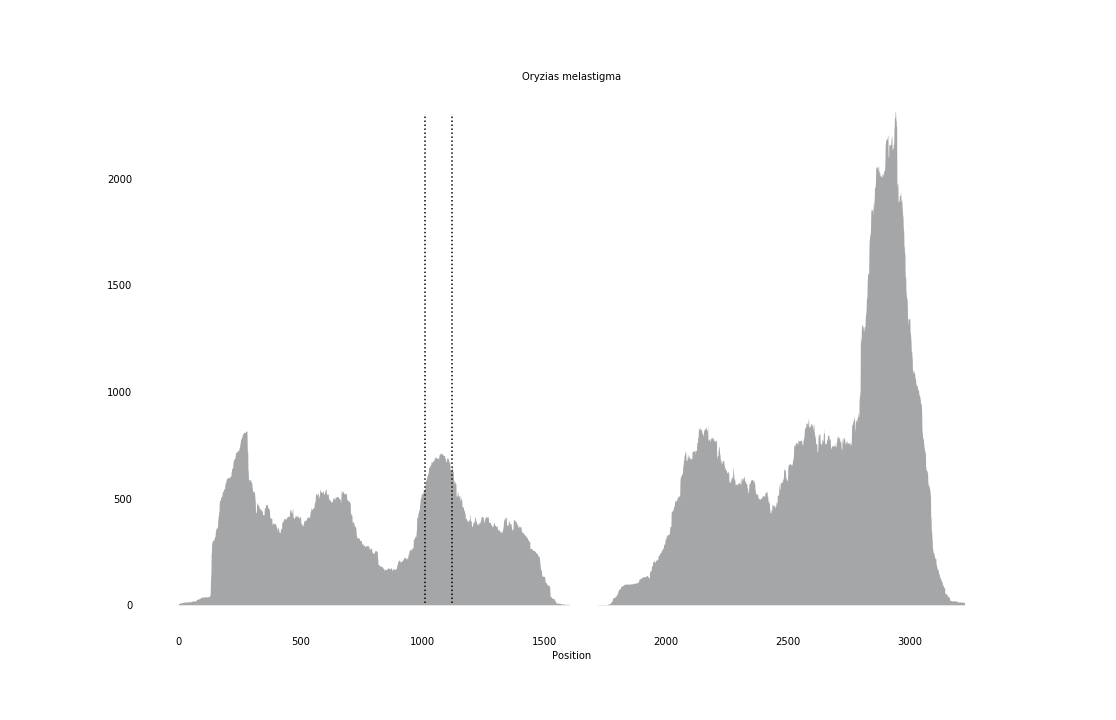

### Owenia_fusiformis_coverage_correct.png

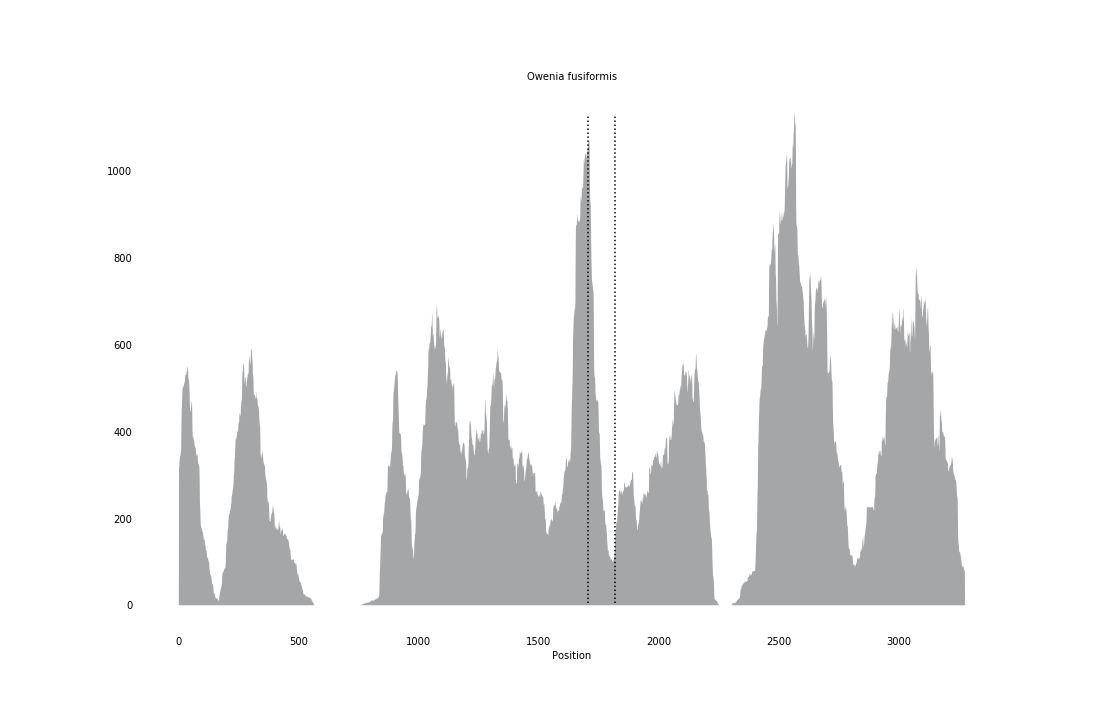

### Panagrellus_redivivus_coverage_correct.png

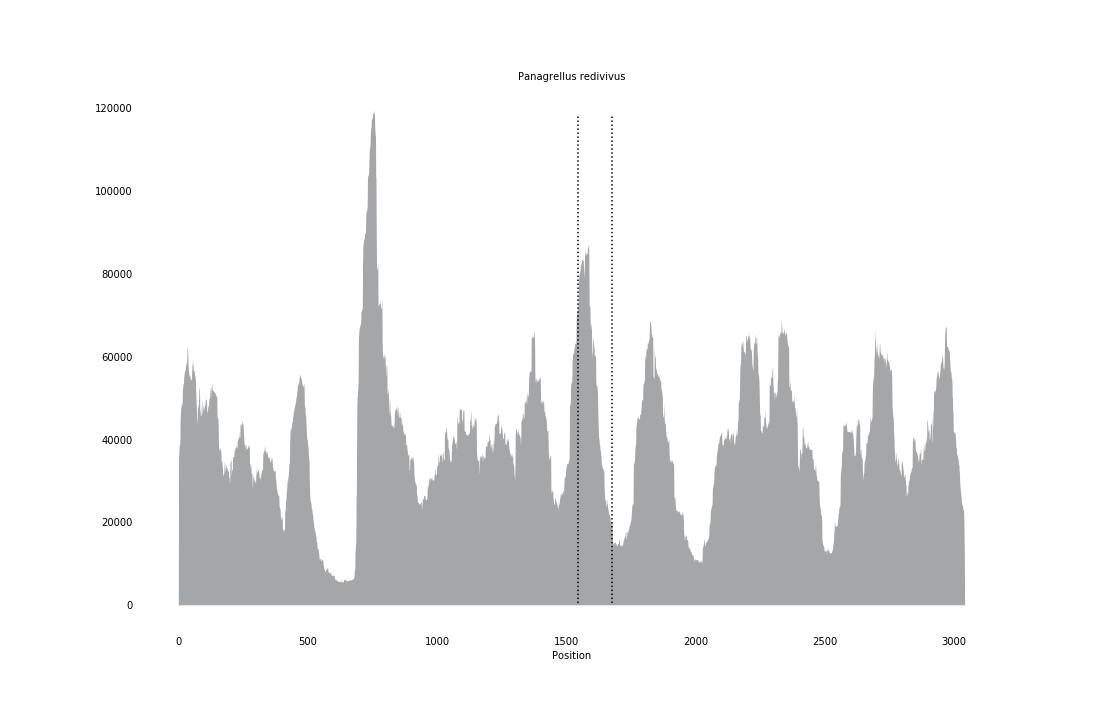

### Papio_anubis_coverage_correct.png

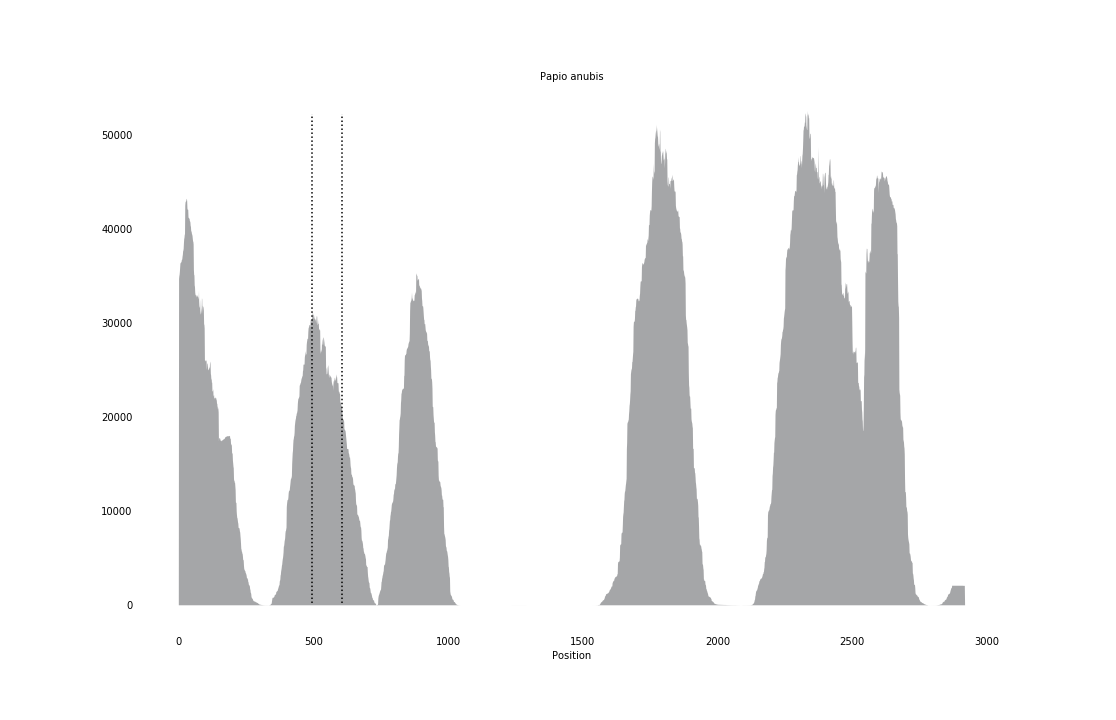

### Paracentrotus_lividus_coverage_correct.png

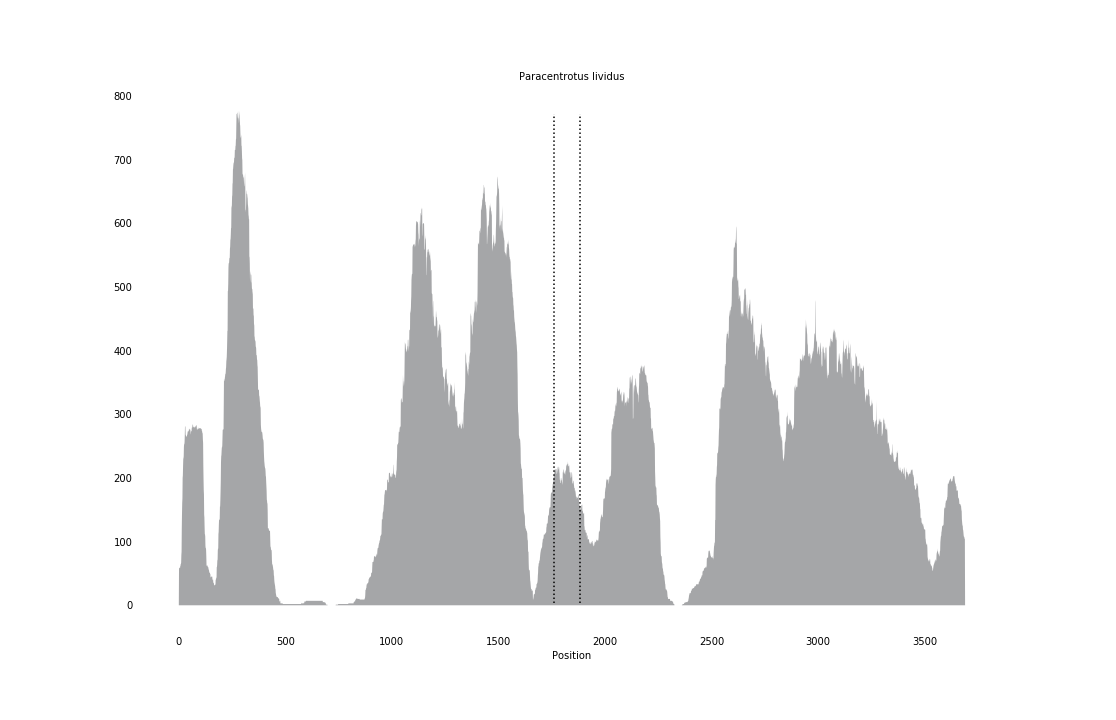

### Paragordius_varius_coverage.png

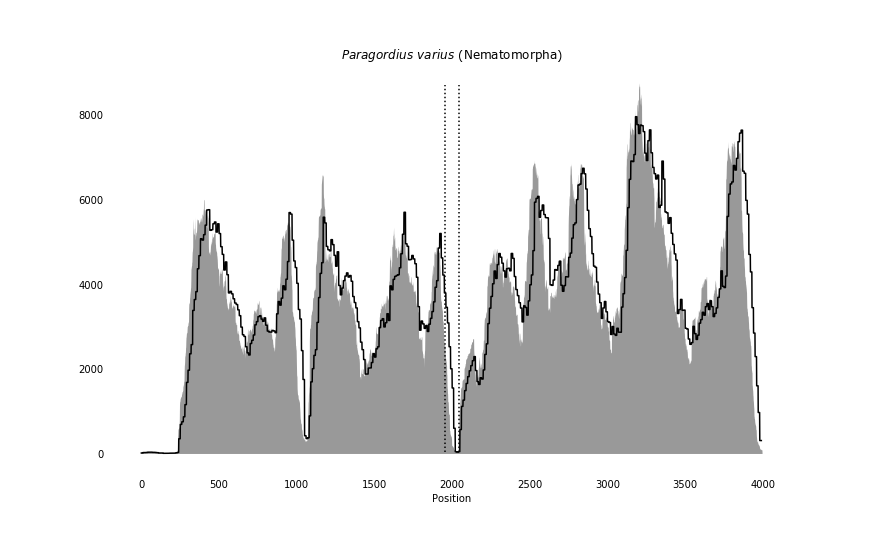

### Paratomella_rubra_coverage.png

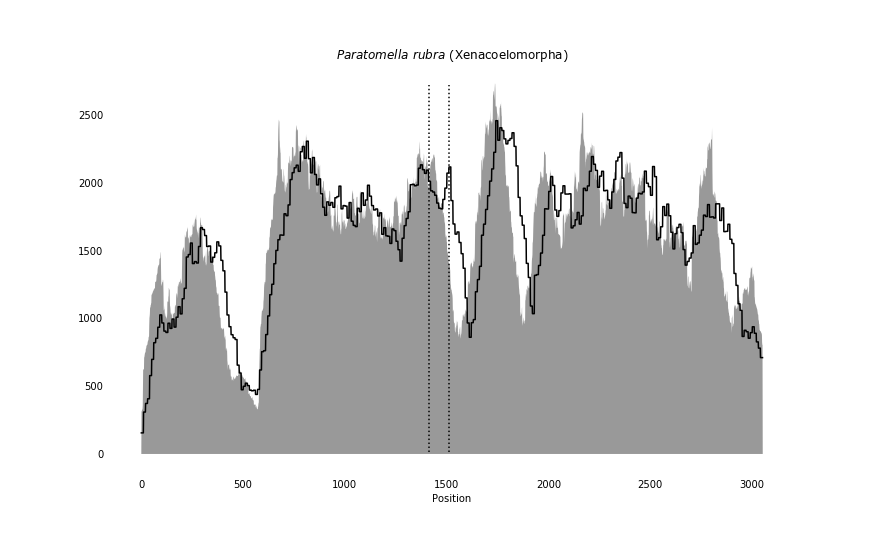

### Pelophylax_nigromaculatus_coverage_correct.png

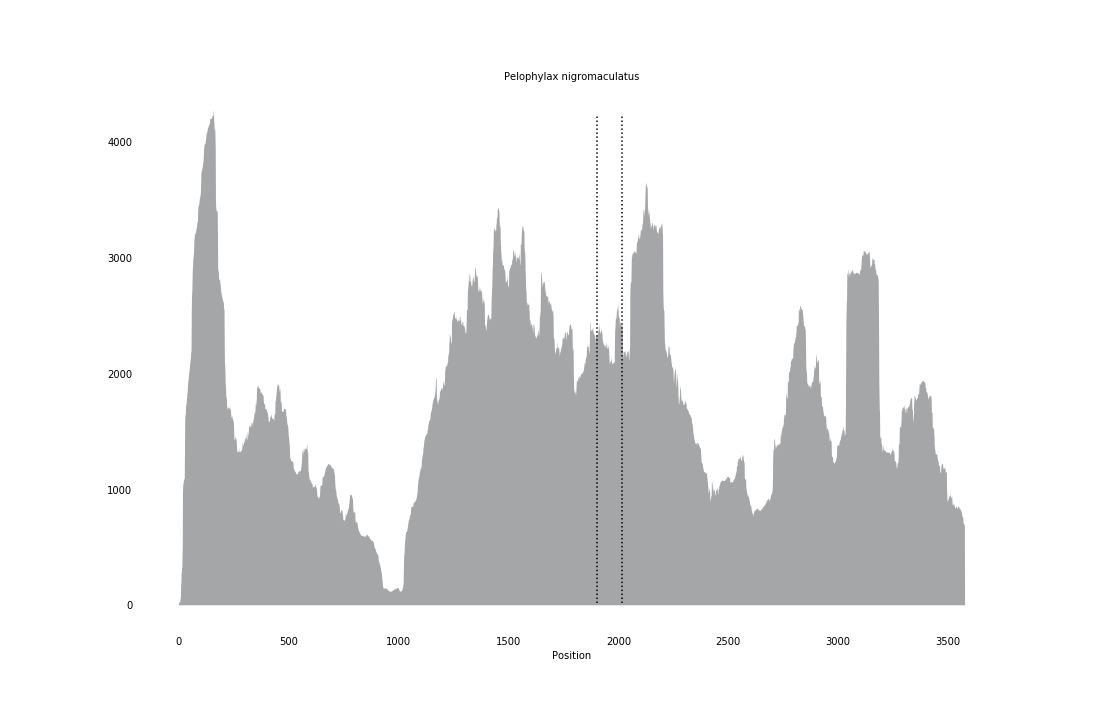

### Petromyzon_marinus_coverage.png

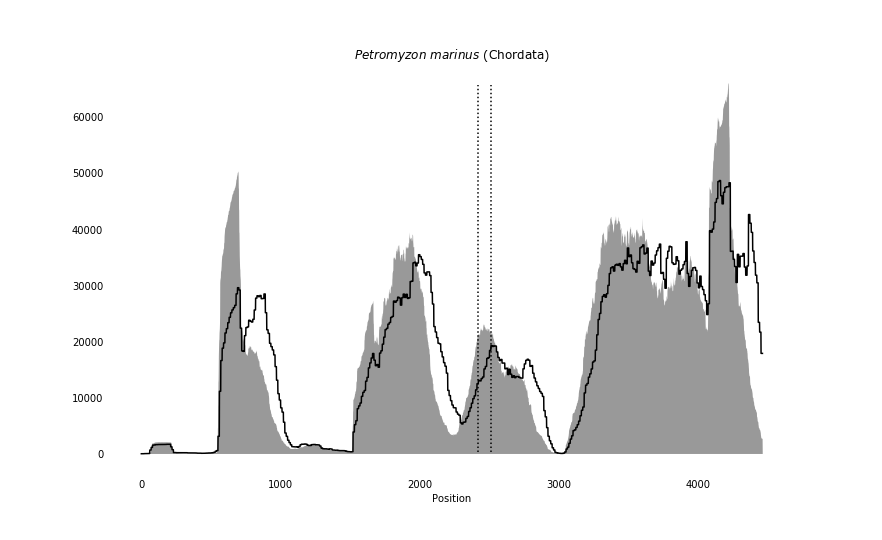

### Phascolion_cryptum_coverage_correct.png

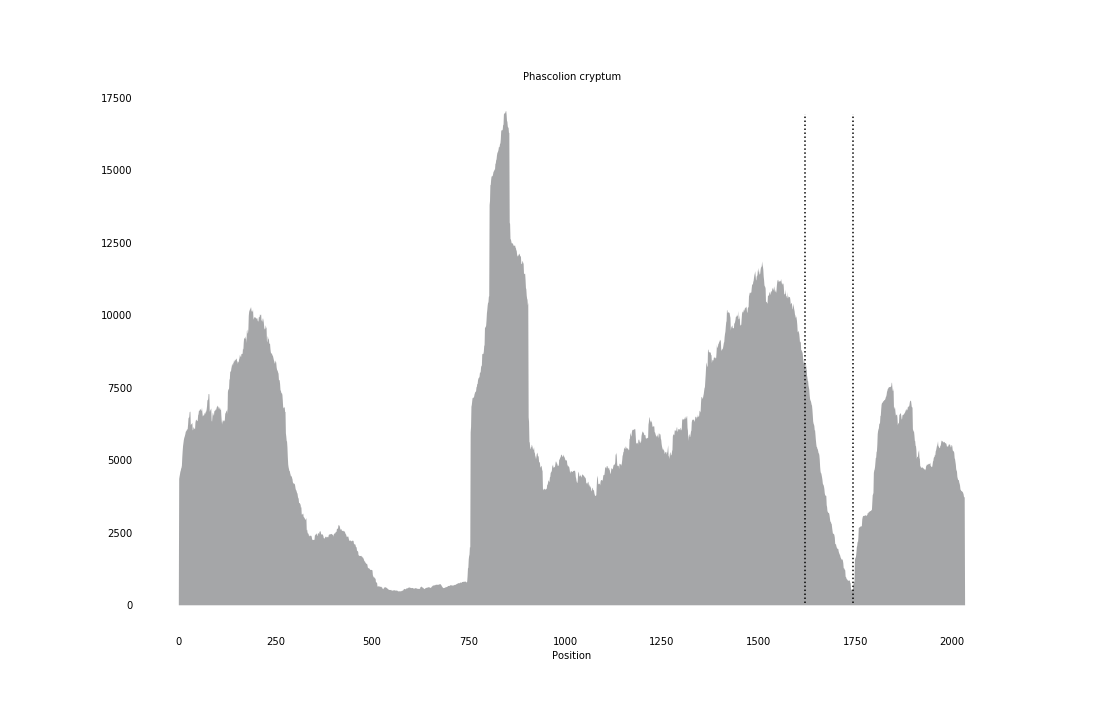

### Phodopus_campbelli_coverage_correct.png

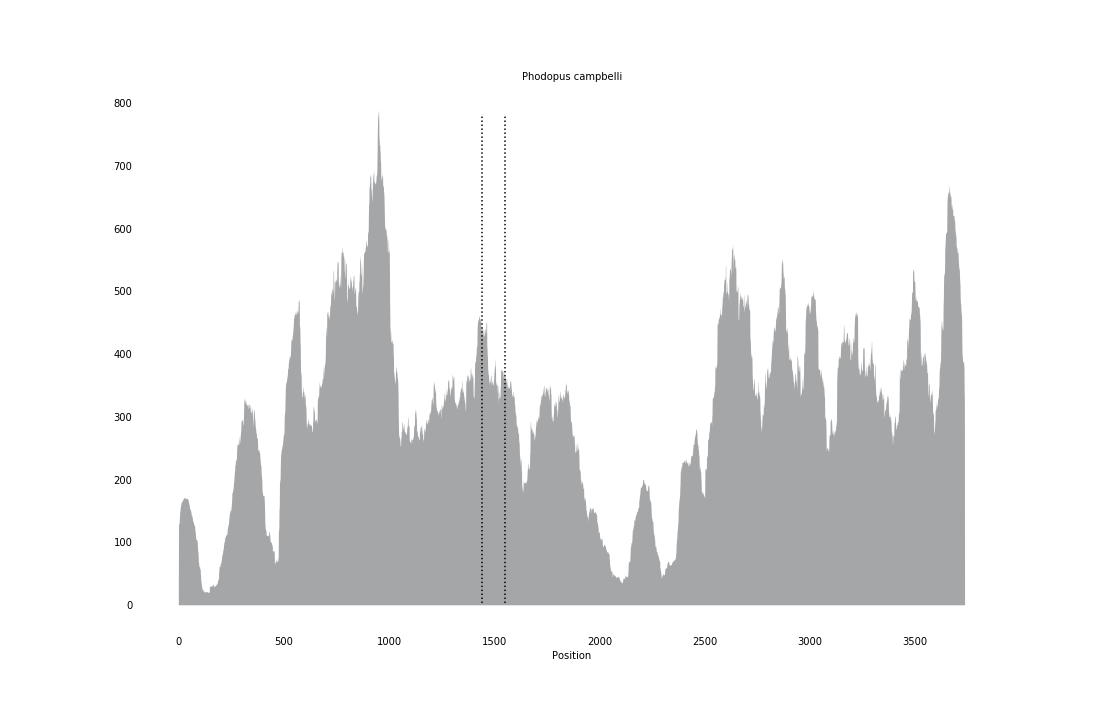

### Plectus_sambesii_coverage.png

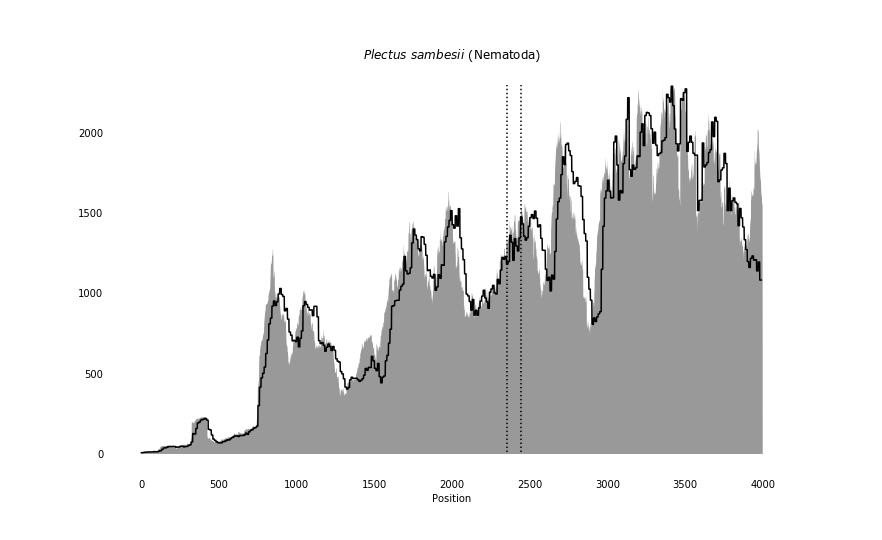

### Priapulus_caudatus_coverage.png

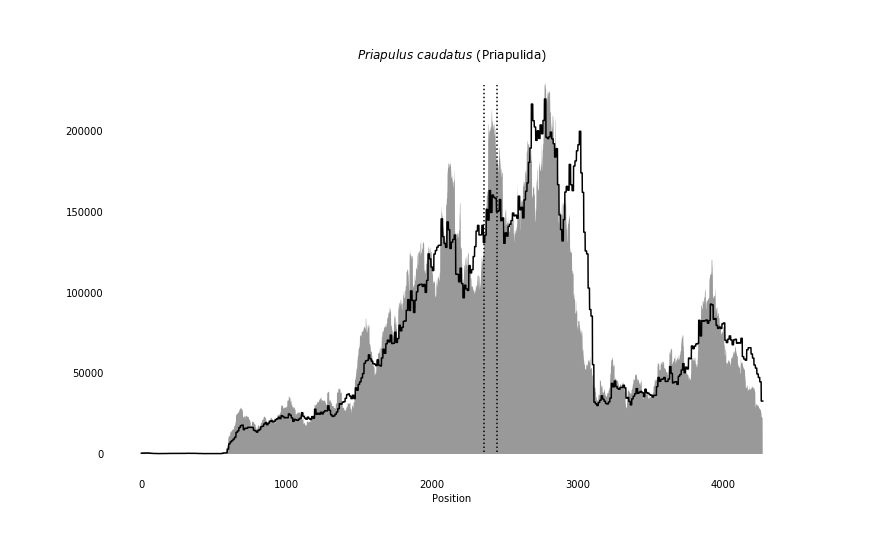

### Ptychodera_flava_coverage.png

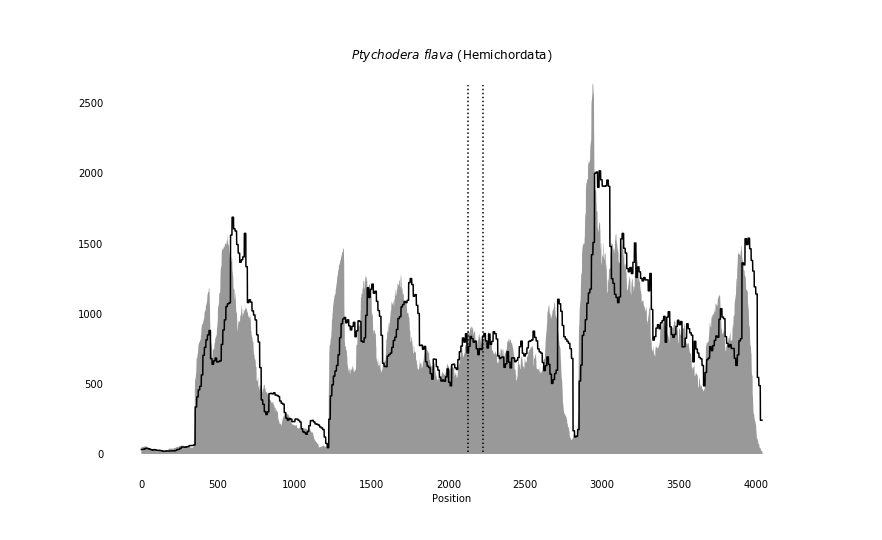

### Pycnophyes_kielensis_coverage.png

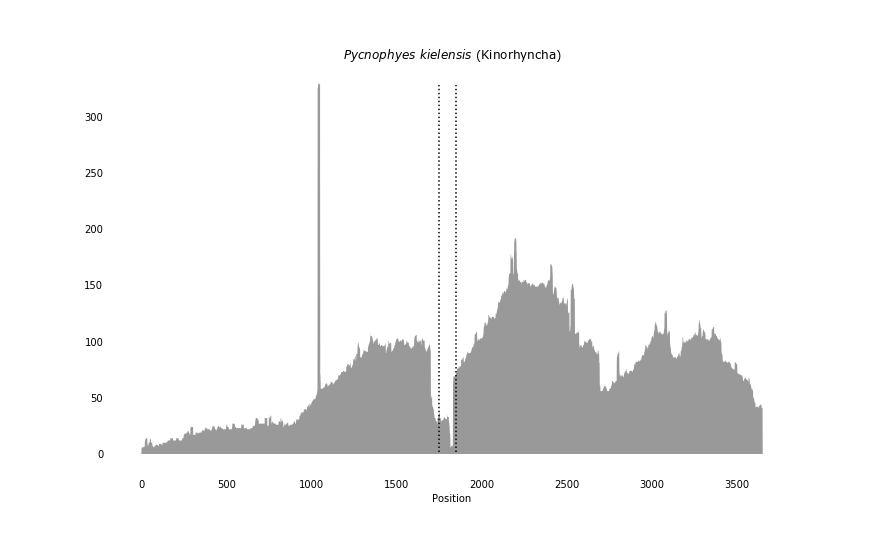

### Rhabdopleura_compacta_coverage.png

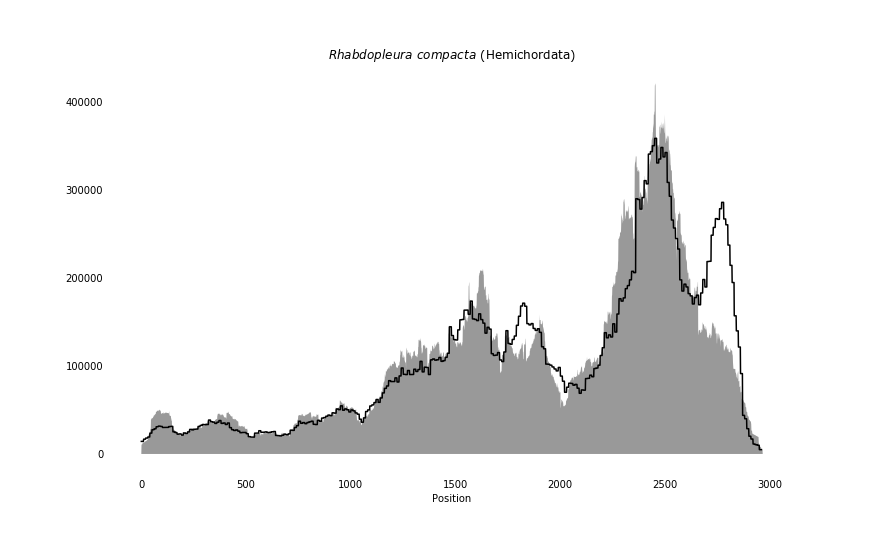

### Romanomermis_culicivorax_coverage.png

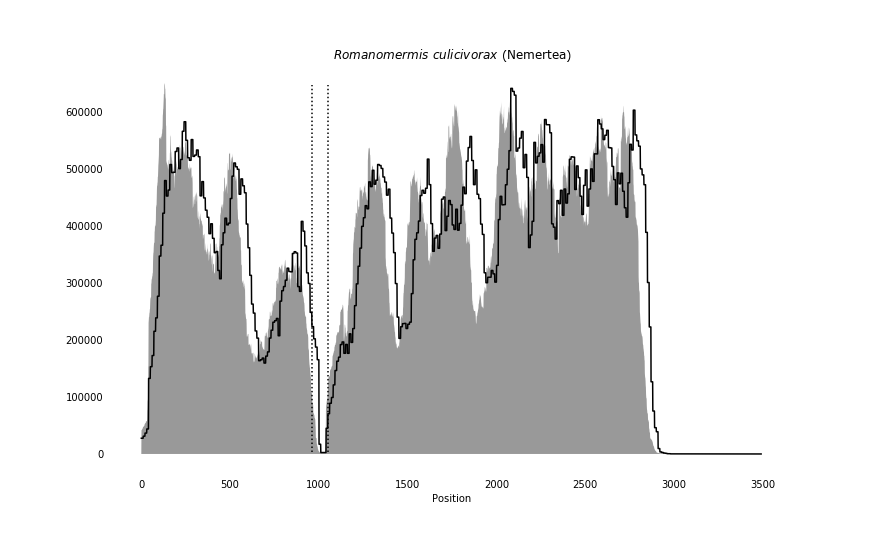

### Sagitta_elegans_coverage.png

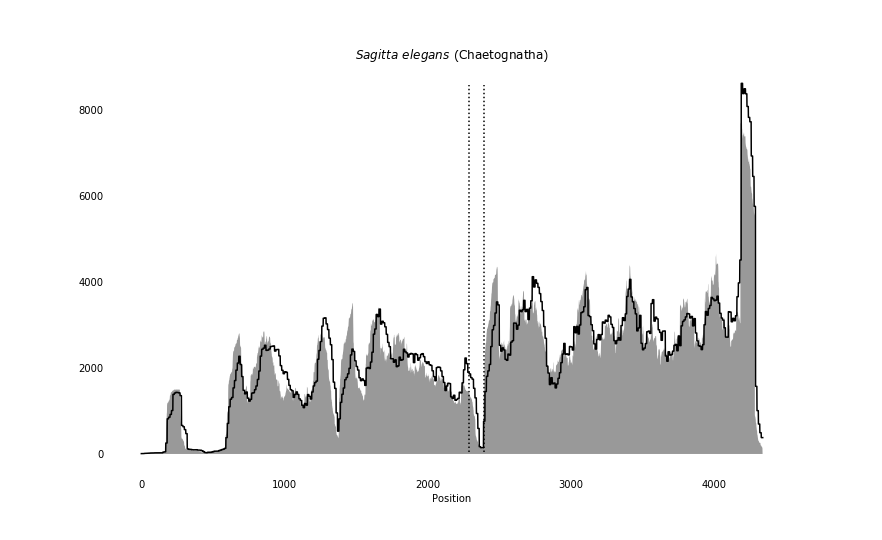

### Salpa_thompsoni_coverage_correct.png

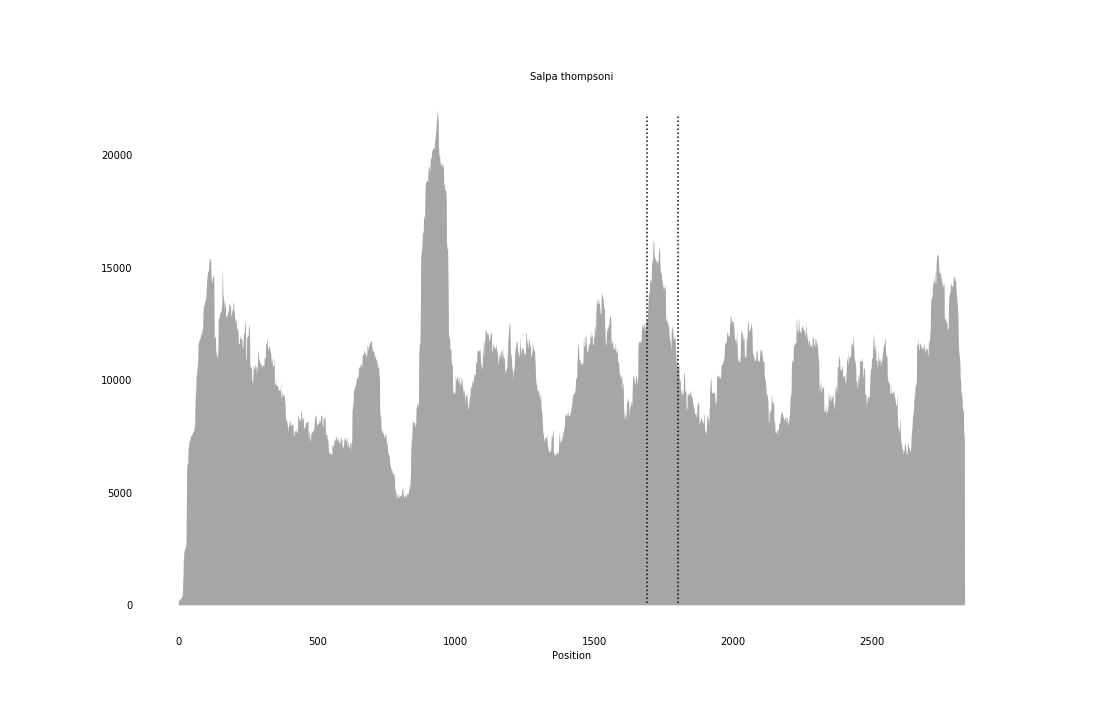

### Salvelinus_alpinus_coverage_correct.png

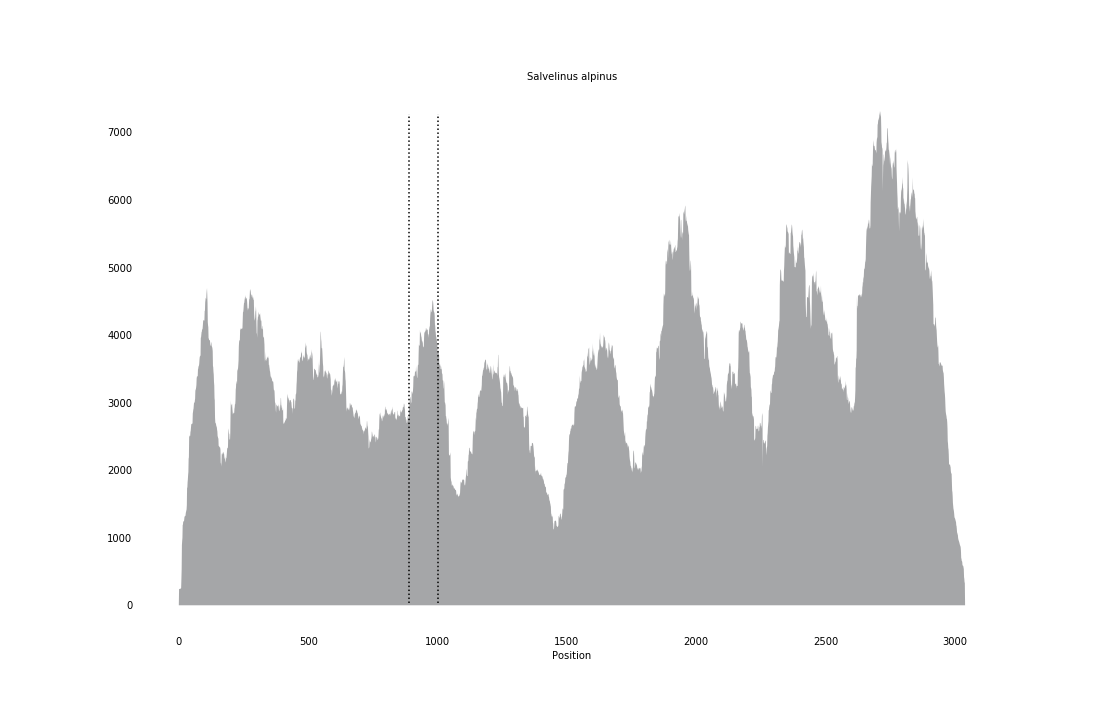

### Salvelinus_fontinalis_coverage_correct.png

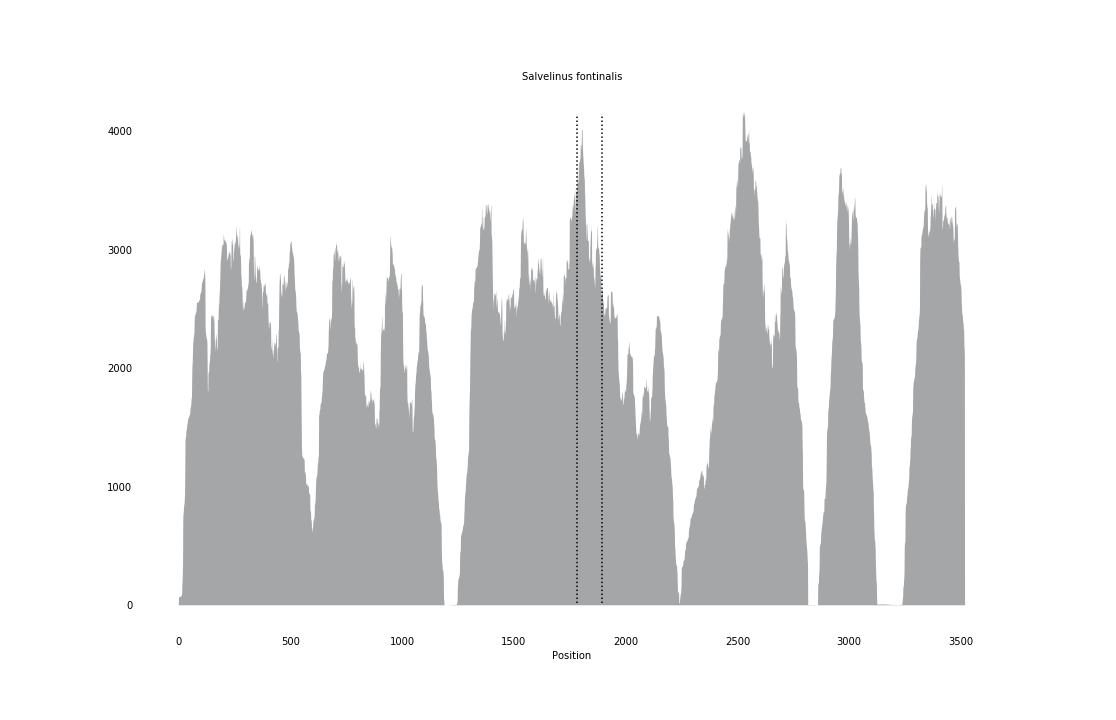

### Scyliorhinus_torazame_coverage_correct.png

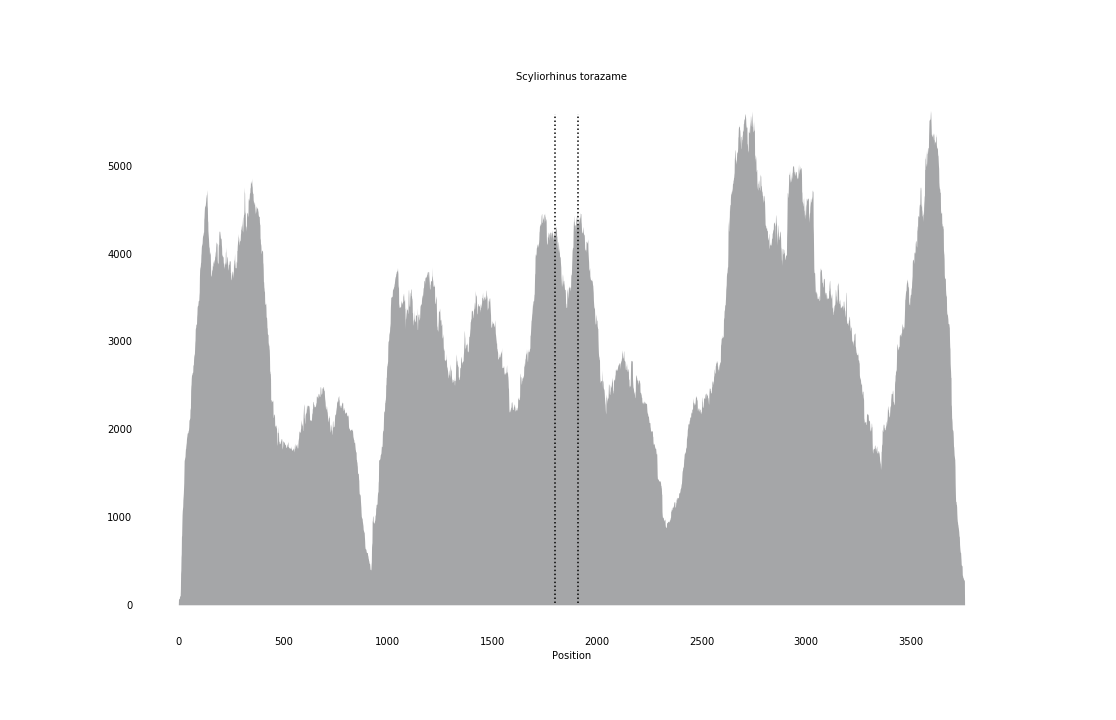

### Sebastes_caurinus_coverage_correct.png

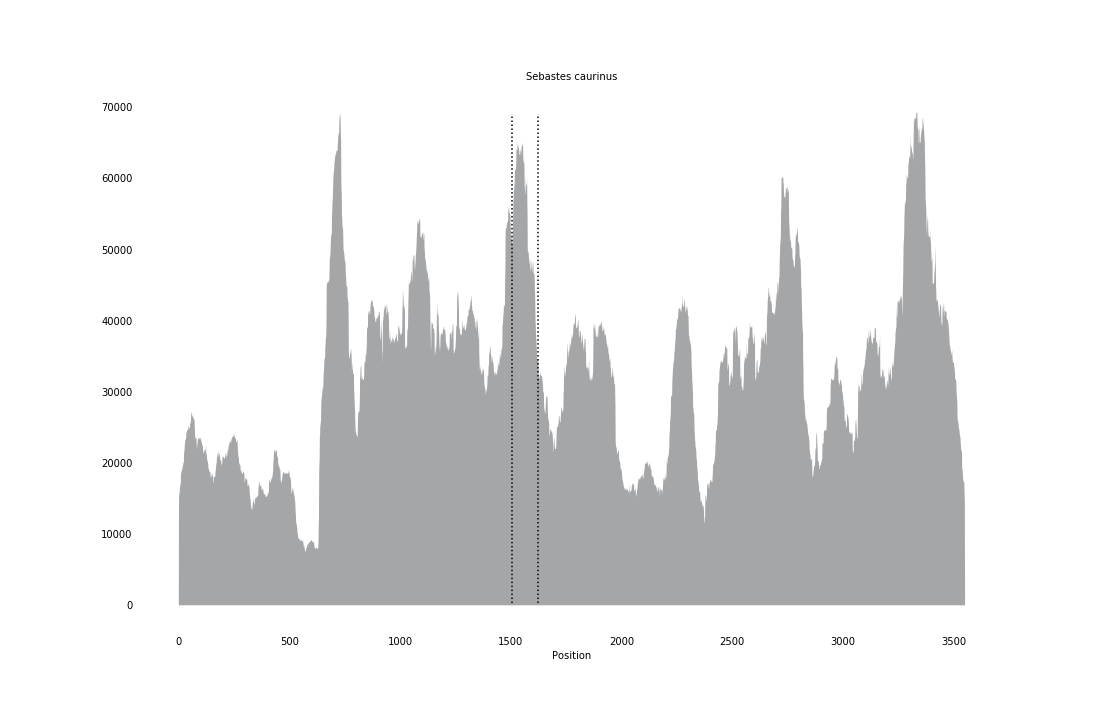

### Sipunculus_nudus_coverage_correct.png

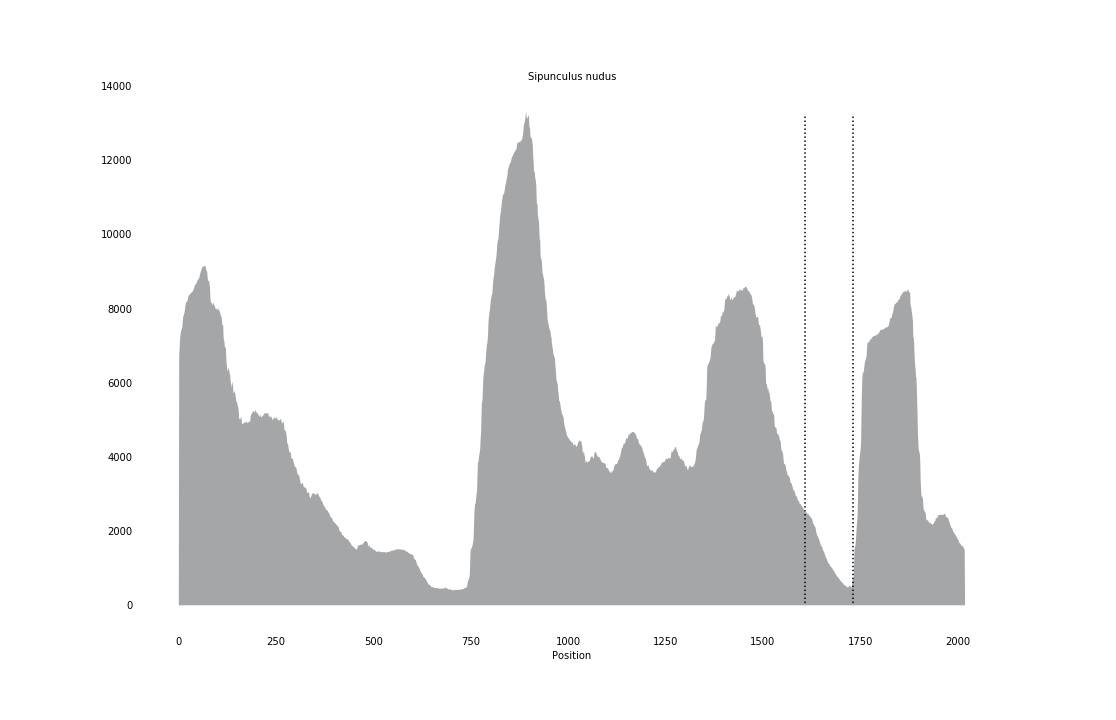

### Strongylocentrotus_purpuratus_coverage.png

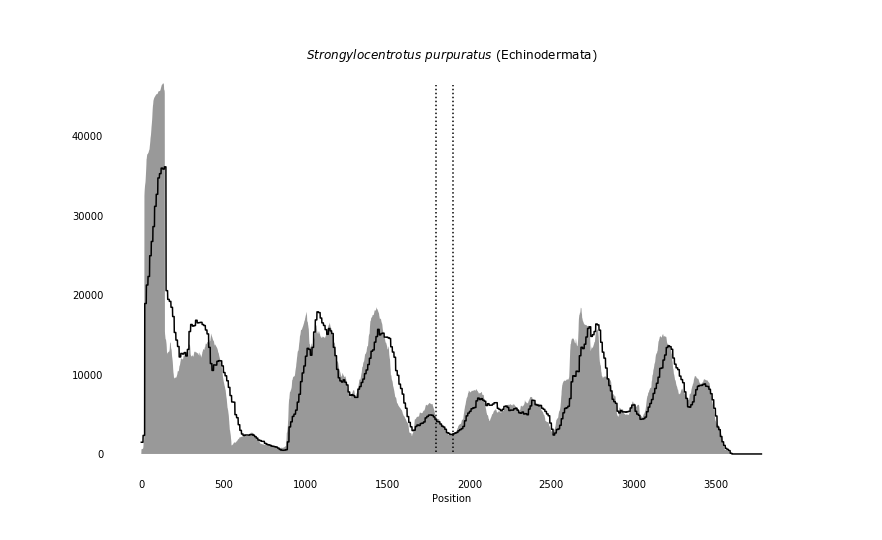

### Symbion_pandora_coverage.png

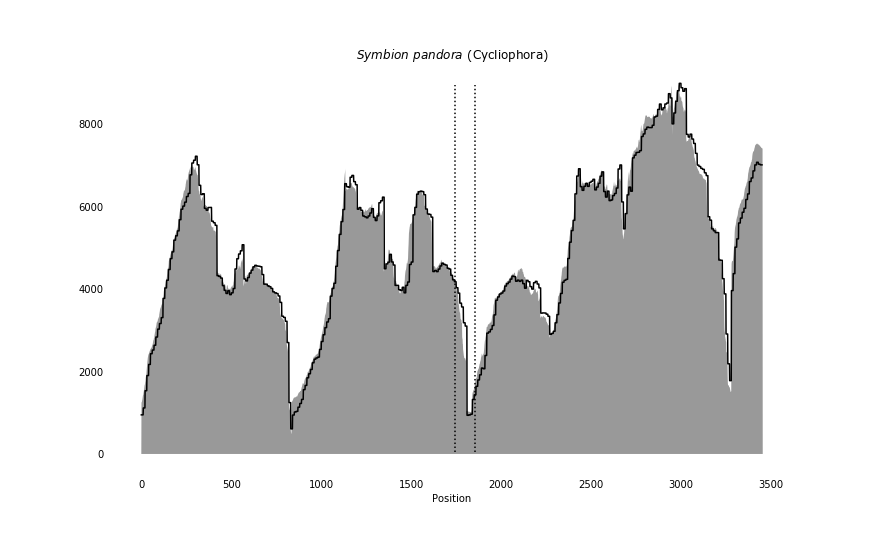

### Symsagittifera_roscoffensis_coverage_correct.png

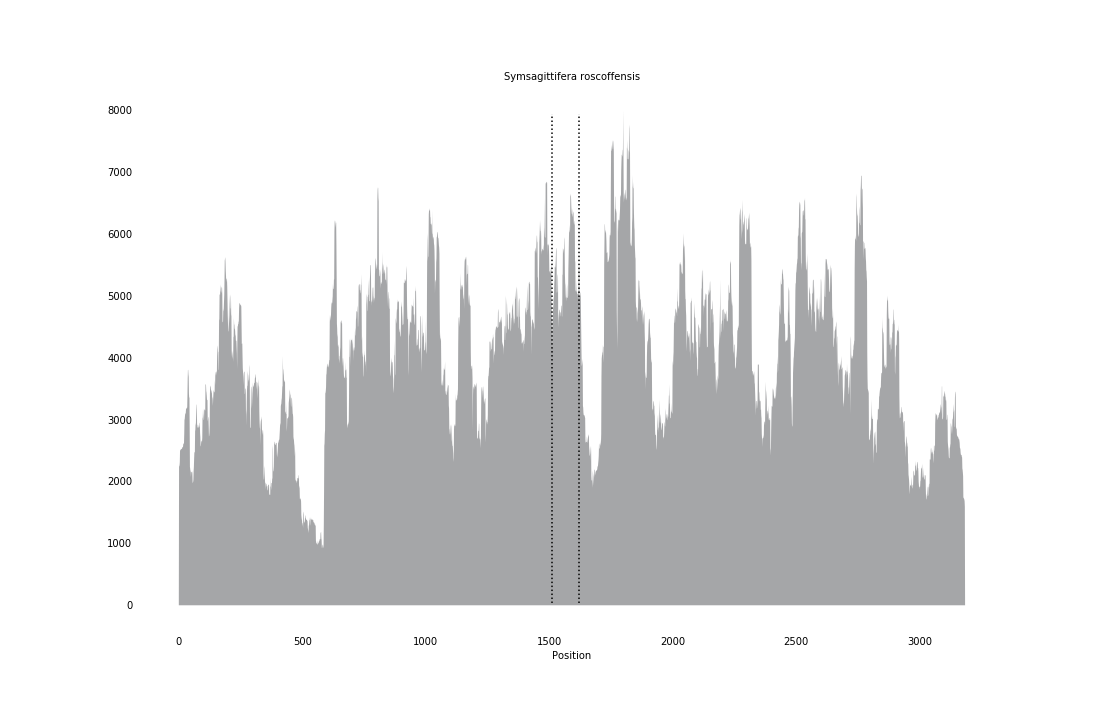

### Terebratalia_transversa_coverage.png

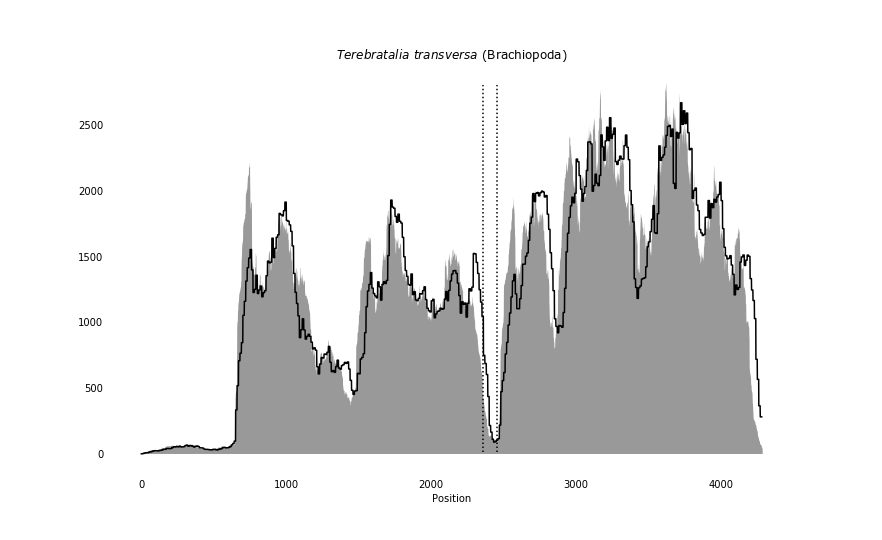

### Trichinella_zimbabwensis_coverage_correct.png

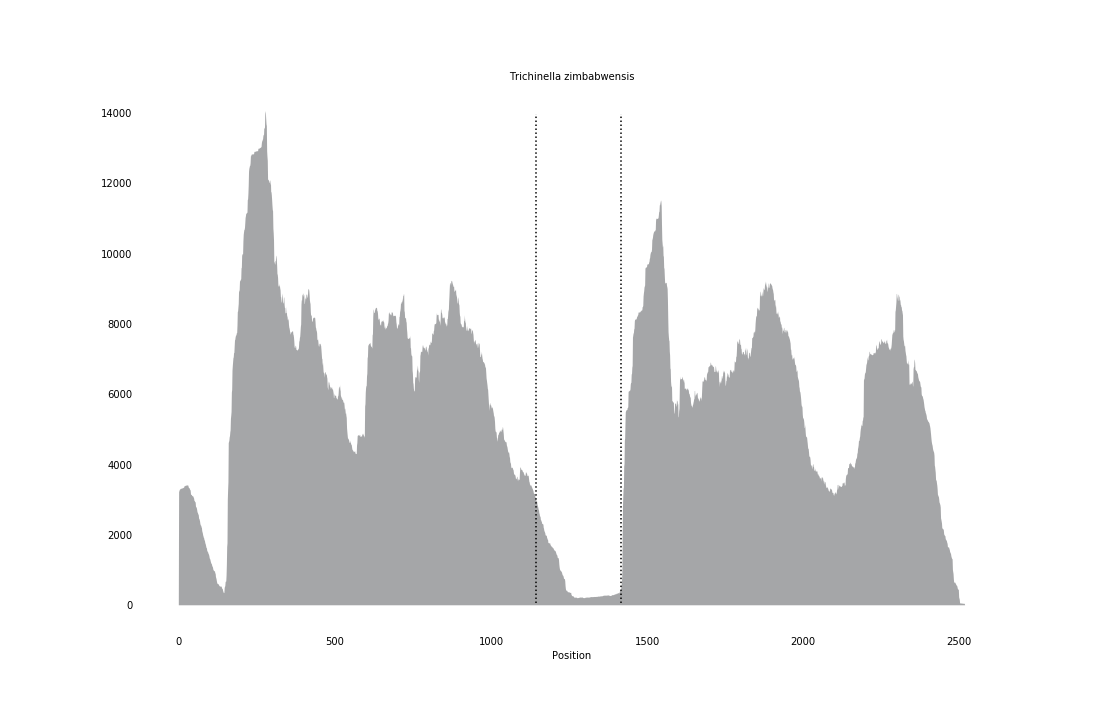

### Umbra_pygmaea_coverage_correct.png

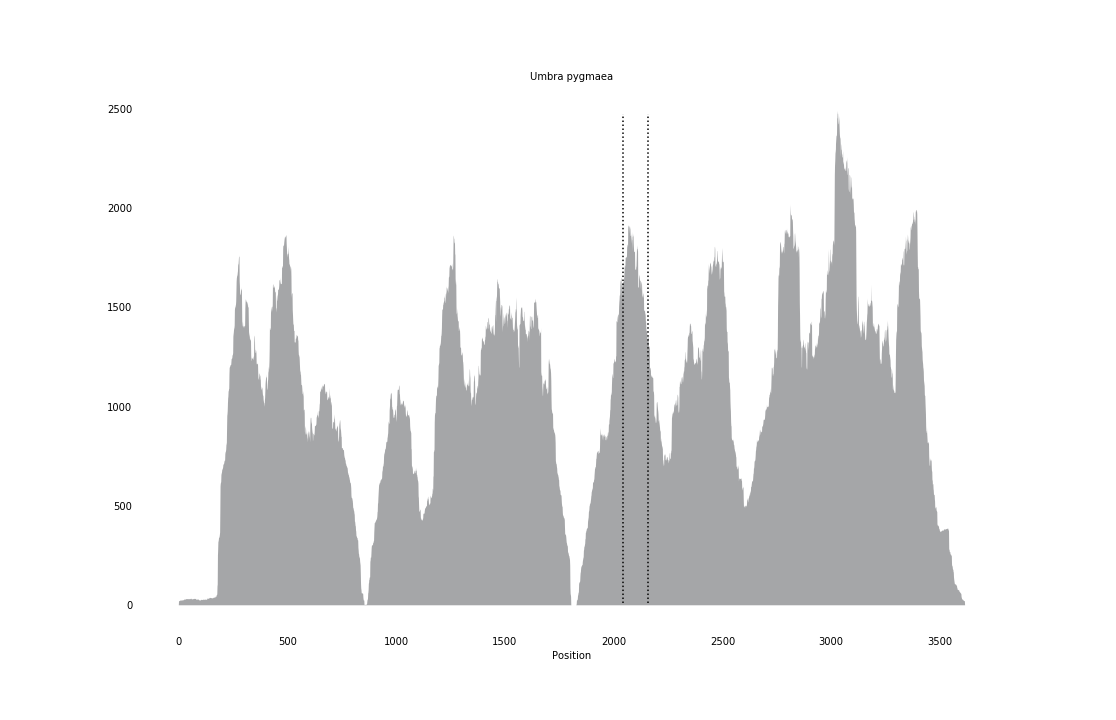

### Xenoturbella_profunda_coverage.png

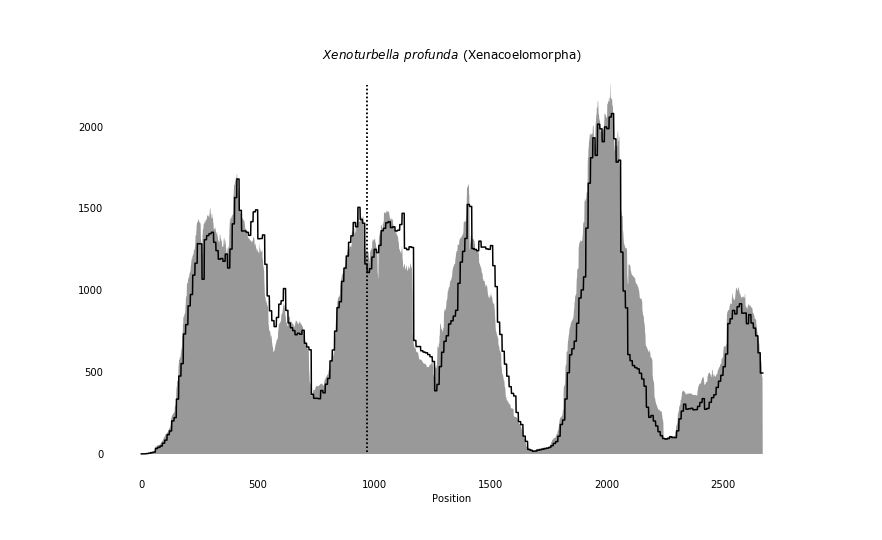
