## Supplementary figures and images for "Computational discovery of hidden breaks in 28S ribosomal RNAs across eukaryotes and consequences for RNA Integrity Numbers"

### adineta_log_fr.png

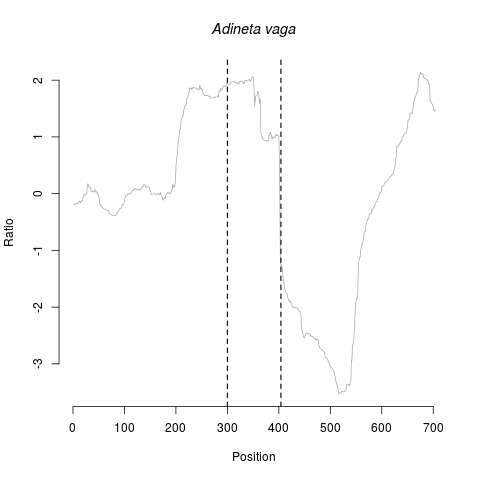

### adineta_log_rf.png

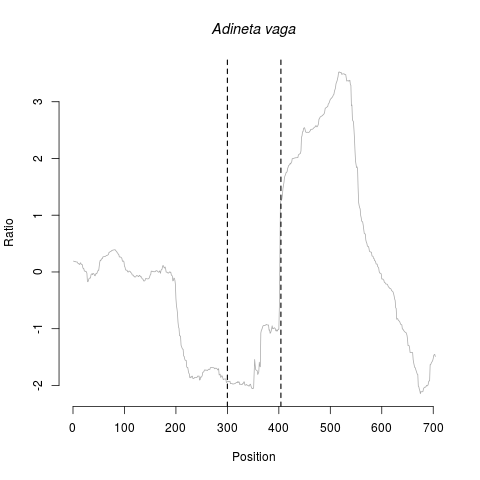

### aporometra_log_fr.png

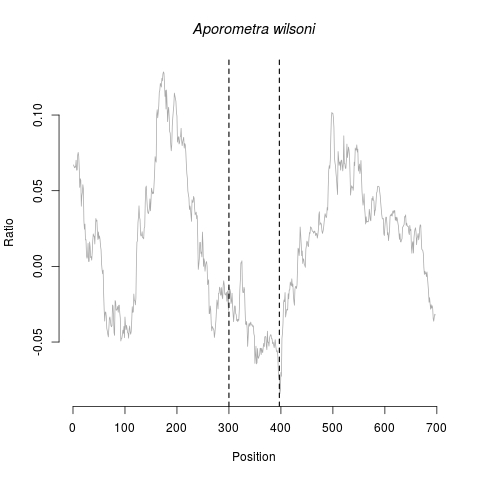

### aporometra_log_rf.png

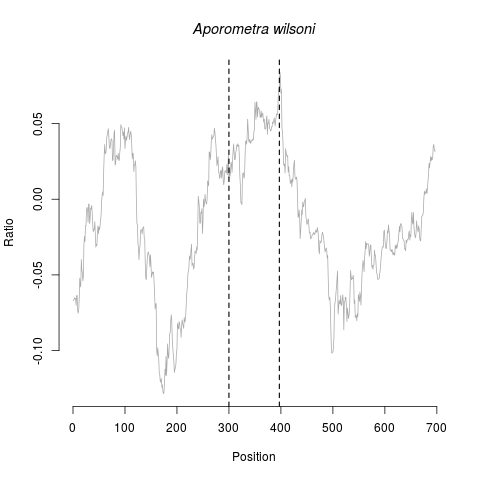

### barentsia_log_fr.png

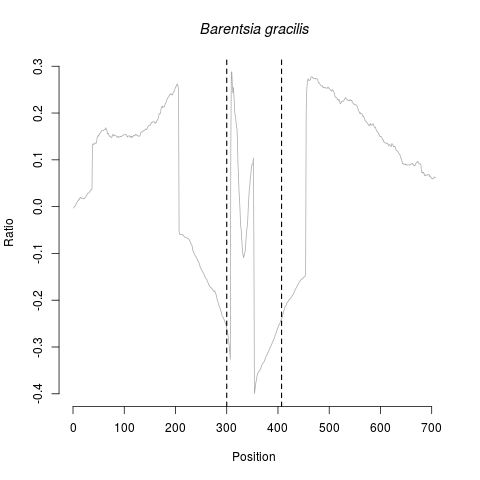

### barentsia_log_rf.png

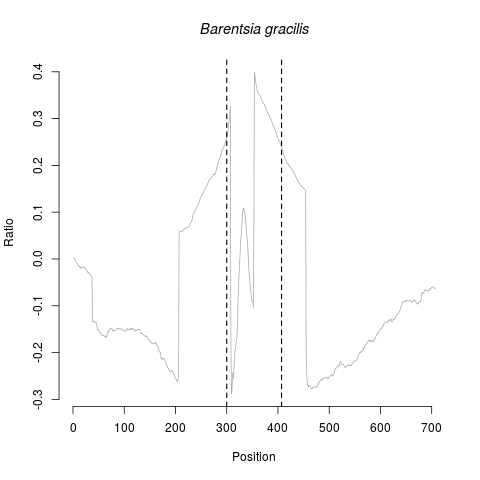

### bombyx_log_fr.png

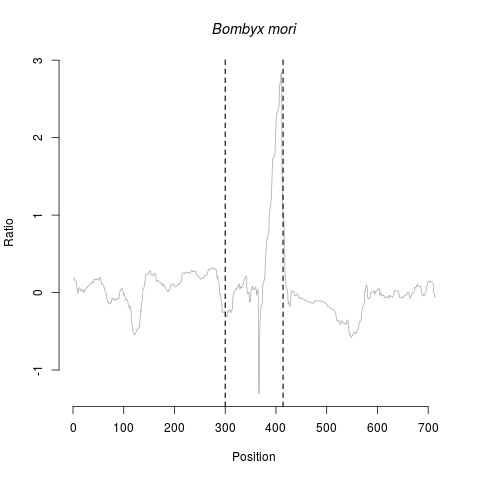

### bombyx_log_rf.png

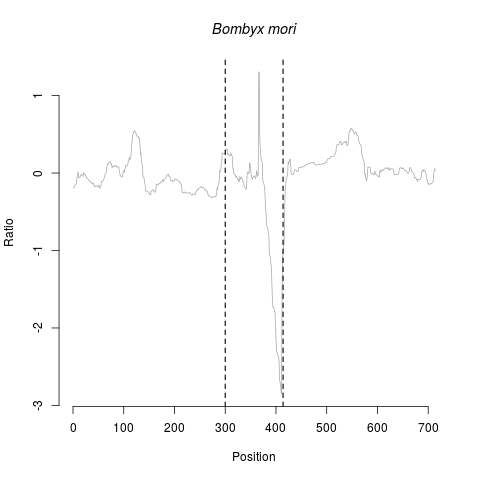

### cerebratulus_log_fr.png

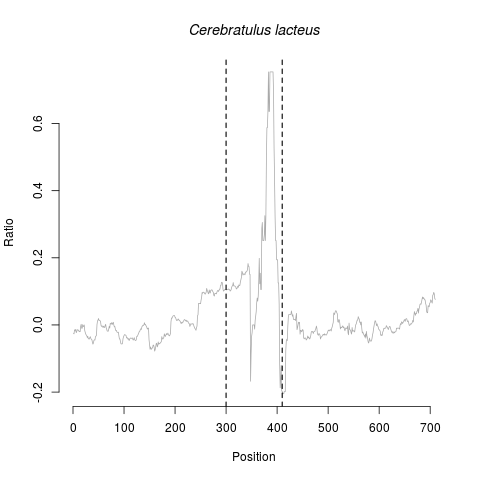

### cerebratulus_log_rf.png

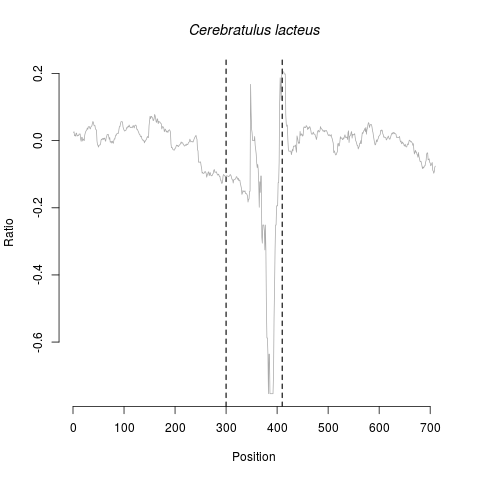

### intoshia_log_fr.png

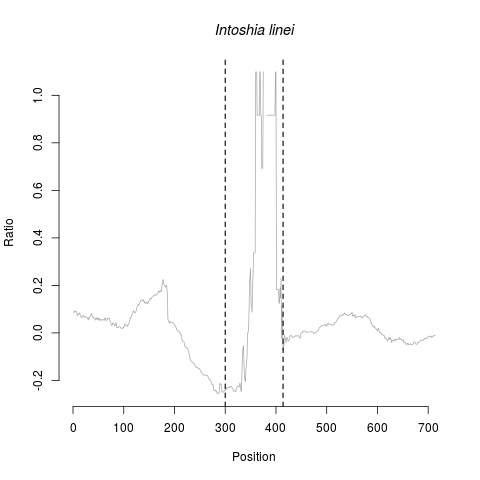

### intoshia_log_rf.png

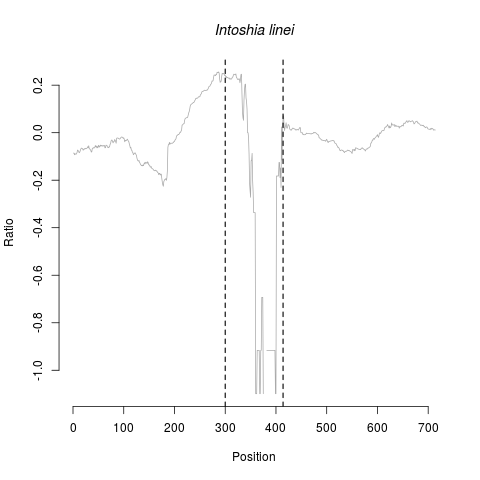

### mesodasys_log_fr.png

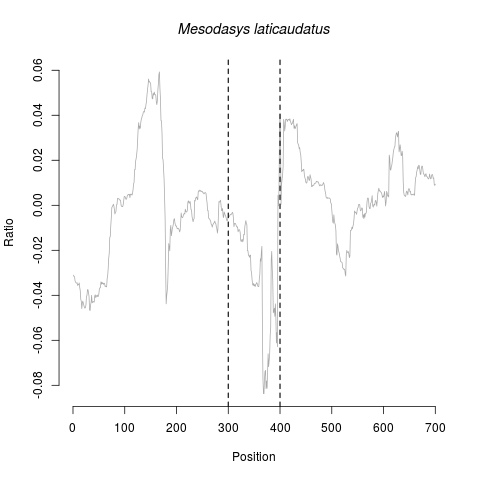

### mesodasys_log_rf.png

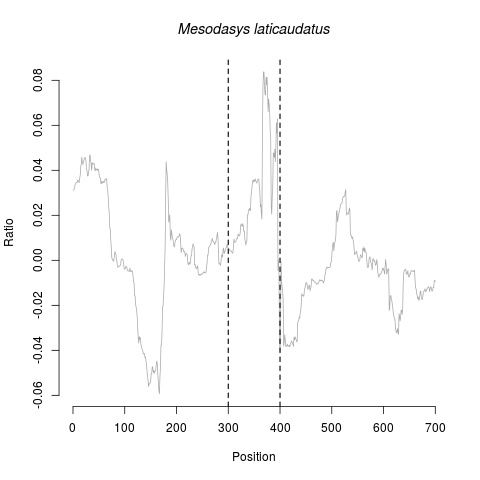

### nemertoderma_log_fr.png

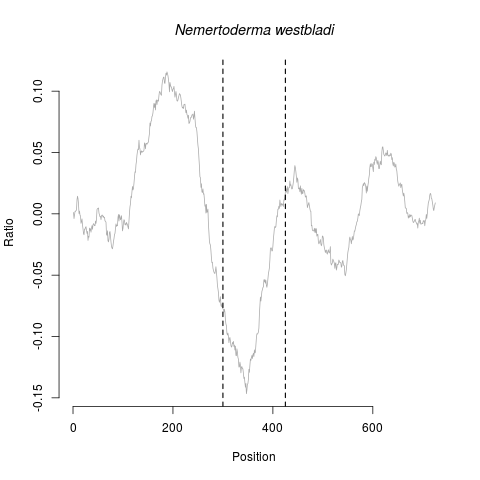

### nemertoderma_log_rf.png

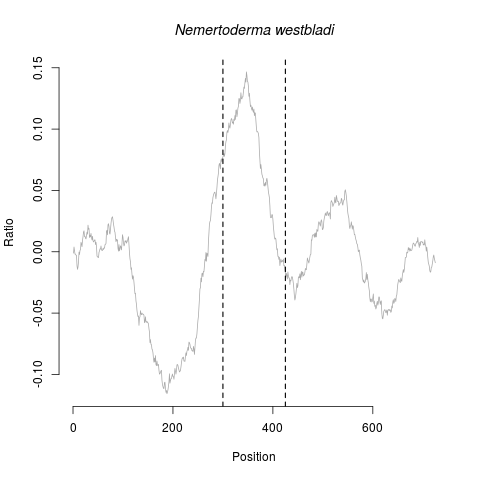

### oreochromis_log_fr.png

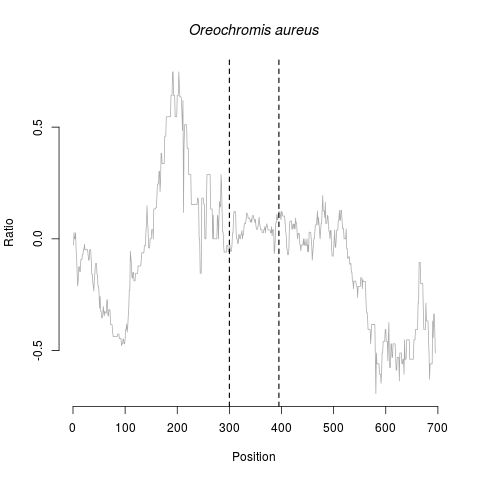

### oreochromis_log_rf.png

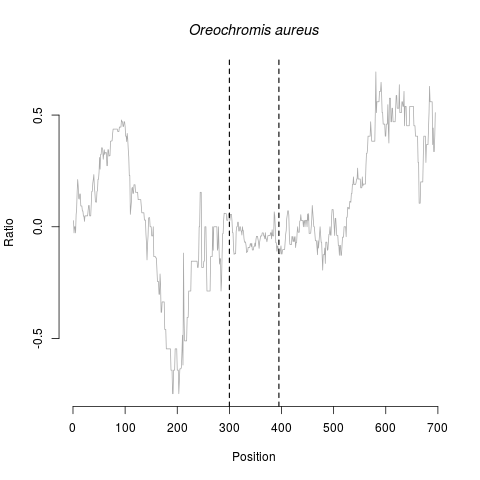

### paragordius_log_fr.png

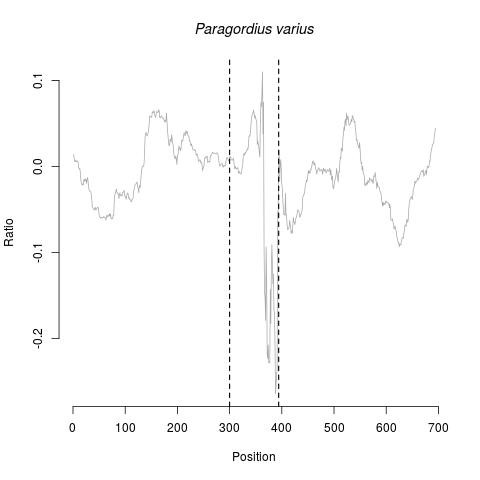

### paragordius_log_rf.png

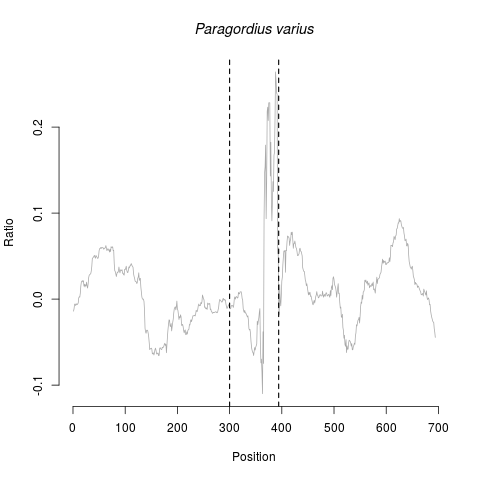

### paratomella_log_fr.png

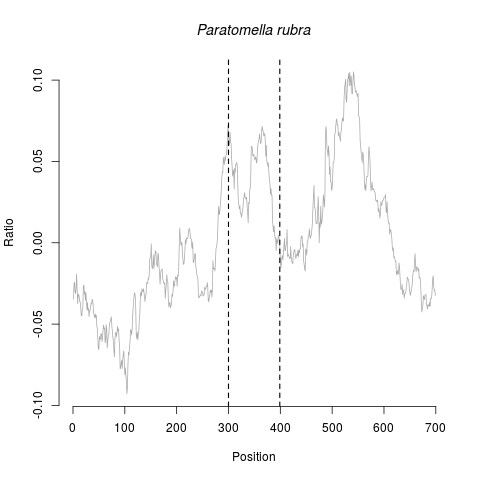

### paratomella_log_rf.png

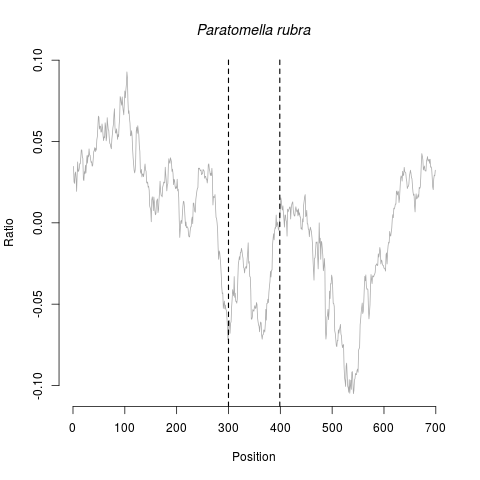

### pleurobrachia_log_fr.png

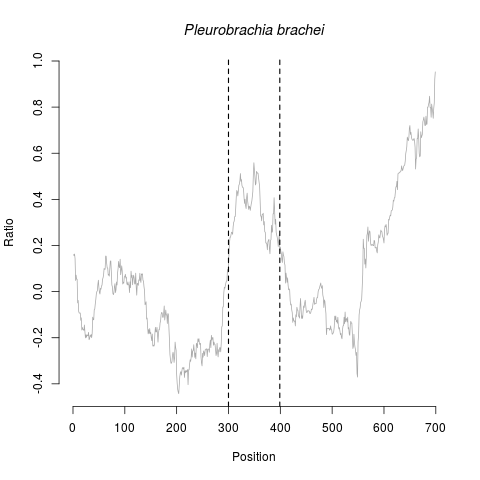

### pleurobrachia_log_rf.png

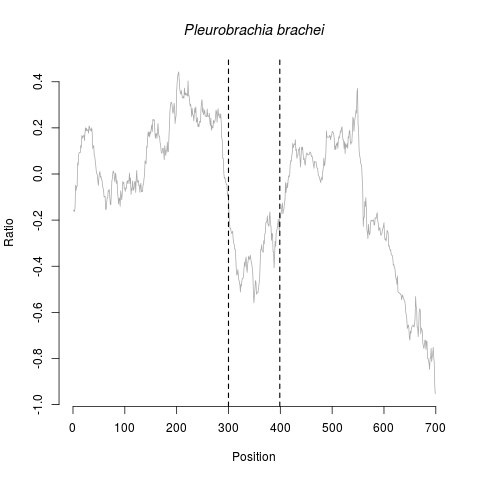

### ptychodera_log_fr.png

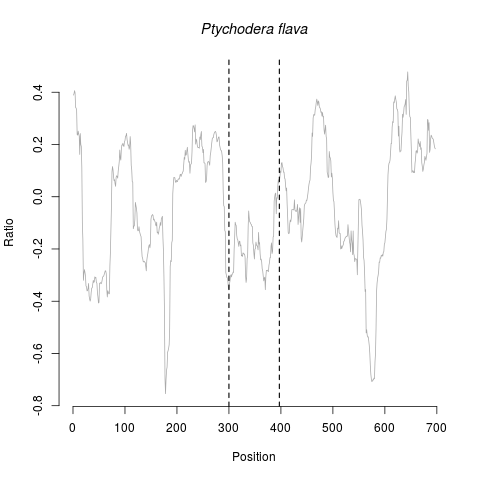

### ptychodera_log_rf.png

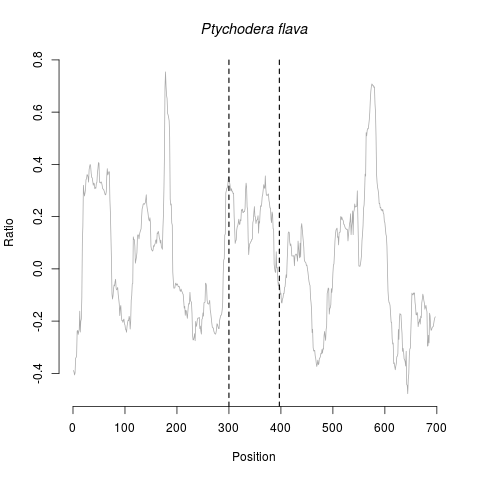

### sagitta_log_fr.png

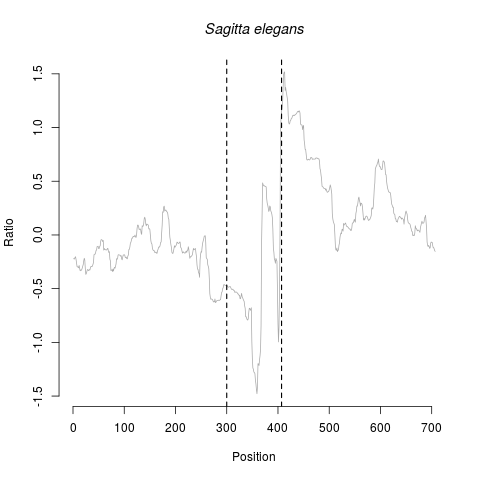

### sagitta_log_rf.png

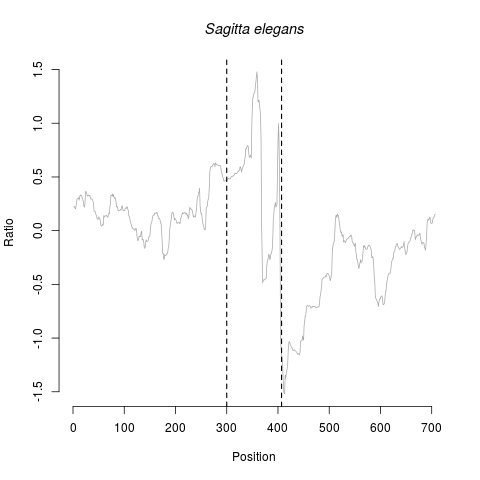

### symbion_log_fr.png

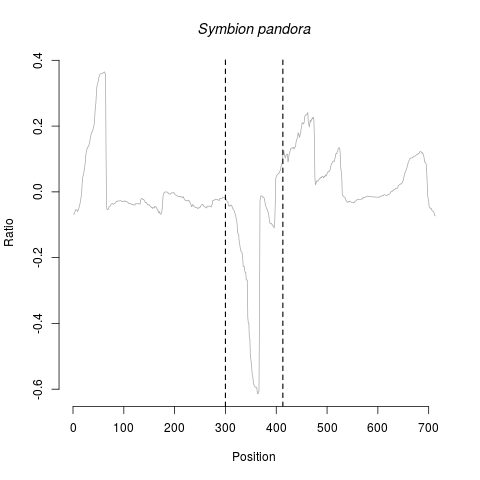

### symbion_log_rf.png

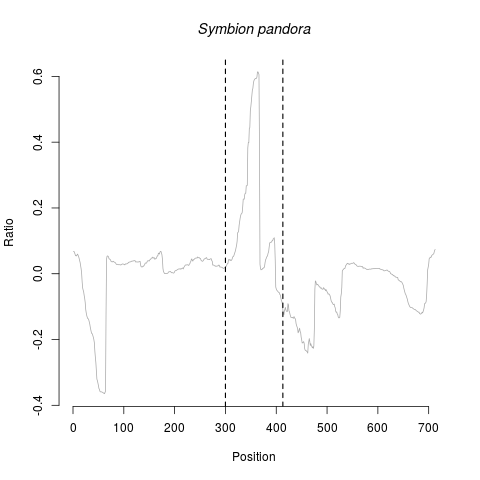

### terebratalia_log_fr.png

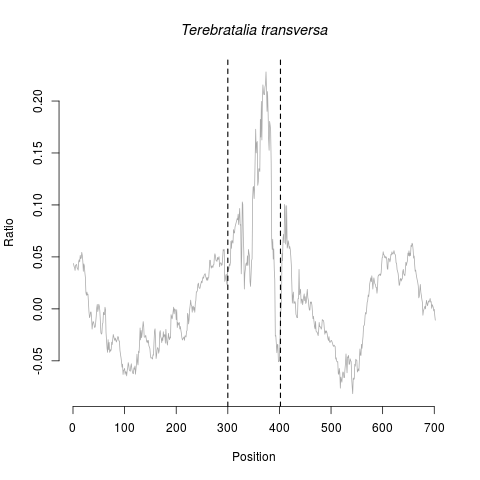

### terebratalia_log_rf.png

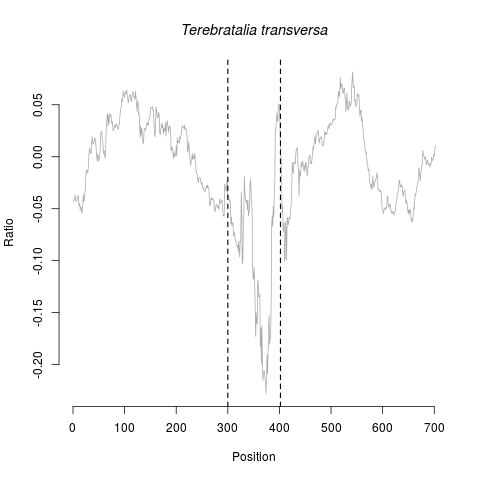

### trichoplax_log_fr.png

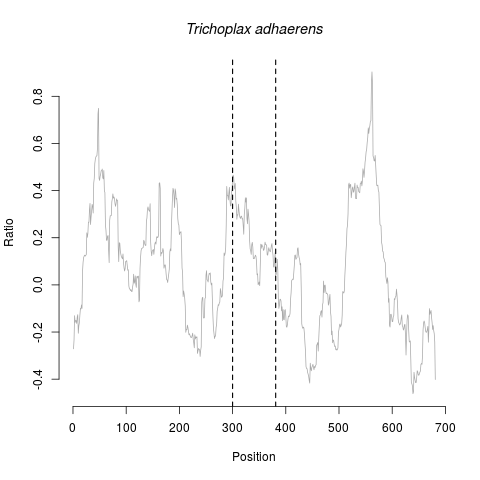

### trichoplax_log_rf.png

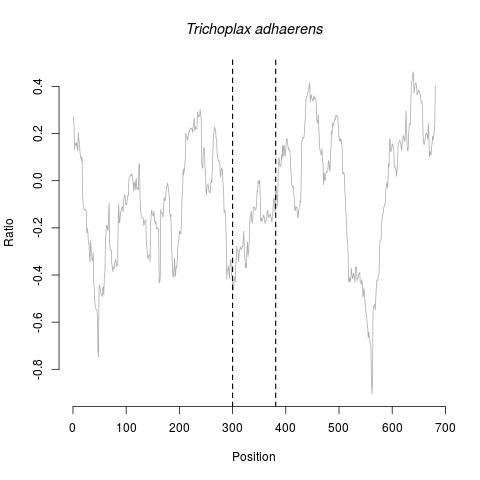
